# Gradient analysis of landscape variation in Norway

**DOI:** 10.1101/2020.06.19.161372

**Authors:** Trond Simensen, Rune Halvorsen, Lars Erikstad

**Author notes:** Abbreviations: A = land-use variable; AIV = area-independent variable; AA = area, measured as proportion of observation unit; Abygg_a = large buildings; AlpA = proportion of area above the forest line; Araaf_a = open areas; Arbar_a = coniferous forest; Arbla_a = mixed boreal forest; Arfull_a = cultivated land; Arlov_a = deciduous forest; Arover_a = surface cultivated land; Asp_n_a = north facing terrain; Asp_s_a = south facing terrain; BA = amount of boreal/alpine landscape; BE = bio-ecological gradient; Bavstn_a = sedimentary rock; BGF = basic geomorphological forms; Bohei_a = boreal heaths; Bomd_a = metamorphic rock; BP = glacier; Bplu_a = plutonic rock; Bpoor_a = lime-poor bedrock geology; BrI = mean BrI in the polygon; Brich_a = lime-rich bedrock; Build_a = built-up area; Bvul_a = volcanic rock; Ccom_m = coastal complexity; C_ek_a = simple coastline; C_kk_a = complex coastline; City_a = town/city area; CLG = complex landscape gradient; COU = count variables; Cr3_r_a = smooth/flat coast; Cr3_u_a = rugged coast; Cre_b_a = steep coast; Cre_f_a = flat coast; Crug3_m = coastal ruggedness, VRM3, mean; Crug9_m = coastal ruggedness, coarse scale, VRM9, mean; Cul_a_s = number of archaeological heritage sites; Cul_b_s = number of ancient rock art sites; Cul_k_s = number of church ruins; Cul_m_s = number of marine cultural heritage sites; Cul_t_s = number of technical heritage sites; Cul_u_s = number of cultural heritage sites outdoors; DCA = detrended correspondence analysis; Discoast = mean distance to coast; Dislake = mean distance to lake; Dismire = mean distance to mire; DN = valley form; DNI = relationship between valley depth and valley width; EDU = ecodiversity distance units; EDU-L = ecodiversity distance units in landscapes; Ekspbe_a = slightly protected coast; Ekspmo_a = slightly exposed coast; Ekspve_a = exposed coast; Er_m = hypsographic index, ER, mean; FlA = amount of flat land; Flat_a = flat terrain; G = basic geo-ecological variables; GE = geo-ecological gradient; Gab_a = built-up; Gab_fi = fisheries-related buildings; Gab_nae = commercial buildings; Glac_a = glacier; GNMDS = global nonmetric multidimensional scaling; Guro_t_a = rugged terrain; H.C. units = half-change units; I = inland landscapes; IA = inland hills and mountains; INA = internal association; ID = inland valleys; IF = fine sediment plains; IfI = mean infrastructure index in the polygon; not used inter alia because of error in terrestrial area correction; IfIu = same as IfI, but calculated by the formula for IfIutv in Erikstad et al. (2014); II = mean lake index in the polygon; Innoy_s = freshwater lake islands; Inns_s = number of lakes; IP = freshwater lake properties; IX = inland plains without dominance of fine sediments; JI = mean agricultural index in the polygon; JK1 = primary soil/ sediment class; JK1–B = exposed bedrock (> 50% of area covered by exposed bedrock); JK1–E = glaciofluvial deposits; JK1–H = marine deposits; JK1–M = thick layer of till; JK1–0 = no soil/sediment class (> 50% not assigned to specific sediment/soil type); JK2 = secondary soil/ sediment class; JK2–B = exposed bedrock (> 25% of area covered by exposed bedrock); JK2–E = glaciofluvial deposits; JK2–EH = marine deposits; JK2–EM = thick layer of till; JK2–E0 = no soil/sediment class (> 25% not assigned to specific sediment/soil type); JP = amount of agriculture; K = coastal landscapes; Kbf_a = exposed bedrock; Kelv_a = glaciofluvial deposits; KF = fjords and other coastal landscape; KL = coast line; Klac_a = lacustrine deposits; Kmar_a = marine deposits; KS = coastal plains; Kskred_a = landslide soil; Ktkmo_a = thick layer of till; KV = groups of characterising variables; collective term for primary key and analysis variables; Lake_a = freshwater lake; Land_a = terrestrial area; LEP = landscape property profiles; LGL = total landscape gradient length; LG-variable = landscape gradient variable; LEC = local complex environmental variable; Lled_a = power lines; LNr = serial number; LT = landscape type; M = marine landscapes; Maro_s = marine islands; Meant_a = moderate slope; MI = mean mire index in the polygon; Mire_a = mire; MP = abundance of mire; NiN = Nature in Norway; OI = amount of infrastructure; ONV = orthogonal key variable; OU = Observation unit; Oyst_i = inverse island size; PCA = principal component analysis; PD = proportional dissimilarity; PKV-group = parsimonious KV-group, see KV; PLC = primary landscape major-type candidate; PNV = primary key variable; pONV’s = the primary orthogonal key variable (see ONV) values ranged on a 0-1 scale; PRA = proportion of area (pixel number variable); R_net_a = river; Rd_anl_a = reindeer husbandry facilities; RDA = redundancy analysis; RE = relief; River_a = large river; rONV = rescaled ONV (see ONV); RR1 = mean RR1 in the polygon given in meters; RR1_m = altitudinal range; rrONV = ranged, rescaled ONV (see ONV); Rug3_m = terrain ruggedness VRM3, mean; SD = standard deviation; S.D. units = standard deviation units; Sefr_a = old buildings; Setr_s = number of summer mountain pasture; SK1–S = sedimentary plains on sorted sediments; SK1–SF = fine-grained sedimentary plains (marine sediments [often characterised by agricultural land use]); SK1–SG = glaciofluvial plains (gravel/clay deposits [often with forests]); SK1–U = plains without or with small proportions of sorted sediments, e.g., coarse, thick layers of till; SK1–UU = wide valley bottoms with ‘esker’, i.e. sinuous or otherwise winding ridges of material deposited by a meandering subglacial stream, etc. (10-50% sorted sediments); SK1–UA = other plains (< 10% sorted sediments); including glaciers, areas dominated by exposed bedrock, etc.; SN = archipelago properties, new; Sn_flekk = patchy open treeless area; Sn_frisk = moist/fresh heath/open areas; Sn_imp = impediment; Sn_lav = lichen heath; Sn_torr = dry heath/open areas; Sn_ureg = unregistered heath/open areas; SP = archipelago properties; SS = sediment sorting; Steep_a = steep terrain/slope; Sti_a = trail, path; Stroml_a = weak ocean current; Stromn_a = normal ocean current; Stroms_a = strong ocean current; Svcat = statistical variable category; TI = total inertia; TP = peaks; TPI = terrain position index; TPI1 = mean TPI1 in the polygon; Tpi1_mp = terrain form TPI1, numerical, mean; Tpi1h_a = convex terrain; Tpi1l_a = depressions; Tpi6l_a = rugged terrain, TPI6; U = bio-ecological variation; VE = Variation explained; Vei_b_a = road; Vkode = variable code; Vmag_a = regulated magazine.

## Abstract

A multitude of landscape characterisation and mapping methods exist, but few methods take into account that landscapes properties vary in a gradual, continuous manner along multiple directions of variation. In this study, we used gradient analytic methods, rooted in ecological continuum theory, to analyse landscape variation throughout Norway. The aim is to explain differences in landscape properties in the simplest possible way, by identifying ‘complex landscape gradients’ (CLGs), i.e. composite gradients of co-occurring landscape elements and properties.

We collected data by stratified sampling of 100 test areas (20×20 km), in which we delineated a total of 3966 observation units (landscape polygons 4–30 km²) based on geomorphological criteria. For each observation unit, 85 landscape variables were recorded. We identified patterns of variation in landscape element composition by parallel use of two multivariate statistical methods, detrended correspondence analysis (DCA) and global nonmetric multidimensional scaling (GNMDS).

The analyses revealed that the most important properties explaining differences in total landscape elements composition was location of the landscape relative to the coastline and coarse-scale landform variation. Most landscape elements had distinct optima within specific segments along broad-scale complex-gradients in landscape properties. A tentative landscape-type hierarchy was built by an iterative procedure by which the amount of compositional turnover in landscape-element composition between adjacent types was standardised. Six ‘major landscape types’ were identified based on geomorphological criteria. Within each major type, we identified a unique set of 2–5 important CLGs, representing geo-ecological, bio-ecological, and land use-related landscape variation. Minor landscape types were obtained by combining segments along two or more CLGs.

The study shows that geological diversity, biological diversity and human land-use are tightly intertwined at the landscape level of ecological complexity, and that predominantly abiotic processes control and constrain both biotic processes and human land use.

## INTRODUCTION

### THE LANDSCAPE LEVEL OF ECOLOGICAL DIVERSITY

Since the 1980s, ecologists have become increasingly aware that the relative importance of factors controlling biotic and abiotic patterns and processes vary with the scale of observation, often being non-linearly related to the gradient from finer towards broader spatial and temporal scales (Swanson 1988, Wiens 1989, McGill 2010b, Zarnetske et al. 2019). Accordingly, any ecological phenomenon (e.g. the diversity of species, ecosystems and landscapes), should be addressed at the spatial and temporal scale at which the phenomenon appears (Estes et al. 2018). Hierarchy theory (Allen & Starr 1982, O’Neil et al. 1986, Urban et al. 1987, King 2005) suggests that ecological systems are implicitly hierarchical, in the sense that the interacting components are organised in ‘levels of organisation’ within a hierarchically organised system. A well-known example of a *biotic* hierarchy is the series cell – organism – population – community (Turner & Gardner 2015). Complex entities at any particular level in such a hierarchy can be explained and studied in terms of entities only one level down in the hierarchy; entities which themselves are likely to be complex enough to need further reduction to their own component parts (King 2005, Allen & Starr 2017). Dawkins (1986) termed the study of such entities ‘hierarchical reductionism’, and pointed out that the kinds of explanations suitable at high levels in the hierarchy may be qualitatively different from the kinds of explanations suitable at lower levels.

Halvorsen et al. (2020) defined ‘ecodiversity’ as ‘the diversity of units defined by biotic as well as abiotic components and their interactions, and the processes that give rise to variation in the structure and composition of these components’. ‘Landscapes’ are often recognised as a separate level of ecological diversity, simultaneously addressing biotic and abiotic variation in heterogeneous areas of kilometres-wide extent (Noss 1990, Allen & Hoekstra 1990, Bailey 2009, Halvorsen et al. 2020). The domain of spatial scales typically applied in landscape characterisation and mapping addresses Nature’s diversity at a scale less detailed than that of, e.g. an ecosystem, and more detailed than that of ecoregions (e.g. Bailey 2014). In this context, landscapes contain biotic and abiotic subsystems such as landforms, ecosystems, meta-ecosystem complexes and other natural and human-induced landscape elements at spatial scales from 10^6^ to 10^10^ m² (often referred to as meso-scale) responding to abiotic and biotic processes occurring over timespans from 10^1^ to 10^4^ years (Delcourt et al. 1982, Dikau 1989).

We define ‘landscape’ as a more or less uniform area characterised by its content of observable, natural and human-induced landscape elements, i.e. natural or human-induced objects or characteristics, including spatial units assigned to types at an ecodiversity level lower than the landscape level, which can be identified and observed on a spatial scale relevant for the landscape level of ecodiversity (Halvorsen et al. 2020). Furthermore, we define ‘landscape element’ as a natural or human-induced object or characteristic, including spatial units assigned to types at an ecodiversity level lower than the landscape level, which can be identified and observed on a spatial scale relevant for the landscape level of ecodiversity (Halvorsen et al. 2020). ‘Landscape types’ are defined as more or less uniform areas characterised by their content of observable, natural and human-induced landscape elements (Halvorsen et al. 2020).

Knowledge about variation at the landscape level of ecodiversity is a prerequisite for knowledge-based spatial planning and management (Marsh 2005). Such knowledge is also considered as essential to fulfil obligations set by international conventions such as the European Landscape Convention (Anonymous 2000) and legal frameworks such as the Norwegian Nature Diversity Act (Anonymous 2009). The primary purpose of the latter is, e.g. to protect ‘biological, geological and landscape diversity’ and promote conservation and sustainable use of the ‘full range of variation of habitats and landscape types’. While the concept of landscape diversity presupposes knowledge about the relative frequency or abundance of discrete entities (landscape types), defining such entities is challenging (Skånes 1997, Simensen et al. 2018). By and large, the composition, structure and functions of landscapes vary in a gradual, continuous manner along multiple ‘directions of gradual variation’ (Halvorsen et al. 2016). For such objects, that are not naturally classified, all type systems are artificial in the sense that they have to be constructed by some set of rules and that many such rule sets may, in principle, be applied (Økland & Bendiksen 1985). Accordingly, a multitude of different methods exist for the characterisation and mapping of ecological diversity at the landscape level (Simensen et al. 2018).

### LANDSCAPE ANALYSIS IN NORWAY

Systematic descriptions and mapping of specific natural phenomena based on type-oriented systems has a long tradition in Norway (e.g. vegetation types: Nordhagen 1917; soil types: Glømme 1935; forest types: Barth 1944; landform types: Rudberg 1960; land-cover types: Einevoll 1965). Although the concept of ‘landscape types’ has been used informally in geographical texts in Norway for a long time (Sømme 1938, Holt-Jensen 2009), comprehensive and evidence-based type systems for the landscape level of ecological diversity, which simultaneously address biotic and abiotic variation, have still not been established. Regional and national landscape characterisation efforts have mostly followed the regional geographic tradition by which the individual character of singular areas such as landscape units or regions is described rather than addressing geographical phenomena in general (i.e. by a type-system; see e.g. Nordisk ministerråd 1984, Nordisk Ministerråd 1987, Puschmann 2005).

Moreover, the few examples of landscape characterisation methods applied at broader scales in Norway have relied strongly on an intuitive, qualitative, expert-based and holistic assessment of the total composition of landscape elements and properties (see, e.g. Nordisk ministerråd 1987, Puschmann 2005). While qualitative descriptions of spatially delineated landscape units are intuitively appealing for communication purposes, the lack of transparency, repeatability and scientific rigour behind expert-based methods will often constrain their usefulness for a broader range of applied and scientific purposes (Simensen et al. 2018). Expert-based methods are prone to a multitude of biases and uncertainties, such as confirmation bias (seeking information that affirms pre-existing beliefs), coverage bias (the extent to which different issues and are reported and considered) and concision bias (selectively focusing on information, losing nuance), possibly leading to inconsistencies and lack of correspondence to observable reality (see e.g. Gelman & Henning 2017). Quantitative analysis, on the other hand, has the great merit of revealing relationships which are inherent in the study material, but otherwise unrecognisable (Whittaker 1966). A high degree of observer independence is a mandatory prerequisite for addressing several specific research questions within landscape ecology and physical geography, such as the study of spatial distribution and abundance of landscape elements, and studies of patterns, structure and processes in the landscape. Alahuhta et al. (2019) and Schrodt et al. (2019) thus maintained that progress in making connections between non-living and living nature requires systematic data collection designed to improve our knowledge of the linkages between biodiversity and geodiversity at several scales and levels. Several Norwegian scientists (e.g. Moen 1999, Strand 2011, Krøgli et al. 2015, and Erikstad et al. 2015) have called for more systematic, observer-independent and repeatable frameworks for the study of the total composition of landscape elements throughout Norway.

### METHODS FOR LANDSCAPE CHARACTERISATION AND MAPPING

Internationally, the recent trend has been towards increasing observer-independence in biophysical landscape characterisation, due to improved availability of area-covering landscape data derived by remote sensing and powerful novel statistical and geographical analysis methods (Simensen et al. 2018, Zarnetske et al. 2019, Yang 2020). A wide variety of ‘landscape metrics’ are now available for analysis of patterns of selected aspects of landscape variation derived from categorical land-cover data (i.e. landscape structure; see Malinowska & Szumacher 2013, Lausch 2015, Hesselbarth et al. 2019). Holistic assessments of the total composition of landscape elements and properties are most often conducted by stepwise, criteria-based GIS overlay techniques, multispectral segmentation, or GIS analysis in combination with multivariate statistical analysis (Simensen et al. 2018, Yang et al. 2020). The latter groups of methods typically include recording of a broad selection of physical landscape attributes within fixed spatial units (e.g. 1×1 km raster cells) followed by multivariate statistical analyses and supervised or unsupervised clustering techniques, to classify or group landscapes with related characteristics (Bailey 2009). A map is produced by drawing lines around cells classified to the same or related classes. Commonly applied clustering methods are Two Way Indicator Species Analysis (TWINSPAN; Hill 1979) and K-means clustering (MacQueen 1967). In Norway, Krøgli et al. (2015) analysed landscape data in geographical grids of 5×5 km and 1×1 km squares, respectively, and identified ten landscape categories with a high degree of internal similarity.

Although classification by numerical clustering procedures have been successful in identifying ‘groups of similar landscapes‘, clustering methods are in general not well suited for the analysis of observation units that vary more or less continuously along gradients, as most often is the case with landscape properties. Much information is lost when multidimensional networks are assigned to clusters within a unidimensional hierarchy because the relationships between the classes at the same level in the hierarchy are hidden (Tuomikoski 1942, Økland & Bendiksen 1985). Since clustering methods do not allow for close examination of the abundance and distribution of landscape elements along multiple gradients and dimensions in the landscape space (cf. Whittaker 1967), the results of cluster analyses of landscape variation are challenging to interpret ecologically (cf. Austin 2002). An additional disadvantage of clustering methods is that the number of clusters, and hence landscape types, must be defined in advance rather than follow from the analysis of data. We argue that alternative approaches are needed to obtain a better understanding of the relationships between geodiversity and biodiversity at the landscape level (cf. Alahuhta et al. 2019, Halvorsen et al. 2020).

### GRADIENT ANALYSIS OF LANDSCAPE VARIATION

An understanding of natural variation based upon knowledge about environmental gradients and species’ responses to these gradients – a gradient perspective on species-environment relationships (Halvorsen 2012) – is supported by evidence from ecosystems all over the world and has served as theoretical foundation for reserach in plant and community ecology for more than 50 years (Gleason 1939, Curtis 1959, Whittaker 1967, Austin 2005). Application of a wide range of methods of gradient analysis (including multivariate gradient analysis, i.e. ordination) has been crucial for disentangling the underlying set of rules that regulate the composition, structure and functions of ecosystems (Økland 1990, McGill 2010b) including microbiological systems (van Elsas et al. 2006). At the landscape level, however, the continuum concept in general, and ordination methods in particular, are rarely applied to understand and describe patterns of variation in landscape element composition. At this level, gradient analysis is usually limited to the use of single continuous variables as an alternative to ‘landscape metrics’ based on categorical landscape data (Cushman et al. 2010, Lausch et al. 2015). Continuous variables at the landscape level can in principle be obtained by two approaches (Lausch 2015): (i) by direct measurements on a continuous scale (e.g. a digital elevation model); or (ii) by deriving continuous variables from categorical land cover data by a moving window approach (McGarigal & Cushman 2005).

A key point in the gradient perspective on species-environment relationships is the concept of the environmental complex-gradient (Whittaker 1956), i.e. that a set of correlated environmental variables act on the species in concert rather than one by one. Numerous studies of species-environment all over the world have underpinned the notion that a few major environmental complex-gradients typically account for a significant fraction of the total variation in species composition that can be explained by variation in the environment (Halvorsen 2013). Moreover, species generally are found within a restricted interval along each major environmental complex-gradient. We hypothesise that the complex-gradient concept, commonly used to understand and describe species’ relationships to the environment, can be extended to the landscape level of ecological diversity. Furthermore, we hypothesise that ecosystem and other landscape elements, just like species, may have distinct optima (maximum probability of occurrence) within specific segments of broad-scale, complex gradients in landscape properties. If this hypothesis is correct, evidence-based and testable type systems can be obtained by combining segments along two or more CLGs based on the amount of compositional turnover (Halvorsen et al. 2020). While the principle of turning a multidimensional space into types by combining gradient intervals has a long history in vegetation ecology (e.g. Tuomikoski 1942, Økland & Eilertsen 1993), similar approaches to the establishment of landscape typologies are uncommon.

In 2016, Halvorsen et al. launched the conceptual framework of ‘Nature in Norway’ (NiN) for a description of landscape variation throughout Norway. NiN was later expanded and developed to a universal system for systematisation of ecological diversity at several scales and levels (EcoSyst; Halvorsen et al. 2020). EcoSyst consists of a set of general principles and methods for systematisation of Nature’s diversity that simultaneously addresses biotic and abiotic variation across different levels of organisation. The EcoSyst framework aims to provide the basis for an evidence-based systematics for all observable aspects of ecodiversity, at spatial scales from microhabitats to landscapes.

A first, pilot, version of a gradient-based landscape type-system was developed within the NiN framework for Nordland county (Erikstad et al. 2015), where 173 landscape variables recorded in 258 observation units were subjected to multivariate analyses to identify ‘landscape gradients’. The study showed that ordination methods allow flexible analyses of landscape variables derived from a wide variety of data sources: (i) count data (e.g. buildings); (ii) areal coverage of various land-cover types; and (iii) continuous variables derived from direct measurements (e.g. remote sensing) or by neighbourhood calculations (e.g. morphometric variables). The study also demonstrated that composite landscape gradients could be obtained from multivariate analyses of landscape element compositional data undertaken to reduce the dimensionality of an n-dimensional landscape-level hyperspace. Nevertheless, the pilot study also revealed a clear need for a standardised procedure to identify parsimonious sets of complex landscape gradients, as well as the need for a standardised method to divide such gradients into segments by transparent and repeatable criteria.

In this study, we expand on the experience gained from the Nordland pilot project and analyse landscape variation by use of a sample of observation units from the entire Norwegian mainland, including coastal areas. We use, for the first time, a novel, gradient-based iterative procedure which involves calculation of ‘ecodiversity distance’, i.e., the extent to which the landscape element composition differs between adjacent candidate types. The current study was conducted as a part of the methodological development of the NiN framework (building on Halvorsen et al. 2016), and has been part of the empirical basis for NiN and, subsequently, the theoretical framework of EcoSyst (Halvorsen et al. 2020).

### AIM AND RESEARCH QUESTIONS

The aims of this study are fivefold:

1. To identify major trends and patterns in landscape variation throughout Norway by applying gradient analysis methods to landscape element compositional data.
2. To study linkages between biodiversity and geodiversity at the landscape level of ecological diversity by analysing the relative importance of co-occurring landscape elements and properties within various functional categories, i.e. geo-ecological, bio-ecological and land-use related landscape variation.
3. To examine if, and in case to what extent, landscape elements have distinct optima along broad-scale, complex gradients in landscape property composition, that is, intervals in which they reach maximum occurrence probability.
4. To explain variation in landscape element composition and properties in the simplest possible way, by identifying a parsimonious set of gradients of co-ordinated variation.
5. To construct the first version of an evidence-based landscape-type hierarchy based on the EcoSyst principles (Halvorsen 2020), based on standardisation of the amount of landscape element compositional turnover between adjacent types.

### STRUCTURE OF THE ARTICLE

Our study was conducted in several stages or phases, the interpreted results from each phase forming the platform on which the next phase was built. The structure of this paper reflects the structure of the study, by presenting the analytic results and their interpretation together for each consecutive phase in a combined ‘Results and interpretation’ chapter. The final chapter contains a more general discussion of the overall results and methodological issues.

## MATERIAL AND METHODS

### STUDY AREA

This study area spans latitudes from 57°57’N to 71°11’N and longitudes from 4°29’E to 31°10’E and comprises the entire mainland of Norway including the coastal zone, but excluding the Svalbard archipelago, Jan Mayen and Bear Island. The range of variation in natural conditions found in Norway includes most of the variation that can be found in the circumboreal zone (Bryn et al. 2018), including both terrestrial, marine, limnic and snow and ice ecosystems (Halvorsen et al. 2016). The study area is characterized by a wide range of climatic variation; all seven temperature-related vegetation zones commonly recognised in northern Europe, from boreo-nemoral to high alpine, occur in Norway (Bakkestuen et al. 2008). Norway has a high mineral and bedrock diversity (Ramberg et al. 2008), and a high diversity of landforms (Gjessing 1978).

In addition to natural variation, the diversity of ecosystems in Norway is enhanced by variation in human land use over time, that has affected the distribution and structure of ecosystems throughout the country. Domestic animals and agriculture spread from the Middle East to Norway around 3700 BC (Emanuelsson 2009), and by 1000 BC agriculture had consolidated in the coastal regions of northern Norway as well. From the sixteenth century, the Sami people started to herd semi-domesticated reindeer (Hansen & Olsen 2004). From this time onwards, most Norwegian ecosystems have, to some extent, been affected by land-use activities such as domestic grazing, outfield fodder collection, heath burning, reindeer husbandry, forestry, and industrial, urban and recreational development (Almås et al. 2004). During the twentieth century, extensive land use based on collection of outfield resources gradually declined. Recent landscape changes in Norway are among others related to urbanisation (with associated changes in land use), intensified land use in central or highly productive areas, abandonment of marginal agricultural areas, rapid forest encroachment in previously open areas, reduction of areas with permanent snow cover and translocation of the forest line into higher altitudes (Bryn et al. 2013, Fjellstad & Dramstad 1999).

Note that references to counties and municipalities in this study follow the administrative division as it was in 2018.

### ESTABLISHMENT OF OBSERVATION UNITS

A total of 100 sample areas, each 20×20 km, were selected using a stratified sampling strategy. A subjective component was included in the sampling procedure to ensure that as much as possible of the known variation in landscape properties in Norway was represented in the data material (Fig. 1). Within each sample area, polygons/observation units (hereafter: OUs) were delineated in terms of spatial landscape units using the criteria from the ‘pilot Nordland project’ (see Erikstad et al. 2015; Halvorsen et al. 2016 and Supplementary material Appendix S1). All 4166 OUs with more than a pre-defined fraction of their total area within the sample area were included in the study (also the parts of the OUs outside the border of the sample area limit were included, as indicated by the rugged outline of the OU clusters in Figs 2–3). All OUs were subjected to initial control, which formed the basis for removing 199 OUs which failed to meet the following two criteria: (1) containing terrestrial land (193 marine OUs); or (2) obvious error in the delimitation (digitisation error, etc.; 6 OUs).

**Figs. 1–3.**
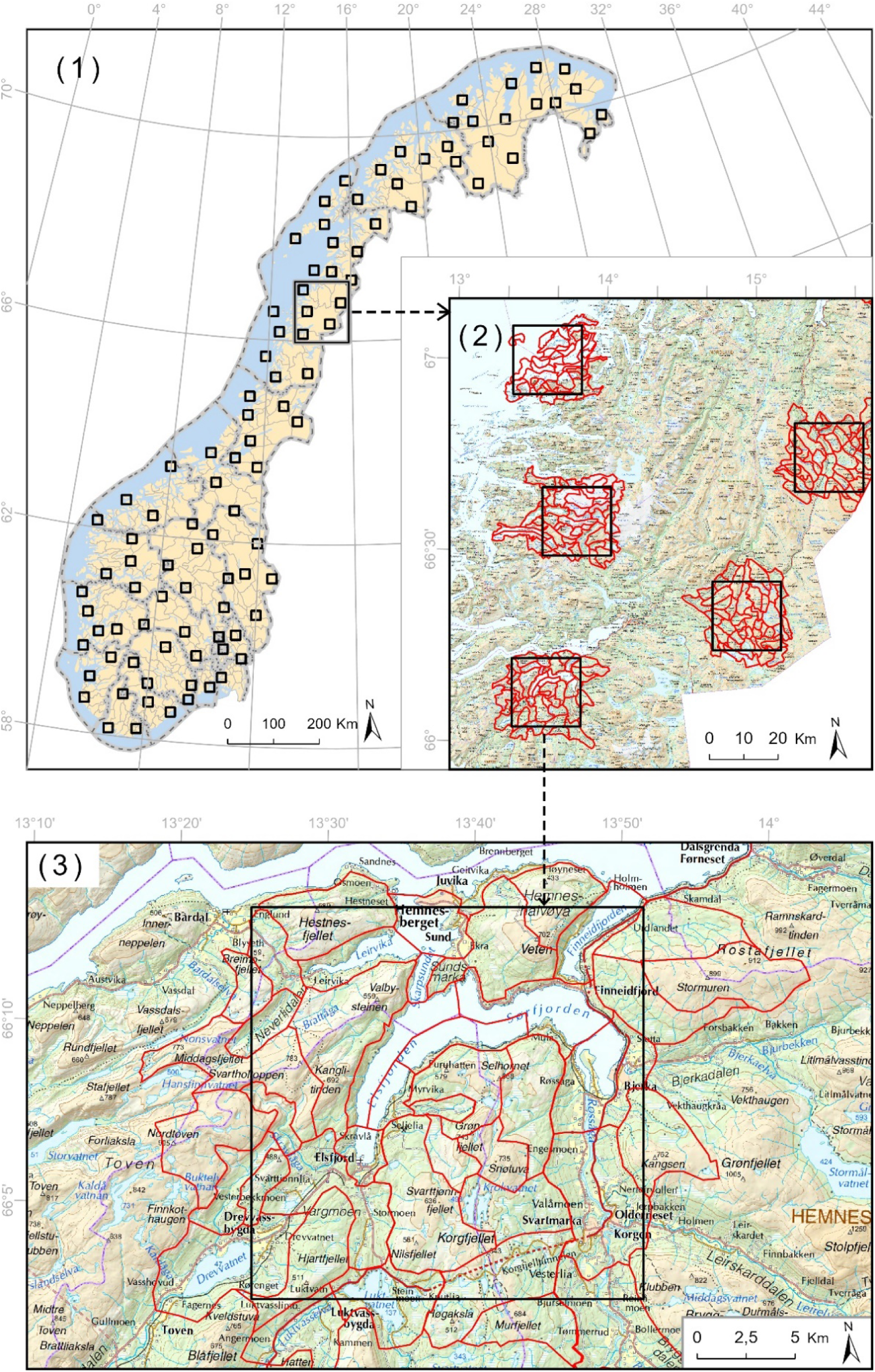
(1) Placement of the 100 sample areas within Norway. (2) Within each sample area, observation units were delineated in terms of spatial landscape units using the criteria from the ‘pilot Nordland project’ (Supplementary material Appendix S1). (3) Example showing delineation of observation units in Hemnes, Norland. Background maps: Norwegian Mapping Authority, 2020.

The data set consisting of the remaining 3967 OUs was subjected to variable transformation, etc. After initial analyses, polygon ID879, which should have been split into two polygons according to the ‘pilot Nordland criteria’, was also removed. The total data set used for the analyses therefore consisted of 3966 OUs.

The OUs were hierarchically nested within sample areas, and cannot be regarded as strictly statistically independent observations. Based on the knowledge that spatial autocorrelation in the data will tend to inflate the rate of Type I errors (rejection of a true null hypothesis), we interpreted all statistical test conservatively, using test results for indication rather than for hypothesis testing in the strict statistical sense (cf. Økland 2007).

### *A PRIORI* CHARACTERISATION OF OBSERVATION UNITS

For the purpose of interpretation of subsequent analyses, all OUs were characterised *a priori* (i.e. before analyses) by assigning to each OU values for up to four types of variables.

(1) *Landscape-type variables* (LT variables), which affiliate the OUs with tentative major-type group (Table 1) and major type at the landscape-type level in NiN version 2.0, i.e. based upon ‘pilot Nordland criteria’ as implemented in the ‘pilot Nordland project’ (Supplementary material, Appendix S1).

**Table 1.** Landscape-type (LT) variables: major-type groups and major types at the landscape level in NiN version 2.0, according to the ‘pilot Nordland project’. The LT variables are used for initial sorting of observation units (OUs).

| Major type group | Major type |
| --- | --- |
| Inland landscape (I) | Valleys (D)<br>Inland plains (S)<br>Hills and mountains (A) |
| Coastal landscape (K) | Fjords (F)<br>Coastal plains (S)<br>Coastal hills and mountains (A) |
| Marine landscape (M) | Marine continental slope (K)<br>Marine valleys (D)<br>Marine coastal plains (S)<br>Marine hills- and mountains (A) |

(2) *Landscape-gradient variables* (LG variables), which are ordered factor variables, one for each landscape gradient identified in the ‘pilot Nordland Project’, upon which the division into minor types in NiN version 2.0 was based. The LG variables (Table 2) provide a more detailed characterisation of the OUs than the LT variables. An LG variable was used to characterise an OU when the OU was part of a data subset in which the variable was relevant to all OU members.

**Table 2.** Landscape-gradient (LG) variables used for initial characterisation of OUs (assignment of OUs to LG segments). ‘Segment terms’ are given for each segment as ordered from 1 to ‘No. of segments’. ‘Normal’ segments are italicised; extreme steps that specifies major types or major-type groups in which the variable is not relevant, are given in bold face. LG variables are characterised as ‘new’ if they result from the analytic process in the present study, and as ‘rep(laced)’ if used in NiN version 2.0 only.

| Code | Term | Segment terms | No. of segments |
| --- | --- | --- | --- |
| DN | Valley shape | open \| relatively open \| relatively narrow and steep \| narrow and steep-sided [valley form] | 4 |
| RE | Relief | flat/irregular plains \| low hills \| hilly | 3 |
| TP | Peaks | *absence* \| presence | 2 |
| BP | Glacier | *absence* \| presence | 2 |
| KL | Coastline (*new*) | *absence* \| presence | 2 |
| SP | Coastal/archipelago properties (*rep.*) | **terrestrial** \| *coast-line landscape* \| large islands \| small islands \| skerries/islets \| **marine** | 6 |
| SN | Coastal/archipelago properties (*new*) | **terrestrial** \| *coast-line landscape* \| large islands (> 20 km<sup>2</sup>) \| medium sized islands (1.5–20 km<sup>2</sup>) \| small islands (0.1–1.5 km<sup>2</sup>) \| skerries/islets \| **marine** | 6 |
| BA | Boreal/alpine landscape | Boreal landscapes \| forest line landscapes \| alpine landscapes | 3 |
| IP | Abundance of freshwater lakes | *low* \| high | 2 |
| MP | Abundance of mire | *low* \| high | 2 |
| OI | Amount of infrastructure | without \| low \| medium \| relatively high \| high (town/city) \| very high [city] | 6 |
| JP | Agricultural land-use intensity | insignificant \| moderate \| high | 3 |
| FIA | Amount of flat land (*new*) | steep (proportion of pixels with slope < 2° less than 0.45) \| relatively steep (prop. 0.45–0.65) \| sloping (prop. 0.65–0.75) \| undulating (prop. 0.75–0.85) \| plain (prop. > 0.85) | 5 |
| SS | Sediment sorting (*new*) | unsorted (<10% of area covered by sorted sediments, i.e., glaciofluvial and marine deposits \| low sorting (10–50% sorted sediments) \| medium sorting (50–75% sorted sediments) \| sediment plain (> 75% sorted sediments) | 4 |

(3) Three categorical (unordered) *landscape-factor variables* (LF variables), which addressed sediment/soil properties, were added to the LT and LG variables for descriptive purposes (Table 3). The three variables were: (i) *Sediment category* (SK1), a hierarchically nested categorical variable with classes defined on the basis of soil variables and additional information on the proportion of sorted sediments. (ii) *Primary soil/sediment class* (JK1), an unordered factor variable with up to 5 classes for dominant soil type. An OU was assigned to a JK1 class when more than 50% of its area was covered by the sediment/soil type in question. (iii) *Secondary soil/sediment class* (JK2), an unordered factor variable with up to 5 classes for co-dominant soil/sediment class. An OU was assigned to a JK2 class when more than 25% of its area was covered by the sediment/soil type in question.

**Table 3.**
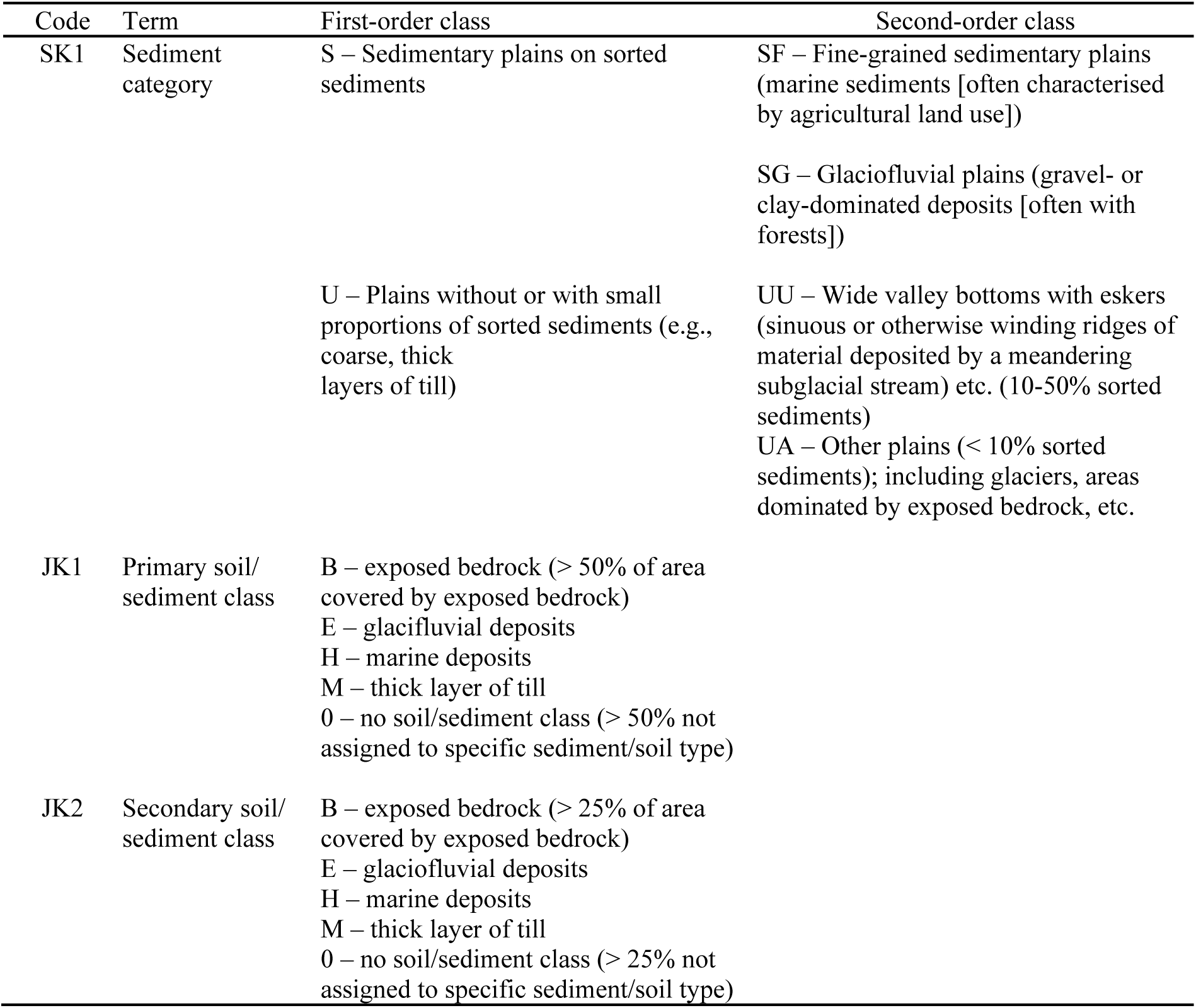
Landscape-factor (LF) variables used for initial characterisation of OUs (assignment of OUs to LF class). Some of the variables are structured as sets of hierarchically nested classes.

(4) *Key variables*, defined in NiN 2.0 as ‘observable landscape properties or variables derived from such properties, which are used for segmentation of a (complex) landscape gradient’, and used in the ‘pilot Nordland project’ (Erikstad et al. 2015) to operationalise the division of complex landscape gradients into segments. The raw key variables, here termed ‘primary key variables’ (Table 4), were obtained by calculating focal statistics (neighbourhood calculation; see e.g. Lovelace et al. 2019) for one or more landscape properties for the 81 100×100 m pixels with more than half of their area contained within a neighbourhood circle of radius 500 m, centered on the focal 100×100 m pixel (Fig. 4). The mean value for each relevant primary key variable in each OU was used for complementary characterisation of OU positions along (complex) landscape gradients.

**Fig. 4.**
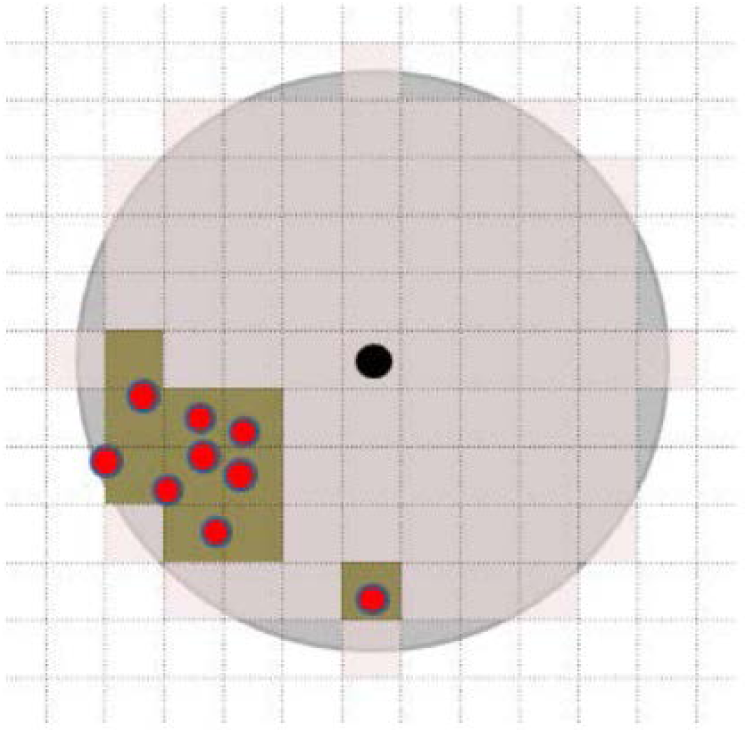
The principle of key variable calculation as a focal statistic (frequency) within 81 standardised grid cells of 100×100 m. The 81 cells (marked with pink shadow) are wholly or partly contained within a circle (marked with grey shadow) with a radius of 500 m around one focus point for which the key variable is calculated (marked with a black dot located in the middle of a cell). In the example, the property the key variable is based on (e.g., presence of buildings) indicated by red dots. In this example, the key variable has the value 10, or alternatively 10/81 = 0.123 if given as frequency.

**Table 4.**
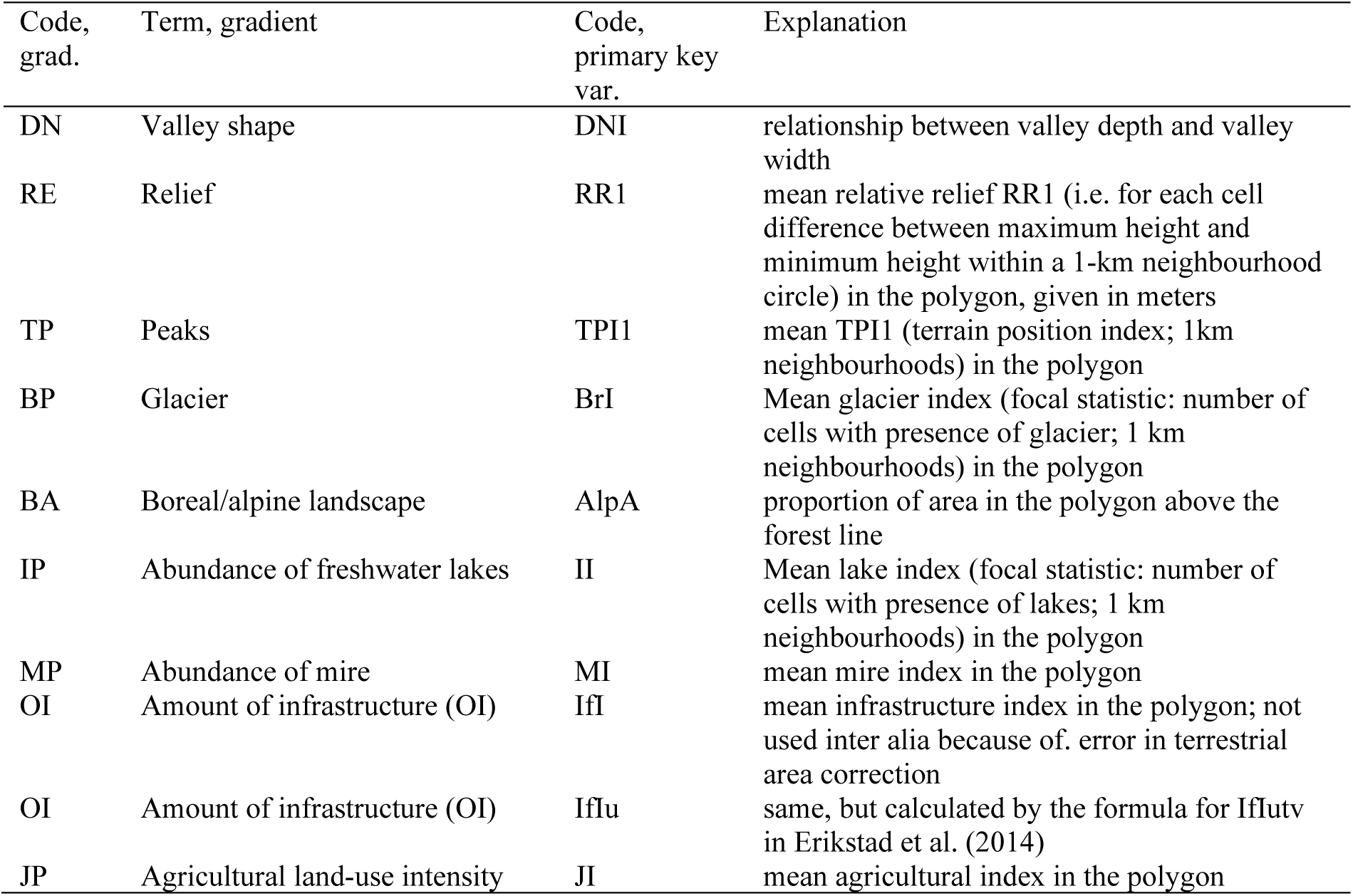
Primary key variables used for complementary characterisation of the observation units (OUs).

### SELECTION AND CATEGORISATION OF ANALYTIC VARIABLES FOR STATISTICAL ANALYSES

For all of Norway, information about properties that were observable on a landscape-relevant scale were collected from available maps and databases. This information was organised into 111 primary descriptive variables, each represented by a binary (0/1) value for each pixel (cell) in a raster with 100 m resolution, covering all of Norway. The binary variables thus indicated presence or absence of the property in each 100×100 m pixel. These primary variables are referred to as ‘ecological basic maps’. Based on these, secondary, or analytic, variables were derived to characterise the OUs of this study. Analytic variables may represent mean values for primary variables, proportions of OU area with value of the primary variable above a given threshold, etc. This is further explained below.

Many of the 111 secondary variables had strong pair-wise correlations. In order to avoid a disproportionate emphasis on properties represented in the data by many, strongly correlated, variables, secondary variables were successively removed until we obtained a set of variables in which no variable pair had a Kendall’s rank correlation coefficient of τ > 0.7. This pre-selection reduced the number of variables to 85 (Table 5). This set of variables, which will be referred to as ‘the total analytic variables set’, contains variables that can be sorted into three statistical variable categories based on their relationship to the primary variables:

**Table 5.**
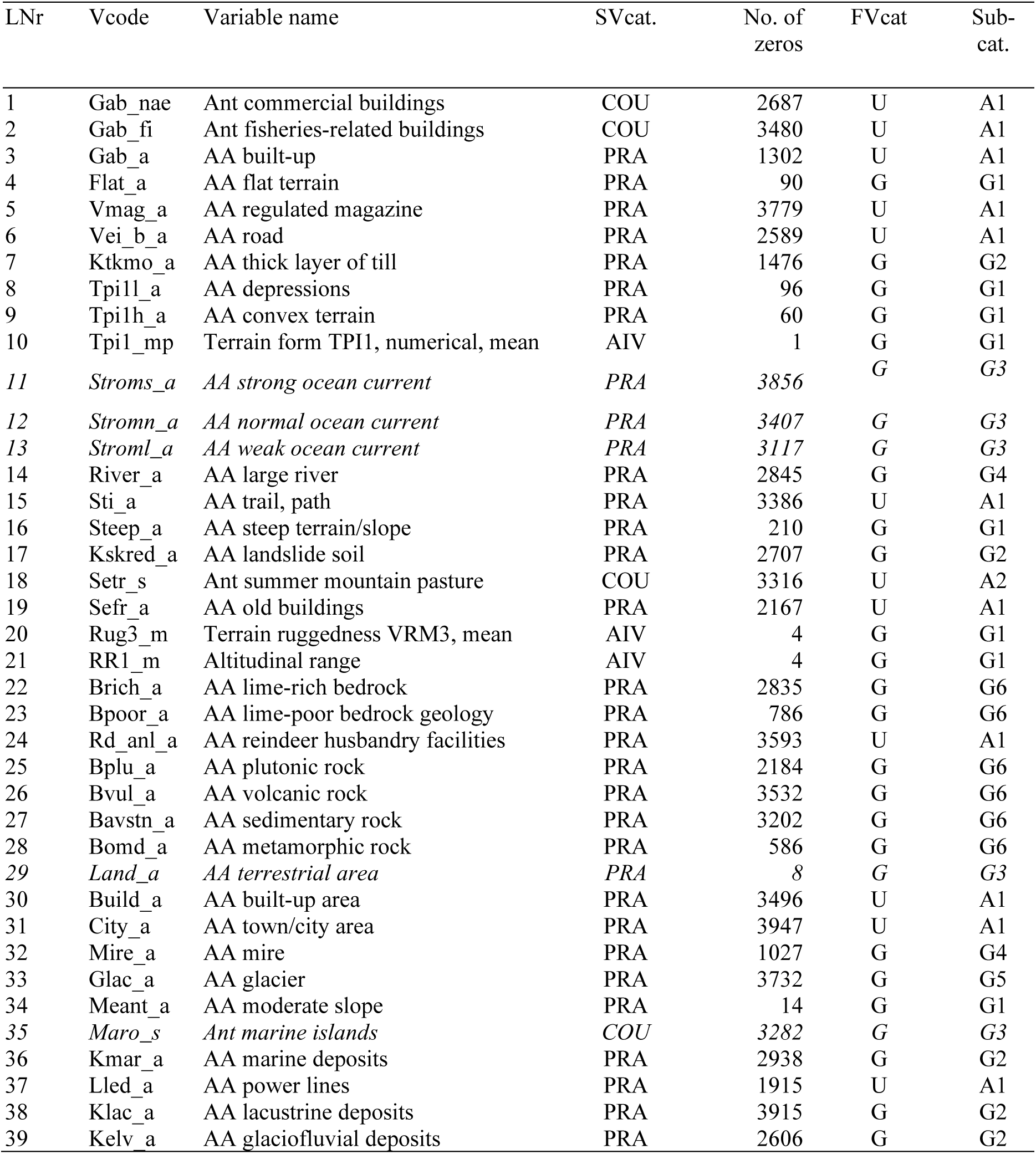
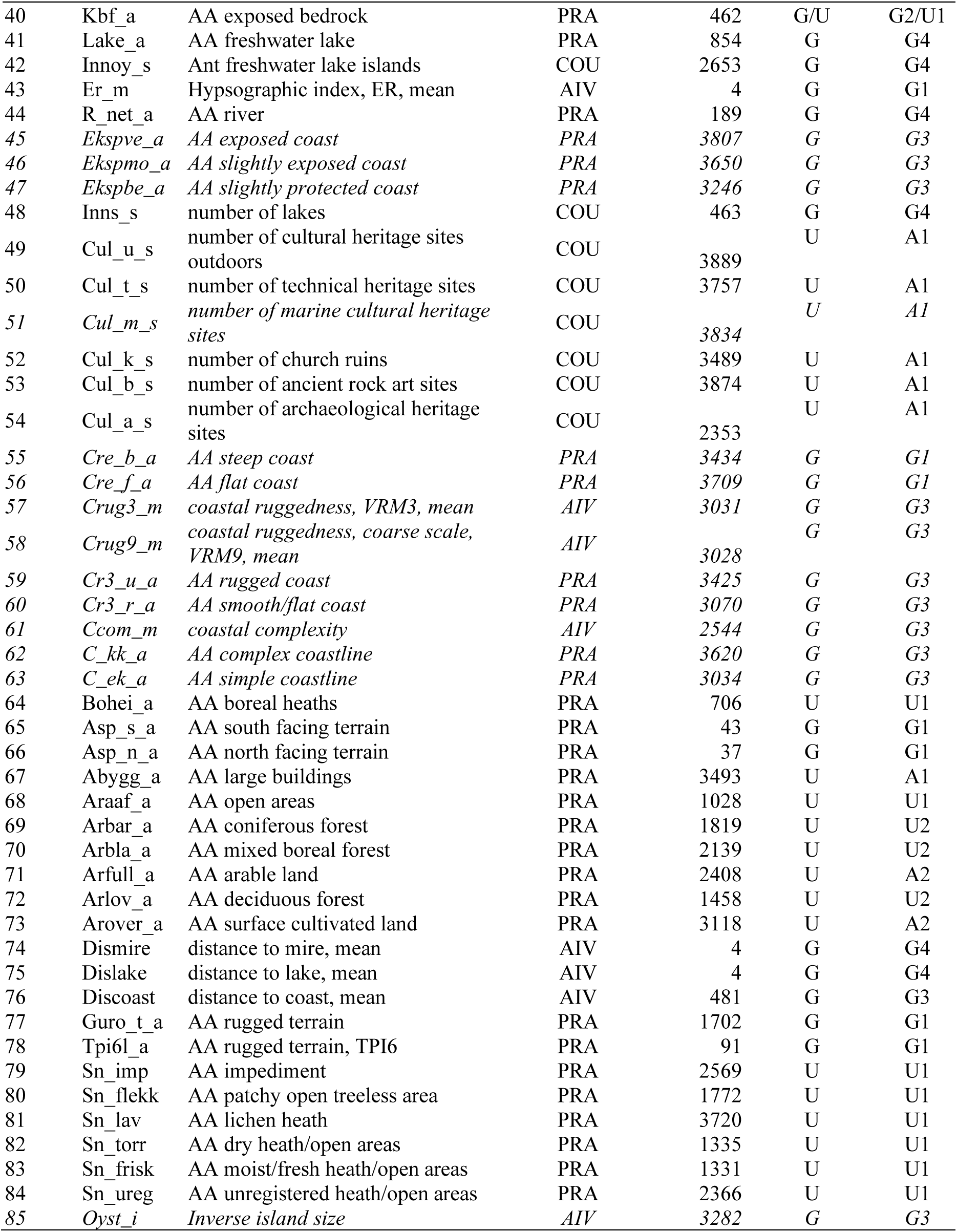
Overview over analytic variables used to characterise the observation units: LNr = serial number; Vcode = variable code, used in all Tables etc; Variable name = full name used in Tables etc. (Ant = Number; AA = Area); SVcat = Statistical variable category; COU = count variables (number/km²); PRA = proportion of area (pixel-number variables); AIV = area-independent variables (variable-specific scales); Count 0 = Number of observations with value 0 (only specified for variable categories O and P); FVcat = Functional variable category [(G = geo-ecological variation; U = bio-ecological variation; A = Land Use Variation)]; Subcategory = Tentative assignment of the variable to a CLG theme [(G1) Topographic variables; (G2) Soils; (G3) The variation from inner to outer coast (‘coastal/archipelago properties’); (G4) Water-related variables; (G5) Glaciers; (G6) Bedrock; (U1) Open area cover; (U2) Forest cover; (A1) Infrastructure; (A2) Agricultural land use]. Variables that are not relevant for OUs without coastline are italicised.

| LNr | Vcode | Variable name | SVcat. | No. of zeros | FVcat | Sub-cat. |
| --- | --- | --- | --- | --- | --- | --- |
| 1 | Gab_nae | Ant commercial buildings | COU | 2687 | U | A1 |
| 2 | Gab_fi | Ant fisheries-related buildings | COU | 3480 | U | A1 |
| 3 | Gab_a | AA built-up | PRA | 1302 | U | A1 |
| 4 | Flat_a | AA flat terrain | PRA | 90 | G | G1 |
| 5 | Vmag_a | AA regulated magazine | PRA | 3779 | U | A1 |
| 6 | Vei_b_a | AA road | PRA | 2589 | U | A1 |
| 7 | Ktkmo_a | AA thick layer of till | PRA | 1476 | G | G2 |
| 8 | Tpi1l_a | AA depressions | PRA | 96 | G | G1 |
| 9 | Tpi1h_a | AA convex terrain | PRA | 60 | G | G1 |
| 10 | Tpi1_mp | Terrain form TPI1, numerical, mean | AIV | 1 | G | G1 |
| 11 | *Stroms_a* | *AA strong ocean current* | *PRA* | *3856* | *G* | *G3* |
| 12 | *Stromn_a* | *AA normal ocean current* | *PRA* | *3407* | *G* | *G3* |
| 13 | *Stroml_a* | *AA weak ocean current* | *PRA* | *3117* | *G* | *G3* |
| 14 | River_a | AA large river | PRA | 2845 | G | G4 |
| 15 | Sti_a | AA trail, path | PRA | 3386 | U | A1 |
| 16 | Steep_a | AA steep terrain/slope | PRA | 210 | G | G1 |
| 17 | Kskred_a | AA landslide soil | PRA | 2707 | G | G2 |
| 18 | Setr_s | Ant summer mountain pasture | COU | 3316 | U | A2 |
| 19 | Sefr_a | AA old buildings | PRA | 2167 | U | A1 |
| 20 | Rug3_m | Terrain ruggedness VRM3, mean | AIV | 4 | G | G1 |
| 21 | RR1_m | Altitudinal range | AIV | 4 | G | G1 |
| 22 | Brich_a | AA lime-rich bedrock | PRA | 2835 | G | G6 |
| 23 | Bpoor_a | AA lime-poor bedrock geology | PRA | 786 | G | G6 |
| 24 | Rd_anl_a | AA reindeer husbandry facilities | PRA | 3593 | U | A1 |
| 25 | Bplu_a | AA plutonic rock | PRA | 2184 | G | G6 |
| 26 | Bvul_a | AA volcanic rock | PRA | 3532 | G | G6 |
| 27 | Bavstn_a | AA sedimentary rock | PRA | 3202 | G | G6 |
| 28 | Bomd_a | AA metamorphic rock | PRA | 586 | G | G6 |
| 29 | *Land_a* | *AA terrestrial area* | *PRA* | *8* | *G* | *G3* |
| 30 | Build_a | AA built-up area | PRA | 3496 | U | A1 |
| 31 | City_a | AA town/city area | PRA | 3947 | U | A1 |
| 32 | Mire_a | AA mire | PRA | 1027 | G | G4 |
| 33 | Glac_a | AA glacier | PRA | 3732 | G | G5 |
| 34 | Meant_a | AA moderate slope | PRA | 14 | G | G1 |
| 35 | *Maro_s* | *Ant marine islands* | *COU* | *3282* | *G* | *G3* |
| 36 | Kmar_a | AA marine deposits | PRA | 2938 | G | G2 |
| 37 | Lled_a | AA power lines | PRA | 1915 | U | A1 |
| 38 | Klac_a | AA lacustrine deposits | PRA | 3915 | G | G2 |
| 39 | Kelv_a | AA glaciofluvial deposits | PRA | 2606 | G | G2 |
| 40 | Kbf_a | AA exposed bedrock | PRA | 462 | G/U | G2/U1 |
| 41 | Lake_a | AA freshwater lake | PRA | 854 | G | G4 |
| 42 | Innoy_s | Ant freshwater lake islands | COU | 2653 | G | G4 |
| 43 | Er_m | Hypsographic index, ER, mean | AIV | 4 | G | G1 |
| 44 | R_net_a | AA river | PRA | 189 | G | G4 |
| 45 | Ekspve_a | AA exposed coast | PRA | 3807 | G | G3 |
| 46 | Ekspmo_a | AA slightly exposed coast | PRA | 3650 | G | G3 |
| 47 | Ekspbe_a | AA slightly protected coast | PRA | 3246 | G | G3 |
| 48 | Inns_s | number of lakes | COU | 463 | G | G4 |
| 49 | Cul_u_s | number of cultural heritage sites outdoors | COU | 3889 | U | A1 |
| 50 | Cul_t_s | number of technical heritage sites | COU | 3757 | U | A1 |
| 51 | Cul_m_s | number of marine cultural heritage sites | COU | 3834 | U | A1 |
| 52 | Cul_k_s | number of church ruins | COU | 3489 | U | A1 |
| 53 | Cul_b_s | number of ancient rock art sites | COU | 3874 | U | A1 |
| 54 | Cul_a_s | number of archaeological heritage sites | COU | 2353 | U | A1 |
| 55 | Cre_b_a | AA steep coast | PRA | 3434 | G | G1 |
| 56 | Cre_f_a | AA flat coast | PRA | 3709 | G | G1 |
| 57 | Crug3_m | coastal ruggedness, VRM3, mean | AIV | 3031 | G | G3 |
| 58 | Crug9_m | coastal ruggedness, coarse scale, VRM9, mean | AIV | 3028 | G | G3 |
| 59 | Cr3_u_a | AA rugged coast | PRA | 3425 | G | G3 |
| 60 | Cr3_r_a | AA smooth/flat coast | PRA | 3070 | G | G3 |
| 61 | Ccom_m | coastal complexity | AIV | 2544 | G | G3 |
| 62 | C_kk_a | AA complex coastline | PRA | 3620 | G | G3 |
| 63 | C_ek_a | AA simple coastline | PRA | 3034 | G | G3 |
| 64 | Bohei_a | AA boreal heaths | PRA | 706 | U | U1 |
| 65 | Asp_s_a | AA south facing terrain | PRA | 43 | G | G1 |
| 66 | Asp_n_a | AA north facing terrain | PRA | 37 | G | G1 |
| 67 | Abygg_a | AA large buildings | PRA | 3493 | U | A1 |
| 68 | Araaf_a | AA open areas | PRA | 1028 | U | U1 |
| 69 | Arbar_a | AA coniferous forest | PRA | 1819 | U | U2 |
| 70 | Arbla_a | AA mixed boreal forest | PRA | 2139 | U | U2 |
| 71 | Arfull_a | AA arable land | PRA | 2408 | U | A2 |
| 72 | Arlov_a | AA deciduous forest | PRA | 1458 | U | U2 |
| 73 | Arover_a | AA surface cultivated land | PRA | 3118 | U | A2 |
| 74 | Dismire | distance to mire, mean | AIV | 4 | G | G4 |
| 75 | Dislake | distance to lake, mean | AIV | 4 | G | G4 |
| 76 | Discoast | distance to coast, mean | AIV | 481 | G | G3 |
| 77 | Guro_t_a | AA rugged terrain | PRA | 1702 | G | G1 |
| 78 | Tpi6l_a | AA rugged terrain, TPI6 | PRA | 91 | G | G1 |
| 79 | Sn_imp | AA impediment | PRA | 2569 | U | U1 |
| 80 | Sn_flekk | AA patchy open treeless area | PRA | 1772 | U | U1 |
| 81 | Sn_lav | AA lichen heath | PRA | 3720 | U | U1 |
| 82 | Sn_torr | AA dry heath/open areas | PRA | 1335 | U | U1 |
| 83 | Sn_frisk | AA moist/fresh heath/open areas | PRA | 1331 | U | U1 |
| 84 | Sn_ureg | AA unregistered heath/open areas | PRA | 2366 | U | U1 |
| 85 | Oyst_i | Inverse island size | AIV | 3282 | G | G3 |

(1) *Pixel-number variables* (PRA) were obtained by counting the number of 100×100 m pixels (grid cells) in which the primary variable in question had a value of 1, i.e. the landscape property in question was recorded as present (‘presence pixels’). Pixel-number variables *y* were expressed as area fractions calculated as:

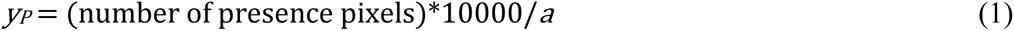

where *a* is the total area of the OU, given in m^2^.

(2) *Count, or density, variables* (COU) were primarily obtained as counts of the actual number of observable objects of a given category in an OU, e.g. the actual number of buildings in the OU (and *not* the number of 100×100 m pixels containing at least one such object). The counts were standardised to densities by dividing the actual counts with OU area *a*, given in km²:

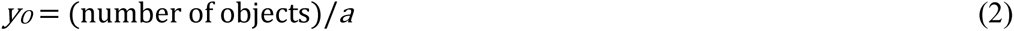

(3) *Area-independent variables* (AIV) were used to characterise the polygons *as such*, recorded on the original, area-independent scale. A-type variables lose their meaning if transformed to area fractions or densities. Examples of such variables are terrain ruggedness, coastline complexity and altitudinal range.

Among the area-independent variables, Oyst_i, ‘inverse island size’, required particular attention. The Oyst_i variable is an index derived from island size, calculated as:

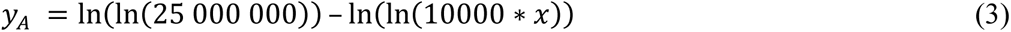

where *x* is the size of the largest island within the OU, given in km². This transformation reversed the direction of the original variable *x*, providing low values for the mainland and high values for the OUs in which the largest islands were small skerries and islets. The value *x* = 2500 was used to represent the mainland as the largest island along the coast of the Norwegian mainland, Hinnøya, comprises 2204 km².

The 85 analytic variables were also, regardless of statistical variable category, sorted into three functional variable categories with subcategories, based on the type of landscape property they describe:

1. *Basic geo-ecological variables* (category G) describe basic natural conditions that typically result in easily observable landscape properties (large-scale geomorphological features such as lakes and rivers, glaciers, and other specific landforms). The G category comprised 6 subcategories: (G1) Topographic variables; (G2) Soils; (G3) The variation from inner to outer coast (‘coastal/archipelago properties’); (G4) Water-related variables; (G5) Glaciers; (G6) Bedrock (transition to category U).
2. *Bio-ecological variables* (category U) describe vegetation properties (open areas, forest cover etc.) and geo-ecological conditions that are not directly observable, but instead manifest themselves in the occurrence or distribution of observable landscape properties (e.g. ‘geological richness’). The U category comprised 2 subcategories: (U1) Open area cover; (U2) Forest cover.
3. *Land-use variables* (category A) describe observable expressions of human activity, typically different types of infrastructure or agriculture. The A category comprised 2 subcategories: (A1) Infrastructure; (A2) Agricultural land use.

### TRANSFORMATION OF ANALYTIC VARIABLES

All analytic variables were transformed to zero skewness (Økland et al. 2001) by the equations:

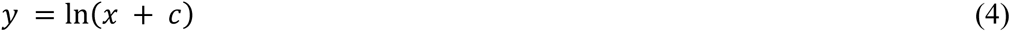

for variables with a right-skewed distribution, and:

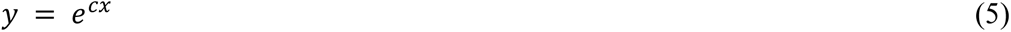

for variables with a left-skewed distribution; *y* is the raw transformed value and *x* the value on the original scale. After transformation, each variable was ranged, i.e. converted to values z on a standard scale from 0 to 1 by the equation:

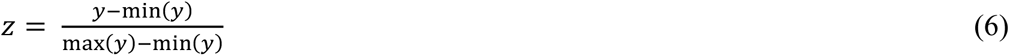

Variables with high frequency of 0 values and hence with a strongly right-skewed distribution (e.g. variables that represent the frequency of rare phenomena) could not be transformed to zero skewness. For variables with more than *ca.* 50% zero observations, skewness is often minimised by transforming all presence observations into one and the same value, that is, by transformation into a binary variable. Such a transformation does, however, incur massive loss of information. As a compromise between the individually desirable but incompatible goals of skewness reduction and preservation of quantitative information in variables with many zero values, we used the value for the transformation constant *c* which resulted in the lowest value of *x* for presence (lowest original value > 0) of z_min_ = 0.200 after transformation and ranging. The highest transformed presence value, z_max_ = 1.000 then became 5 × z_min_ (the lowest presence value).

Transformed and ranged variables can be back-transformed to the original scale by solving equations (4) or (5) inserted in (6) for *x*:

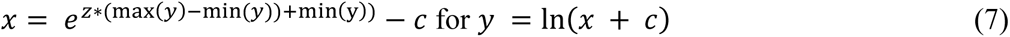

and

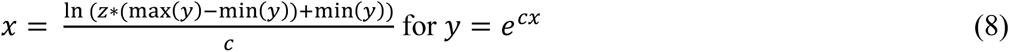

Relationships between original (untransformed) and transformed values for analytic variables that are particularly relevant for assignment of OUs to segments along complex landscape gradients, delineation of major types, etc., are given in Tables 6 and 7.

**Table 6.**
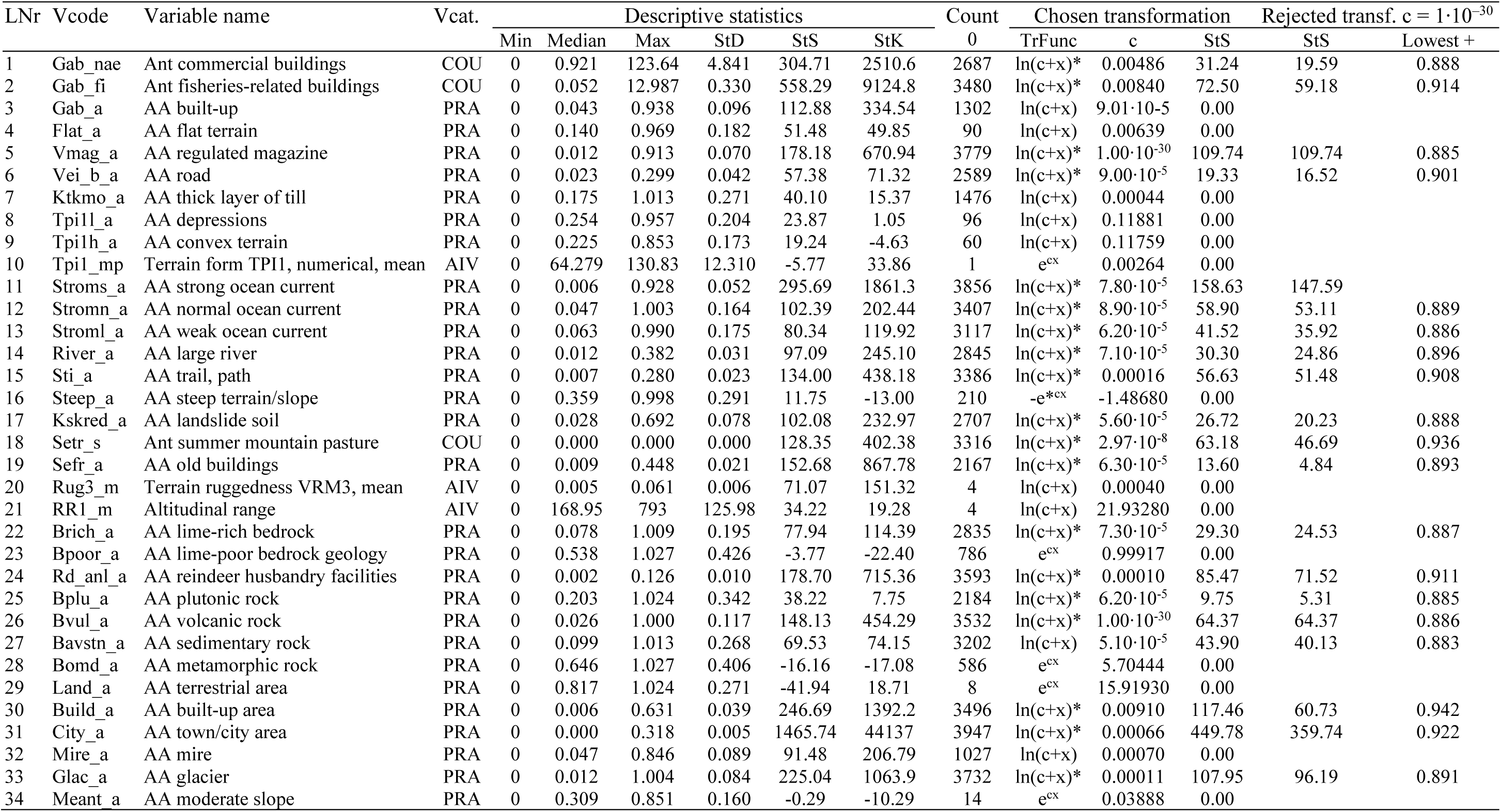
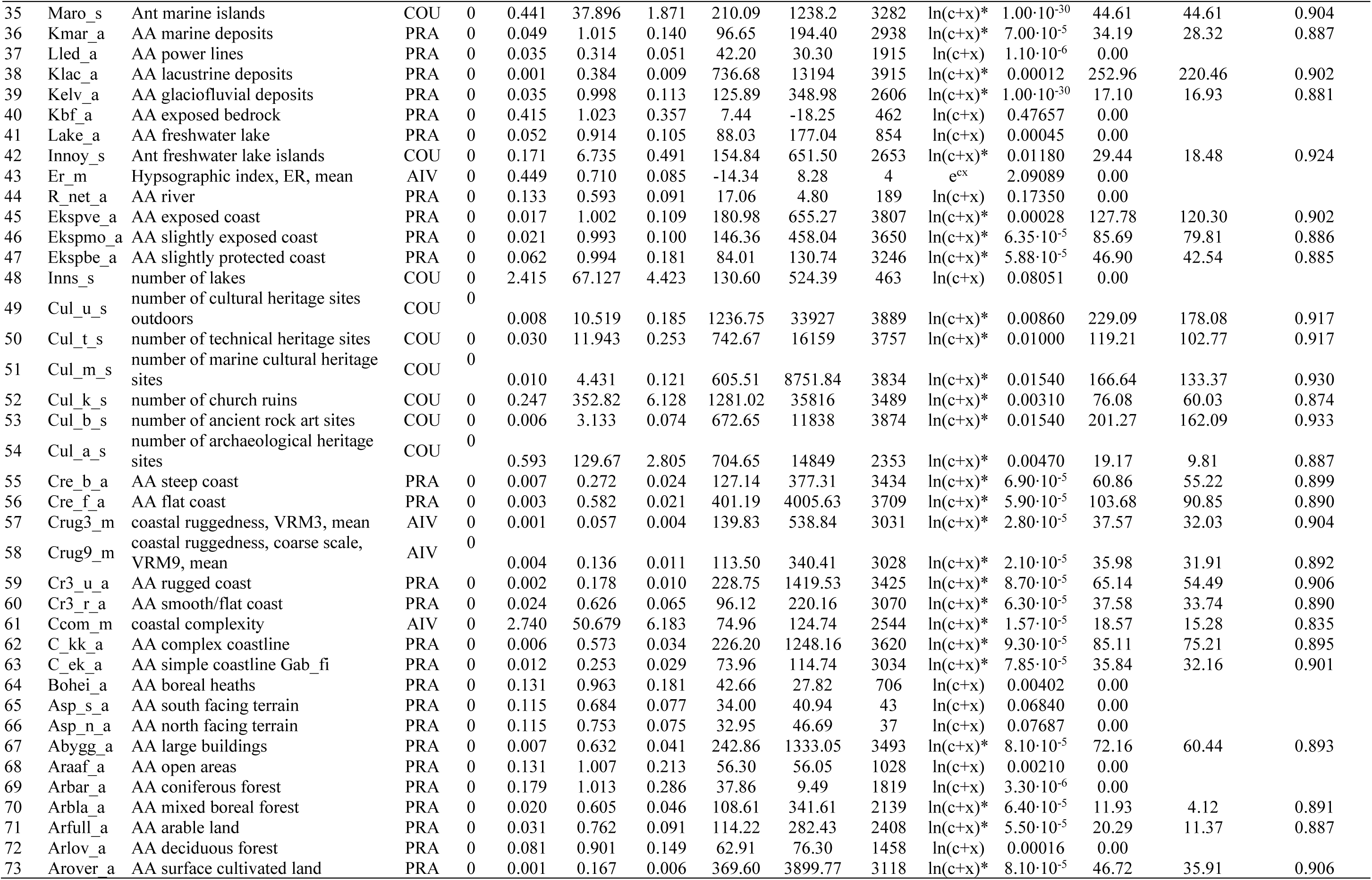
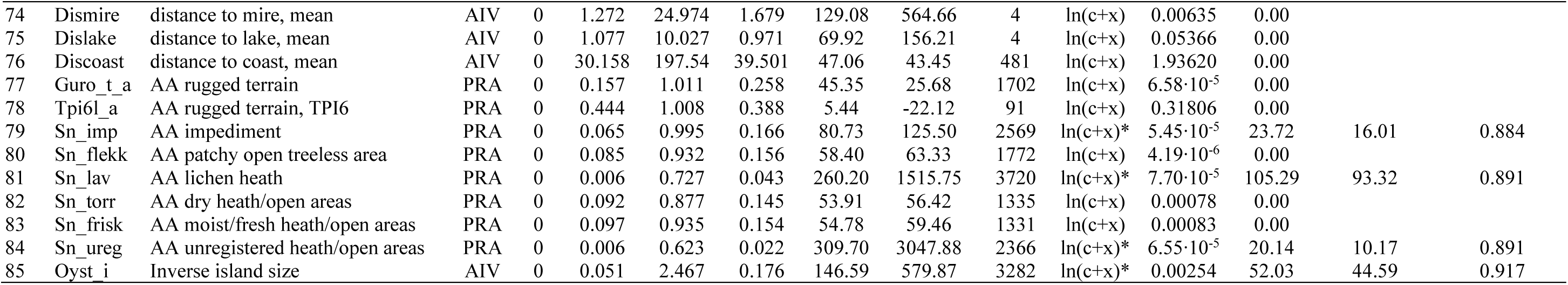
Descriptive statistics and transformations of the analytic variables in Table 5: LNr = serial number; Vcode = variable code; Vcat. = variable category; COU = count variables; PRA = pixel number variables, i.e. proportion of area; AIV = area-independent variables; Descriptive statistics = ; COU; 0 = Number of observations with value 0; TrFunc = chosen transformation function; c = constant; StS = standardised skewness, Lowest + = value for lowest degree of presence of the object in question, after transformation with c = 1·10^−30^.

**Table 7.** Original and transformed values for proportion of areas for soil types.

| Untransformed values (proportion of area) |  | 0.01 | 0.05 | 0.10 | 0.25 | 0.50 |
| --- | --- | --- | --- | --- | --- | --- |
| Transformed values | Thick layers of till (Ktkmo_a) | 0.4093 | 0.6126 | 0.7016 | 0.8195 | 0.9089 |
|  | Landslide soil (Kskred_a) | 0.5510 | 0.7212 | 0.7948 | 0.8520 | 0.9656 |
|  | Marine deposits (Kmar_a) | 0.5186 | 0.6859 | 0.7581 | 0.8538 | 0.9261 |
|  | Lacustrine deposits (Klac_a) | 0.5499 | 0.7479 | 0.8336 | 0.9469 | — |
|  | Glaciofluvial deposits (Kelv_a) | 0.5435 | 0.7029 | 0.7717 | 0.8626 | 0.9314 |
|  | Exposed bedrock (Kbf_a) | 0.0181 | 0.0870 | 0.1662 | 0.3679 | 0.6258 |

### ORDINATION OF LANDSCAPE PROPERTY MATRICES FOR IDENTIFICATION OF COMPLEX LANDSCAPE GRADIENTS

#### Ordination methods

Ordination methods were used to summarise the main gradients of coordinated variation in landscape properties among the observation units, as characterised by the 85 analytic variables. Ordination analyses were performed by a data- and result-driven procedure, starting with the total data set (3966 OUs), and followed by analyses of subsets. The splitting into subsets was based upon interpretation of the results of previous ordination analyses in terms of tentative major-type groups, major types and groups of OUs of special interest.

Ordination methods sort observation units (OUs) so that observation units with similar characteristics (i.e. similar values for the analytic variables) are placed near each other in the conceptual geometric space spanned by the ordination axes and observation units which differ in many respects are placed further apart (Økland 1990, Legendre & Legendre 2012). Moreover, ordination methods sort OUs along axes of decreasing importance, i.e. in order of decreasing explained variation (in the variables). Note that ‘explained variation’ is used here in a strict statistical sense, i.e. with respect to a statistical measure of variation that may or may not be ecologically meaningful (cf. Økland 1999).

All data sets that were subjected to ordination analysis were, in principle, analysed in the same way, by parallel ordination (van Son & Halvorsen 2014), using the methods DCA (*detrended correspondence analysis*; Hill 1979, Hill & Gauch 1980) and GNMDS (*global nonmetric multidimensional scaling*; Kruskal 1964a, 1964b, Minchin 1989).

DCA was performed by use of standard parameters in the function decorana in the R-package *vegan* version 2.4-0 (Oksanen et al. 2016). The DCA axes were characterised by their ‘eigenvalue’ and gradient length, expressed as S.D. units (*standard deviation units*).

GNMDS was performed by use of the following *vegan* parameter settings: vegdist with dissimilarity measure = proportional dissimilarity (PD = ‘Bray-Curtis dissimilarity’; Czekanowski 1909; Bray & Curtis 1957; Økland 1990); isomapdist with epsilon = 0.5 [changes the dissimilarity matrix to a matrix with geodetic distances (Mahecha et al. 2007) by replacing all PD-values > 0.5 with the sum of reliable PD-values for ‘shortest distance’ between the compared OUs (Swan 1970; Williamson 1978; De’ath 1999; Bouttier et al. 2003)]; monoMDS with maximum number of iterations (maxit) = 1000, convergence ratio for stress reduction (smin) = 1·10^−7^ and stopping criterion for lowest stress-value (sfgrmin) = 1·10^−7^. A minimum of 100 GNMDS solutions, all from random start configurations, were obtained. The GNMDS solution with lowest stress was compared with the solution with second-most lowest stress by a Procrustes test (Oksanen et al. 2016), by which the two ordinations are compared after origo-translation, mirroring, and linear rescaling of axes. A correlation coefficient (the Procrustres sum of squares) was used as a measure of the degree of correspondence between the compared ordinations. A permutation test (protest) was used to test the hypothesis: ‘the degree of correspondence between the two ordinations is not better than between two random point configurations’. The test was conducted by use of 999 permutations. A significant test (rejection of the hypothesis at the α = 0.001 level) was taken as a necessary, but not sufficient, condition for ordination results to be potentially meaningful. The graphic representation of the results of Procrustes analysis, by which the origo-translated, mirror-imaged and linearly rescaled axes of one of the compared ordinations was plotted in the first ordination (the plot(procrustes)command) was used to identify OUs with unstable location (i.e. very different positions in the two ordinations). The fraction of OUs with unstable location was used together with the procrustres sum of squares to judge the stability and reliability of ordinations.

The solution with lowest stress was accepted as the best solution when, based on the results of the Procrustes test and the number of OUs with unstable location, it was considered as essentially equal to the ordination with the second lowest stress. In the opposite case, new GNMDS ordinations were obtained, based upon new random initial configurations, until either (1) two similar, lowest-stress solutions were found and the best was accepted, or (2) no best solution was obtained from 1000 random initial configurations.

The best (lowest stress) solution was subjected to varimax rotation by PCA and rescaled to half-change (H.C.) units using the postMDS function of *vegan*. Varimax rotation translates the ordination space so that GNMDS axis 1 goes through the longitudinal axis of the cloud of OUs in the ordination space (i.e. captures the maximum possible amount of variation in the ordination space, expressed as sum of squared OU scores), axis 2 is perpendicular to axis 1 and passes through the longest axis that is orthogonal to the first axis, etc. Rescaling to H.C. units scales the axes in units that express landscape property compositional turnover. One H.C. unit corresponds to an average difference in landscape property composition of 0.5 units on the PD scale.

For each analysed data set or subset, GNMDS ordination was first performed with number of dimensions k = 4 (i.e. to produce a four-dimensional GNMDS solution). The best 4-dimensional solution was then compared to the DCA solution (DCA always finds four axes) by calculating pair-wise Kendall’s nonparametric correlation coefficients τ between all pairs of axes. If all GNMDS axes had paired correlations with at least one DCA axis greater than τ = 0.4, we considered the 4-dimensional GNMDS ordination to be confirmed by the DCA ordination. In such cases, with 4 confirmed ordination axes, the 4-dimensional GNMDS ordination was taken as basis for further interpretation. If the 4-dimensional GNMDS ordination was not confirmed by DCA, the entire process was repeated for k = 3 and, unless the three axes of the 3-dimensional GNMDS ordination were confirmed by 3 or 4 DCA axes, the process was repeated once more and the 2-dimensional GNMDS ordination compared to DCA. For all analysed data sets, both axes in the 2-dimensional GNMDS ordination were confirmed by DCA.

In the few cases where the GNMDS ordination process had to be terminated without a confirmed ordination result, the DCA ordination was used as the basis for further interpretation with great caution. The purpose of parallel ordination is to avoid subjecting to interpretation ordination results that have high probability of being burdened with mathematical artefacts. Such artefacts are not likely to be present when two very different methods, such as DCA and GNMDS, give closely similar results (Økland 1996, van Son & Halvorsen 2014). The few cases without a confirmed GNMDS ordination result were always obtained for large data sets (i.e. with many OUs). In these cases, all ordinations had about the same stress value, *ca.* 0.27, and the Procrustes figure (Fig. 5) revealed that the ordination results were unrelated. In these cases, the OUs were randomly distributed along the ordination axes within a sphere-shaped point cloud (exemplified by Fig. 6). Such a result, which, as far as we know, has not previously been reported in the literature, probably represents a case where the relationships in the data matrix are so complex that the GNMDS iteration procedure fails ‘to break out of’ the initial configuration.

**Figs. 5–6.**
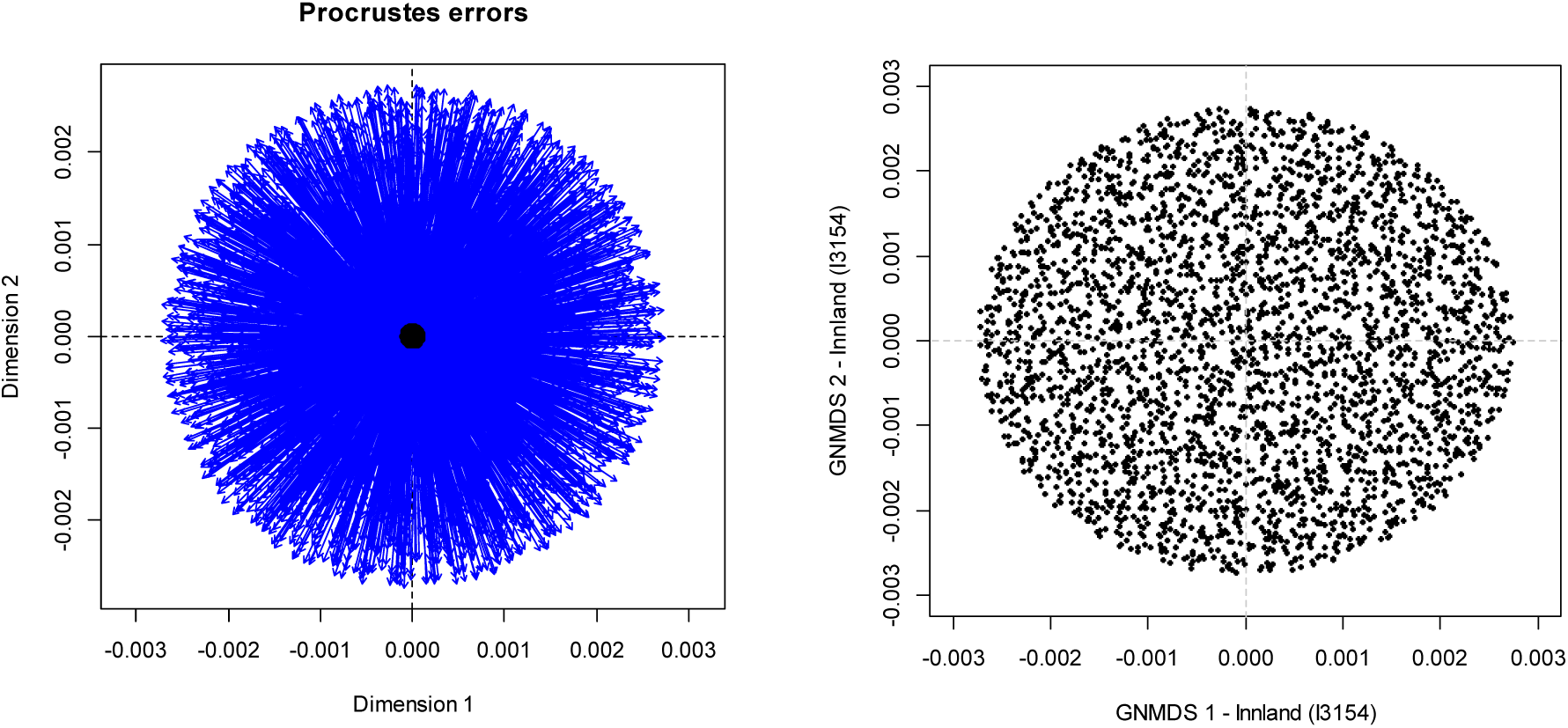
Procrustes figures. (5) Comparison between two ordinations, by which blue lines connect positions of OUs in the second ordination to positions of the same OU in the first ordination, for an unsuccessful GNMDS ordination of a large dataset; (6) Unsuccessful GNMDS ordination of a large dataset with the OUs randomly distributed in a sphere-shaped point cloud.

R version 3.2.2 (R Core Team 2018) with the package *vegan*, version 2.4-1 (Oksanen et al. 2016) was used for all statistical analyses.

#### Interpretation of ordination results

Ordinations summarise correlations between the analytic variables and, at the same time, reveal relationships between observation units (OUs). Because the variables in this case express landscape properties and because identification of gradients in landscape properties is the purpose of the analyses, ordination results are, strictly speaking, not in need of being interpreted using extraneous data (i.e. data other than those used to obtain the ordination results). In this respect, the ordination analysis of landscape element compositional data in this study differs from ordinations of vegetation data which require interpretation by use of external environmental information.

Nevertheless, all ordination results obtained in this study were subjected to a formalised interpretation, the main purpose of which being to gain a best possible understanding of the complex, co-ordinated patterns of variation in the landscape data sets as a basis for identifying complex landscape gradients (CLGs). Furthermore, the interpretation assisted division of the CLGs into standard segments and the building of a system of landscape types.

Standard tools were used for the interpretation of ordinations. For all confirmed ordination axes, non-parametric correlations (Kendall’s τ) between the axis and each individual key variable and analytic variable were calculated. Because most landscape data sets are large in terms of the number of OUs they contain, most correlation coefficients are significantly different from 0. Furthermore, *p* values for the correlation test of independence between variables are burdened with uncertainty due to the nested sampling design (and associated spatial autocorrelation).We therefore report correlation coefficients only, as indices of the strength of relationships between variables. The following terms were used to characterise correlations:

|τ| > 0.6 : very strong
0.5 < |τ| < 0.6 : strong
0.4 < |τ| < 0.5 : quite strong
0.3 < |τ| < 0.4 : noticeable

Noticeable and stronger correlations (|τ| > 0.3) are marked with colours on a scale from light orange to dark brown in correlation tables. Furthermore, the term ‘correlated with’ is used throughout the text to mean |τ| > 0.3 (the alternative, |τ| < 0.3, is referred to as ‘weakly correlated’).

Several graphic tools were used in the interpretation. Vectors pointing in direction of the largest increase in variable value were identified for key variables and selected analytic variables using the envfit function in *vegan*. For factor variables (e.g. soil types), the envfit function returns the centroid for the location of OUs belonging to each class. As a measure of the strength of the relationship between each vector and OU locations in the ordination diagram, the fraction of variation in the ordination diagram explained by the vector, r², was calculated. Furthermore, ordination diagrams were provided with isolines for selected variables, calculated by use of the ordisurf function in *vegan*. Symbols of different shapes and colours for different landscape types and landscape gradient segments were used in the ordination diagrams in order to illustrate the variation in landscape properties.

#### Identification of complex landscape gradients

Interpreted ordination results were used to identify *candidate* major landscape types as maximally well-defined groups of OUs in ordination diagrams. As for the ecosystem type level in NiN (Halvorsen et al. 2016), the candidate types had to satisfy additional criteria, e.g. with respect to possession of important complex landscape gradients (CLGs), to be accepted as major types.

#### Identification of CLG-candidates

Candidates for complex landscape gradients (CLGs) were identified, separately for each major type, based on ordination analyses of data subsets consisting of all OUs tentatively affiliated with the major type in question. The CLG candidates thus identified were major-type and data-set specific CLGs, which were subsequently divided into standard segments [i.e. intervals of a complex-gradient, each comprising at least 1 ecodiversity distance unit (EDU) of variation in the key characteristic within the major type in question; cf. Halvorsen et al. (2020)].

Groups of characterising variables (KVs; collective term for primary key and analysis variables) were considered primary CLG candidates when containing at least one KV that was at least quite strongly correlated (|τ| > 0.4) with at least one confirmed ordination axis. This is referred to as the basic criterion (1) for recognition of an CLG. For groups of KV’s that satisfied this basic criterion, the CLG candidate included all KV’s that (i) belonged to one and the same functional variable subcategory; (ii) had approximately the same pattern of correlations with the ordination axis (or axes); and (iii) were quite strongly correlated (|τ| > 0.4) with this or these. A CLG candidate thus consisted of a group of primary key and/or analytic variables. For CLG candidates that, according to criterion (iii) for inclusion of variables, would include 8 or more variables with |τ| > 0.5 with the ordination axis (or axes), the stronger demand (iv) on included variables, that only variables with |τ| > 0.5 with the axis was included, replaced (iii). This reduced bias towards subcategories represented by many variables (such as infrastructure).

In order to ensure that functional subcategories could be identified as CLG candidates even if they were represented by a small number of KVs, a relaxed criterion (2) replaced criterion (1) for subcategories represented by 3 or fewer variables. According to criterion (2), a sufficient condition for being considered as a CLG candidate was that the variable had a ‘noticeable’ correlation (|τ| > 0.3) with the axis (or axes). For CLG candidates identified by use of (2), variables were included if they satisfied criteria (i) and (ii), while (iii) was replaced by (v), that the CLG candidate should include all variables with a noticeable correlation (|τ| > 0.3) with the axis (or axes) in question.

All CLG candidates identified by the criteria (1) or (2) were subjected to a judgement of homogeneity to ensure that they consisted of variables that, in addition to representing one and the same functional subcategory, made up a group of sufficiently strongly correlated variables. If not, the CLG candidate was split into two or more CLG candidates, each consisting of one or more variables. Splitting into two groups was enacted when the characterising variables could be divided into subgroups in such a way that no pair of variables, one variable from each subgroup, was pairwise strongly correlated (|τ| > 0.4) with each other. The corollary of this splitting criterion is that variables that were quite strongly correlated [noticably correlated in the case of criterion (2)] with at least one other variable from the same subcategory constitute one CLG candidate. Furthermore, variables that were not correlated with at least one other variable from the same subcategory made up a CLG candidate on their own.

For each CLG candidate so identified, the analytic and key variables were sorted from highest to lowest |τ| value with the ordination axis (or axes) in question.

#### Identification of parsimonious complex variable groups and calculation of explained variation for each CLG candidate

The requirement for CLG candidates to be represented by groups of correlated variables (KV groups) necessarily implied that the variation explained (in statistical sense) by the variables in a group overlap extensively. In order to obtain a realistic estimate of the relative amount of the variation in a data set that was explained (still in statistical sense) by a CLG candidate, useful as a measure of the relative importance of the CLG candidate (as a source of variation at the landscape type level), we obtained for each CLG candidate a parsimonious KV-group (PKV group) consisting of the KV variables that provided a significant and independent contribution to explaining variation in the analytic data set in question. The variable selection process used to obtain the parsimonious KV-group is explained below.

The PKV group, and estimates for the variation it explained, were obtained using constrained ordination by the Redundancy Analysis (RDA) method (Rao 1964, ter Braak 1985). RDA is the constrained variant of Principal Component Analysis (PCA), an ordination method that finds axes which successfully maximises the explained sum of squares (‘linear variation’) in a data set. In constrained ordination, the axes are constrained to express the variation *also explained* by a set of constraining variables (Økland 1996). For a PKV group with *s* variables, a maximum number of *s* constrained ordination axes can be extracted. These are ordered by decreasing variation explained, as expressed by the eigenvalues of the axes. The sum of the variation explained by each of the *s* constrained ordination axes was used as an estimate for the total variation explained by a CLG candidate, represented by its PKV group.

For each CLG candidate in each major type, the KV group was reduced to a PKV group by forward selection of variables (e.g. Blanchet et al. 2008). Forward selection is an iteration process that starts with calculating, for each of the *m* variables in the group, how much of the variation in landscape-element composition the variable explains. The variation explained (VE) by a variable was found by an RDA analysis with the variable in question as the only constraining variable, as VE_i_ = λ_1_/TI, where λ_1_ is the eigenvalue of the constrained ordination axis and TI is total inertia, that is, the sum of the eigenvalues of all PCA axes that can be extracted for the data set in question. TI is the total sum of squares for all analytic variables [after centering (to mean = 0) and standardisation (to SD = 1)]. For PCA with centred and standardised variables, TI = *m*, the number of analytic variables in the data set. For simplicity, all variations explained were standardised to TI = 1. After the variation explained by each KV had been found, the KV with the highest VE was incorporated in the PKV group. In the second round of the iteration procedure, the analysis was repeated with the remaining *m* – 1 KVs as constraining variables and the KV selected in the first iteration cycle as ‘conditioning variable’ in *m* – 1 RDA analyses. In each of these RDA analyses, the method first removed the variation explained by the chosen KV and then calculated how much of the residual variation (the variation remaining after the variation explained by the selected KV is accounted for) that was explained by each of the remaining variables. After all variables had been subjected to RDA analyses, the variable which explained the largest proportion of the residual variation was selected. In principle, the iteration process can continue until there are no variables left or until no variation in the data set remains unexplained. In practice, however, the process is stopped when none of the remaining variables provides a sufficiently large independent contribution to explain the residual variation. To avoid inclusion of variables that are highly correlated with variables already included in the PKV group, we adopted the default practice in forward selection, to set a lower threshold value for the (residual) explained variation that a variable must explain to be included. This limit can, in principle, be determined by use of statistical tests. However, given the large data sets analysed in this study (i.e. with many OUs), the null hypothesis that the explained variation is not significantly higher than expected of a random variable, will be rejected with low *p* values in almost all cases. In addition, testing is made difficult by the fact that we do not distinguish between constraining variables and analytic variables (the latter is contained in the data set subjected to RDA and therefore contributes 1/*m* of TI). We instead used as a lower limit ω_0_ = *g* · λ_0_ where *g* is a constant and λ_0_ is the expected variation explained by a random variable, expressed as a fraction of the residual variation. In round 1 of the iteration process, i.e. before any variable has been included in the PKV group and the residual variation (before the round) equals the total inertia, ω_0_ was set equal to:

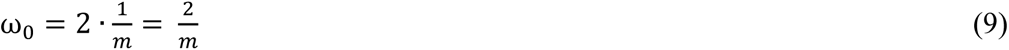

The residual variation after iteration cycle *j*, when *j* variables have been included in the PKV group and the variation explained so far (the sum of the eigenvalues of the *j* constrained axes) is λ*_j_*, is 1 – λ*_j_*. In cycle *j* + 1, the threshold value is given by the equation:

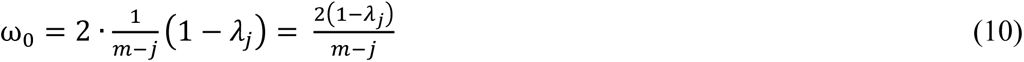

The parameter value 2 was selected after careful assessment of the results for the first analysed data sets. For data sets with fewer than 200 OUs, with a more prominent stochastic component of variation, the threshold value 2.5 was used.

The variable selection process was stopped when no more KVs met the threshold criterion. The CLG candidate was represented by its PKV group (consisting of *s* variables) in subsequent analyses. The variation explained by the CLG candidate was obtained as λ_s_, the sum of the eigenvalues of the *s* constrained ordination axes extracted with the PKV group as conditioning variables. The first axis of the RDA ordination of the PKV group is the linear combination of the *s* variables in the PKV group which explained the maximum amount of variation in the data set. This axis is termed the CLG candidate’s primary key variable.

#### Forward selection among CLG candidates for a set of orthogonal CLGs

An important lesson from the ‘pilot Nordland project’ (Erikstad et al. 2015) was that the large number of landscape gradients that were considered to be important for each major type, and hence used to define minor types by the standard NiN procedure (see Halvorsen et al. 2020: Appendix S2), resulted in an unmanageably high number of minor types, unsuited for practical descriptive purposes. The practical solution chosen for the ‘pilot Nordland project’ was to aggregate minor types subjectively into types at a level between minor and major types. This procedure did, however, not comply with the standards of the EcoSyst framework (of which NiN version 2 shall be an implementation; Halvorsen et al. 2020). Therefore, in order to construct a system of minor types at the landscape level that may be theoretically well founded and practically useful, the principle adopted at the ecosystem level in NiN version 2 of major type-independent local complex-gradient concepts was not adopted at the landscape level. Instead, the set of *t* accepted CLG candidates was reduced to a set of *u* orthogonal (i.e. independent) CLG’s. Forward selection (as described above) was used, but with CLG candidates (each represented by its parsimonious KV group) as units in the selection process instead of individual KVs. CLG selection was undertaken successively for the three functional variable categories, first the basic geo-ecological variables (category G) which describe the fundamental conditions for soil and vegetation; thereafter bio-ecological variation (category U), which describe basic conditions for human exploitation of the landscape; and, finally, land-use variables (category A).

In the first phase of this selection process, the CLG candidate in category G with the largest amount of explained variation λ_s_ was chosen. Then, for each of the other CLG candidates in the G category (represented by their PKV group), RDA was used to calculate how much of the variation in the data set that was explained *in addition to* the variation already explained by the first chosen CLG. After a careful assessment of preliminary results, 6% (0.06) variation (in the data set) explained was chosen as a lower threshold value for inclusion of a CLG candidate in the group of orthogonal CLGs. For data sets with fewer than 200 OUs, with a more prominent stochastic component of variation, the value 0.08 was used. When no more CLG candidates met the 0.06/0.08 criterion, the selection process was stopped. Thereafter, the procedure was repeated for CLG candidates in the bio-ecological U category, with all selected CLGs in the G category, represented by their PKVs, as conditioning variables. The 0.06/0.08 criterion for inclusion of additional CLGs was applied. Finally, the procedure was repeated for CLG candidates in the A category, with all selected CLGs in the G and U categories, represented by their PKVs, as conditioning variables.

#### Orthogonal primary key variables for each CLG

The set of *u* CLGs found by forward selection of CLG candidates represents a parsimonious set of orthogonal CLGs for the major type. Since each CLG is represented by a set of *s* = 1–4 variables, these orthogonal CLGs constitute a series of orthogonal subspaces of dimension *s* within the data-dimensional space. CLGs were operationalised to one stepwise gradient each by reduction of each of these subspaces to one dimension. In line with the procedure described above, RDA was used to find the vector in data-dimensional space that captured the maximum possible variation in the *s*-dimensional space spanned by the CLG. This vector, referred to as the orthogonal key variable (ONV) for the CLG, was given by the first RDA axis in the constrained ordination of the PKV group of the CLG, with all variables included in all PKV-groups of all previously identified CLGs as conditioning variables.

The ONVs were extracted as actual RDA score vectors from the respective constrained ordinations by the vegan function scores using the following parameter options: display = “lc” [extracts scores that are linear combinations of the constraining variable (‘*linear combinations* = *“lc scores”’*), while the alternative ‘*weighted average scores = “wa scores”* extracts scores after one iteration cycle towards the first PCA axis for the data set in question]; scaling = 1 (provides *correlation biplot scaling* of scores, which optimises the relationship between the cosine of the angle between the analytic variable vectors in ordination space and their Pearson’s product-moment correlation coefficient).

After extraction, the ONVs were ranged onto a standard 0–1 scale by the procedure used to transform analytic variables. The term ONV is used also for these ranged ‘primary ONVs’.

#### Estimation of landscape distance

The requirements for a standardised division of complex landscape gradients (CLGs), which is a prerequisite for a testable division into landscape types, were the same as the requirements for a standardised division of local complex environmental variables at the NiN (and EcoSyst) ecosystem level (Halvorsen et al. 2016: chapter B2; Halvorsen et al. 2020: Appendix S2): (1) An explicitly defined unit for measuring the difference in landscape-element composition between units along a CLG; (2) a method for quantifying this difference; (3) a method for calculating the magnitude of landscape variation along a CLG (the gradient length); and (4) a method for dividing a CLG into standard segments. Definitions and methodologies for standardised division of LECs (see Halvorsen et al. 2016: chapters B2 and Appendices 2–5; Halvorsen et al. 2020: Appendix S2) are used as templates for the corresponding definitions and methods at the landscape level. Here, therefore, only differences relative to the corresponding methodology for the ecosystem level are explained.

The parallel to the species as key characteristic at the ecosystem level is landscape elements (defined as ‘natural or human-induced objects or characteristics, including spatial units assigned to types at an ecodiversity level lower than the landscape level, which can be identified and observed on a spatial scale relevant for the landscape level of ecodiversity’). The parallel to the species composition, which is used to quantify differences between ecosystem-type candidates, is the landscape-element composition, defined as ‘the landscape elements occurring within a specific area, quantified by an appropriate performance measure’. In this study, the landscape-element composition of each OU was represented by the 85 analytic variables, quantified by use of the transformed and ranged performance scale.

In EcoSyst, the unit of compositional turnover of the key characteristic at an ecodiversity level along a complex variable in the key source of variation at this level, is termed the ‘ecodiversity distance unit’ (EDU; Halvorsen et al. 2020). Adapted to the landscape level (landscape-element composition along a CLG), the unit is referred to as the ecodiversity distance unit in landscapes (EDU–L).

Two fundamentally different approaches to estimation of differences in landscape-element composition between two landscape-type candidates are available: (1) indirectly, based upon the candidates’ positions along confirmed DCA or GNMDS ordination axes (Halvorsen et al. 2016: Appendix 2g); and (2) directly, based upon a measure of dissimilarity calculated between the units that are compared (Halvorsen et al. 2016: Appendix 2h). The arguments for choosing alternative (2) for the ecosystem level are directly transferable to the landscape level (see Halvorsen et al. 2016: Appendix 3). Furthermore, the much shorter gradient lengths of the DCA and GNMDS ordinations of landscape-type OUs (all ordination axes: < 2.5 S.D. units in DCA and < 2 H.C. units in GNMDS) than between ecosystem types (Halvorsen et al. 2015) favours (2) because the tendency for non-linearity of the proportional dissimilarity (PD; see Halvorsen et al. 2016: Appendix 3h) measure decreases with decreasing gradient length. PD was therefore used to quantify the variation in landscape-element composition also at the landscape-type level.

The so-called ‘generalisation challenge’ in calculating differences in species composition is also present when differences in landscape-element composition are quantified. This challenge implies that the expected inequality in landscape-element composition between OUs located at the same position along a CLG differs from zero because of random variation (‘noise’) in landscape-element composition. Accordingly, measurements of compositional differences between OUs cannot be added and subtracted unless the ‘inequality between repeats’, or ‘internal association’, i.e. the compositional difference between OUs placed at the same point along the underlying CLG (INA; Whittaker 1952, 1960, Økland 1990), can be quantified. At the ecosystem level, the generalisation challenge is resolved by using generalised species-list data to calculate ecodiversity distance units in ecosystems (EDU–E; Halvorsen et al. 2020). Such data contain ‘smooth’ (‘noise-free’) species abundance data for each candidate ecosystem type, derived from models of species’ abundance distributions along local complex environmental gradients. These models are obtained by consensus among experts, following a standard procedure which makes use of all available data as well as informal knowledge (see Halvorsen et al. 2016: chapter B2c). Our knowledge about variation at the landscape-type level is, however, so fragmentary that the method for using generalised data sets assembled by expert judgement was inapplicable.

Instead, we chose an alternative, five-step procedure for estimation of ecodiversity distance in landscapes:

1. Division of CLG’s ONV into segments (ONV segments) into which the OUs are filed.
2. Calculation of landscape element profiles (LEPs) for each ONV segment. The landscape element profiles are the parallels to the generalised species lists at the ecosystem level.
3. Calculation of proportional dissimilarity (PD) between the LEPs.
4. Estimation of the ‘inequality between repeats’ (INA) from the PD values and calculation of landscape distances (LD) between ONV segments.
5. Estimation of the gradient length of the CLG, in EDU–L units.

(1) *Segmentation of CLGs by their ONV.* Ideally, we intend (i) to divide each ONV into segments that are as small as possible, in order to be able to describe the variation in landscape element composition along the ONV in the greatest possible detail; (ii) to make all ONV segments equally wide (i.e. span similar landscape-element compositional turnover along the CLG) and thus to make them ideal minor-type candidates; and (iii) that each ONV segment is represented by as many OUs as possible to reduce random variation as much as possible and, at the same time, increase the degree of generalisation. Conditions (i) and (iii) are incompatible, calling for a best possible compromise. Furthermore, condition (ii) is incompatible with both of (i) and (iii) because the distribution of OUs along most ONVs is unimodal (more or less skewed). Accordingly, equal segment width (measured on the 0–1 scale of the ranged ONV) typically implies that segments near CLG ends contain very few OUs.

Based on the above considerations, the following criteria were used to divide ONVs into segments: (i) No segment should contain less than *n* = 25 OUs. (ii) The number of intervals *q* = 8. (iii) Each segment should have the same width, measured in the units by which the ONV is scaled. The criteria are arranged hierarchically, so that criterion (i) has priority over criterion (ii) which, in turn, has priority over criterion (iii). Application of the criteria can be exemplified by the affiliation of the 118 OUs in major-type candidate IF (inland sediment plains) to segments. Regardless of which CLG is addressed, criterion (i) sets an upper limit of four (*q* = 4) for the number of segments, criterion (i) overruling criterion (ii). While criterium (iii) suggests that the four segments comprise OUs in the ranges 0–0.25, 0.25–0.5, 0.5–0.75 and 0.75–1 ONV units, criterion (iii) may be overruled by criterion (i), implying that segments near ONV endpoints comprise the 25 OUs with the lowest, respective highest, ONV scores. Thereafter, the remaining OUs are distributed on two segments of equal width (criterion iii) if not again overruled by criterion (i) which demands a minimum of 25 OUs in each segment.

(2) *Calculation of LEPs for each ONV segment*. For each of the *q* ONV segments, the mean value of each (zero-skewed transformed and ranged) analytic variable was calculated. The vector of mean values for each analytic variable in the set of OUs that represent the ONV segment, the ‘centre of gravity-vector’, was used as landscape element profile.

(3) *Calculation of PD between the LEPs*. The PD (proportional dissimilarity index) is the one among the dissimilarity indices in common use, e.g. in vegetation ecology, which is best suited for calculating ecodiversity distance in ecosystems (Økland 1986). This holds true also for other situations in which the response variables (here: the analytic variables) are in general linearly or unimodally related to the underlying complex gradients (here: CLGs). In its simplest form, calculated for pairs of ONV intervals *j* and *k*, PD is given by the formula

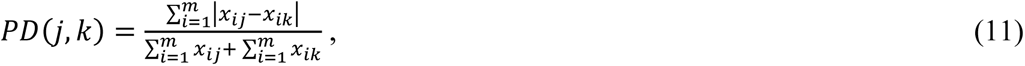

where *x_ij_* is the value of variable *i* in ONV segment *j*, *x_ik_* is the value of variable *i* in ONV segment *k*, and *m* is the total number of variables (i.e. the total number of variables with variation in the major-type candidate in question). The PD index is bounded below by 0 and above by 1; the value 0 indicates that the landscape-element composition of two ONV segments is identical (i.e. that each variable has the same mean value in segments *j* and *k*), the value 1 represents the situation (which in landscape contexts is completely unrealistic) that one of the two compared LEPs has the value 0 while the other has a positive value for each variable.

PD was calculated between all pairs of ONV segments *j* and *k* from 1 to *q* = 8, *j* ≠ *k*.

(4) Estimation of ‘inequality between repeats’ (INA) based on estimated PD values and calculation of landscape distances (LD) between ONV segments. PD values between completely generalised vectors (without random variation) of variables that vary linearly along a gradient, are additive (Økland 1986). In mathematical terms, this means that PD differences between three ONV segments, represented by centres of gravity A, B and C, where A < B < C along the gradient, satisfy the equation:

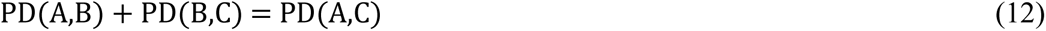

If the LEP profiles satisfied demands for complete generalisation and additivity, PD could be used directly as a measure of landscape distance between centres of gravity of ONVs, after linear rescaling of PD values to EDU-L units (i.e. by multiplication with a constant *w*). Inspection of landscape-element compositional data for all ONVs in all major-type candidates (see Results) did, however, show that PD(A,C) > PD(A,B) + PD(B,C) almost throughout and, thus, that the requirement for (complete) generalisation is never fulfilled. Moreover, the results show that the linearity requirement is fulfilled to varying degrees. Extreme cases by which PD(A,C) < PD(A,B) + PD(B,C), that is, that the similarity in landscape element composition between A and C is greater than between A and B or between B and C, also exist. This calls for correction of raw PD values for INA, which was accomplished in accordance with the following line of reasoning:

Let LD(A,C) denote the underlying, true landscape distance (after correction for INA) between centres of gravity of segments A and C. It follows from the definition of landscape distance EDU-L that

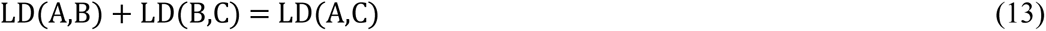

Inadequate generalisation, however, means that all PD values can be expressed as a sum of LD and INA, so that:

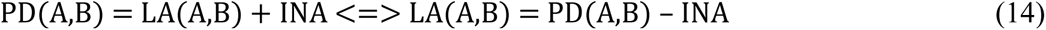

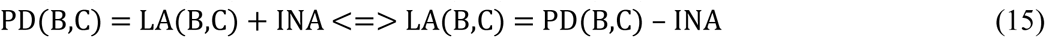

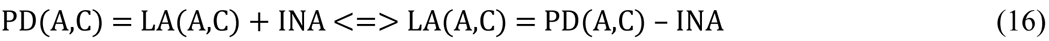

Inserting (14–16) into (13), provides an estimate for INA:

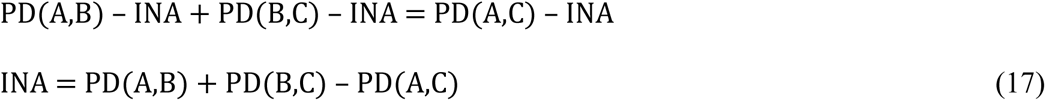

Equation (17) can be used for all combinations of A, B and C in the data set. When *q* = 8, the number of unique combinations of three segments with centres of gravity A, B and C is 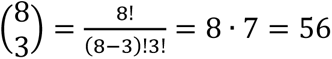. Accordingly, 56 single estimates for INA are provided. Three principally different methods for obtaining one estimate of INA (and for correcting the matrix of between-segment PD values) from these 56 single INA estimates were evaluated.

(i) *Method I (global INA correction; assuming linear variation and constant INA along the gradient)*. If the magnitude of stochastic variation in variable values does not vary along the CLG (as represented by the ONV) and all variables vary linearly along this gradient, *the mean value of the 56 single INA estimates* is the obvious choice of estimator for INA because it is based on the largest possible number of observations.

However, preliminary analyses (e.g. of the coastal plains data set) showed that the assumption of constant INA is untenable for most CLGs and most major-type candidates. The main reason for this (which applies to all data sets to a greater or lesser extent) is that segments near CLG ends are represented by fewer OUs than segments near the middle of the gradient and, accordingly, that the magnitude of stochastic variation (and INA) is inversely related to the number of OUs in each segment. If we assume constant INA along the entire CLG, the pairwise distances between the centre of gravity of ONV segments near the centre of the gradient will be ‘overcorrected’ while distances near the gradient endpoints are ‘undercorrected’. In extreme cases, LD estimates between central ONV segments become very small (and often negative) while LD estimates between endpoint segments become disproportionately large. The result of LD calculations by Method I are reported, but not used in the subsequent analyses.

(ii) *Method II (local INA correction; assuming linear variation, but allowing for variation in INA along the gradient)*. Given that INA varies along the gradient, single INA estimates have to be used to provide separate estimates of INA for different parts of the gradient. Let us assume that the gradient has been divided into 8 segments with points of gravity ABCDEFGH, each represented by an LEP. Equation (17) provides two INA estimates based on PD values for relationships between B and C, based upon relationships with the two adjacent segments, respectively, AB and CD:

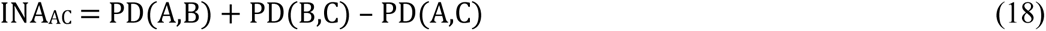

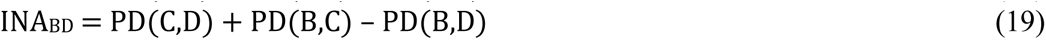

Method II uses the average of INA_AC_ and INA_BD_ as an estimate of INA in the BC range, and finds LA(B,C) directly from the equation

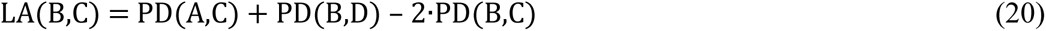

The local INA estimate is then:

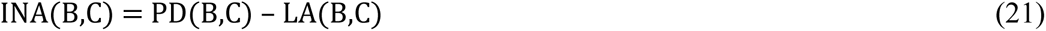

For the outermost ranges, only one INA estimate is available, e.g. for INA_AB_ given by

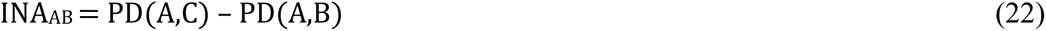

Results of analyses of landscape element profile data sets have revealed some cases of very large differences between the INA and landscape distance estimates based on Methods I and II. Thorough review of some of these cases (e.g. CLG KA for data set IX399) has shown that LD estimates can be directly misleading due to nonlinear variation along the gradient (see above for explanation and below for review of the example. This motivated for a third method, which was used for all calculations of landscape distance.

(iii) *Method III (local INA correction; assuming nonlinear variation along the gradient)*. This method estimates LA(*x*, *y*) as the mean difference in PD for PD (*z*, *x*) and PD (*z*, *y*) over all *z* ≠ *x*, *y*, where *x* and *y* are the segments to be compared. Thus, for *q* = 8, we obtain:

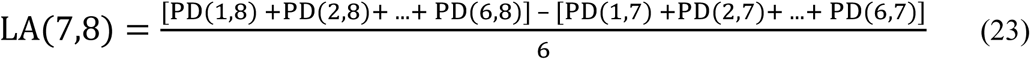

Method III was first actualised by analysis of CLG 1 (distance to the coast) for data set IX399 (other inland plains), which contained parts with strongly nonlinear variation in landscape-element composition. This resulted in large differences between gradient-length estimates (sums of LDs for all intervals between the CLG endpoints; see below) by the three methods: I: 0.0655; II: 0.5698; and III: 0.2959. Method I provided an estimate of INA of 0.1647 ± 0.0680 (mean ± S.D.), which resulted in negative LD values between ONV segments 3 and 4, 4 and 5, and 5 and 6. Method II clearly overestimated gradient length. Inspection of the PD matrix (e.g. Fig. 7) reveals the reason for the failure of Method II: that the landscape element composition of segment 1 is more similar to the composition of segments 6–8 than to segment 5. Similarly, segment 8 is more similar to segments 3–5 than to segment 6. The effect of nonlinear variation along the gradient is corrected for by Method III, which for this gradient yields gradient length estimates intermediate between those provided by the two other methods.

**Fig. 7.**
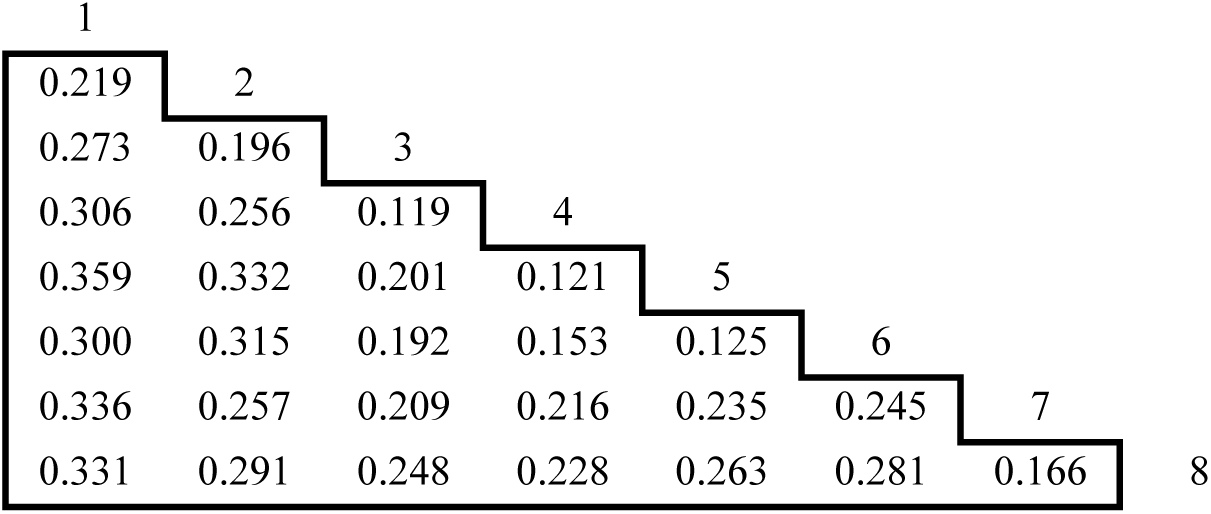
Uncorrected matrix of PD dissimilarities between 8 LEPs for CLG KA (Distance to coast) for major-type candidate ‘Other inland plains’ (IX) based on analysis of data set IX399.

(5) *Estimation of total landscape gradient length*. Total landscape-gradient length (LGL), on the corrected PD inequality scale, was calculated by summing up LD values for all intervals 12, 23, 34, 45, 56, 67 and 78 into which the ONV was divided. Additionally, based on the same reasoning as for the ecosystem level (Halvorsen et al. 2016, 2020), we assume that the LEPs represent the centre of gravity in each interval (centre of the interval), i.e. that the underlying gradient extends beyond the centre of gravity of the intervals 1 and 8 by half the LD to the centres of gravity of the neighbouring segments. Accordingly, the LGL was estimated by the equation:

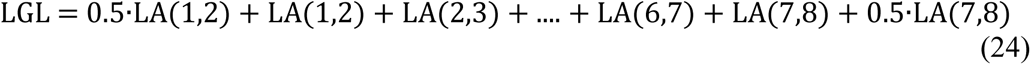

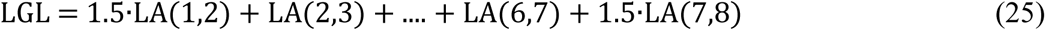

The LD and LGL estimates are given in PD units. In order to divide CLGs into standard segments, comparable among CLGs, LDs and LGLs in PD units were converted to ecodiversity distance units in landscapes (EDU–L units). Just like for the corresponding EDU–E unit for ecosystems, there is no *a priori* theoretical reason for choosing a specific definition of 1 EDU–L. The amount of compositional turnover corresponding to 1 EDU does, however, have important practical implications since the number of segments into which each CLG is divided determines the number of minor types. For example, halving the EDU results in a 16-fold increase in the number of minor types in a major type with 4 important CLGs. Based on the experience from analysis of all landscape data sets, we pragmatically defined 1 EDU–L unit = 0.08 PD units. LD and LGL results are, in most cases, reported using the original measurement scale in PD units.

#### Rescaling of ONVs and standardised division into segments

Each ONV, representing one CLG in one given major-type candidate, was rescaled by use of the results of LEP dissimilarity analyses according to the following procedure:

1. A new measurement scale from 0 to LGL was established for each ONV by first placing the centre of gravity of ONV segment 1, TP1, at 0.5 · LD(1,2). Then the centre of gravity TP2 of ONV segment 2 was placed at 0.5 · LD(1,2) + LD(1,2) = 1.5 · LD(1,2), TP3 at 1.5 · LD(1,2) + LD(2,3) etc.
2. Borders between the intervals, GR12, GR23 etc., on the new measurement scale were then estimated by ‘smoothing’ to further reduce the effect of stochasticity in landscape element composition by the equations:

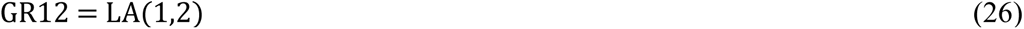

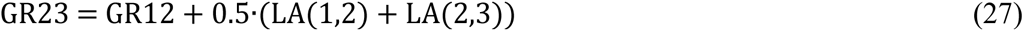

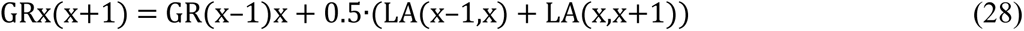

1. Positions of OEs along an ONV relative to this new scale were found by linear interpolation within each segment: Let IGRxy denote positions of borders between ONV segments *x* and *y* along the ONV as originally scaled from 0 to 1 and let *y* be the position of one specific OE on this scale. Within each interval, a linear rescaling was then done by converting the position *t* of an OU, given on the original measurement scale, into a new position z on the PD measurement scale from 0 to LGL using the equation:

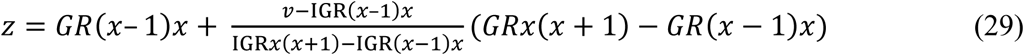 The vector Z is denoted the rescaled ONV (rONV).
2. For use in some analyses, the rONV vector was ranged to a scale from 0 to 1 to compare relative placements along the rONVs. These ranged, rescaled ONVs are referred to as rrONVs.

A standard segment along a CLG, represented by an rONV, is defined as an interval along the CLG/ONV with a minimum extent of at least 1 EDU–L unit (= 0.08 PD units). This definition accords with the general definition of standard segments in EcoSyst (Halvorsen et al. 2020): ‘one in a set of intervals into which a complex-gradient is divided, that is made up by one, two or more elementary segments, each comprising at least 1 ecodiversity distance unit (EDU) of variation in the key characteristic within the major type in question’. As with the standardised division of LECs, the number of major-type adapted segments is set to

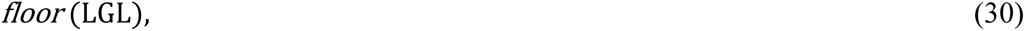

that is, the number of EDU-L between gradient endpoints rounded down to the nearest integer number. Accordingly, a gradient length of at least 2 EDU–L units (> 0.16 PD units) is required for dividing a CLG (rONV) into two segments (and two series of minor types), at least 0.24 PD-units is required for division into three segments, etc.

#### Interpretation and manual pruning of results

After gradient-length estimates had been obtained and all CLGs were divided into segments, the results were subjected to interpretation and manual pruning by guidelines parallel to those applied to the ecosystem system level (see Halvorsen et al. 2016: chapter B4c; Halvorsen et al. 2020: Appendix S2): Combinations of segments with very few OUs were not accepted as minor types unless they represented interpretable, distinct although rare landscapes (e.g. highly urbanised landscapes, mountain summits, glaciers etc.). No fixed limit on OU frequency was set; this issue was discussed separately for each major-type candidate as part of the interpretation of results. OUs initially affiliated with declined minor-type candidates were transferred to neighbouring minor types in CLG space by the ‘snippet criterion’ as follows: First, the CLGs were arranged in order of decreasing importance (decreasing variation explained or gradient length). Secondly, each OU was affiliated with the type with the same gradient-segment combination, except for the least important CLG. The procedure was continued until only accepted minor types were left.

### OPERATIONALISATION OF rONVs TO KEY VARIABLES FOR PRACTICAL IDENTIFICATION OF MINOR LANDSCAPE TYPES

A fundamental principle in the ‘pilot Nordland project’, which is carried over to NiN version 2, is that the correlative analyses of variation in landscape-element composition in large data sets is just a first (but crucial) step on the road towards a system of landscape types. After thorough interpretation, the results are discussed and operationalised. The first step towards operationalisation of the analytic results, was to ‘translate’ the rONV representation of CLGs to key variables or other explicit criteria that could be used in practice for drawing borders between standard segments. In some cases, these key variables or criteria can be used directly to draw boundaries between spatial landscape units, in other cases, further iterations are needed for ‘operationalisation’, e.g. by modelling or by defining operationalised key variables (OpNV). In accordance with the stated aim, the present study includes a discussion of potential OpNVs for each CLG. The use of these key variables to obtain a landscape-type map for Norway is treated in a companion paper (Simensen et al., in prep.).

## RESULTS AND INTERPRETATION

### IDENTIFICATION OF MAJOR-TYPE GROUPS

#### Ordination of Tot3966 – the total data set

The 3966 OUs were distributed on landscape-type variables (major types in the ‘pilot Nordland project’) as follows: KF (fjords): *n* = 349; KS (coastal plains): *n* = 521; KA (coastal hills and mountains): *n* = 3; ID (Valleys): *n* = 1017 and; IA (inland hills and mountains): *n* = 2076. Note that the IS (inland plains) major type was not recognised in Nordland and was therefore not included among landscape-type variables.

The total data set was too large to be ordinated by GNMDS, and unconfirmed DCA ordination results are therefore reported. The DCA ordination had two strong axes, as indicated by a considerable reduction of eigenvalues and gradient lengths from DCA axis 2 to DCA axis 3 (Table 8). The ratio of gradient lengths for DCA axes 2 and 1 was 0.945. Further description and interpretation was therefore based upon DCA axes 1 and 2.

**Table 8.** Eigenvalues and gradient lengths for the DCA ordination of the total dataset Tot3966. S.D. units = standard deviation units

|  | DCA axis 1 | DCA axis 2 | DCA axis 3 | DCA axis 4 |
| --- | --- | --- | --- | --- |
| Eigenvalue | 0.2154 | 0.1651 | 0.0714 | 0.0532 |
| Gradient length (S.D. units) | 2.4362 | 2.3029 | 1.6761 | 1.8097 |

The ordination diagram for DCA axes 1 and 2 (Fig. 8) had no visible artefacts and separated coast from inland and, accordingly, fjord (KF) from valley (ID); see Fig. 9. In order to deduce precise criteria for the border between coast and inland (e.g. at the major-type group level), OUs with special combinations of characteristics (in the borderland between coast and inland) were studied in more detail (Table 9; Figs 10–17). With few exceptions, OUs without coastline were clearly separated from OUs with coastline also in the sparse area in the ordination diagram (Fig. 10). This also applied to OUs which, due to inaccuracy in the delineation of OUs, possessed a small element of coastline (Figs 10, 14); also these OUs possessed ‘coastal characteristics’. The only group in Table 9 that was divided into almost equally large subgroups on either side of the sparse area, was group 2 of Table 9 (Fig. 11): for IA (inland hills and mountains) without coastline (KL = 1) on a large island (SN = 2). OUs on smaller islands (group 3; SN = 3; largest island < 20 km^2^; Fig. 12), on the other hand, possessed characteristics more typical of coastal landscapes, regardless if a coastline was present in the OU or not.

**Fig. 8.**
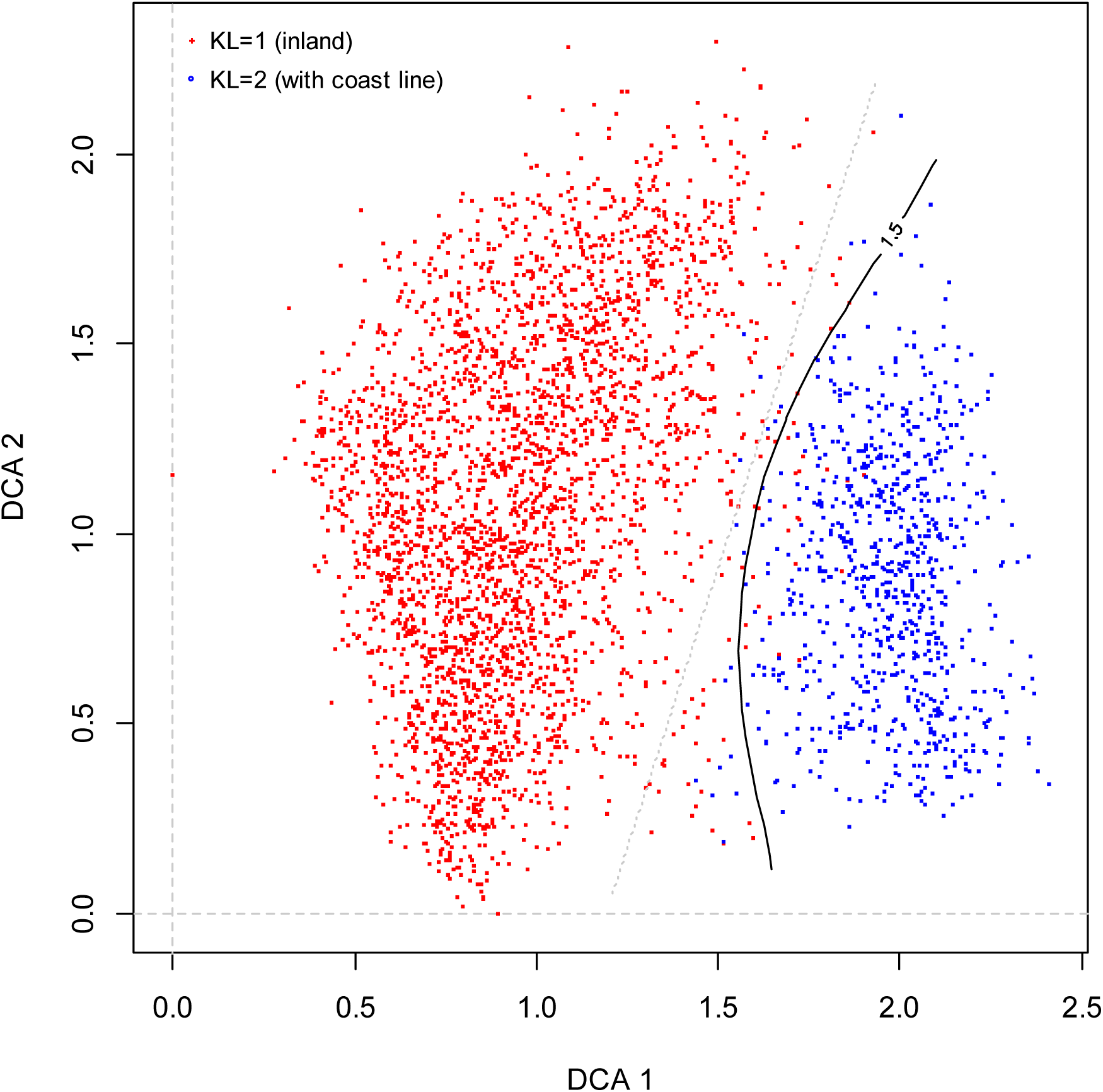
DCA ordination of the Tot3966 data set (all 3966 observation units), axes 1 and 2 (scaled in standard deviation units). KL = 1: inland observation units without coastline (red); KL = 2: observation units without coastline (blue). The black, continuous ‘1.5’ line marks equal probability (0.5) for an OU to be affiliated with inland and coastal landscapes.

**Fig. 9.**
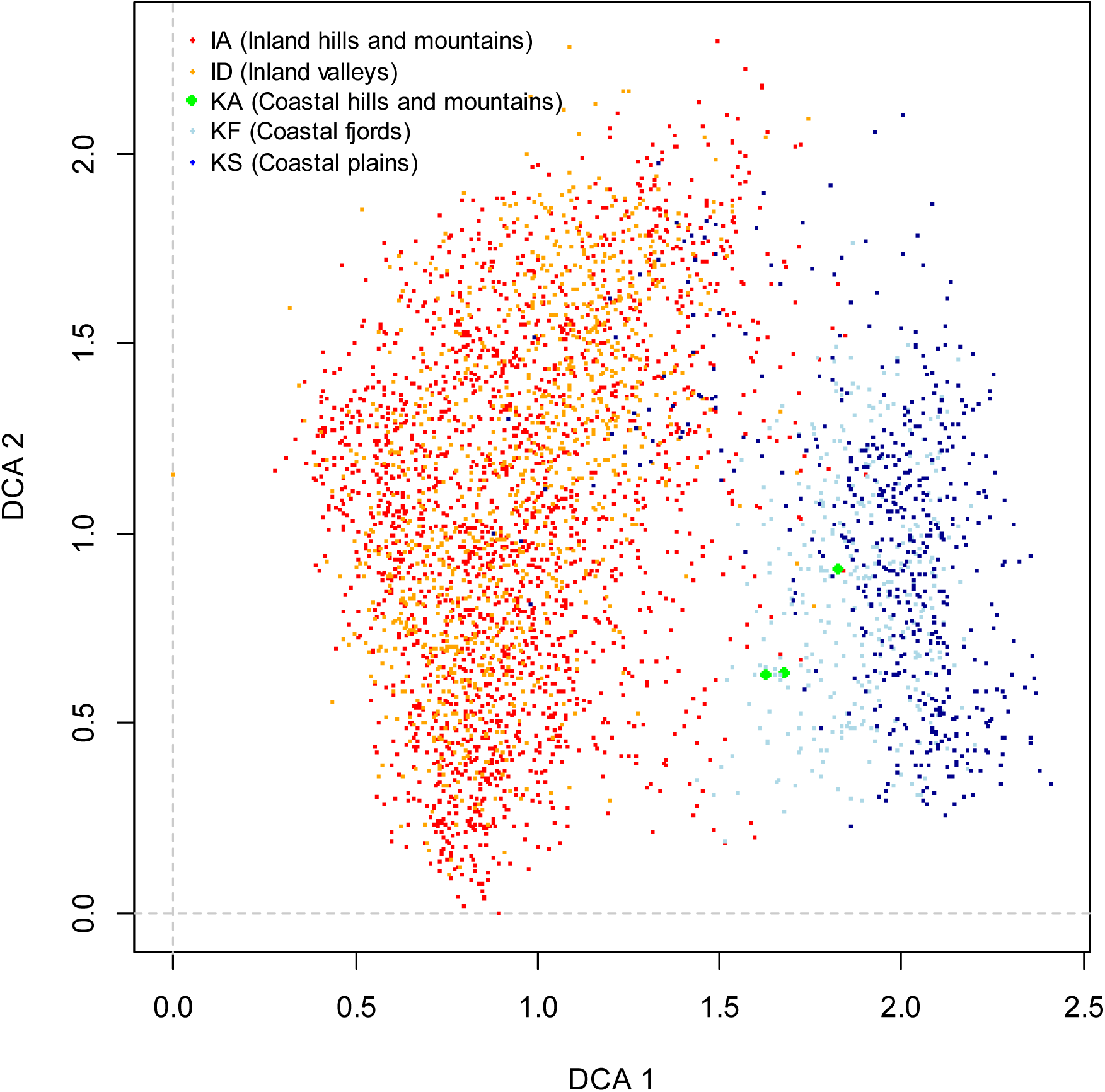
DCA ordination of the Tot3966 data set (all 3966 observation units), axes 1 and 2 (scaled in S.D. units). Colours indicate affiliation to major-types of the ‘pilot Nordland project’ (LT variables; see Table 1).

**Figs. 10–15.**
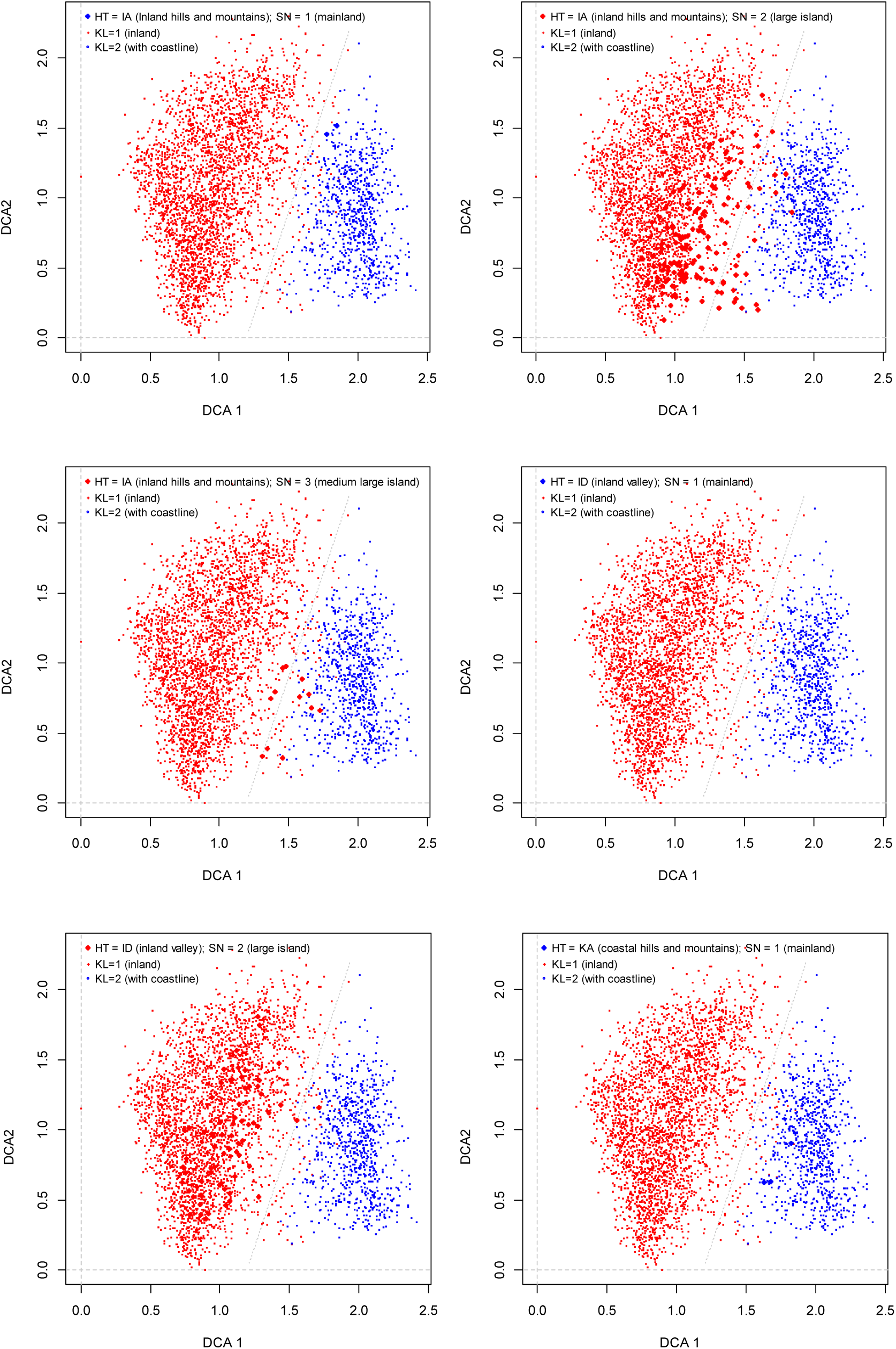
Distribution of observation units with specific properties (LT-variable, location on island, presence or absence of coastline; large dots) in the transition zone between inland and coastal landscapes in the DCA ordination diagram, axes 1 and 2, for the Tot3966 data set (scaled in S.D. units). Red dots = inland OUs (without coastline; KL = 1); blue dots = coastal OUs (with coastline; KL = 2). Affiliation to LT-variables (LT variables, see Table 1): IA = inland hills and mountains; ID = Inland valleys; KA = coastal hills and mountains. SN = 1: mainland OU; SN = 2: observation unit on large island (> 20 km2); SN = 3: observation unit on small or medium-sized island (1.5–20 km^2^).

**Table 9.** Characterisation of observation units in the transition zone between inland and coastal landscapes in the DCA ordination diagram of the Tot3966 data set (Fig. 8). IA = inland hills and mountains; ID = (inland) valleys; KA = coastal hills and mountains; KS = coastal plains; KL = 1: inland observation units without coastline (red dots in Fig. 8); KL = 2: observation units with coastline (blue dots in Fig. 8); SN = 1: observation unit on the Norwegian mainland; SN = 2: observation unit on large island (> 20 km^2^); SN = 3: observation unit on small or medium-sized island (1.5–20 km^2^).

| Group | Grouping criteria | No. of OUs | Description | Interpretation |
| --- | --- | --- | --- | --- |
| 1 | IA<br>&KL = 2<br>&SN = 1 | 2 | Inland hill and mountain OUs on the mainland with coastline (resulting from delineation inaccuracies) | Both clearly belongs to the coastal group |
| 2 | IA<br>&KL = 1<br>&SN = 2 | 150 | Inland hills and mountain OUs on island $> 20 \text{ km}^2$, without coastline | Not surprisingly, these OUs are not distributed over the full range of variation across the sparse region area (in the ordination diagram) between inland and coast. The subset with these 150 OUs should be further analysed to reveal detailed patterns |
| 3 | IA<br>&KL = 1<br>&SN = 3 | 12 | Inland hill and mountain OUs on island $1.5\text{--}20 \text{ km}^2$, without coastline. | These OUs are located on the 'coastal side' or very close to the coastal side of the sparse region and make up a group with inland polygons on an island, which should be analysed further |
| 4 | ID<br>&KL = 2<br>&SN = 1 | 1 | Valley OU on the mainland with coastline (probably resulting from erroneous delineation) | Like IA OUs and groups 1, this OU clearly belongs to the coastal group |
| 5 | ID<br>&KL = 1<br>&SN = 2 | 70 | Inland valley OUs on island $> 20 \text{ km}^2$, without coastline | With two exceptions, these OUs are located on the 'inland side' of the sparse area. Their positions differ clearly from ID & KL = 2 & SN = 2; showing that the occurrence of a coastline is crucial for a valley polygon to have 'coastal character' |
| 6 | KA | 3 | Three coastal hill and mountain OUs (KL = 2 & SN = 1) | Clearly belonging to the coastal group |
| 7 | KS<br>&KL = 1<br>&SN = 0 | 31 | OU's from coastal plains without coastline, on the mainland | With 3 (–4) exceptions, all of these OUs belong to the inland group and have high DCA-axis 2 scores, high areal cover of mires and lakes, and possess infrastructure |
| 8 | KS<br>&KL = 1<br>&SN = 2 | 33 | OU's from coastal plains without coastline, on large island. | With 3 exceptions, all of these OUs clearly belong to the inland group |

*Conclusion.* Two data subsets, consisting of OUs with uncertain affiliation to coast or inland major-type groups, were identified and analysed further:

**X162**: 162 OUs in LT-type IA (without coastline), located on island
**X64**: 64 OUs in LT-type KS (coastal plains), located on island

The OUs with coastline, erroneously assigned to inland LT types, of groups 1 and 4, were removed before further analysis.

#### Examination of data subset X162 – OUs assigned to LT-type IA (inland hills and mountains) located on islands

Of the 162 OUs in data subset X162, 31 were located to the right of the dotted line through the sparse region in the Tot3966 DCA ordination diagram (Fig. 16), thus having ‘coastal characteristics’, while the remaining 131 OUs were located to the left of that line. Using absence of coastline as a strict criterion for dividing OUs into ‘inland’ and ‘coast’ would therefore imply misclassification of 31/162 = 19.1% of OUs in the X162 subset, given that the dotted line in the sparse region of Figs 8–16 was taken as the true border between ‘inland’ and ‘coast’ groups. A strong but not fully linear relationship between the Oyst_i (inverse island size) variable and OU positions was found across the sparse region (Fig. 16): the dotted line roughly corresponded to Oyst_i = 0.6, i.e. island size ≈ 216 km^2^. If the Oyst_i variable was used to sort subset X162 OUs on inland and coastal groups, the lowest misclassification rate (proportion of OUs that were misclassified to the inland and coastal groups, based upon Oyst_i values), 14.8%, would be obtained for Oyst_i = 0.733, i.e. by including OUs of islands < 10 km^2^ without coastline in the ‘coastal group’ (Fig. 17).

**Figs. 16–17.**
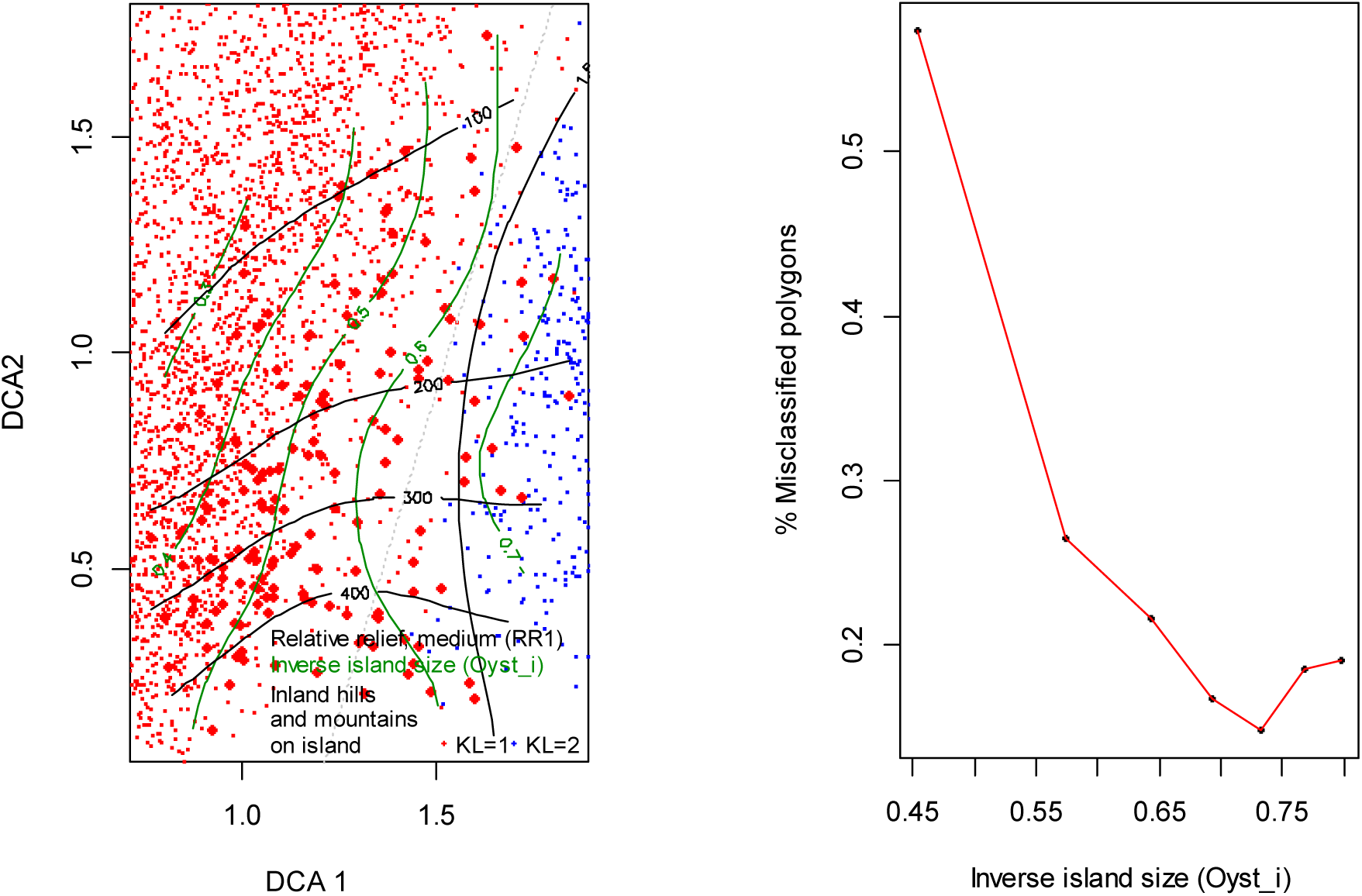
(16) Close-up showing the distribution of the 162 OUs in the X162 data subset (large, red dots) in the transition zone between inland (without coastline, left; red dots) and coastal landscapes (with coastline; blue dots) in the DCA ordination diagram, axes 1 and 2, for the Tot3966 data set (scaled in S.D. units). Isolines for medium relative relief (analytic variable RR1_m; black lines) and inverse island size (Oyst_i; green lines) are shown. OUs affiliated to LT-type IA, situated on islands, are indicated by large red dots. (17) Proportion of OUs misclassified (i.e., OUs without coastal line that are erroneously classified as coast), as function of inverse island size (Oyst_i index), provided that inverse island size is used to differentiate between coastal and inland major-type groups. The minimum value of 14.8% was obtained for Oyst_i = 0.733, which corresponds to an island size of 10 km^2^.

*Conclusion*. The improvement of precision in assignment of island OUs without coastline obtained by splitting these OUs onto ‘inland’ vs. ‘coastal’ groups in the DCA ordination of the Tot3966 data set by transferring OUs on islands < 10 km^2^ to coastal landscapes, is marginal. Hence, we judge the disadvantages of a more complex definition of these two major landscape-type groups larger than the advantages of a marginally reduced misclassification rate. We therefore conclude that island OUs without coastline should be affiliated with the inland group of OUs.

#### Examination of data subset X64 – OUs assigned to LT-type KS (coastal plains) located on islands

Of the 64 OUs in data subset X64, 7 were located to the right of the dotted line through the sparse region in the Tot3966 DCA ordination diagram (Fig. 18) and the remaining 57 OUs were located to the left of that line. Neither island size nor primary key variables were correlated with positions of these coastal-plain OUs in the ordination diagram (Fig. 18; results of correlation analyses not shown).

**Fig. 18.**
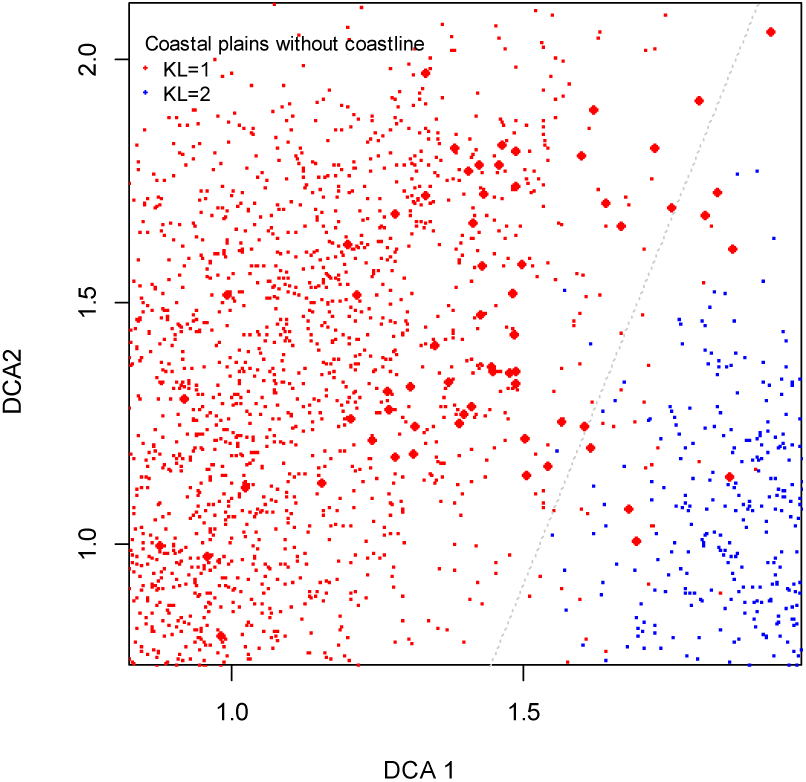
Distribution of the 64 observation units of the X64 data subset (coastal plains without coastline; large dots) in the transition zone between inland and coastal landscapes in the DCA ordination diagram, axes 1 and 2, for the Tot3966 data set (scaled in S.D. units). Red dots = inland OUs (without coastline; KL = 1); blue dots = coastal OUs (with coastline; KL = 2).

*Conclusion*. Island OUs assigned to LT-type coastal plains should be affiliated with the inland group of OUs.

#### Conclusion on division into major-type groups

The analyses support a principal division of OUs into two major-type groups; inland landscapes (I) and coastal landscapes (K). There are good reasons to keep strictly marine landscapes (i.e., without coastline and terrestrial land) as a third major-type group (M).

The results of the analyses of Tot3966 and subsets X162 and X64 support separate analyses of one coastal data set (**K810**) and one inland data set (**I3150**) as the next step in the analysis. Six inland OUs with traces of coastline or incomplete records of some variables were removed before further analysis.

### IDENTIFICATION OF MAJOR-TYPE CANDIDATES

#### Ordination of K810 – the coastal landscapes data subset

All four axes of the 4-dimensional GNMDS ordination of the K810 data subset (Fig. 20) were confirmed by the corresponding DCA axis (Fig. 19; Table 10). The gradient lengths indicated two strong axes (Table 10; axis 1 and 2, respectively). The GNMDS ordination had Procrustes SS = 0.0092 and no unstable OUs (i.e. OUs that obtained widely different positions in the two lowest-stress GNMDS solutions). The first two GNMDS ordination axes separated OUs assigned to ‘pilot Nordland’ major types coastal plains (LT-variable KS) and fjord (KF) along a line from the upper left to the lower right corners of Fig. 21. The black, continuous line in Fig. 21 indicates the transition from preponderance of KS OUs (lower left) to preponderance of KF OUs (upper right). Taking this line as the truth, misclassification rates were 47/350 = 13.4% for KF and 53/457 = 11.6% for KS OUs, respectively.

**Figs. 19–20.**
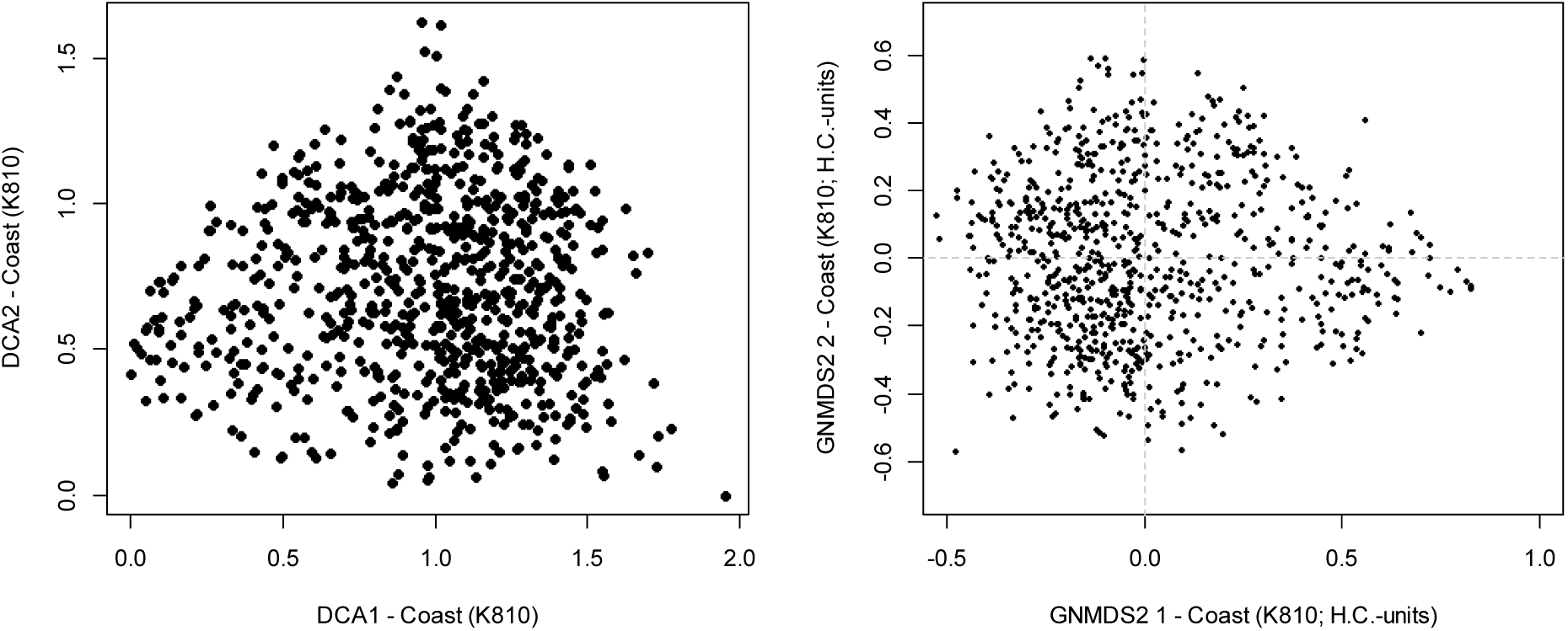
DCA and GNMDS ordinations of the K810 data subset (810 observation units in coastal landscapes), axes 1 and 2. (19) DCA ordination axes are scaled in standard deviation (S.D.) units. (20) GNMDS ordination axes are scaled in half-change (H.C.) units.

**Fig. 21.**
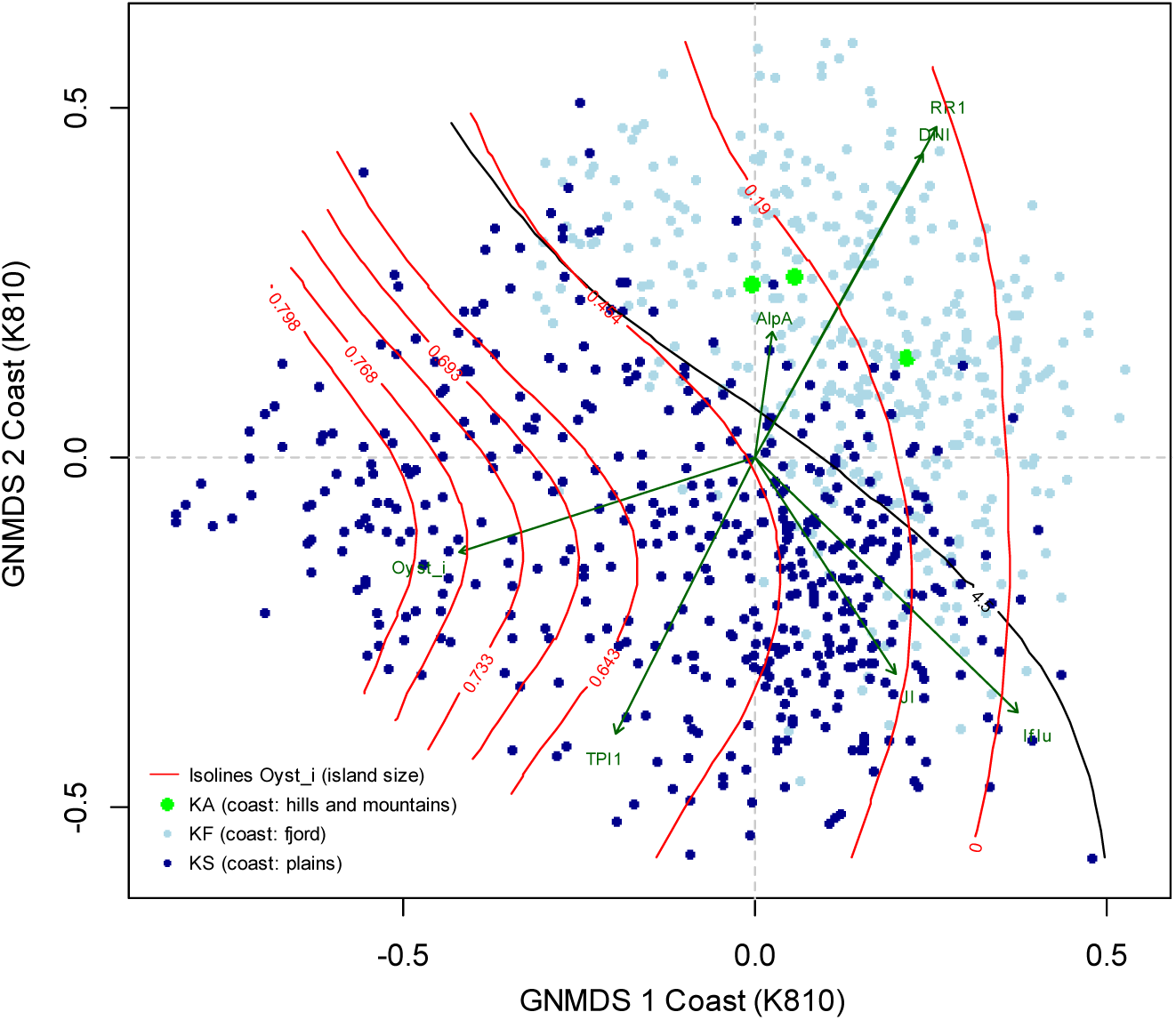
GNMDS ordination of the K810 data subset (810 observation units in coastal landscapes), axes 1 and 2 scaled in half-change (H.C.) units; colours show affiliation to major-types of the ‘pilot Nordland project’ (LT-variables; see Table 1); isolines for island size (variable Oyst_i); and vectors pointing in the direction of maximum increase of Oyst_i and selected primary key variables (see Table 4 for explanation). The black, continuous ‘4.5’ line marks equal probability (0.5) for an OU to be affiliated with KS and KF.

**Table 10.** Kendall’s rank correlation coefficients (τ), eigenvalues and gradient lengths for parallel DCA and GNMDS ordinations of the subset K810 (coastal landscapes).

|  | GNMDS axis 1 | GNMDS axis 2 | GNMDS axis 3 | GNMDS axis 4 |
| --- | --- | --- | --- | --- |
| Strongest correlated DCA axis | DCA 1 | DCA 2 | DCA 3 | DCA 4 |
| $\tau$ (with strongest correlated DCA axis) | 0.8102 | 0.8046 | 0.5555 | 0.5174 |
| DCA-axis eigenvalue | 0.1166 | 0.0880 | 0.0424 | 0.0387 |
| Gradient length of DCA-axis (S.D. units) | 1.9564 | 1.6602 | 1.2810 | 1.2003 |
| Gradient length of GNMDS axis (H.C. units) | 1.3520 | 1.1630 | 0.9110 | 0.9450 |

Isolines for Oyst_i in Fig. 21 established island size as an important variable for explaining variation among KS OUs, while island size was less important in fjords. The three OUs assigned to ‘pilot Nordland’ major type KA (coastal hill and mountain landscapes) were placed inside the cluster of KF OUs. This suggests that there is no fundamental difference between the side of a fjord and a mountain coast, most likely because, at the outset, there is no logical reason why a steep fjord side and a steep coastline not situated in a fjord-like landform should differ in other landscape features than, perhaps, terrain roughness. Furthermore, this indicates that the criterion adopted in the ‘pilot Nordland project’, to divide fjord segments into separate OUs for the two sides when they have different landscape element composition and the width of the fjord exceeds 1 km, should be carried over to NiN version 2.

*Conclusion*. The analyses support a principal distinction between coastal plains and fjords, based on the criteria that were used to distinguish two major types in the ‘pilot Nordland project’ (Appendix S1). The sparse data material does not support discrimination of coastal hills and mountains as a major type on its own.

#### Ordination of I3150 – the inland landscapes data subset

After removal of analytic variables that express coastal characteristics, 67 variables were retained in the inland data subset I3150. This data subset was too large to be ordinated by GNMDS, and unconfirmed DCA ordination results are therefore reported (Table 11). DCA axis 1 had an eigenvalue more than twice as large as that of DCA axis 2, which in turn had an eigenvalue 1.8× that of DCA axis 3. The gradient length of DCA axis 1 was ca. 1.5× that of DCA axis 2. This indicated existence of one strong and one less strong, but potentially interpretable, gradient in landscape-element composition in the I3150 data subset. The DCA ordination diagram for axes 1 and 2 (Fig. 22) did not clearly separate the three relevant major types of the ‘pilot Nordland project’: inland hills and mountains (IA), (inland) valleys (ID) and coastal plains (KS), i.e. landscapes close to the coast but without coastline. A slight tendency for concentration of IA-affiliated OUs at low DCA axis 1 scores (to the left) and ID- and KS-affiliated OUs to the right in Fig. 22 could, however, be observed. Affinity to IA, ID or KS did, however, only explain 3.7% of the variation in I3150 (ANOVA: F_2,3147_ = 60.35, r^2^ = 0.0369, P < 0.0001).

**Fig. 22.**
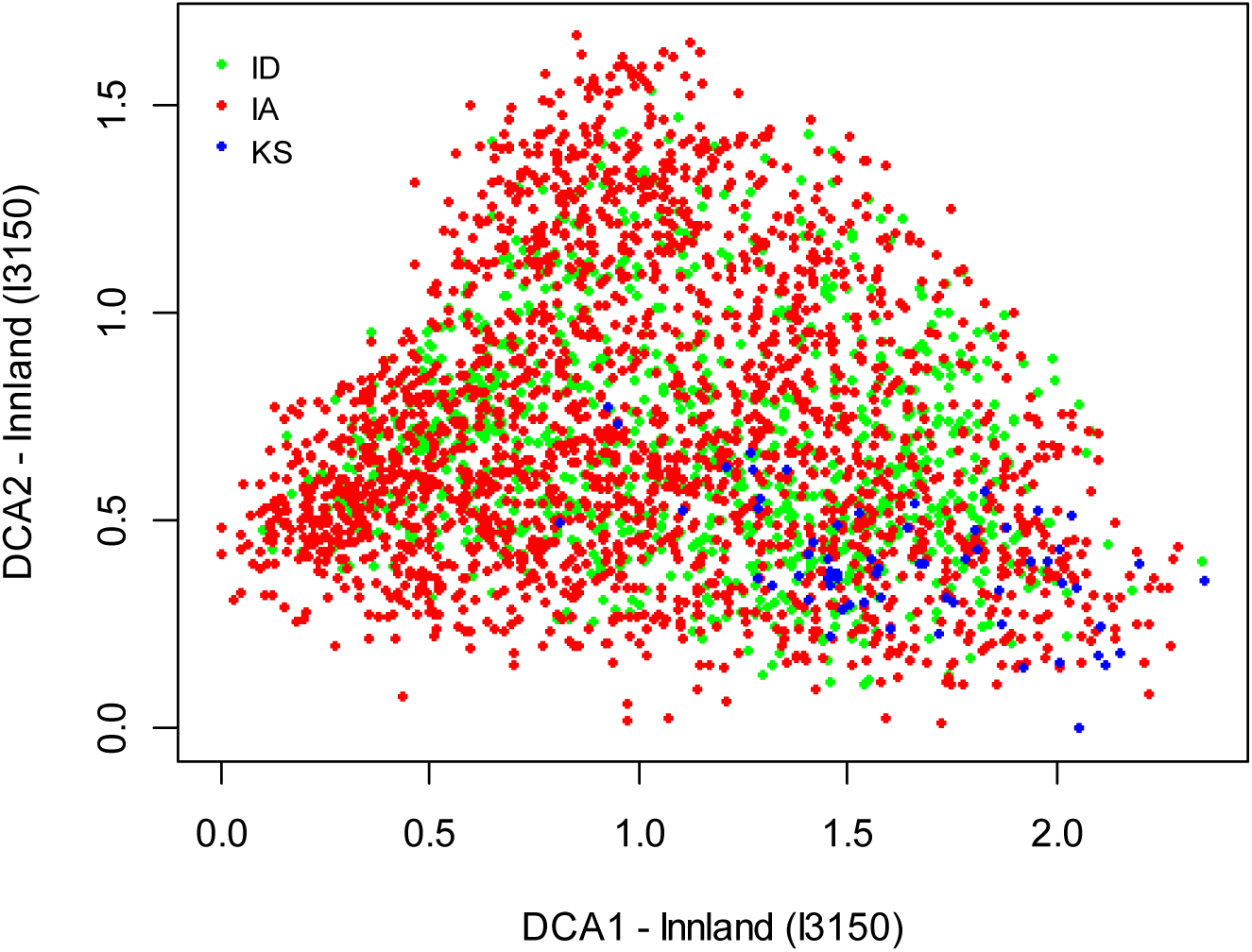
DCA ordination of the I3150 data subset (3150 observation units from inland landscapes), axes 1 and 2 (scaled in S.D. units). Colours show affiliation to major-types of the ‘pilot Nordland project’ (LT-variables; see Table 1).

**Table 11.** Eigenvalues and gradient lengths for the DCA ordination of the subset I3150, inland landscapes.

|  | DCA axis 1 | DCA axis 2 | DCA axis 3 | DCA axis 4 |
| --- | --- | --- | --- | --- |
| Eigenvalue | 0.2066 | 0.0913 | 0.0510 | 0.0548 |
| Gradient length (S.D. units) | 2.3513 | 1.6664 | 1.5127 | 1.5018 |

Correlations with primary key and analytic variables established DCA axis 1 as an infrastructure gradient, while DCA axes 2–4 had no obvious interpretation in terms of explanatory variables (Table 12). Furthermore, the correlation analysis also suggested that variation in the intensity of human exploitation was conditioned on underlying geo-ecological variation. Geo-ecological variables correlated with DCA axis 1 (τ values in parentheses) were (Table 14): areal coverage of rugged terrain (Guro_t_a; τ = –0.4391); relief (RR1; τ = – 0.4169); areal coverage of flat terrain (Flat_a; τ = 0.4032); areal coverage of steep terrain (Steep_a; τ = –0.3890); mean terrain ruggedness (Rug3_m; τ = –0.3554); and areal coverage of terrain depressions (TPI1h_a; τ = –0.3151). This supported interpretation of DCA axis 1 also as a gradient from a rough and coarse-scaled, alpine terrain towards a more gentle, less ‘rugged’ terrain. Glaciers, as indicated by variables BrI and Glac_a, and several soil variables, were also correlated with DCA 1 (τ> 0.30; see Table 12). While glaciers were concentrated to low DCA axis 1 scores, the opposite was true for mires.

**Table 12.**
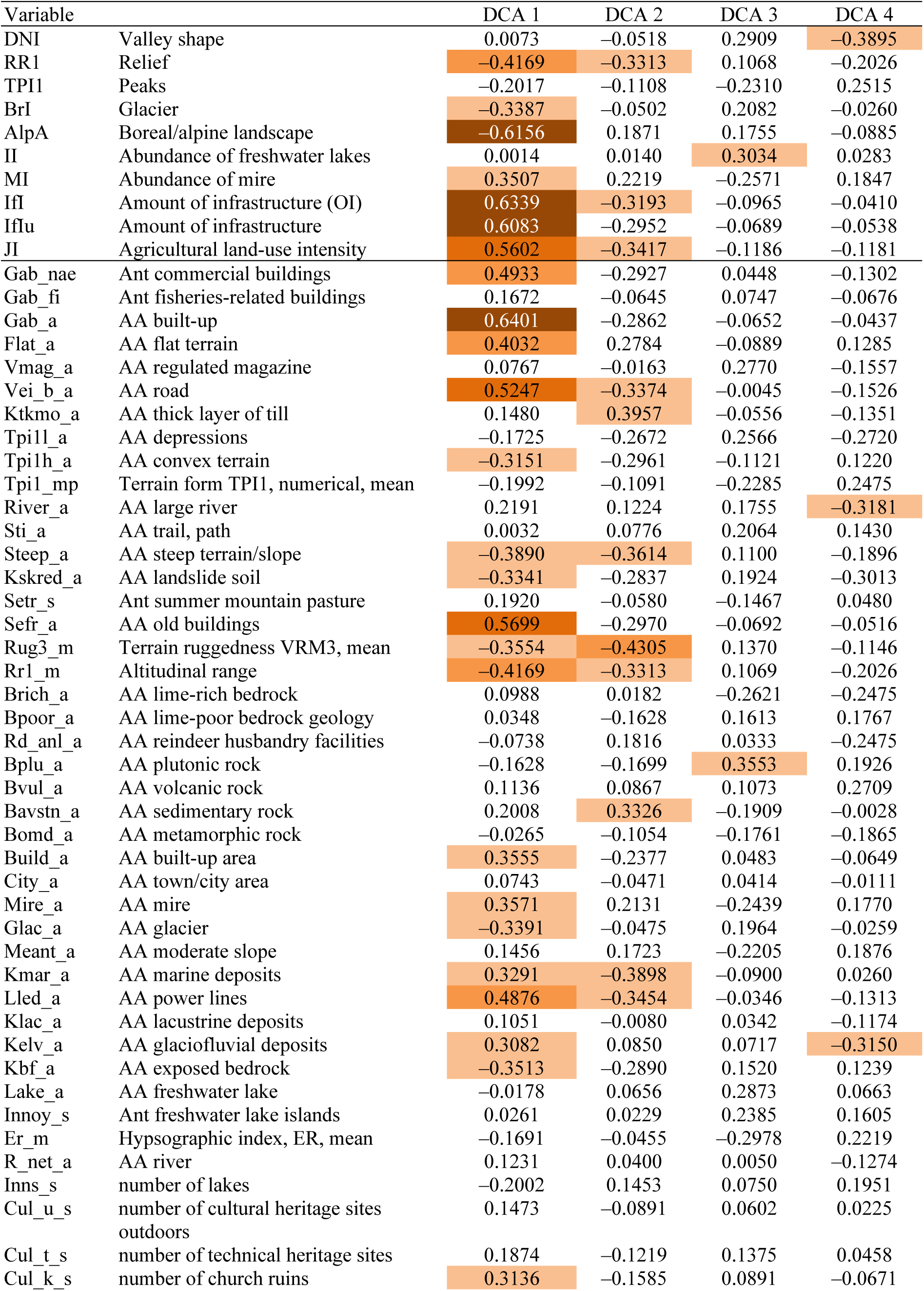
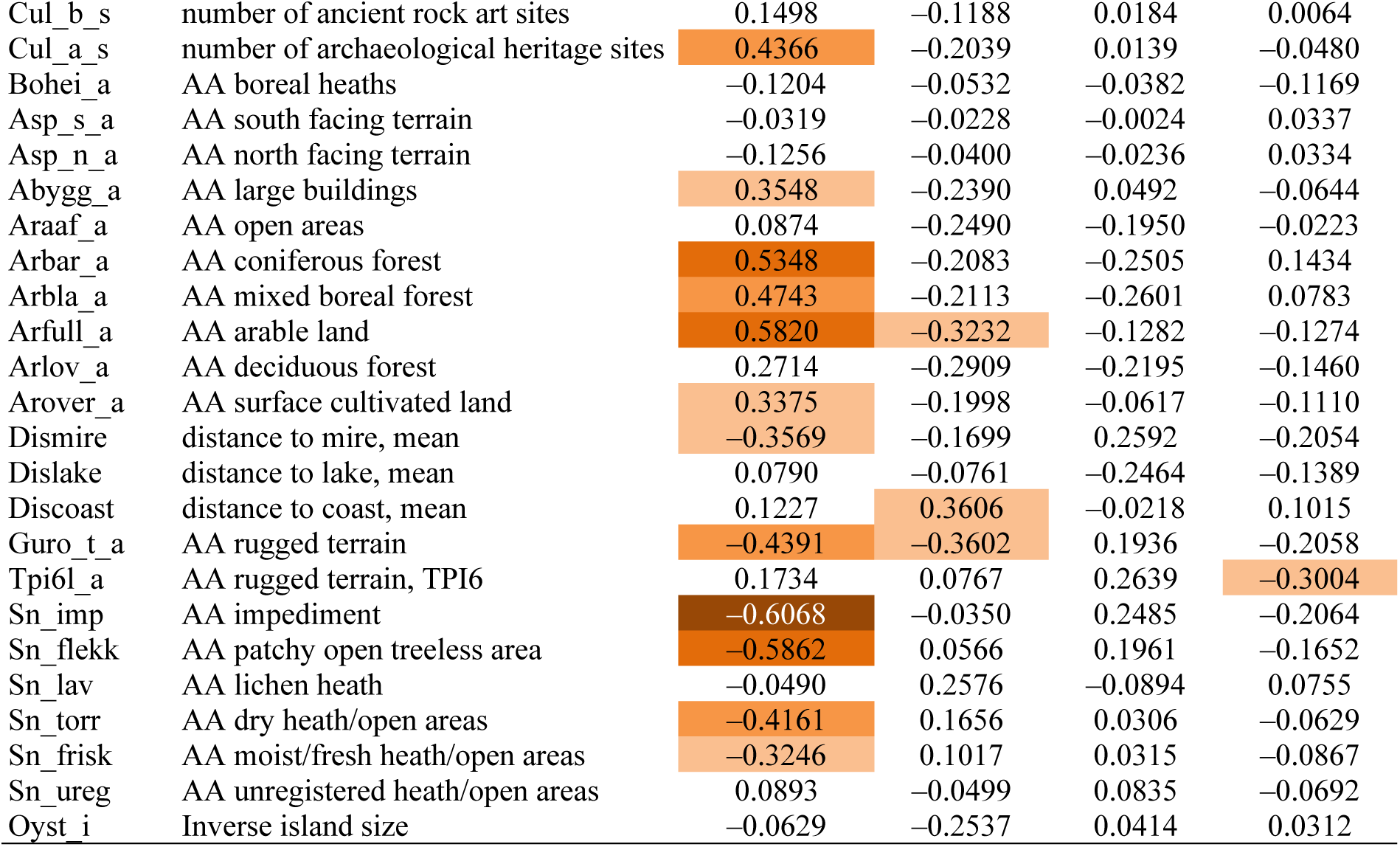
Kendall’s rank correlation coefficients (τ) between DCA ordination axes 1–4 for the inland landscapes data subset (I3150) and the and the 10 primary key variables (rows 1–10) and the 67 analytic variables used to characterise the 3150 observation units.

**Table 13.** Eigenvalues and gradient lengths for the DCA ordination of geo-ecological variables in the inland dataset I3150G. S.D. units = standard deviation units.

|  | DCA axis 1 | DCA axis 2 | DCA axis 3 | DCA axis 4 |
| --- | --- | --- | --- | --- |
| Eigenvalue | 0.1430 | 0.0725 | 0.0599 | 0.0632 |
| Gradient length (S.D. units) | 1.9927 | 1.7371 | 1.5389 | 1.6465 |

**Table 14.** Kendall’s rank correlation coefficients (τ) between DCA ordination axes 1–4 for the inland landscapes data subset (I3150) based upon 10 primary key variables (rows 1–10) and the 35 basic geo-ecological variables characterising the 3150 observation units.

| Variable |  | DCA 1 | DCA 2 | DCA 3 | DCA 4 |
| --- | --- | --- | --- | --- | --- |
| DNI | Valley shape | 0.0724 | 0.3643 | –0.3911 | –0.2986 |
| RR1 | Relief | 0.6338 | 0.2401 | –0.1329 | 0.1285 |
| TPI1 | Peaks | 0.2259 | –0.2105 | 0.2929 | 0.3543 |
| BrI | Glacier | 0.3240 | 0.0884 | –0.1266 | 0.0962 |
| AlpA | Boreal/alpine landscape | 0.2892 | 0.0412 | –0.1191 | 0.1729 |
| II | Abundance of freshwater lakes | 0.0001 | –0.1046 | –0.0647 | –0.2658 |
| MI | Abundance of mire | –0.4007 | –0.2133 | 0.0393 | –0.0794 |
| IfI | Amount of infrastructure | –0.2107 | 0.0701 | 0.0878 | –0.1420 |
| IfIu | Amount of infrastructure | –0.2076 | 0.0800 | 0.0622 | –0.1421 |
| JI | Agricultural land-use intensity | –0.2082 | 0.1612 | 0.1222 | –0.1098 |
| Flat_a | AA flat terrain | –0.5885 | –0.1469 | 0.0992 | –0.1615 |
| Ktkmo_a | AA thick layer of till | –0.3582 | 0.0329 | –0.3162 | –0.0194 |
| Tpi1l_a | AA depressions | 0.3654 | 0.2946 | –0.2096 | –0.1671 |
| Tpi1h_a | AA convex terrain | 0.5098 | –0.0609 | 0.1959 | 0.3138 |
| Tpi1_mp | Terrain form TPI1, numerical, mean | 0.2229 | –0.2075 | 0.2878 | 0.3504 |
| River_a | AA large river | –0.2657 | 0.2983 | –0.3481 | –0.3556 |
| Steep_a | AA steep terrain/slope | 0.6380 | 0.2309 | –0.1087 | 0.1152 |
| Kskred_a | AA landslide soil | 0.4642 | 0.2835 | –0.1899 | 0.0049 |
| Rug3_m | Terrain ruggedness VRM3, mean | 0.6675 | 0.1649 | –0.0190 | 0.0732 |
| Rr1_m | Altitudinal range | 0.6338 | 0.2401 | –0.1329 | 0.1285 |
| Brich_a | AA lime-rich bedrock | –0.0871 | 0.2513 | –0.1620 | 0.3392 |
| Bpoor_a | AA lime-poor bedrock geology | 0.1049 | –0.1502 | 0.1375 | –0.2784 |
| Bplu_a | AA plutonic rock | 0.2785 | –0.2505 | –0.0668 | –0.1936 |
| Bvul_a | AA volcanic rock | –0.1183 | –0.2945 | –0.3170 | –0.0252 |
| Bavstn_a | AA sedimentary rock | –0.4445 | –0.1730 | –0.1694 | 0.2524 |
| Bomd_a | AA metamorphic rock | 0.1111 | 0.3456 | 0.1947 | 0.0279 |
| Mire_a | AA mire | –0.4003 | –0.2043 | 0.0380 | –0.0912 |
| Glac_a | AA glacier | 0.3236 | 0.0899 | –0.1224 | 0.1102 |
| Meant_a | AA moderate slope | –0.2058 | –0.1482 | 0.1160 | 0.1047 |
| Kmar_a | AA marine deposits | –0.0811 | 0.0869 | 0.3773 | –0.0744 |
| Klac_a | AA lacustrine deposits | –0.1129 | 0.1467 | 0.0356 | –0.0854 |
| Kelv_a | AA glaciofluvial deposits | –0.3204 | 0.3230 | –0.2468 | –0.3037 |
| Lake_a | AA freshwater lake | –0.0175 | –0.1526 | –0.0467 | –0.2746 |
| Innoy_s | Ant freshwater lake islands | –0.0102 | –0.1890 | 0.0147 | –0.2994 |
| Er_m | Hypsographic index, ER, mean | 0.1563 | –0.1593 | 0.2698 | 0.4076 |
| R_net_a | AA river | –0.1108 | 0.1439 | –0.1204 | –0.1789 |
| Inns_s | Number of lakes | 0.0509 | –0.2489 | 0.0887 | –0.0836 |
| Asp_s_a | AA south facing terrain | 0.0541 | –0.0235 | 0.0346 | 0.0101 |
| Asp_n_a | AA north facing terrain | 0.1351 | –0.0338 | 0.0468 | 0.0848 |
| Dismire | Distance to mire, mean | 0.3603 | 0.2195 | –0.0653 | 0.0814 |
| Dislake | Distance to lake, mean | –0.0241 | 0.2160 | –0.0089 | 0.2302 |
| Discoast | Distance to coast, mean | –0.3044 | –0.1234 | –0.1865 | –0.0662 |
| Guro_t_a | AA rugged terrain | 0.6483 | 0.2276 | –0.1159 | 0.0692 |
| Tpi6l_a | AA rugged terrain, TPI6 | –0.1741 | 0.2612 | –0.3445 | –0.3808 |
| Oyst_i | Inverse island size | 0.1817 | –0.1184 | 0.1729 | –0.0940 |

*Conclusion*. Besides identifying a clear main gradient in landscape-element composition related to land use and observable infrastructure, the ordination results (DCA axis 1) support existence a rough, geomorphologically based subdivision of inland landscapes into valleys and ‘non-valley landscapes’. However, the variation in landscape element composition in the inland data set I3150, as revealed by DCA ordination of all 67 analytic variables, appears so complex that important structural patterns may be hidden in this complexity. Also, the fact that the ordination of the I3150 data subset indicated that the dominant infrastructure gradient is conditioned on variation in geo-ecological (geomorphological) properties, may indicate that variation in landscape-element composition relative to basic geomorphological variation is concealed behind, or overshadowed by, land-use related variation. A new ordination of I3150, using the basic geo-ecological analytic variables (category G) only, was therefore undertaken, in order to clarify the basis for dividing inland landscape into major types.

#### Ordination of I3150G – the inland landscapes data subset, using basic geo-ecological analytic variables only

The reduction from 67 to 35 variables did not change the fact that the I3150 data subset was too large to be ordinated by GNMDS, and results of an unconfirmed DCA ordination are therefore reported (Table 13). DCA axis 1 had an eigenvalue almost twice as large as that of DCA-axes 2–4, while the gradient lengths of the four DCA axes were in the same range (Table 13). The OUs were evenly distributed in the space spanned by DCA ordination axes 1 and 2, without obvious artefacts (a very weak tongue effect can be traced on DCA axis 2).

Analysis of correlations between DCA axes and the primary key and analytic variables (Table 14) revealed a very strong relationship between DCA axis 1 and terrain ruggedness (relief-related variables) and a distinct relationship between DCA axis 2 and valley shape (DNI variable, expressing the relationship between valley depth and valley width). The results of this ordination were examined graphically in greater detail (Figs 23–28), with the aim of finding a supported division into major types.

**Fig. 23-28.**
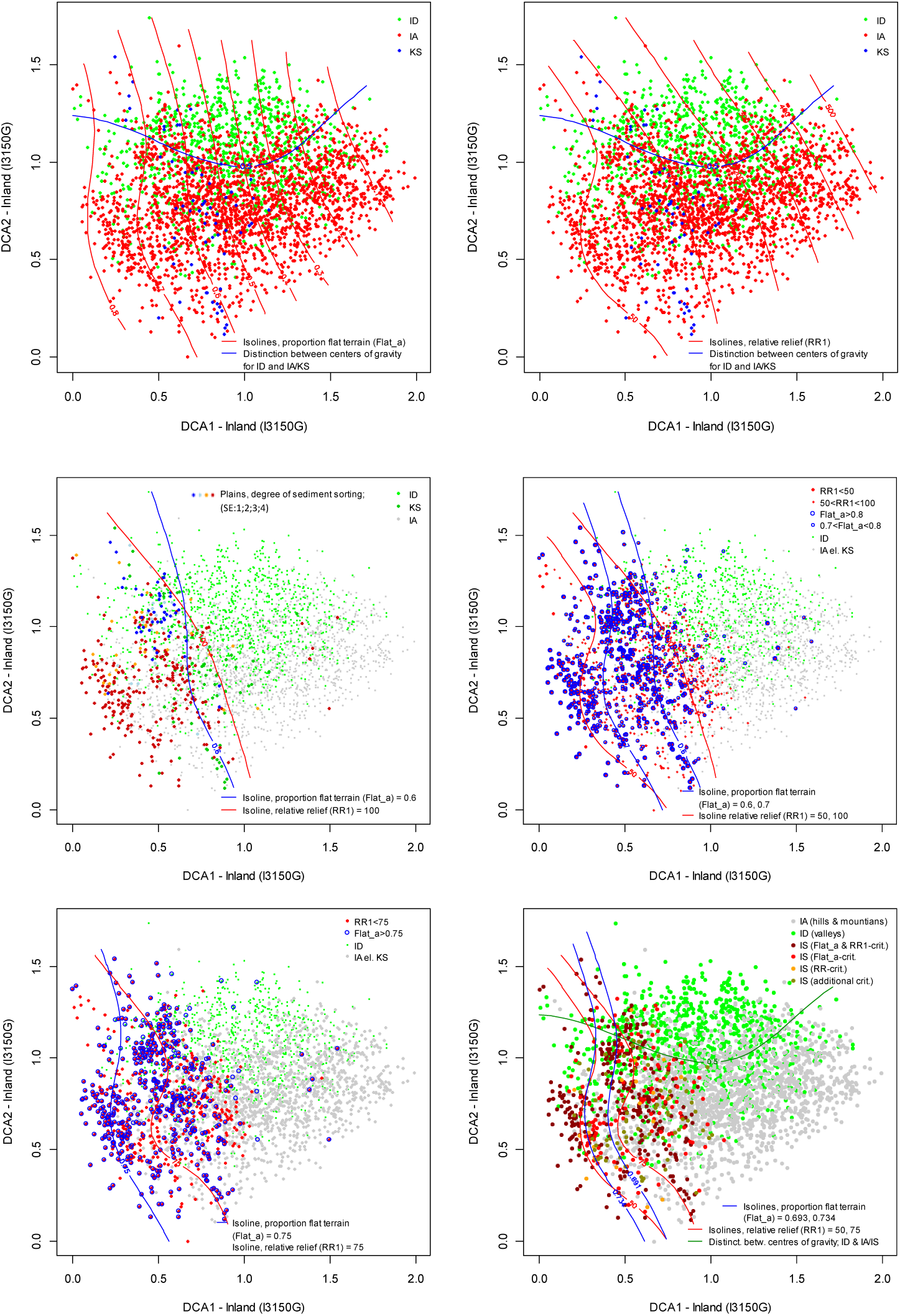
Distribution of observation units with specific properties in the DCA ordination diagram, axes 1 and 2, for the inland data set (I3150G) (scaled in S.D. units). ID = inland valleys; IA = Inland hills- and mountains; IS = Inland plains; KS = Coastal plains; Flat_a = proportion of area with flat terrain; RR1 = relative relief (altitudinal range); SE = sediment sorting; SE1 = unsorted (< 10% of area covered by sorted sediments, i.e., glaciofluvial and marine deposits); SE2 = low degree (10–50%) of sorted sediments; SE3 = medium degree (50–75%) of sorted sediments; SE4 = sediment plain (> 75% sorted sediments).

Fig. 23 shows a relatively sharp differentiation between valleys (green symbols) and other landscapes (hill and mountain landscapes and plains) along DCA axis 2 (the shift from preponderance of red or blue vs. green dots in Fig. 23 is indicated by the blue line). Despite the relatively abrupt shift, none of the variables were more than noticeably correlated with DCA axis 2 (Table 14). This indicates that valley landscapes differ from other inland landscapes first and foremost in the overall composition of landscape elements.

Isolines for Flat_a (area of flat land; Fig. 23) and relative relief (RR; Fig. 24) have more or less identical pattern in the DCA ordination, demonstrating a very strong gradient common to all inland landscapes from plains to the most extremely steep and rugged terrain. This gradient is omnipresent, regardless of major geomorphological form (distinct valley, hill, mountain or plain). OUs affiliated with the ‘pilot Nordland’ major type coastal plains, without coastline (blue dots in Fig. 23), neither segregated from hill and mountain landscapes nor from valleys in the ordination diagram but tended to occur in a broad band from low to high scores along DCA axis 2 for relatively low to intermediate scores along DCA axis 1. The almost complete lack of ‘inland coastal plain’ OUs near the low-score end of DCA axis 1 (Fig. 23) suggests existence of inland landscapes with features more typical of plains, such as lower relative relief and higher proportion of flat terrain, than the ‘inland coastal plain’. This result thus supports separation of inland plains as a major type on its own, parallel to the separation of coastal plains (KS) from fjords (KF) in the coastal major-type group.

The degree of sediment sorting (landscape-factor variable SK1: category S; cf. Table 3) increased towards the left in the ordination diagram (low DCA axis 1 scores), i.e. towards OUs with characteristics typical of plains (Fig. 25) such as low relative relief and high proportion of flat terrain (Figs 23–24). Fig. 25 shows that OUs with a high proportion of their area covered by flat terrain also had relative relief << 100 m, suggesting that these two characteristics, separately or in combination, can be used to separate plains from hills/mountain landscapes. Detailed examination of selected OUs from different parts of Norway (not shown) did not show clear patterns with respect to position in the ordination diagram on one hand and sediment category SK1 or relative relief RR1 on the other.

Close examination of Fig. 26, designed to fine-tune criteria for separating plains from hills and mountains on the basis of relative relief and proportion of flat terrain (note that the values for Flat_a of 0.7 and 0.8 correspond to back-transformed area proportions of 100×100 m pixels with slope < 2° of approx. 20% and 35%, respectively; see the chapter ‘Transformation of analytic variables’) shows (i) that RR = 50 is too strict to be used as a separating criterion; most likely because the RR1 index is vulnerable to edge effects; (ii) RR = 100 is too liberal, affiliating with plains many OUs that do not comply with a common understanding of a plain (e.g. at Romerike, SE Norway); and (iii) that Flat_a = 0.75 is likely to be the one among the three potential criteria that provides the sharpest borderline (note that all large, but not all small blue rings in Fig. 26 have Flat_a > 0.75). In summary, Fig. 26 provides support for a division into plains on one hand and hills and mountains on the other hand, but fails to provide clear criteria for drawing the boundary.

Figs 27 and 28 take this one step further, exploring RR1 > 75 *and/or* Flat_a > 0.75 (i.e. more than 27% flat terrain) as potential criteria for separation of plains from hills and mountains. Placement of isolines for RR1 = 75 and Flat_a = 0.75 far to the left of the points of gravity in the clouds of blue and red dots is a result of the presence of many grey dots (OUs affiliated with IA) among the blue and red ones. This suggests that a demand on plains that RR1 < 75 *and* Flat_a > 0.75 will fail to separate non-valley inland OUs into two well-delimited groups that capture the overall pattern of variation in landscape-element composition. This calls for a more careful examination of OUs at the transition from plains to rugged landscapes.

The number of OUs that satisfy the condition ‘RR1 < 75 & Flat_a > 0.75’ is 396. The five OUs located far to the right in the ordination diagram (tentatively having characteristics of rugged landscapes; see Figs 26–27), are the only five out of the 396 OUs that have BP = 2, i.e. that contain plateau glaciers (ice caps). This suggests that OUs with prominent glaciers should be grouped with hill and mountain landscapes regardless if the surface is flat or sloping. Based upon Fig. 28, which shows the distributions of OUs with different *combinations* of relative relief and proportion of area with flat terrain in the ordination diagram, we examined the geographical distribution of OUs located in the borderland between plains and hills/mountains to answer three questions (Figs 29–34):

**Figs. 29–32.**
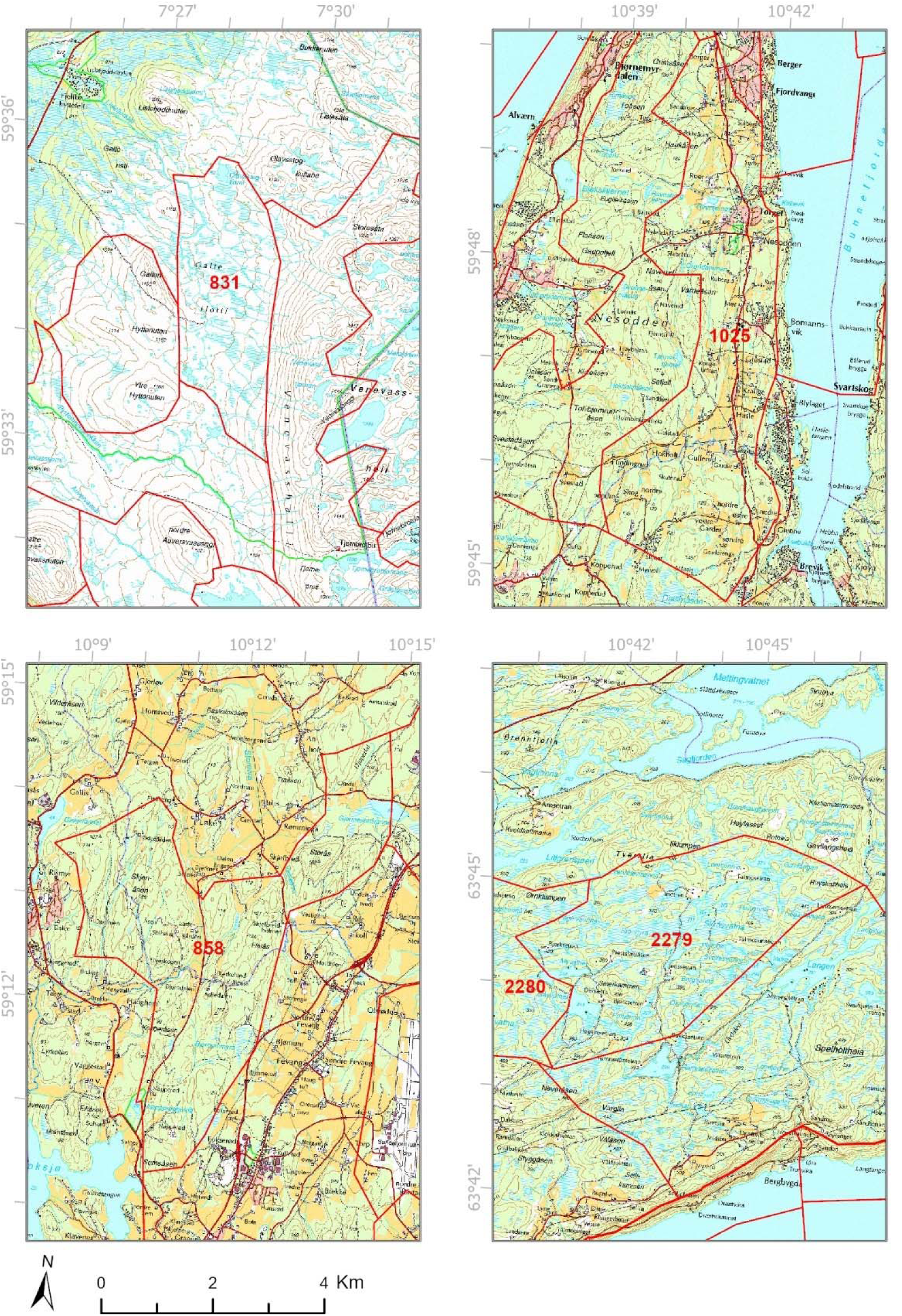
(29) Observation unit (OU) Id 831 Hovden protected landscape, Bykle municipality; Flat_a = 0.7008 and RR1 = 73. (30) OU Id 1025 Nesodden municipality, Flat_a = 0.7424, RR1 = 63). (31) OU Id 858 Torp, Sandefjord, Flat_a = 0.6768 and RR = 64. (32) OU Id 2279 Fosen, Leksvik, Flat_a = 0.6028 and RR = 73.

(i) Should smooth, gently sloping terrain be classified as plains? Our view is that it should not, because a plain should by definition be flat. Two examples of OUs of this kind that, in our opinion, should not be assigned to plains are Id 831 (Fig. 29; Aust-Agder: Bykle: Hovden landscape protection area), characterised by Flat_a = 0.7008 and RR1 = 73, and Id 1025 (Fig. 30; Akershus: Nesodden: comprising the eastern slope of the Nesodden peninsula), with Flat_a = 0.7424 and RR1 = 63. The values for Flat_a and RR1 for these OUs suggest that a stricter criterion for inclusion among plains than RR1 < 75 *or* Flat_a < 0.75 is needed.

(ii) Should rugged low-relief terrain be classified as plains? Our *a priori* answer to this question is also negative. Relevant examples are OU Id 858 (Fig. 31; Vestfold: Sandefjord: forested area west of Torp airport), with Flat_a = 0.6768 and RR = 64; and Id 2279 (Fig. 32) and Id 2280 (Fig. 33) situated on a plateau in the inland of Fosen peninsula (Nord-Trøndelag: Leksvik), with Flat_a = 0.6028 and RR = 73, and Flat_a = 0.5996 and RR1 = 75, respectively. This indicates that RR < 75 is not a sufficient criterion for characterising an OU as belonging to plains.

**Figs. 33–34.**
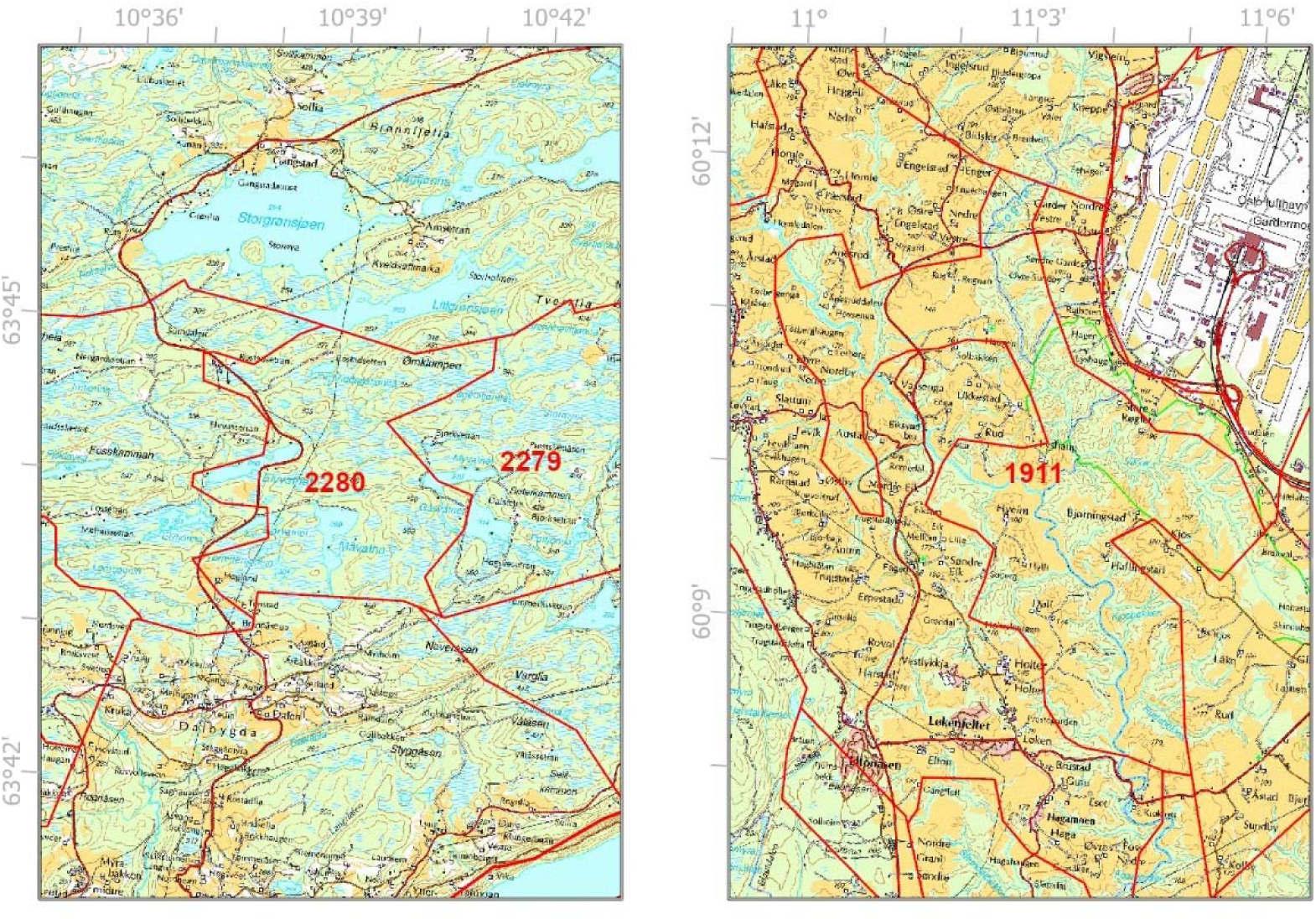
(33) Observation unit (OU) Id 2280 Fosen, Leksvik, Flat_a = 0.5996 and RR1 = 75. (34) OU Id 1911, Gardermoen, Flat_a = 0.6522; RR1 = 47.

(iii) Should sediment plains with ravines be classified as plains? Our immediate answer is ‘yes’, provided that the ravine valleys are not particularly deep. OU Id 1911 (Fig. 34; Akershus: Ullensaker: Gardermoen, just S of the airport) exemplifies this (Flat_a = 0.6522 and RR1 = 47). This may indicate that RR1 < 50 may serve as an absolute criterion for assigning OUs to plains.

Fig. 28 shows how 520 inland OUs, assigned to an inland plain candidate major type IS by four different criteria, are distributed in the ordination diagram. Ordered hierarchically, these four criteria provide the following groups:

ISfr (307 OUs with Flat_a > 0.734 and RR < 50)
ISf (120 OUs with Flat_a > 0.734 and RR > 50)
ISr (19 OUs with RR < 50 and Flat_a < 0.734)
ISx (74 OUs with 0.691 < Flat_a < 0.734 and 50 < RR < 75)

Of the remaining inland OUs, 1012 belonged to inland valleys and 1618 to inland hills and mountains.

*Conclusion*. The analyses support separation of valleys from other inland landscapes on the basis of the dominant terrain form (valley or non-valley). The analyses also support a tentative separation of inland plains from inland hills and mountains provided that at least one of the following criteria is met: (i) Proportion of area occupied by pixels with flat terrain (slope < 2°) > 25% (Flat_a > 0.734); (ii) RR1 < 50, or (iii) the combination of 20–25% flat terrain (0.691 < Flat_a < 0.734) and RR1 < 75.

#### Ordination of IS520 – the inland plains data subset

Two confirmed ordination axes were found for the IS520 data subset (Table 15). The best two-dimensional GNMDS solution (Fig. 36), identified from 1000 starting configurations, was used for further analysis and interpretation despite considerable instability, as evident from the high Procrustes SS value of 3.871 and the four unstable OUs. Both axes had gradient lengths within the same range (i.e. relative length > 0.8× that of the longest axis). No visible artefacts could be seen in the ordination diagrams (Figs 35–36). Very strong correlations between GNMDS axis 1 and land-use variables (infrastructure as well as agriculture, cf. Table 16), support the interpretation that the main gradient in landscape-element composition primarily expresses variation in characteristics related to human influence, and to a gradient from lowlands to mountain plains. Furthermore, the strong correlation between this axis and the proportion of OU area covered by marine deposits (Kmar_a) may indicate an underlying geo-ecological structure related to dominant soil type (Fig. 36, Table 16). The 520 OUs were distributed onto soil classes (JK1) as follows; B (exposed bedrock): *n* = 60; E (glaciofluvial deposits): *n* = 37; H (marine deposits): *n* = 63; M (thick layer of till): *n* = 306; 0 (unsorted sediments): *n* = 54.

**Figs. 35–36.**
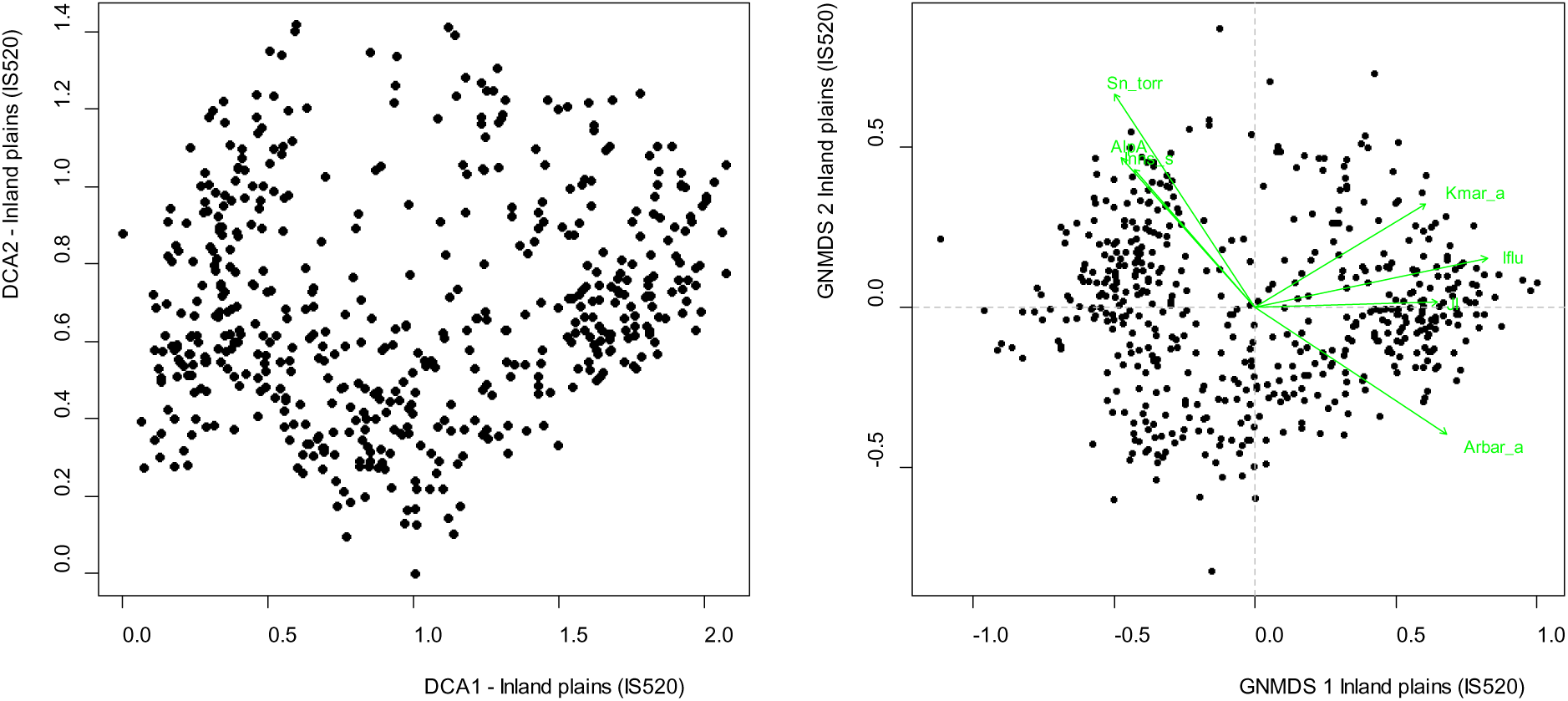
DCA and GNMDS ordinations of the IS520 data subset (520 observation units in inland plains), axes 1 and 2. (35) DCA ordination, axes are scaled in standard deviation (S.D.) units. (36) GNMDS ordination, axes are scaled in half-change (H.C.) units. Selected variables are represented by green arrows pointing in the direction of maximum increase of the variable (relative lengths of arrows are proportional to the correlation between the recorded variable and the ordination axis). Variable names are abbreviated according to Tables 4 and 5.

**Table 15.**
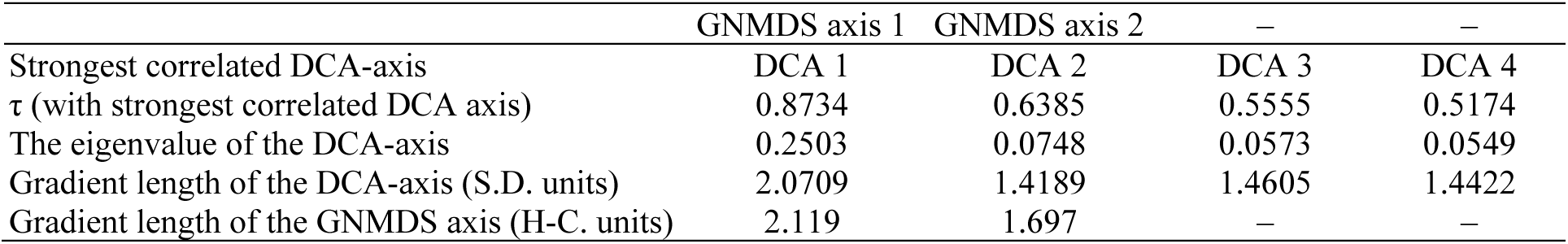
Kendall’s rank correlation coefficients (τ), eigenvalues and gradient lengths for parallel DCA- and GNMDS-ordinations of the subset IS520 Inland plains. S.D. units = standard deviation units; H.C. units = half-change units.

**Table 16.** Kendall’s rank correlation coefficients (τ) between GNMDS ordination axes 1–2 for the inland plains data subset (IS520) and the 9 primary key variables (rows 1–10) and the 65 analytic variables used to characterise the 520 observation units.

| Variable |  | GNMDS1 | GNMDS 2 |
| --- | --- | --- | --- |
| RR1 | Relief | –0.0035 | 0.0018 |
| FIA | Amount of flat land | 0.0755 | –0.0177 |
| TPI1 | Peaks | –0.0920 | 0.0236 |
| AlpA | Boreal/alpine landscape | –0.5498 | 0.2980 |
| II | Abundance of freshwater lakes | –0.1585 | 0.1631 |
| MI | Abundance of mire | –0.1605 | –0.1245 |
| IfI | Amount of infrastructure (OI) | 0.7527 | 0.0612 |
| IfIu | Amount of infrastructure | 0.7349 | 0.0851 |
| JI | Agricultural land-use intensity | 0.6718 | 0.0579 |
| Flat_a | AA flat terrain | 0.0756 | –0.0176 |
| Ktkmo_a | AA thick layer of till | –0.3273 | –0.2524 |
| Tpi1l_a | AA depressions | 0.1772 | 0.0636 |
| Tpi1h_a | AA convex terrain | 0.0228 | 0.0915 |
| Tpi1_mp | Terrain form TPI1, numerical, mean | –0.0905 | 0.0299 |
| River_a | AA large river | 0.0616 | 0.0365 |
| Steep_a | AA steep terrain/slope | 0.1156 | 0.0550 |
| Kskred_a | AA landslide soil | 0.0273 | 0.1240 |
| Rug3_m | Terrain ruggedness VRM3, mean | 0.1672 | 0.1341 |
| Rr1_m | Altitudinal range | –0.0034 | 0.0018 |
| Brich_a | AA lime-rich bedrock | 0.1359 | 0.1256 |
| Bpoor_a | AA lime-poor bedrock geology | 0.1764 | 0.0722 |
| Bplu_a | AA plutonic rock | 0.0792 | 0.4321 |
| Bvul_a | AA volcanic rock | 0.0021 | –0.1149 |
| Bavstn_a | AA sedimentary rock | –0.2114 | –0.0306 |
| Bomd_a | AA metamorphic rock | 0.0810 | –0.1768 |
| Mire_a | AA mire | –0.1574 | –0.1130 |
| Meant_a | AA moderate slope | 0.0283 | 0.0259 |
| Kmar_a | AA marine deposits | 0.5373 | 0.1420 |
| Klac_a | AA lacustrine deposits | 0.1678 | 0.0131 |
| Kelv_a | AA glaciofluvial deposits | 0.2448 | –0.1105 |
| Lake_a | AA freshwater lake | –0.2566 | 0.1766 |
| Innoy_s | Ant freshwater lake islands | –0.1161 | 0.2080 |
| Er_m | Hypsographic index, ER, mean | –0.0193 | –0.0562 |
| R_net_a | AA river | 0.1073 | 0.1105 |
| Inns_s | Number of lakes | –0.4176 | 0.2210 |
| Asp_s_a | AA south facing terrain | 0.0952 | –0.0133 |
| Asp_n_a | AA north facing terrain | –0.0837 | 0.1008 |
| Dismire | Distance to mire, mean | 0.1640 | 0.0929 |
| Dislake | Distance to lake, mean | 0.3416 | –0.2026 |
| Discoast | Distance to coast, mean | –0.0372 | –0.3935 |
| Guro_t_a | AA rugged terrain | 0.0260 | 0.1256 |
| Tpi6l_a | AA rugged terrain, TPI6 | 0.0939 | –0.0745 |
| Oyst_i | Inverse island size | 0.0911 | 0.2613 |
| Kbf_a | AA exposed bedrock | 0.2142 | 0.2479 |
| Bohei_a | AA boreal heaths | –0.0739 | 0.3032 |
| Arbar_a | AA coniferous forest | 0.5065 | –0.2672 |
| Arbla_a | AA mixed boreal forest | 0.5237 | –0.2393 |
| Arlov_a | AA deciduous forest | 0.4541 | –0.1456 |
| Sn_imp | AA impediment | –0.2524 | 0.0795 |
| Sn_flekk | AA patchy open treeless area | –0.4511 | 0.3324 |
| Sn_lav | AA lichen heath | –0.1880 | 0.1521 |
| Sn_torr | AA dry heath/open areas | –0.5502 | 0.3872 |
| Sn_frisk | AA moist/fresh heath/open areas | –0.3939 | 0.3731 |
| Sn_ureg | AA unregistered heath/open areas | 0.3418 | 0.2029 |
| Gab_nae | Ant commercial buildings | 0.6440 | 0.1072 |
| Gab_fi | Ant fisheries-related buildings | 0.1206 | 0.0536 |
| Gab_a | AA built-up | 0.7353 | 0.0787 |
| Vmag_a | AA regulated magazine | 0.0728 | 0.0440 |
| Vei_b_a | AA road | 0.6486 | 0.0814 |
| Sti_a | AA trail, path | 0.0360 | 0.0972 |
| Setr_s | Ant summer mountain pasture | 0.1828 | −0.1441 |
| Sefr_a | AA old buildings | 0.6985 | 0.0319 |
| Rd_anl_a | AA reindeer husbandry facilities | −0.1455 | −0.0049 |
| Build_a | AA built-up area | 0.5129 | 0.0694 |
| City_a | AA town/city area | 0.1231 | 0.0277 |
| Lled_a | AA power lines | 0.6016 | 0.1013 |
| Cul_u_s | number of cultural heritage sites outdoors | 0.2293 | 0.0530 |
| Cul_t_s | number of technical heritage sites | 0.2179 | 0.0560 |
| Cul_k_s | number of church ruins | 0.3990 | 0.0967 |
| Cul_b_s | number of ancient rock art sites | 0.2097 | 0.0723 |
| Cul_a_s | number of archaeological heritage sites | 0.5907 | 0.1401 |
| Abygg_a | AA large buildings | 0.5149 | 0.0693 |
| Araaf_a | AA open areas | 0.4622 | 0.0455 |
| Arfull_a | AA arable land | 0.6554 | 0.0255 |
| Arover_a | AA surface cultivated land | 0.2741 | 0.1382 |

Further exploration of the patterns expressed in the GNMDS ordination (Figs 37–40) revealed two almost non-overlapping groups of OUs: (1) plains on coarse sediments and/or bare rock (JK1 classes B, M and 0); and (2) plains on fine sediments (JK1 classes E and H). Fine-sediment plains formed a distinct cluster near the high-score end of GNMDS axis 1 (Fig. 40), also characterised by high values for the infrastructure (Iflu; Fig. 37) and agricultural land-use indices (Fig. 38). This shows that agriculture and other types of human exploitation are more or less confined to fine-sediment plains. The NiN principle that major (landscape) types shall differ with respect to complex-variable groups suggests splitting of inland plains into two major types.

**Figs. 37–40.**
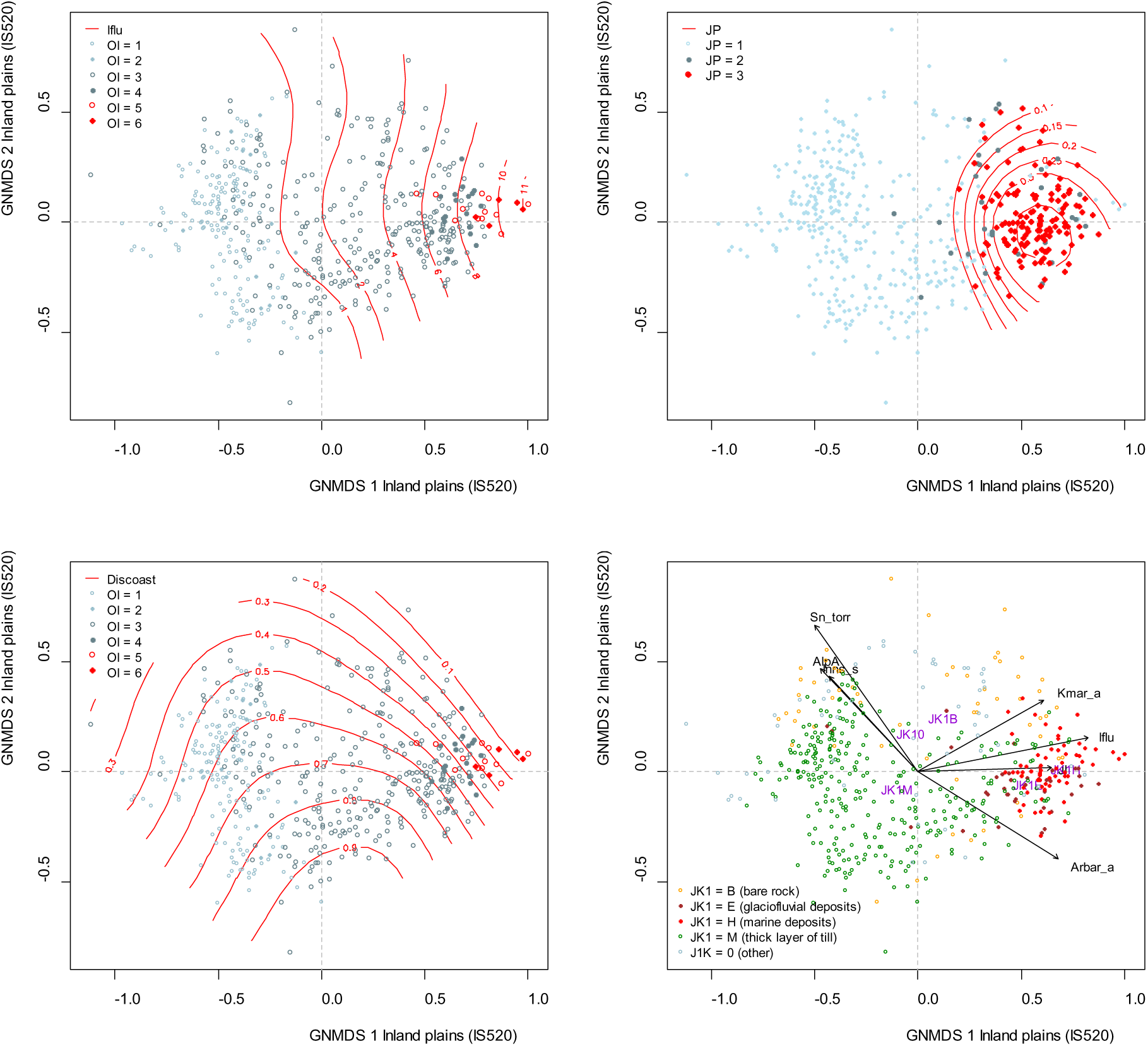
Distribution of observation units with specific properties in the GNMDS ordination diagram, axes 1 and 2, for the inland plains data set (IS520). Axes are scaled in half-change (H.C.) units. Selected variables are represented by arrows pointing in the direction of maximum increase of the variable (relative lengths of arrows are proportional to the correlation between the recorded variable and the ordination axis). Variable names are abbreviated according to Tables 4 and 5. (37) Isolines for the infrastructure index (IfIu); OI = stepwise variation in amount of infrastructure from low (OI 1–2) via medium OI 3–4) to high amount of infrastructure, including towns and/or cities (OI 5–6). (38) Isolines (red) for agricultural land-use intensity (JP). JP = stepwise variation in agricultural land-use intensity from low (JP 1) via medium (JP 2) to high (JP 3). (39) Variation in distance to coast (Discoast) and amount of infrastructure (OI). (40) Variation in soil types/sediment sorting (see legend).

Fig. 40 indicates that marine sediments have a particularly strong structuring effect on landscape-element composition. This is substantiated by examination of OUs dominated by bare rock (JK1 = B) and co-dominated by marine sediments (JK2 = M), which cluster with JK1 = M OUs.

*Conclusion*. The results lend support to a division of inland plains into two major types; (1) fine-sediment plains (dominated by marine and glaciofluvial deposits); and (2) ‘other plains’. Based on ordination results for the IS530 data subset, inland plains were assigned to fine-sediment plains if at least one of the following five criteria were satisfied: (1) proportion of area with marine deposits > 50%; (2) proportion of area with glaciofluvial sediments > 50%; (3) if no sediment type covers 50%, that the total proportion of area with fine sediments (marine deposits and glaciofluvial sediments) > 50%; (4) proportion of area with marine deposits > 20% *and* cover of exposed bedrock > 50%; (v) proportion of area with marine deposits > 10%, *and* no dominant soil class (i.e. no single soil class covers more than 50% of OU area).

Three OUs with positions in the ordination diagram that did not match their sediment composition were excluded from further analyses of the inland plains data subsets.

### IDENTIFICATION OF CLG CANDIDATES WITHIN EACH MAJOR-TYPE CANDIDATE

#### Coastal plains (KS)

A total of 457 OUs were affiliated with the coastal plain (KS) major-type candidate. All four axes in the 4-dimensional GNMDS ordination of the KS457 data subset were confirmed by the corresponding DCA axes (Table 17). The GNMDS ordination had Procrustes SS = 0.0363 and seven unstable OUs. The OUs were evenly spread out in the space spanned by DCA (Fig. 41) and GNMDS (Fig. 42) ordination axes 1 and 2, and no visible artefacts could be observed. Gradient lengths of both GNMDS and DCA axes and eigenvalues of the latter showed three relatively strong axes (i.e. relative length > 0.7× that of the longest axis; Table 17). Correlations between key and analytic variables on one hand, and GNMDS axis 1 scores on the other, supported by vector (Figs 42–43) and isoline diagrams (Figs 44–49) revealed that the first axis was very strongly related to land-use intensity (infrastructure and agricultural land use; see Figs 44 and 45, respectively), with strong relationships also to the geo-ecological gradient from inner to outer coast (Fig. 47, Table 18).

**Fig. 41.**
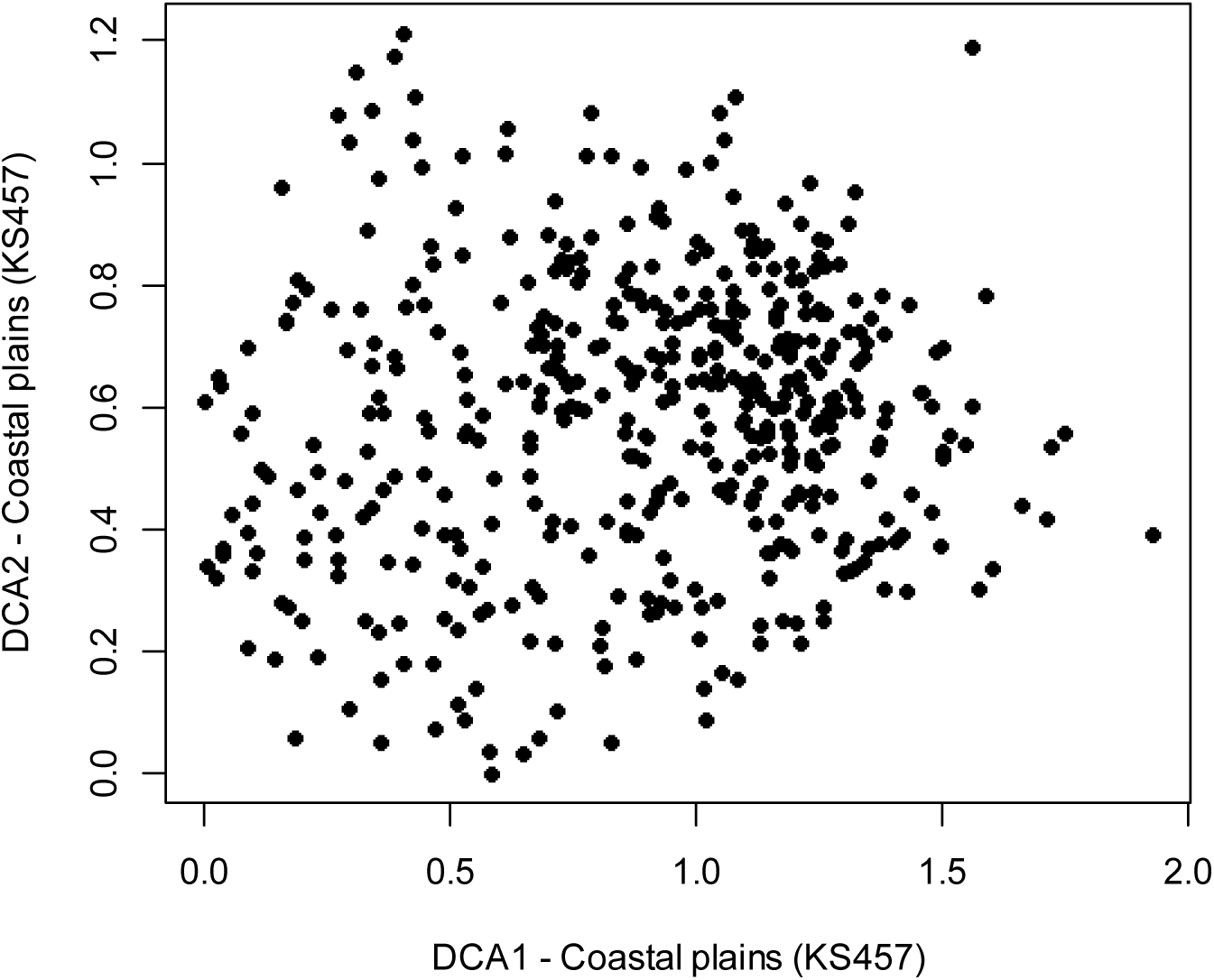
DCA ordination of the KS457 data subset (457 observation units in coastal plains), axes 1 and 2. Axes are scaled in standard deviation (S.D.) units.

**Figs. 42–45.**
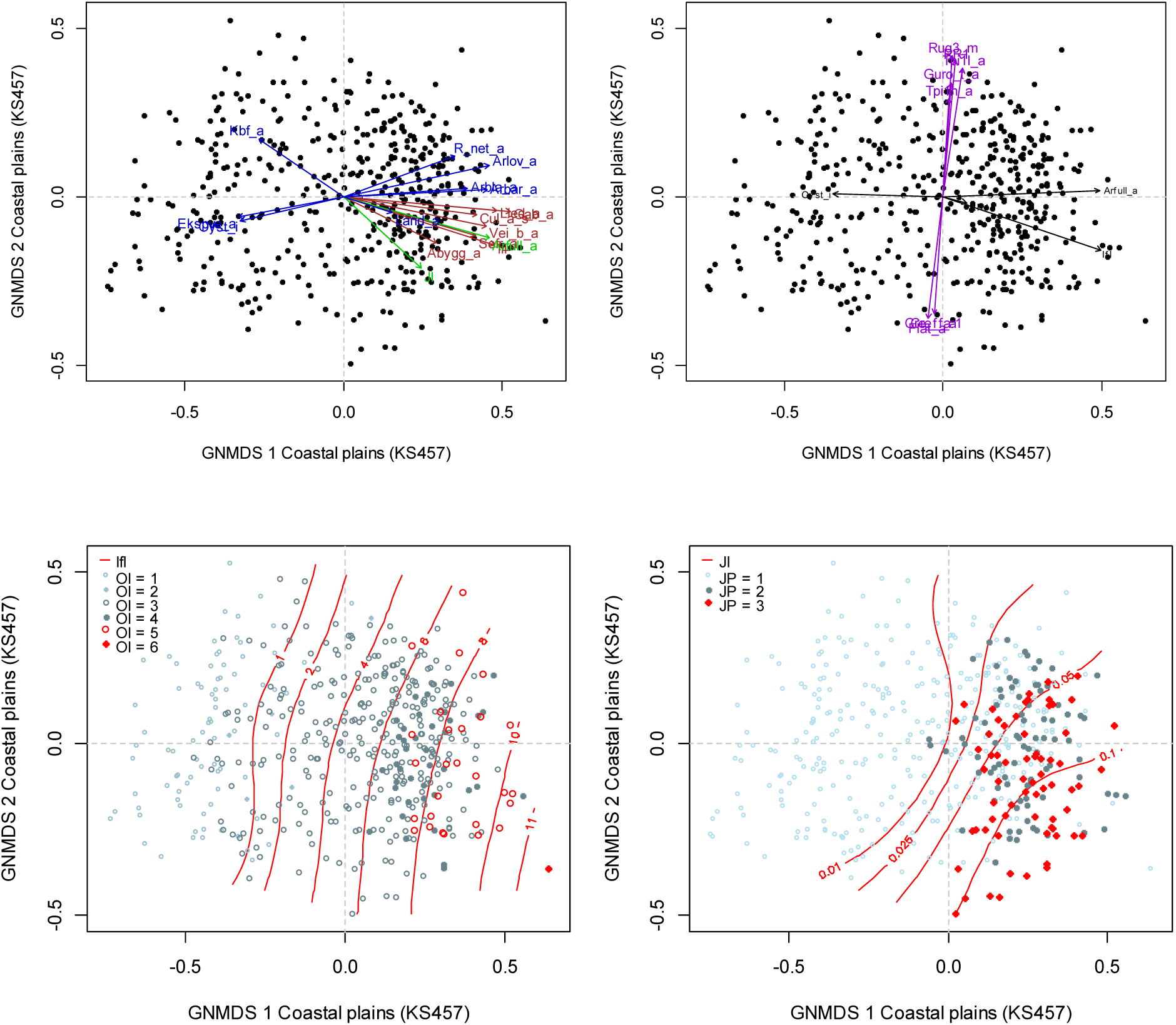
Distribution of observation units with specific properties in the GNMDS ordination diagram, axes 1 and 2, for the coastal plains data set (KS457). Axes are scaled in half-change (H.C.) units. (42–43) Selected variables are represented by arrows pointing in the direction of maximum increase of the variable (relative lengths of arrows are proportional to the correlation between the recorded variable and the ordination axis). Variable names are abbreviated according to Table 5. (44) Isolines (red) for the infrastructure index (Ifl). OI = stepwise variation in amount of infrastructure from low (OI 1–2) via medium (OI 3–4) to high amount of infrastructure, including towns and/or cities (OI 5–6). (45) Isolines (red) for agricultural land-use intensity (JI). JP = stepwise variation agricultural land-use intensity from low (JP 1) via medium (JP 2) to high (JP 3).

**Table 17.** Kendall’s rank correlation coefficients (τ), eigenvalues and gradient lengths for parallel DCA and GNMDS ordinations of the subset KS457 Coastal plains. S.D. units = standard deviation units; H.C. units = half-change units.

|  | GNMDS axis 1 | GNMDS axis 2 | GNMDS axis 3 | GNMDS axis 4 |
| --- | --- | --- | --- | --- |
| Strongest correlated DCA axis | DCA 1 | DCA 2 | DCA 3 | DCA 4 |
| $\tau$ (with strongest correlated DCA axis) | 0.8656 | 0.7326 | 0.5307 | 0.4988 |
| DCA-axis eigenvalue | 0.1272 | 0.0537 | 0.0458 | 0.0350 |
| DCA-axis gradient length (S.D. units) | 1.9271 | 1.2110 | 1.4277 | 1.0566 |
| GNMDS-axis gradient length (H.C. units) | 1.3798 | 1.0210 | 0.9796 | 0.9300 |

**Table 18.**
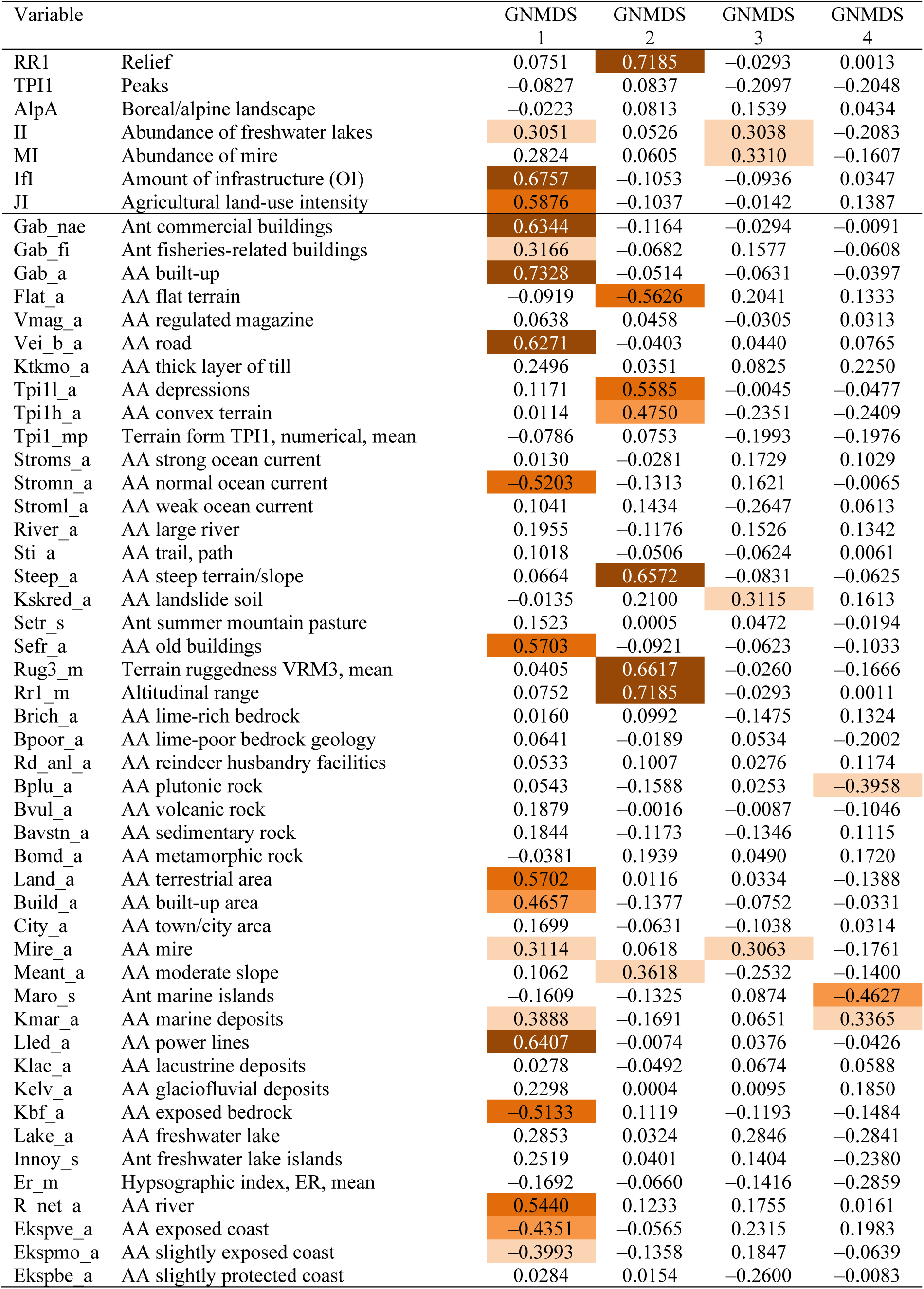
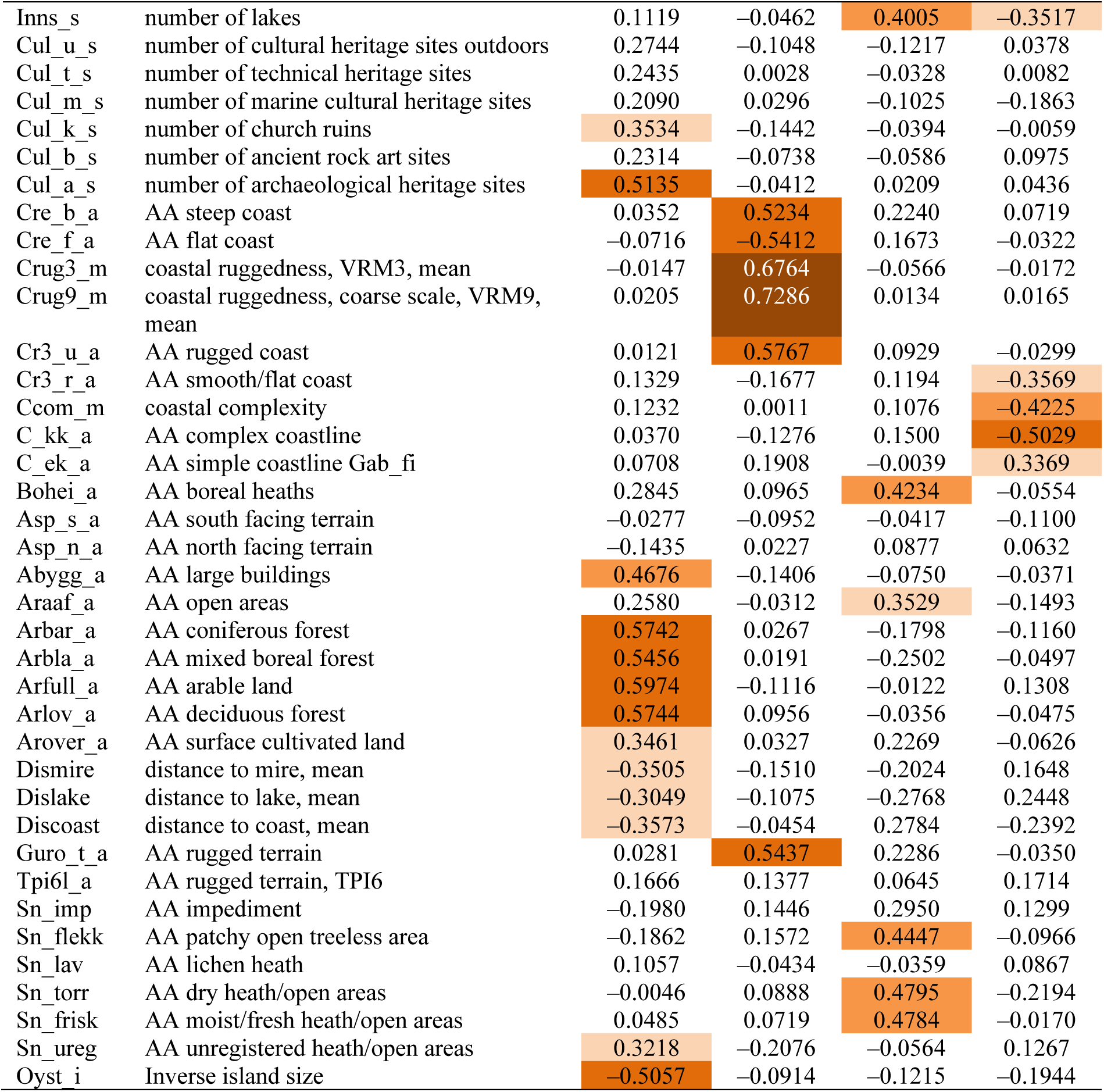
Kendall’s rank correlation coefficients (τ) between GNMDS ordination axes 1–4 for the coastal plains data subset (KS 457) and the 7 primary key variables and the 84 analytical variables used to characterise the 457 observation units.

The OU (Id 2916) that occupied the extreme position along GNMDS axis 1 (lower right corner, Fig. 44) was affiliated with OI class 6 (‘big city’), actually comprising parts of the centre of Oslo, the largest city in Norway. Most bio-ecological variables (functional category U) were strongly correlated with GNMDS axis 1, increasing from 0 (or near 0) for low GNMDS axis 1 scores towards higher GNMDS axis 1 scores (Fig. 42). An example of the pattern of variation of a category-U variable is given by the proportion of area covered by deciduous forest (Arlov_a; Fig. 46). The proportion of OU area covered by bare rock decreased with increasing island size (Fig. 48, Table 18). A similar pattern along GNMDS axis 1 was obtained for the LG-variable coastal/archipelago properties (SN; see Table 2). Fig. 47 shows a clear (inverse) island-size gradient from low to higher GNMDS axis 1 scores. The segregation of OUs along GNMDS axis 1 according to island size was particularly clear-cut for islands smaller than ca. 20 km^2^ (Oyst_i > 0.7; SN ≥ 3). This opens for the possibility that the complex gradient expressed on GNMDS axis 1 actually consists of more than one CLG.

**Figs. 46–49.**
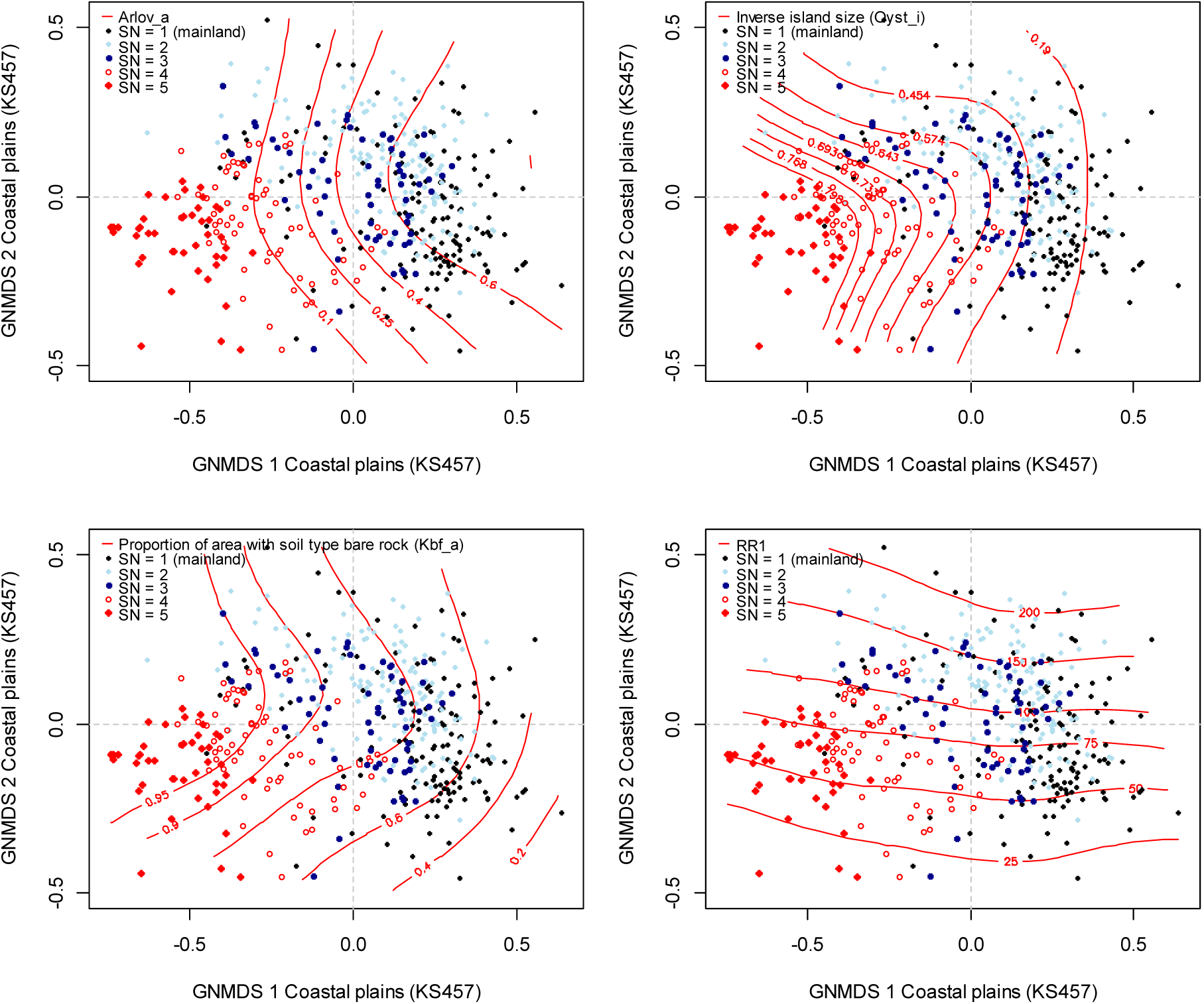
Distribution of observation units with specific properties in the GNMDS ordination diagram, axes 1 and 2, for the coastal plains data set (KS457). Axes are scaled in half-change (H.C.) units. Variation in coastal/archipelago properties (SN) and isolines are shown for (46) proportion of area with deciduous forest (Arlov); (47) inverse island size (Oyst_i); (48) proportion of area with bare rock (Kbf_a); and (49) mean values for relative relief (RR1). SN 1 = terrestrial coast-line landscape; SN 2 = large islands (> 20 km^2^); SN 3 = medium sized islands (1.5–20 km^2^); SN 4 = small islands (0.1–1.5 km^2^); SN 5 = skerries/islets (< 0.1 km^2^).

Among the 69 OUs in the KS457 data subset without infrastructure (IfI = 0), 56 possessed a largest island that was smaller than 1 km^2^. Islands with extensive infrastructure were generally much larger; an exception being Id 4038 which had Oyst_i = 0.8647 and IfI = 9.34. This OU comprises the SW part of Nøtterøy (Vestfold), which has been subjected to extensive summer-cabin development.

Figs 44 and 45 show that agricultural land use is conditioned on a minimum of infrastructure; JP ≥ 2 implies OI ≥ 3. A closer examination revealed a unimodal relationship between JI and IfI (Fig. 50), which implies that peak agricultural land-use intensity was associated with a well-developed infrastructure, decreasing towards cities (Figs 44–45; Fig. 50).

**Fig. 50.**
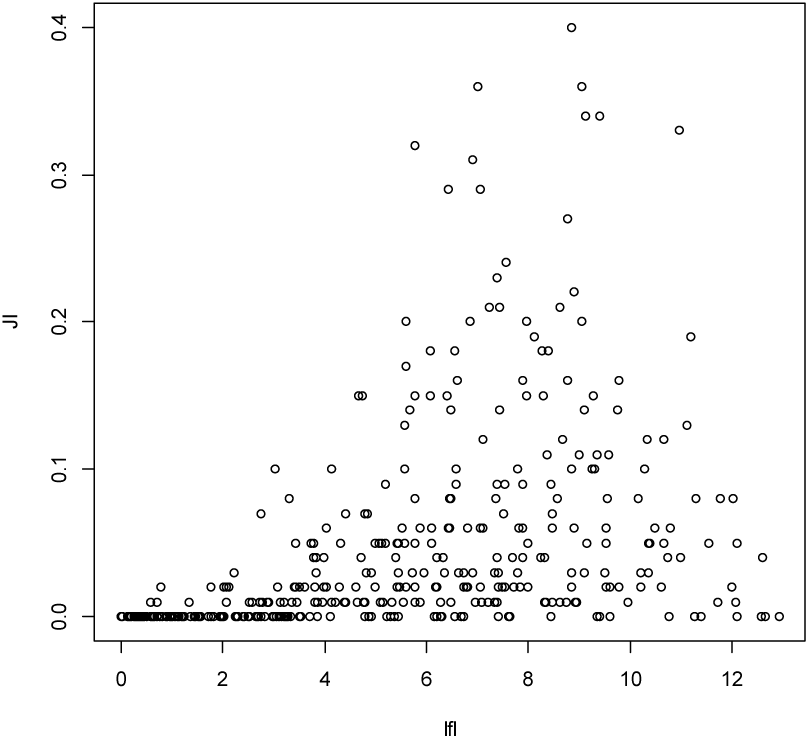
Relationship between values for the infrastructure index (Ifl) and the agricultural land-use intensity index (JI) for the 457 observation units in coastal plains (KS457).

The second GNMDS axis was strongly related to terrain shape; values of relative relief, ruggedness and related variables increased towards high GNMDS axis 2 scores (Fig. 43, 49). Furthermore, the proportions of flat vs steep 100×100 m pixels decreased and increased, respectively, along this axis. The strong signal from topographic variables in coastal plains (Fig. 49) at a first glance contrasts the general idea of a plain.

GNMDS axis 3 was correlated, but not strongly, with variables that expressed presence and areal cover of mires, lakes and boreal heaths (Fig. 51; Table 18). Mires and lakes became more prominent with increasing island size and were more or less absent from small islands (Figs 52–53).

**Figs. 51–54.**
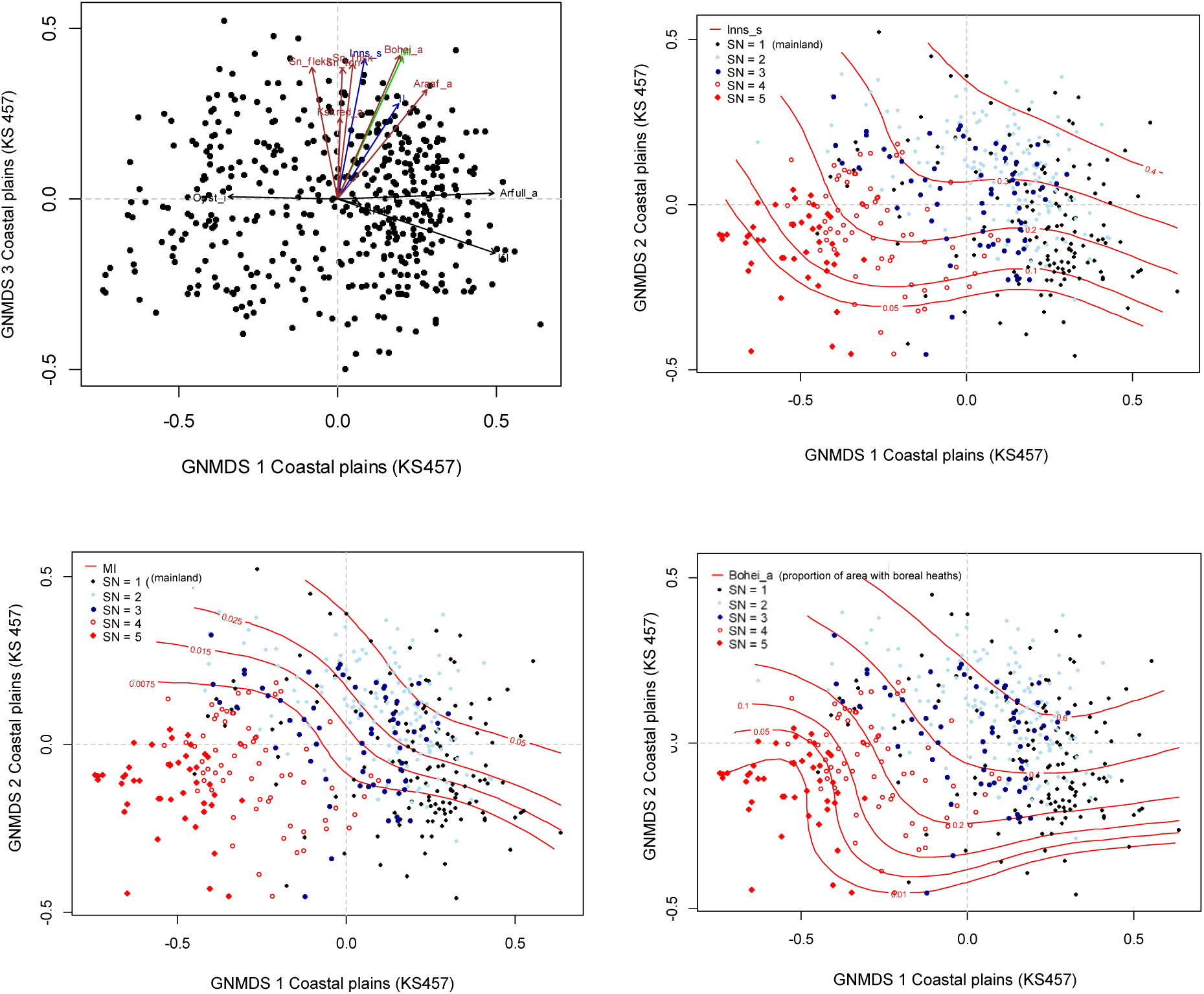
Distribution of observation units with specific properties in the GNMDS ordination diagram, axes 1 and 2, for the coastal plains data set (KS457). Axes are scaled in half-change (H.C.) units. (51) Selected variables are represented by arrows pointing in the direction of maximum increase of the variable (relative lengths of arrows are proportional to the correlation between the recorded variable and the ordination axis). Variable names are abbreviated according to Table 4 and 5. (52–54) Variation in coastal/archipelago properties (SN) and isolines for (52) the number of lakes (Inns_s); (53) the mire index (MI); and (54) the proportion of area covered by boreal heaths (Bohei). SN 1 = terrestrial coast-line landscape; SN 2 = large islands (> 20 km^2^); SN 3 = medium sized islands (1.5–20 km^2^); SN 4 = small islands (0.1–1.5 km^2^); SN 5 = skerries/islets (< 0.1 km^2^).

GNMDS axis 4 was correlated with variables that express coastal complexity, e.g. the number of marine islands (Table 18, Fig. 55). The noticeable correlation of GNMDS axis 4 with proportion of area covered by marine deposits illustrates that sediments are mainly deposited on larger land units (Fig. 55).

**Fig. 55.**
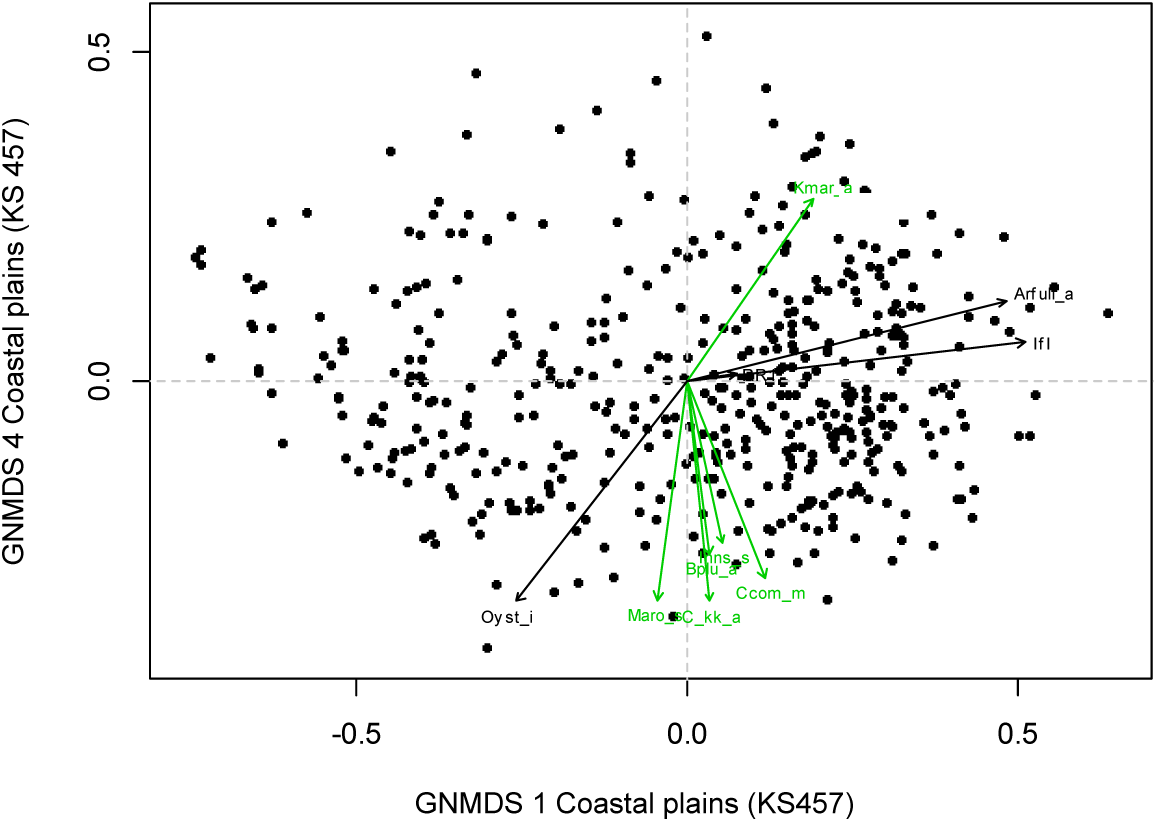
Distribution of observation units with specific properties in the GNMDS ordination diagram, axes 1 and 4, for the coastal plains data set (KS457). Axes are scaled in half-change (H.C.) units. Selected variables are represented by arrows pointing in the direction of maximum increase of the variable (relative lengths of arrows are proportional to the correlation between the recorded variable and the ordination axis). Variable names are abbreviated according to Tables 4 and 5.

Variation in dominant soil/sediment class within the KS457 data subset was explored to assess if coastal plains, like inland plains, should be divided further. The 457 OUs were distributed onto dominant (> 50% cover) soil classes (JK1) as follows; B (> 50% exposed bedrock): *n* = 346; H (marine deposits): *n* = 63; 0 (unsorted sediments): *n* = 61. In addition, 83 OUs belonged to JK2-class (> 25% cover) H. The distribution of JK1 classes in the GNMDS ordination diagram for axis 1 and 2 showed a weak relationship between JK-class and GNMDS axis 1 (Fig. 56) and did not support further division of coastal plains.

**Fig. 56.**
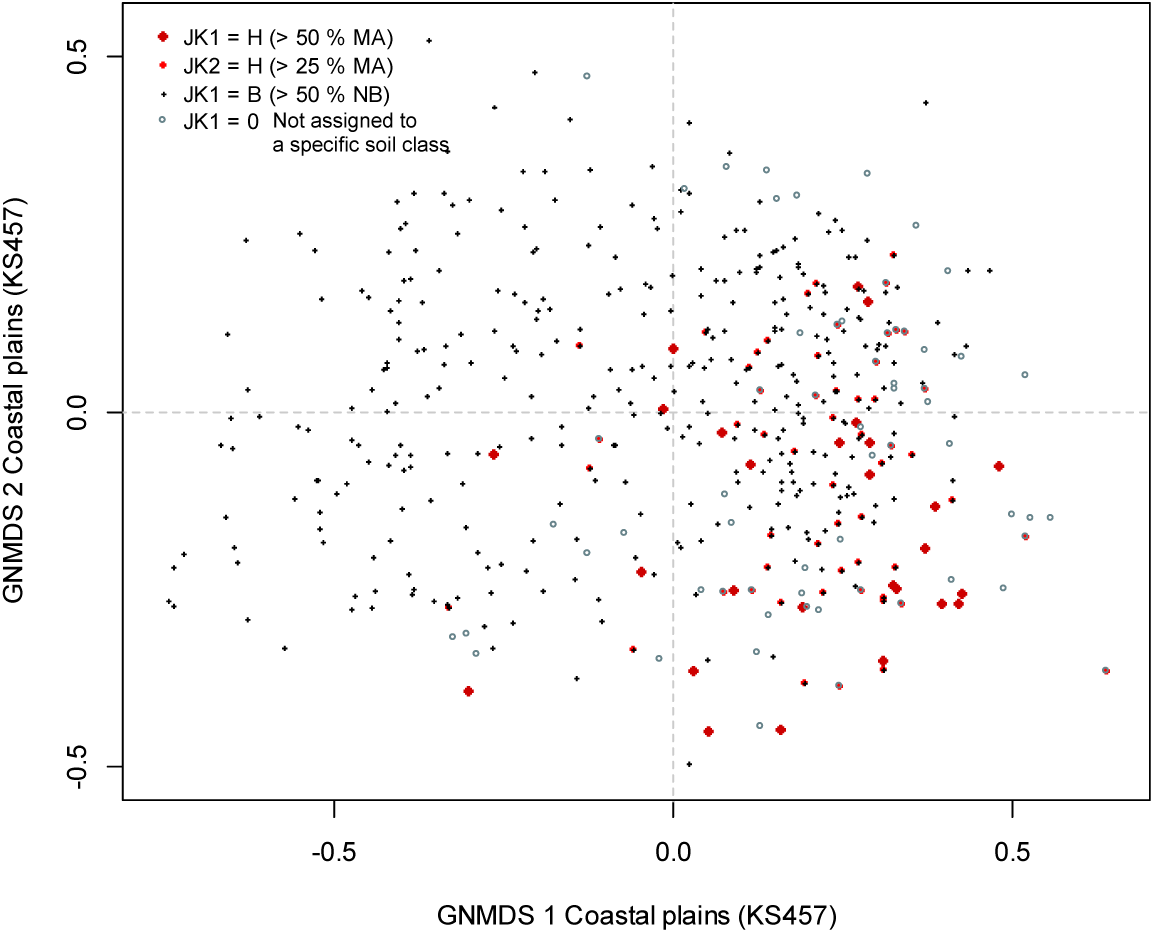
Distribution of observation units belonging to different sediment categories in the GNMDS ordination diagram, axes 1 and 2, for the coastal plains data set (KS457). Axes are scaled in half-change (H.C.) units. JK = primary soil/sediment class. Percentages refer to proportion of area: H = marine deposits (MA); NB = bare rock.

Parallel RDA analyses with soil-class factor variables and relevant continuous analytic variables as alternative ways of representing variation in dominant soil properties showed that the analytic variables Kmar_a and Kbf_a explained more variation in the KS457 data subset (0.0600 and 0.0597) than the factor variables JK1 and JK2 (0.0429 and 0.0344, respectively). We therefore used the analytic variables to represent the CLG candidate soil/sediment class.

*Conclusion*. Based on ordination analyses and interpretation, we identified the following 9 CLG candidates in the three functional variables categories G, U and A (thematically slightly marginal variables are highlighted in grey; the absolute value of the correlation coefficient with the relevant GNMDS axis is given in parentheses):

(G1) Relief (**RE**; GNMDS axis 2): Crug9_m (0.7286) RR1 (0.7185), Crug3_m (0.6764), Rug3_m (0.6617), Steep_a (0.6572), Cr3_u_a (0.5767), Flat_a (0.5626), Tpi1l_a (0.5585), Guro_t_a (0.5437), Cre_f_a (0.5412), Cre_b_a (0.5234), Tpi1h_a (0.4750)
(G2) Inner-outer coast (**IYK**; GNMDS axis 1/2): Land_a (0.5702), Stromn_a (0.5203), Oyst_i (0.5057), R_net_a (0.5440), Ekspve_a (0.4351)
(G3) Abundance of wetlands (**VP**; GNMDS axis 1/3/4: Inns_s (0.4005), Dismire (0.3505), MI (0.3310), II (0.3051), Dislake (0.3049)
(G4) ‘True archipelago properties’ (**SgP**; GNMDS axis 4): C_kk_a (0.5029), Maro_s (0.4627), Ccom_m (0.4225)
(G5) Soil/sediment type (**JA**; GNMDS axis 1/4): Kbf_a (0.5133), Kmar_a (0.3888)
(U1) Vegetation (forest) cover (**SkP**; GNMDS axis 1): Arlov_a (0.5744), Arbar_a (0.5742), Arbla_a (0.5456)
(U2) Open area cover (**AP**; GNMDS axis 3): Sn_frisk (0.4784), Sn_torr (0.4765), Sn_flekk (0.4447), Bohei_a (0.4234), Araaf_a (0.3529)
(A1) Infrastructure (**OI**; GNMDS axis 1): Gab_a (0.7328), IfI (0.6757), Gab_nae (0.6344), Lled_a (0.6407), Vei_b_a (0.6271), Sefr_a (0.5703), Cul_a_s (0.5135), Abygg_a (0.4676)
(A2) Agricultural land-use intensity (**JP**; GNMDS axis 1): Arfull_a (0.5974), JI (0.5876), Arover_a (0.3461)

#### Fjords and other coastal landscapes (KF)

A total of 353 OUs were affiliated with the major-type candidate fjords and other coastal landscapes (KS). The three axes in the 3-dimensional GNMDS ordination of the KF353 data subset were confirmed by DCA axes, with three strong axes in both ordinations (i.e. relative length > 0.792× that of the longest axis; Table 19). Initially, however, the 4-dimensional GNMDS was used as a basis for further analyses and graphical interpretation. Because the first three axes in 4-dimensional GNMDS are almost identical to the corresponding axes in 3-dimensional GNMDS, the figures and interpretation were not re-made for three dimensions. The GNMDS ordination had Procrustes SS = 0.0003 and no unstable OUs. Visible artefacts could neither be observed in the spaces spanned by DCA (Fig. 57) nor by GNMDS (Fig. 58) ordination axes 1 and 2, but the OUs were more spread out towards the periphery in both.

**Fig. 57.**
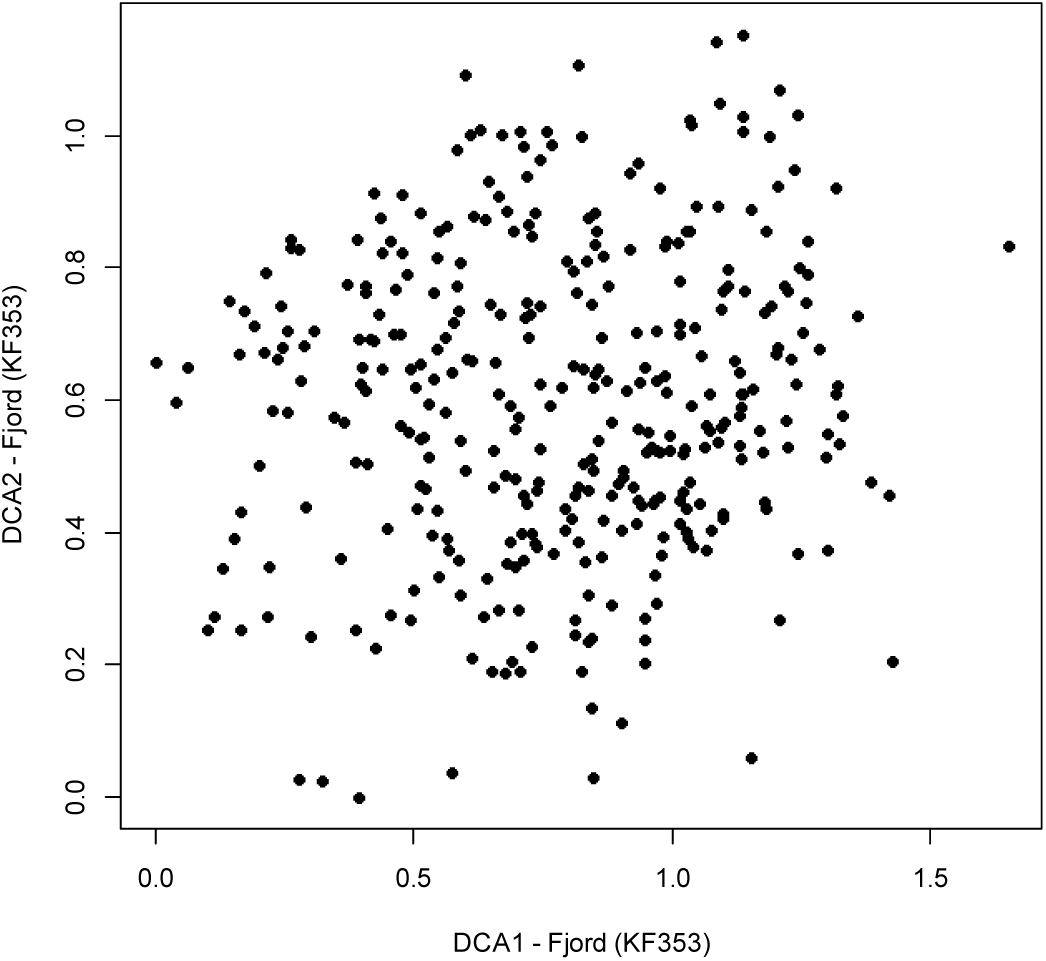
DCA ordination of the KF353 data subset (353 observation units in coastal fjords), axes 1 and 2, scaled in standard deviation (S.D.) units.

**Figs. 58–63.**
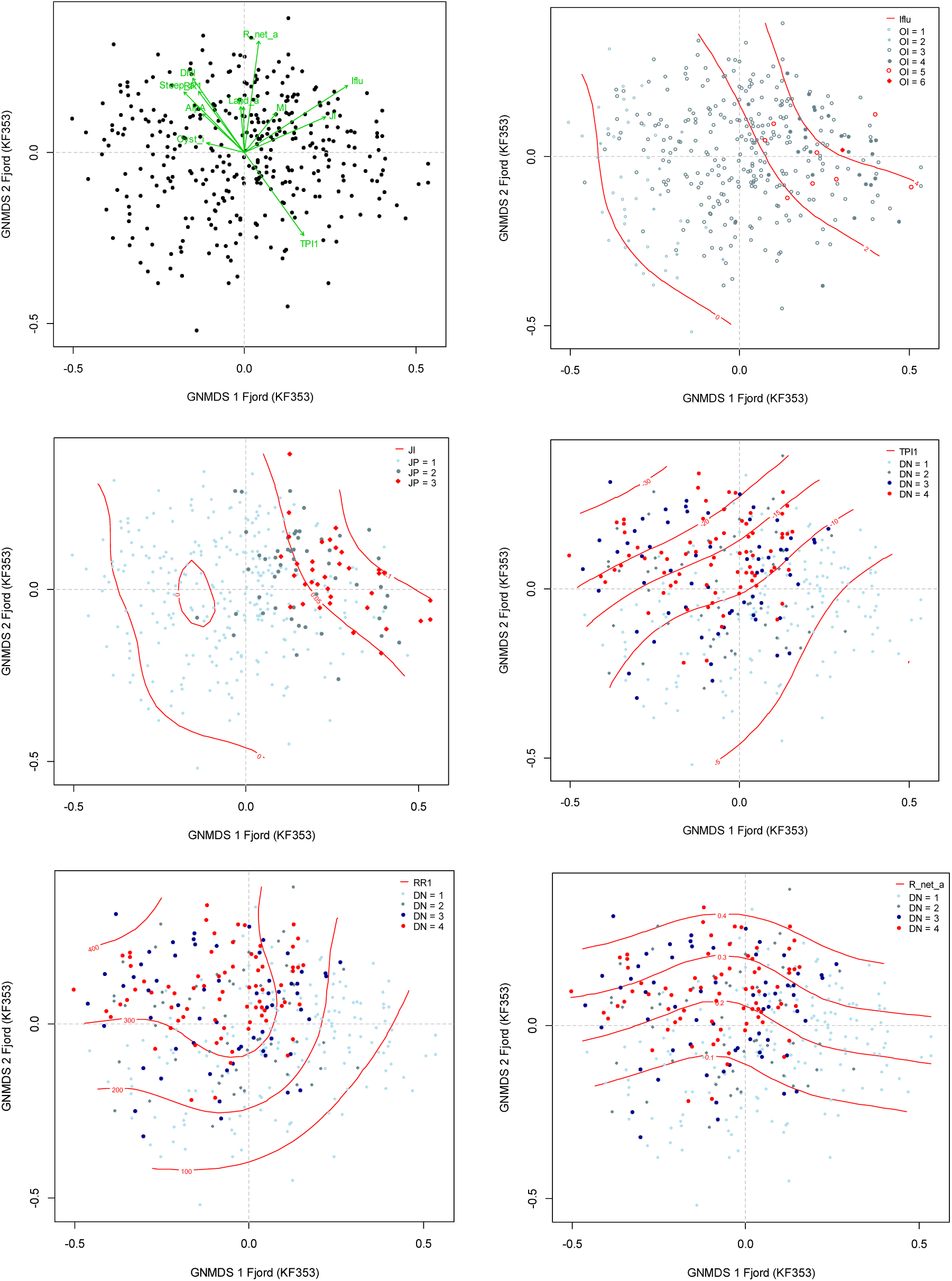
Distribution of observation units with specific properties in the GNMDS ordination diagram, axes 1 and 2, for the coastal fjords data set (KF353). Axes are scaled in half-change (H.C.) units. (58) Selected variables are represented by arrows pointing in the direction of maximum increase of the variable (relative lengths of arrows are proportional to the correlation between the recorded variable and the ordination axis). Variable names are abbreviated according to Table 4 and 5. (59) Isolines (red) for the infrastructure index (Ifl). OI = stepwise variation in amount of infrastructure from low (OI 1–2) via medium (OI 3–4) to high amount of infrastructure, including towns and/or cities (OI 5–6). (60) Isolines (red) for agricultural land-use intensity (JI). JP = stepwise variation in agricultural land-use intensity from low (JP 1) via medium (JP 2) to high (JP 3). (61–63) Valley shape (DN) and isolines (red) for (61) the terrain position index (TPI1); (62) relative relief (RR1); and (63) proportion of 100×100 m pixels with rivers (R_net_a). DN 1 = open valley; DN 2 = relatively open valley; DN 3 = relatively narrow and steep valley; DN 4 = narrow and steep-sided valley.

**Table 19.** Kendall correlation coefficients (τ), eigenvalues and gradient lengths for parallel DCA and GNMDS ordinations of the subset KF353 Coastal fjords. S.D. units = standard deviation units; H.C. units = half-change units.

|  | GNMDS axis 1 | GNMDS axis 2 | GNMDS axis 3 | – |
| --- | --- | --- | --- | --- |
| Strongest correlated DCA axis | DCA 1 | DCA 3 | DCA 2 | DCA 4 |
| $\tau$ (with strongest correlated DCA axis) | 0.7609 | –0.4842 | –0.4124 | – |
| DCA-axis eigenvalue | 0.0963 | 0.0463 | 0.0507 | 0.0377 |
| DCA-axis gradient length (S.D. units) | 1.6490 | 1.2387 | 1.1533 | 1.1055 |
| Correlation with axis in 4-dimensional GNMDS | 0.9683 | 0.9415 | –0.9137 | – |
| GNMDS-axis gradient length (H.C. units) | 1.0385 | 0.9130 | 0.8228 | – |

Correlations between key and analytic variables on one hand (Table 20), and GNMDS axis 1 scores on the other, supported by vector diagrams (Fig. 58) and isoline diagrams (Figs 59–60), revealed that the first axis was strongly related to land-use intensity (infrastructure and agricultural land use; Figs 59 and 60, respectively). Few cities (Fig. 59; OI = 5 & 6) were located in coastal fjords and there was hardly any difference in location along axis 1 between OI = 4, OI = 5 and OI = 6. OUs with moderate land-use intensity (OI = 3; Ifl between 6 and 12) were spread along the entire GNMDS1 axis. Agricultural land-use intensity was clearly related to infrastructure; the isolines for these variables followed the same pattern (Figs 59– 60).

**Table 20.** Kendall’s rank correlation coefficients (τ) between GNMDS ordination axes 1–3 for the coastal fjords data subset (KF353) and the 9 primary key variables (rows 1–9) and the 84 analytical variables used to characterise the 353 observation units.

| Variable |  | GNMDS1 | GNMDS 2 | GNMDS 3 |
| --- | --- | --- | --- | --- |
| DNI | Valley shape | –0.2935 | 0.3231 | –0.3939 |
| RR1 | Relief | –0.2866 | 0.2670 | –0.4495 |
| TPI1 | Peaks | 0.3221 | –0.3638 | 0.2842 |
| BrI | Glacier | –0.0752 | 0.1067 | –0.0619 |
| AlpA | Boreal/alpine landscape | –0.2752 | 0.3293 | 0.1110 |
| II | Abundance of freshwater lakes | –0.0275 | 0.2011 | 0.1375 |
| MI | Abundance of mire | 0.1478 | 0.2422 | 0.3736 |
| IfIu | Amount of infrastructure | 0.6132 | 0.2589 | –0.0495 |
| JI | Agricultural land-use intensity | 0.5870 | 0.1452 | –0.0108 |
| Gab_nae | Ant commercial buildings | 0.5774 | 0.1542 | –0.0941 |
| Gab_fi | Ant fisheries-related buildings | 0.2735 | 0.2353 | 0.0147 |
| Gab_a | AA built-up | 0.6417 | 0.1690 | –0.0689 |
| Flat_a | AA flat terrain | 0.2067 | –0.2209 | 0.3840 |
| Vei_b_a | AA regulated magazine | 0.5237 | 0.1729 | –0.0592 |
| Ktkmo_a | AA road | 0.1808 | 0.3641 | 0.0649 |
| Tpi1l_a | AA thick layer of till | –0.3384 | 0.3591 | –0.3152 |
| Tpi1h_a | AA depressions | –0.1050 | –0.0268 | –0.3931 |
| Tpi1_mp | AA convex terrain | 0.3149 | –0.3556 | 0.2785 |
| Stroms_a | Terrain form TPI1, numerical, mean | –0.0068 | –0.0181 | 0.0835 |
| Stromn_a | AA normal ocean current | –0.0456 | –0.2636 | 0.1268 |
| Stroml_a | AA weak ocean current | –0.0532 | –0.3031 | –0.0860 |
| River_a | AA large river | 0.2774 | 0.1697 | 0.0193 |
| Sti_a | AA trail, path | 0.0844 | 0.0622 | –0.0609 |
| Steep_a | AA steep terrain/slope | –0.2617 | 0.2264 | –0.5080 |
| Kskred_a | AA landslide soil | –0.0830 | 0.4301 | –0.1519 |
| Setr_s | Ant summer mountain pasture | 0.1925 | 0.0931 | –0.0536 |
| Sefr_a | AA old buildings | 0.5149 | 0.1793 | –0.1716 |
| Rug3_m | Terrain ruggedness VRM3, mean | –0.2982 | 0.2357 | –0.4482 |
| Rr1_m | Altitudinal range | –0.2867 | 0.2671 | –0.4496 |
| Brich_a | AA lime-rich bedrock | 0.2321 | –0.1740 | 0.0569 |
| Bpoor_a | AA lime-poor bedrock geology | 0.0328 | 0.2478 | –0.1588 |
| Rd_anl_a | AA reindeer husbandry facilities | –0.1202 | 0.0897 | 0.1179 |
| Bplu_a | AA plutonic rock | –0.2029 | 0.1144 | 0.0375 |
| Bvul_a | AA volcanic rock | 0.0043 | –0.0419 | –0.0784 |
| Bavstn_a | AA sedimentary rock | 0.1994 | –0.1468 | 0.0812 |
| Bomd_a | AA metamorphic rock | 0.1539 | –0.0321 | –0.0651 |
| Land_a | AA terrestrial area | 0.0599 | 0.5645 | 0.0099 |
| Build_a | AA built-up area | 0.4336 | 0.0801 | –0.1015 |
| City_a | AA town/city area | 0.1249 | –0.0100 | –0.0339 |
| Mire_a | AA mire | 0.1199 | 0.2207 | 0.4056 |
| Glac_a | AA glacier | –0.0469 | 0.1270 | –0.0526 |
| Meant_a | AA moderate slope | 0.1671 | –0.1395 | 0.3505 |
| Maro_s | Ant marine islands | 0.0180 | –0.0816 | –0.0714 |
| Kmar_a | AA marine deposits | 0.3925 | –0.0802 | 0.3169 |
| Lled_a | AA power lines | 0.4719 | 0.3002 | –0.0171 |
| Klac_a | AA lacustrine deposits | 0.0321 | 0.0646 | –0.0287 |
| Kelv_a | AA glaciofluvial deposits | 0.0124 | 0.0076 | 0.0159 |
| Kbf_a | AA exposed bedrock | –0.3874 | –0.3352 | –0.0415 |
| Lake_a | AA freshwater lake | –0.0440 | 0.2401 | 0.2569 |
| Innoy_s | Ant freshwater lake islands | –0.0245 | 0.1217 | 0.0756 |
| Er_m | Hypsographic index, ER, mean | 0.1069 | –0.2349 | –0.1375 |
| R_net_a | AA river | 0.1170 | 0.5457 | 0.0032 |
| Ekspve_a | AA exposed coast | -0.1582 | -0.0292 | 0.1631 |
| Ekspmo_a | AA slightly exposed coast | -0.2067 | -0.0668 | 0.2049 |
| Ekspbe_a | AA slightly protected coast | -0.0282 | -0.4804 | -0.0046 |
| Inns_s | number of lakes | -0.0660 | 0.3044 | 0.3180 |
| Cul_u_s | number of cultural heritage sites outdoors | 0.1059 | -0.0400 | -0.0399 |
| Cul_t_s | number of technical heritage sites | 0.1208 | 0.0519 | -0.1371 |
| Cul_m_s | number of marine cultural heritage sites | 0.1301 | -0.0385 | -0.2859 |
| Cul_k_s | number of church ruins | 0.3203 | 0.1190 | -0.1024 |
| Cul_b_s | number of ancient rock art sites | 0.1064 | 0.0033 | -0.0349 |
| Cul_a_s | number of archaeological heritage sites | 0.3940 | 0.1322 | 0.0966 |
| Cre_b_a | AA steep coast | -0.3209 | 0.2235 | -0.3240 |
| Cre_f_a | AA flat coast | 0.2128 | -0.0270 | 0.2524 |
| Crug3_m | coastal ruggedness, VRM3, mean | -0.3576 | 0.0482 | -0.4030 |
| Crug9_m | coastal ruggedness, coarse scale, VRM9, mean | -0.4056 | 0.1749 | -0.3871 |
| Cr3_u_a | AA rugged coast | -0.2631 | 0.1591 | -0.3325 |
| Cr3_r_a | AA smooth/flat coast | 0.2841 | 0.2182 | 0.2729 |
| Ccom_m | coastal complexity | 0.0143 | 0.2909 | -0.0169 |
| C_kk_a | AA complex coastline | 0.0497 | 0.0684 | 0.0033 |
| C_ek_a | AA simple coastline Gab_fi | -0.0701 | 0.2310 | -0.0837 |
| Bohei_a | AA boreal heaths | -0.1887 | 0.4684 | 0.3439 |
| Asp_s_a | AA south facing terrain | 0.0271 | 0.0265 | 0.0275 |
| Asp_n_a | AA north facing terrain | -0.0353 | 0.0182 | 0.0283 |
| Abygg_a | AA large buildings | 0.4378 | 0.0787 | -0.0947 |
| Araaf_a | AA open areas | 0.1487 | 0.3915 | 0.1526 |
| Arbar_a | AA coniferous forest | 0.4186 | -0.0363 | -0.3911 |
| Arbla_a | AA mixed boreal forest | 0.3152 | -0.0646 | -0.4176 |
| Arfull_a | AA arable land | 0.6094 | 0.1460 | -0.0271 |
| Arlov_a | AA deciduous forest | 0.3206 | 0.2897 | -0.0821 |
| Arover_a | AA surface cultivated land | 0.3180 | 0.2506 | 0.0249 |
| Dismire | distance to mire, mean | -0.1495 | -0.1830 | -0.2071 |
| Dislake | distance to lake, mean | 0.1137 | -0.3216 | -0.1229 |
| Discoast | distance to coast, mean | -0.1845 | 0.1603 | 0.3474 |
| Guro_t_a | AA rugged terrain | -0.3248 | 0.3130 | -0.3868 |
| Tpi6l_a | AA rugged terrain, TPI6 | -0.0185 | 0.1246 | 0.1364 |
| Sn_imp | AA impediment | -0.3329 | 0.3194 | 0.1489 |
| Sn_flekk | AA patchy open treeless area | -0.3361 | 0.2917 | 0.1959 |
| Sn_lav | AA lichen heath | -0.0227 | -0.0659 | -0.0706 |
| Sn_torr | AA dry heath/open areas | -0.2746 | 0.3152 | 0.2630 |
| Sn_frisk | AA moist/fresh heath/open areas | -0.2238 | 0.4015 | 0.4020 |
| Sn_ureg | AA unregistered heath/open areas | 0.3033 | 0.1759 | 0.0698 |
| Oyst_i | Inverse island size | -0.1837 | 0.0622 | 0.3072 |

The second GNMDS axis was strongly related to proportion of terrestrial area (Land_a), presence of river networks (R_net_a) and partially also to terrain shape (Table 20; Figs 58 and 63). The relatively short vector arrows for terrestrial area (Land_a) and river networks (R_net_a), running parallel to GNMDS axis 2, indicated relatively weak co-variation with other analysis variables.

GNMDS axis 3 was clearly related to relief (Table 20; Fig. 64). Figs 58 and 62 show that amount of infrastructure (IfIu) and agricultural land-use intensity (JI) on one hand, and the relief-related variables (DNI, RR1 and TPI1) on the other hand, were independent CLGs; the vector arrows were more or less perpendicular to each other. Fig. 65 show that narrow and steep-sided fjords (DN = 3–4) differed from other OUs. The relatively low number of OUs (*n* = 21) with high abundance of mire (MP = 2; Fig. 66) indicate that amount and cover of mires are less important properties in fjords. Number of lakes (Inns_s) and mires (Mire_a) were correlated with both of GNMDS axes 2 and 3, and strongly correlated with each other (τ = 0.4390). The weak signals from the variable ‘island size’ neither support a separation of the mainland from islands in fjords nor suggests that gradients from outer to inner coast are important in the fjord major type.

**Figs. 64–67.**
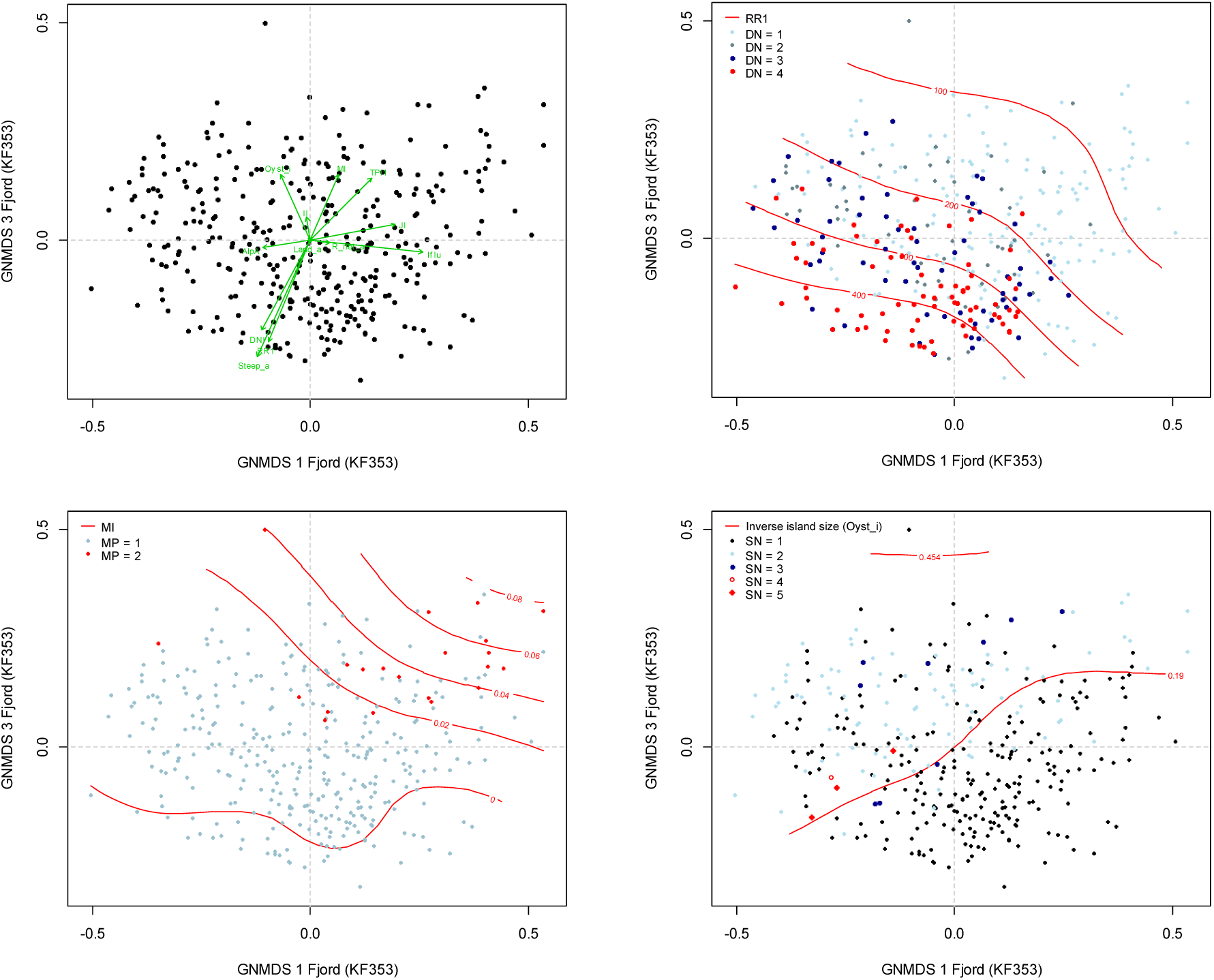
Distribution of observation units with specific properties in the GNMDS ordination diagram, axes 1 and 3, for the coastal fjords data set (KF353). Axes are scaled in half-change (H.C.) units. (64) Selected variables are represented by arrows pointing in the direction of maximum increase of the variable (relative lengths of arrows are proportional to the correlation between the recorded variable and the ordination axis). Variable names are abbreviated according to Tables 4 and 5. (65) Isolines (red) for relative relief (RR1). DN = variation in valley shape: DN 1 = open valley; DN 2 = relatively open valley; DN 3 = relatively narrow and steep valley; DN 4 = narrow and steep-sided valley. (66) Isolines for the mire index (MI). MP = variation in mire abundance: MP 1 = low abundance of mire; MP 2 = high abundance of mire. (67) Isolines for inverse island size (Oyst_i). SN = variation in coastal/archipelago properties: SN 1 = terrestrial coast-line landscape; SN 2 = large islands (> 20 km^2^); SN 3 = medium sized islands (1.5–20 km^2^); SN 4 = small islands (0.1–1.5 km^2^); SN 5 = skerries/islets (> 0.1 km^2^).

*Conclusion*. Based on ordination analyses and interpretation, we identified the following 9 CLG candidates in the three functional variables categories G, U and A (thematically slightly marginal variables are highlighted in grey; the absolute value of the correlation coefficient with the relevant GNMDS axis in parentheses):

(G1) Relief (**RE**; GNMDS axis 3/2/1): Steep_a (0.5080), RR1 (0.4495), Rug3_m (0.4482), Kskred_a (0.4301), Crug9_m (0.4056), Crug3_m (0.4030)
(G2) Abundance of wetlands (**VP**; GNMDS axis 3): Mire_a (0.4056), MI (0.3736), Inns_s (0.3180)
(G3) Coastal exposure (**KX**; GNMDS axis 2): R_net_a (0.5457), Ekspbe_a (0.4804); Stroml_a (0.3031)
(G4) Inner-outer coast (**IYK**; GNMDS axis 3): Discoast (0.3474), Oyst_i (0.3072)
(U1) Open area cover (**AP**; GNMDS axis 2): Bohei_a (0.4684), Sn_frisk (0.4015), Araaf_a (0.3915), Kbf_a (0.3874), Sn_imp (0.3329), AlpA (0.3293), Sn_torr (0.3152),
(U2) Vegetation (forest) cover (**SkP**; GNMDS axis 1/3): Arbar_a (0.4186), Arbla_a (0.4176), Arlov_a (0.3206)
(A1) Infrastructure (**OI**; GNMDS axis 1): Gab_a (0.6417); IfIu (0.6132), Gab_nae (0.5774); Vei_b_a (0.5237), Sefr_a (0.5149), Lled_a (0.4719); Abygg_a (0.4378); Build_a (0.4336)
(A2) Degree of agricultural intensity (**JP**; GNMDS axis 1): Arfull_a (0.6094), JI (0.5870), Arover_a (0.3180)

*Comments:* G3 and G4 (KX and IYK) are interpreted as two separate CLG candidates because the two groups of variables have different correlation patterns (Land_a was not included in this group because we regard it as hardly expressing a characteristic of the landscape itself). The KX group has |τ| > 0.3 with GNMDS axis 2, the IYK group has |τ| > 0.3 with GNMDS axis 3. No pair of variables, one from each group, had |τ| > 0.15.

#### Inland fine-sediment plains (IS)

A total of 118 OUs were affiliated with the inland fine-sediment plains (IF) major-type candidate. The two axes in the 2-dimensional GNMDS ordination of the IF118 data subset were confirmed by the corresponding DCA axes (Table 21). A possible but weak tongue-effect towards the right could be observed in the DCA ordination diagram for axes 1 and 2 (Fig. 68).

**Fig. 68.**
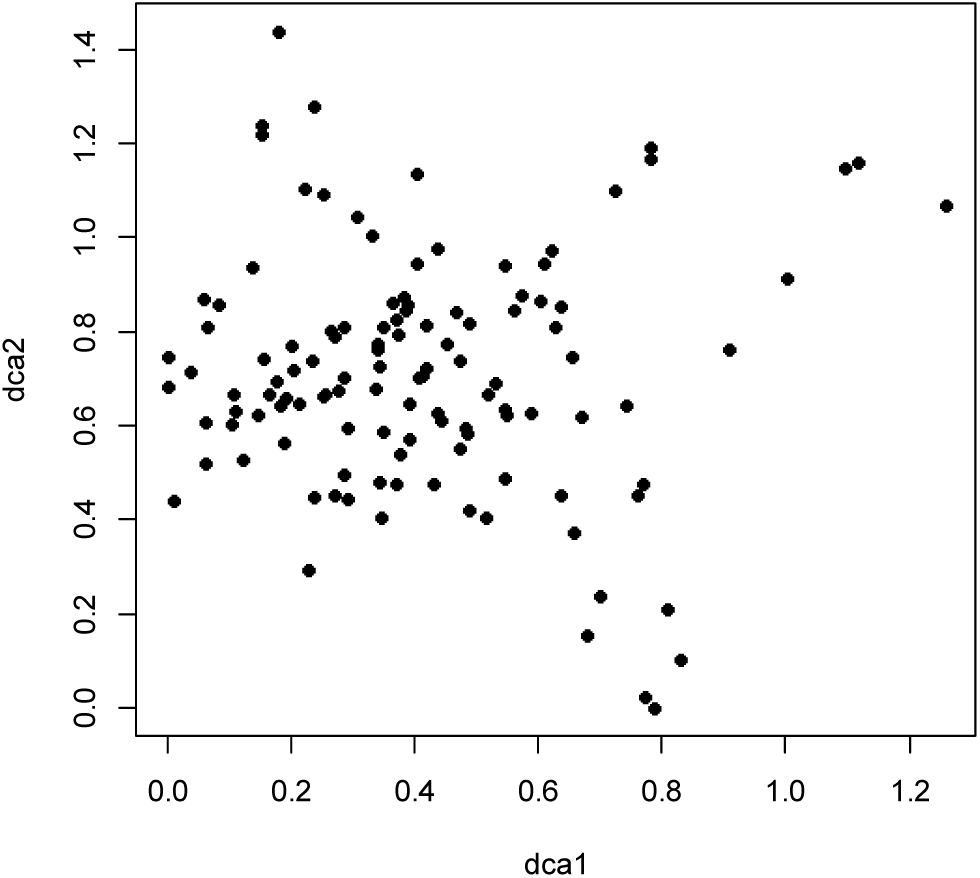
DCA ordination of the IF118 data subset (118 observation units in inland fine-sediment plains), axes 1 and 2, scaled in standard deviation (S.D.) units.

**Table 21.** Kendall’s rank correlation coefficients (τ), eigenvalues and gradient lengths for parallel DCA and GNMDS ordinations of the 118 observation units in the sub-dataset IF118, Inland fine-sediment plains. S.D. units = standard deviation units; H.C. units = half-change units.

|  | GNMDS axis 1 | GNMDS axis 2 | – | – |
| --- | --- | --- | --- | --- |
| Strongest correlated DCA-axis | DCA 1 | DCA 2 | DCA 3 | DCA 4 |
| $\tau$ (with strongest correlated DCA-axis) | 0.4312 | 0.4818 | – | – |
| DCA-axis eigenvalue | 0.0671 | 0.0786 | 0.0583 | 0.0370 |
| DCA-axis gradient length (S.D. units) | 1.2574 | 1.4403 | 1.3754 | 1.0017 |
| GNMDS-axis gradient length (H.C. units) | 1.0384 | 0.7696 | – | – |

The gradient length of GNMDS axis 2 was 0.741× the length of the longest axis; Table 21). The GNMDS ordination had Procrustes SS = 0.0002 and no unstable OUs. The OUs were relatively evenly spread out in the space spanned by GNMDS ordination axes 1 and 2, with OUs more sparsely spread out towards the periphery (Fig. 69).

**Fig. 69–72.**
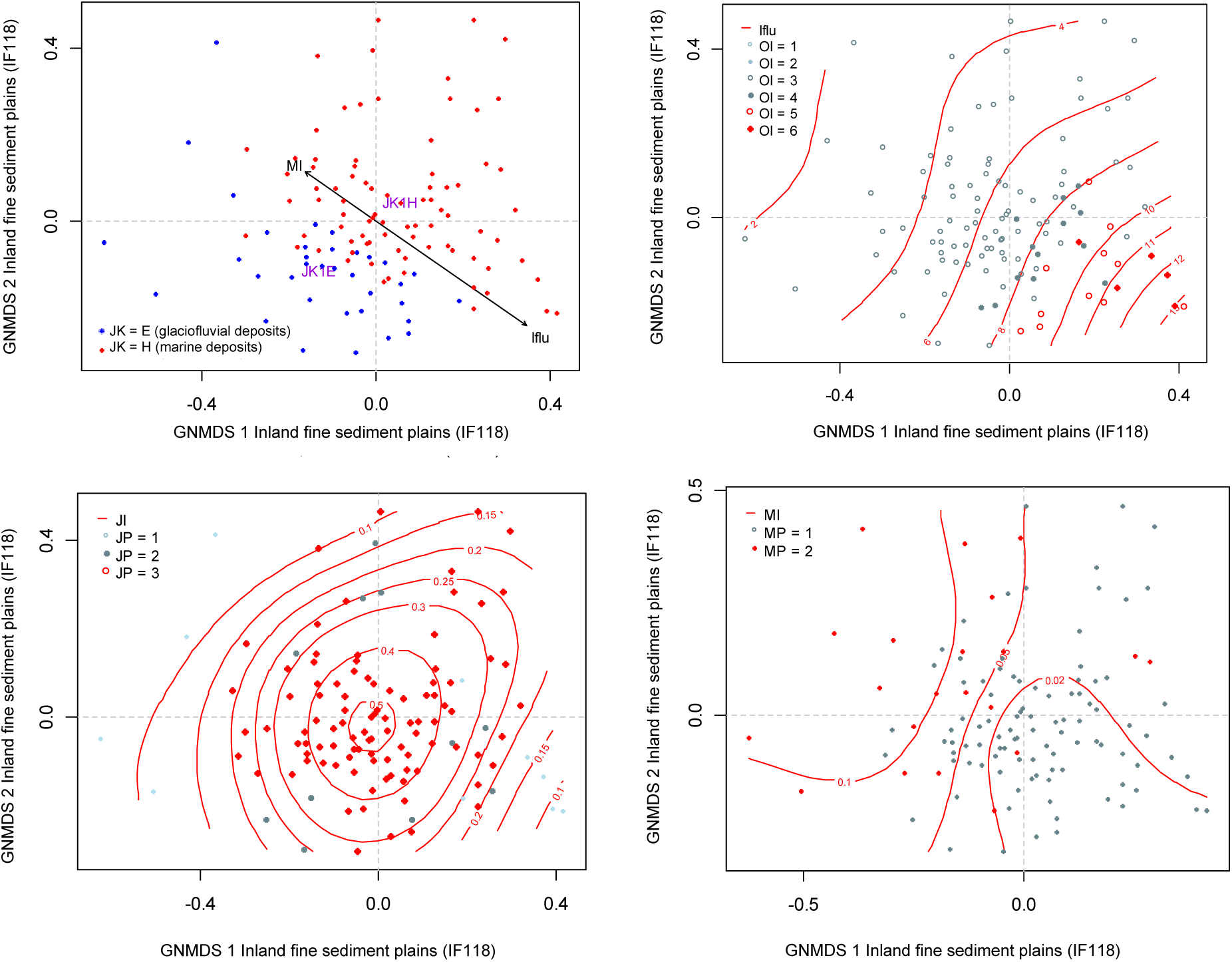
GNMDS ordinations of the 118 observation units in inland fine-sediment plains, axes 1 and 2, scaled in half-change (H.C.) units. (69) Abundance of mires (MI) and the infrastructure index (Iflu) are represented by arrows pointing in the direction of maximum increase of the variable (relative lengths of arrows are proportional to the correlation between the recorded variable and the ordination axis). Binary variables are represented by the abbreviated name in purple text, placed at the centroid of observation units belonging to the class in question. JK = primary soil/sediment category; JK1E = glaciofluvial deposits; JKH = marine deposits. (70) Isolines (red) for the infrastructure index (Iflu). OI = stepwise variation in the amount of infrastructure from low (OI 1–2) via medium (OI 3–4) to high amount of infrastructure, including towns and/or cities (OI 5–6). (71) Isolines (red) for agricultural land-use intensity (JI). JP = stepwise variation in agricultural land-use intensity from low (JP 1) via medium (JP2) to high (JP 3). (72) Isolines for the mire index (MI). MP = variation in mire abundance: MP1 = low abundance of mire; MP2 high abundance of mire.

The GNMDS-ordination (Fig. 69) show a clear division between with glaciofluvial (lower left) and marine (upper right) deposits. Correlations between key and analytic variables on one hand, and GNMDS axis 1 and 2 on the other hand, were in general weaker than for the other major-type candidates, but Table 22 show that the first axis was strongly related to infrastructure (Table 22; Fig. 70). Variation in infrastructure was more noticeable within OUs on marine sediments. The isoline diagram and distribution of OUs in Fig. 71 show that high agricultural land-use intensity (JI = 2) was a common property of all OUs affiliated with this major-type candidate. Table 22 and Figs 69 and 72 show that mire abundance was negatively related to the amount of infrastructure.

**Table 22.** Kendall’s rank correlation coefficients (τ) between GNMDS ordination axes 1–2 for the inland fine-sediment plains data subset (IF118) and the 10 primary key variables (rows 1–10) and the 66 variables used to characterise the 118 observation units.

| Variable |  | GNMDS1 | GNMDS 2 |
| --- | --- | --- | --- |
| RR1 | Relief | 0.1772 | 0.3190 |
| FIA | Amount of flat land | -0.2096 | -0.3164 |
| TPI1 | Peaks | 0.1446 | 0.0025 |
| AlpA | Boreal/alpine landscape | -0.1235 | 0.1235 |
| II | Abundance of freshwater lakes | 0.0555 | 0.1303 |
| MI | Abundance of mire | -0.3770 | 0.1613 |
| IfI | Amount of infrastructure (OI) | 0.5342 | -0.3516 |
| IfIu | Amount of infrastructure | 0.5654 | -0.3427 |
| JI | Agricultural land-use intensity | -0.0204 | -0.1332 |
| JKn | Soil type | -0.3508 | -0.4192 |
| Flat_a | AA flat terrain | -0.2084 | -0.3159 |
| Ktkmo_a | AA thick layer of till | -0.1308 | 0.0560 |
| Tpi1l_a | AA depressions | 0.1315 | 0.3130 |
| Tpi1h_a | AA convex terrain | 0.2419 | 0.3396 |
| Tpi1_mp | Terrain form TPI1, numerical, mean | 0.1444 | -0.0062 |
| River_a | AA large river | -0.1874 | -0.0273 |
| Steep_a | AA steep terrain/slope | 0.1453 | 0.2746 |
| Kskred_a | AA landslide soil | 0.0697 | -0.0490 |
| Rug3_m | Terrain ruggedness VRM3, mean | 0.2080 | 0.3634 |
| Rr1_m | Altitudinal range | 0.1772 | 0.3190 |
| Brich_a | AA lime-rich bedrock | 0.3053 | -0.1136 |
| Bpoor_a | AA lime-poor bedrock geology | -0.0811 | 0.0727 |
| Bplu_a | AA plutonic rock | 0.3241 | 0.2537 |
| Bvul_a | AA volcanic rock | 0.0007 | -0.0086 |
| Bavstn_a | AA sedimentary rock | 0.2738 | -0.1600 |
| Bomd_a | AA metamorphic rock | -0.2771 | 0.0525 |
| Mire_a | AA mire | -0.3514 | 0.2244 |
| Meant_a | AA moderate slope | 0.2075 | 0.2701 |
| Kmar_a | AA marine deposits | 0.2469 | 0.1005 |
| Klac_a | AA lacustrine deposits | -0.0431 | 0.0104 |
| Kelv_a | AA glaciofluvial deposits | -0.3790 | -0.4535 |
| Lake_a | AA freshwater lake | -0.0414 | 0.1906 |
| Innoy_s | Ant freshwater lake islands | -0.0165 | 0.1787 |
| Er_m | Hypsographic index, ER, mean | 0.0465 | -0.2279 |
| R_net_a | AA river | -0.1447 | 0.1392 |
| Inns_s | number of lakes | -0.0841 | 0.1726 |
| Asp_s_a | AA south facing terrain | -0.0058 | -0.2306 |
| Asp_n_a | AA north facing terrain | -0.0159 | 0.2611 |
| Dismire | distance to mire, mean | 0.3256 | -0.2134 |
| Dislake | distance to lake, mean | 0.0228 | -0.1309 |
| Discoast | distance to coast, mean | -0.4519 | -0.2053 |
| Guro_t_a | AA rugged terrain | -0.0917 | 0.2648 |
| Tpi6l_a | AA rugged terrain, TPI6 | -0.1641 | 0.0745 |
| Oyst_i | Inverse island size | -0.0309 | 0.2096 |
| Kbf_a | AA exposed bedrock | 0.2615 | 0.3929 |
| Bohei_a | AA boreal heaths | 0.1435 | 0.0393 |
| Arbar_a | AA coniferous forest | -0.3596 | -0.0054 |
| Arbla_a | AA mixed boreal forest | 0.0042 | -0.1743 |
| Arlov_a | AA deciduous forest | 0.1174 | -0.1314 |
| Sn_imp | AA impediment | -0.1235 | 0.1235 |
| Sn_flekk | AA patchy open treeless area | -0.0827 | 0.2011 |
| Sn_lav | AA lichen heath | -0.0024 | 0.0938 |
| Sn_torr | AA dry heath/open areas | -0.0535 | 0.2435 |
| Sn_frisk | AA moist/fresh heath/open areas | 0.0213 | 0.1742 |
| Sn_ureg | AA unregistered heath/open areas | 0.4117 | -0.3501 |
| Gab_nae | Ant commercial buildings | 0.4808 | -0.3389 |
| Gab_fi | Ant fisheries-related buildings | 0.1112 | 0.0461 |
| Gab_a | AA built-up | 0.5340 | -0.3465 |
| Vmag_a | AA regulated magazine | -0.0783 | 0.1518 |
| Vei_b_a | AA road | 0.3114 | -0.3735 |
| Sti_a | AA trail, path | 0.1885 | -0.0940 |
| Setr_s | Ant summer mountain pasture | 0.0119 | -0.0320 |
| Sefr_a | AA old buildings | 0.3950 | -0.3796 |
| Rd_anl_a | AA reindeer husbandry facilities | -0.1774 | 0.0181 |
| Build_a | AA built-up area | 0.5420 | -0.3395 |
| City_a | AA town/city area | 0.2385 | -0.2117 |
| Lled_a | AA power lines | 0.1238 | -0.0284 |
| Cul_u_s | Number of cultural heritage sites outdoors | 0.3522 | -0.2133 |
| Cul_t_s | number of technical heritage sites | 0.2535 | -0.1461 |
| Cul_k_s | number of church ruins | 0.4154 | -0.2792 |
| Cul_b_s | number of ancient rock art sites | 0.3609 | -0.0212 |
| Cul_a_s | number of archaeological heritage sites | 0.3734 | -0.0919 |
| Abygg_a | AA large buildings | 0.5372 | -0.3444 |
| Araaf_a | AA open areas | 0.2869 | -0.3124 |
| Arfull_a | AA arable land | -0.0719 | -0.1973 |
| Arover_a | AA surface cultivated land | 0.1563 | 0.0452 |

The continuous variables exposed bedrock (Kbf_a) and glaciofluvial deposits (Kelv_a) were more strongly related to the GNMDS axes than the factor variables for soil types (JK1 and KJ2). Thus, we conclude that these mutually correlated variables (Kbf and Kelv; τ = – 0.4082) are better suited for subsequent analyses with the aim of identifying CLG candidates. However, this does not prevent use of dominance of marine vs glacial sediments in the subsequent operationalisation process.

*Conclusion*. Based on ordination analyses and interpretation, we identified the following 9 CLG candidates in the three functional variables categories G, U and A (thematically slightly marginal variables are highlighted in grey; the absolute value of the correlation coefficient with the relevant GNMDS axis in parentheses; note that most CLG candidates have a weak definition basis):

(G1) Soil type (‘abundance of glaciofluvial sediments’; **JA**; GNMDS axis 1/2): Kelv_a (0.4535), Bbf_a (0.3929)
(G2) Distance to coast (**KA**; GNMDS axis 1): Discoast (0.4519)
(G3) Abundance of mire (**MP**; GNMDS axis 1): MI (0.3770), Mire_a (0.3513), Dismire (0.3256)
(G4) Relief (**RE**; GNMDS axis 2): Rug3_m (0.3634), RR1 (0.3190), FlA (0.3164), Tpi1h_a (0.3396), Tpi1l_a (0.3159)
(G5) Bedrock (**BA**; GNMDS axis 1): Bplu_a (0.3241)
(G6) Geological richness (**GR**; GNMDS axis 1): Brich (0.3053)
(U1) Open area cover (**AP**; GNMDS axis 1/2): Sn_ureg (0.4117), Araaf_a (0.3124) (U2) Coniferous forest cover (**SkP**; GNMDS axis 1): Arbar_a (0.3596)
(A1) Infrastructure (**OI**; GNMDS axis 1): IfIu (0.5654), Build_a (0.5420), Abygg_a (0.5372), Gab_a (0.5340), Gab_nae (0.4808), Cul_k_s (0.4154)

*Comment:* Bplu_a and Brich_a are treated as separate CLG candidates because the variables are not correlated with each other (τ = 0.0730).

#### Inland plains without dominance of fine sediments (IX)

A total of 399 OUs were affiliated with the ‘other inland plains’ (IX) major-type candidate. The two axes in the two-dimensional GNMDS ordination of the IX399 data subset were confirmed by the corresponding DCA axes (Table 23; Figs 73–74). The GNMDS ordination had Procrustes SS = 0.1490 and two unstable OUs. No visible artefacts could be observed in the in the ordination space spanned by GNMDS ordination axes 1 and 2 (Fig. 74). A tendency for reduced OU density towards higher scores along both axes was observed. Relative gradient lengths of GNMDS and DCA axes, respectively, were in the same range (0.79–0.85; cf. Table 23).

**Fig. 73.**
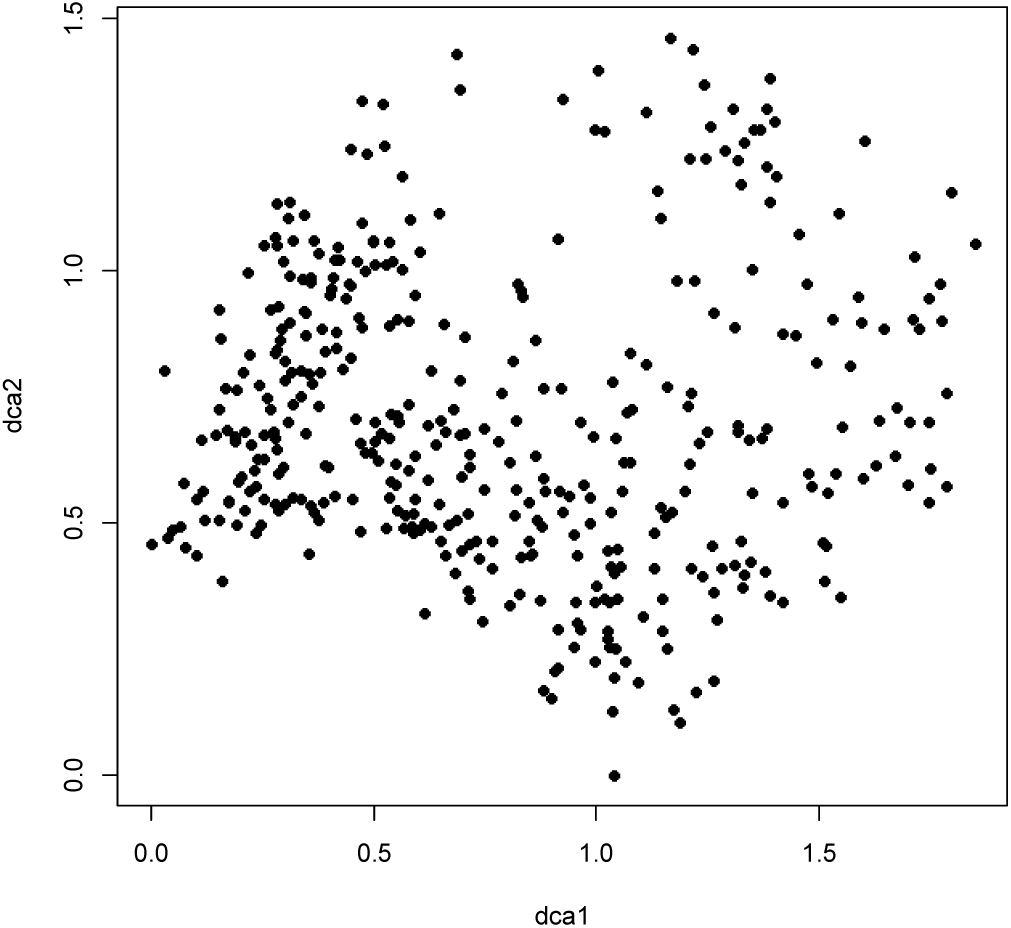
DCA ordination of the IX399 data subset (399 observation units of other inland plains), axes 1 and 2, scaled in standard deviation (S.D.) units.

**Figs. 74–79.**
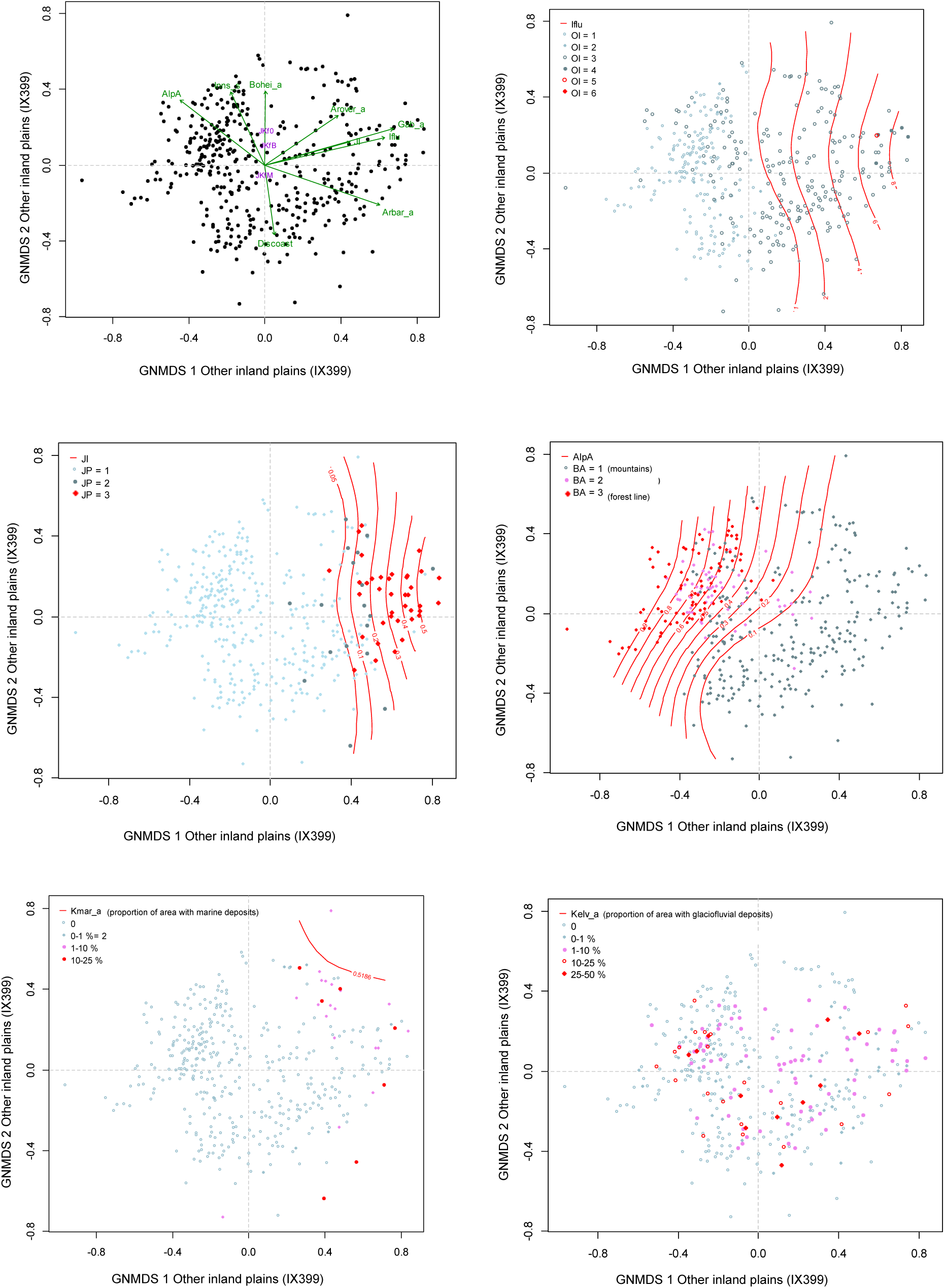
Distribution of observation units with specific properties in the GNMDS ordination diagram, axes 1 and 2, for the ‘other inland plains’ data set (IX399). (74) Selected variables are represented by green arrows pointing in the direction of maximum increase of the variable (relative lengths of arrows are proportional to the correlation between the recorded variable and the ordination axis). Binary variables are represented by the name, abbreviated according to Tables 4 and 5, and placed at the centroid for the presence class (purple text). (75) Isolines (red) for the infrastructure index (Iflu). OI = stepwise variation in amount of infrastructure from low (OI 1–2) via medium (OI 3–4) to high amount of infrastructure, including towns and/or cities (OI 5–6). (76) Isolines (red) for agricultural land- use intensity (JI). JP = stepwise variation in agricultural land-use intensity from low (JP 1) via medium (JP2) to high (JP 3). (77) Isolines (red) for the proportion of OU area situated above the forest line (AlpA). BA = stepwise variation along the gradient from boreal to alpine landscapes from areas dominated by forest (BA 1) via areas located at the forest line (BA 2) to mountainous areas above the forest line (BA 3). (78) Proportion of area covered by with marine deposits (Kmar_a), shown by coloured symbols and isolines (red). (79) Proportion of area with glaciofluvial deposits (Kelv_a), shown by coloured symbols.

**Table 23.**
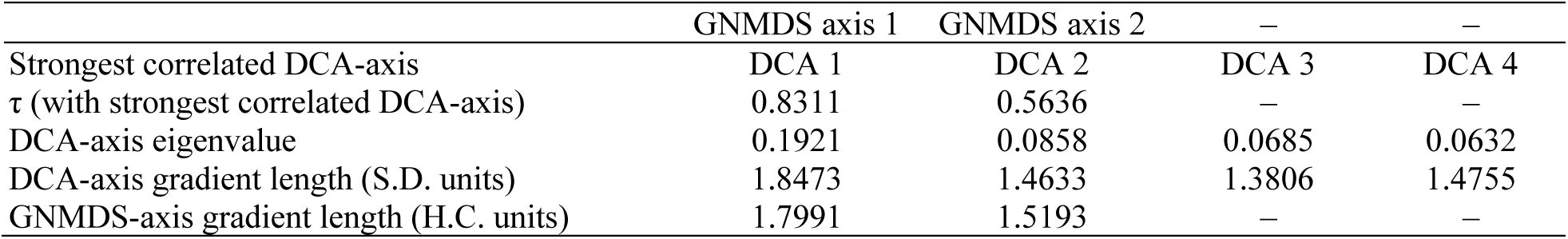
Kendall’s rank correlation coefficients (τ), eigenvalues and gradient lengths for parallel DCA- and GNMDS-ordinations of the subset IX399, Other inland plains. S.D. units = standard deviation units; H.C. units = half-change units.

|  | GNMDS axis 1 | GNMDS axis 2 | – | – |
| --- | --- | --- | --- | --- |
| Strongest correlated DCA-axis | DCA 1 | DCA 2 | DCA 3 | DCA 4 |
| $\tau$ (with strongest correlated DCA-axis) | 0.8311 | 0.5636 | – | – |
| DCA-axis eigenvalue | 0.1921 | 0.0858 | 0.0685 | 0.0632 |
| DCA-axis gradient length (S.D. units) | 1.8473 | 1.4633 | 1.3806 | 1.4755 |
| GNMDS-axis gradient length (H.C. units) | 1.7991 | 1.5193 | – | – |

Correlations between key and analytic variables on one hand, and GNMDS axis 1 scores on the other (Table 24), supported by vector diagrams (Fig. 74) and isoline diagrams (Figs 75–78) revealed a first axis that was very strongly related to a gradient in vegetation cover from boreal to alpine OUs (Fig. 77), with strong relationships also to increasing land-use intensity (infrastructure and agricultural land use; Figs 75 and 76, respectively). However, the variation in the amount of infrastructure was generally low within this major-type candidate; only 6 OUs had above intermediate infrastructure (Fig. 75; OI > 4). The same pattern applied to agricultural land-use intensity; only 39 OUs had high agricultural land-use intensity. These OUs made up a cluster at the right-hand (i.e. high-score) end of GNMDS 1 (Fig. 76; JI = 3). Very few OUs belonging this major-type candidate had large proportional cover of marine deposits (Fig. 78), while OUs with glaciofluvial deposits (Fig. 79) were evenly distributed throughout the ordination space. The few OUs with more than 10% marine deposits were located in the periphery of the ordination space (high GNMDS 1 scores; Fig. 78), indicating a transition towards inland fine-sediment plains.

**Table 24.** Kendall’s rank correlation coefficients (τ) between GNMDS ordination axes 1–2 for the other inland plains data subset (IX399) and the 9 primary key variables (rows 1–9) and the 65 variables used to characterise the 399 observation units.

| Variable |  | GNMDS1 | GNMDS 2 |
| --- | --- | --- | --- |
| RR1 | Relief | –0.0025 | –0.0866 |
| FIA | Amount of flat land | 0.0467 | 0.0595 |
| TPI1 | Peaks | –0.1298 | –0.0675 |
| AlpA | Boreal/alpine landscape | –0.5235 | 0.3069 |
| II | Abundance of freshwater lakes | 0.0232 | 0.1911 |
| MI | Abundance of mire | 0.2023 | –0.0826 |
| Ifl |  | 0.6766 | 0.0821 |
| Iflu | Amount of infrastructure | 0.6322 | 0.1207 |
| JI | Agricultural land-use intensity | 0.5853 | 0.1259 |
| Flat_a | AA flat terrain | 0.0467 | 0.0596 |
| Ktkmo_a | AA thick layer of till | –0.0894 | –0.2357 |
| Tpi1l_a | AA depressions | 0.1054 | –0.0080 |
| Tpi1h_a | AA convex terrain | –0.0491 | –0.0223 |
| Tpi1_mp | Terrain form TPI1, numerical, mean | –0.1188 | –0.0523 |
| River_a | AA large river | 0.0548 | 0.1186 |
| Steep_a | AA steep terrain/slope | 0.0885 | –0.0289 |
| Kskred_a | AA landslide soil | 0.0565 | 0.1445 |
| Rug3_m | Terrain ruggedness VRM3, mean | 0.0592 | 0.0376 |
| Rr1_m | Altitudinal range | –0.0023 | –0.0867 |
| Brich_a | AA lime-rich bedrock | 0.1933 | 0.1648 |
| Bpoor_a | AA lime-poor bedrock geology | 0.2089 | 0.0676 |
| Bplu_a | AA plutonic rock | 0.0413 | 0.3845 |
| Bvul_a | AA volcanic rock | 0.0860 | –0.1186 |
| Bavstn_a | AA sedimentary rock | –0.0964 | –0.0026 |
| Bomd_a | AA metamorphic rock | 0.0185 | –0.1843 |
| Mire_a | AA mire | 0.2049 | –0.0725 |
| Meant_a | AA moderate slope | –0.0115 | –0.0690 |
| Kmar_a | AA marine deposits | 0.2891 | 0.1477 |
| Klac_a | AA lacustrine deposits | 0.1778 | 0.0512 |
| Kelv_a | AA glaciofluvial deposits | 0.1405 | 0.0017 |
| Kbf_a | AA exposed bedrock | 0.0888 | 0.1613 |
| Lake_a | AA freshwater lake | –0.0426 | 0.2421 |
| Innoy_s | Ant freshwater lake islands | 0.0700 | 0.2640 |
| Er_m | Hypsographic index, ER, mean | –0.1138 | –0.1051 |
| R_net_a | AA river | 0.0856 | 0.2056 |
| Inns_s | number of lakes | –0.2281 | 0.3306 |
| Asp_s_a | AA south facing terrain | 0.0926 | 0.0300 |
| Asp_n_a | AA north facing terrain | –0.0301 | 0.0677 |
| Dismire | distance to mire, mean | –0.2108 | 0.0658 |
| Dislake | distance to lake, mean | 0.1307 | –0.2695 |
| Discoast | distance to coast, mean | 0.1634 | –0.3255 |
| Guro_t_a | AA rugged terrain | 0.0224 | 0.1084 |
| Tpi6l_a | AA rugged terrain, TPI6 | 0.1127 | –0.0221 |
| Oyst_i | Inverse island size | 0.1976 | 0.3032 |
| Bohei_a | AA boreal heaths | –0.0110 | 0.3610 |
| Araaf_a | AA open areas | 0.4935 | 0.0761 |
| Arbar_a | AA coniferous forest | 0.5631 | –0.1949 |
| Arbla_a | AA mixed boreal forest | 0.5056 | –0.2135 |
| Arlov_a | AA deciduous forest | 0.4468 | –0.1197 |
| Sn_imp | AA impediment | –0.2839 | 0.0278 |
| Sn_flekk | AA patchy open treeless area | –0.4642 | 0.3064 |
| Sn_lav | AA lichen heath | –0.1487 | 0.1891 |
| Sn_torr | AA dry heath/open areas | –0.4942 | 0.4344 |
| Sn_frisk | AA moist/fresh heath/open areas | –0.2566 | 0.4494 |
| Sn_ureg | AA unregistered heath/open areas | 0.1051 | 0.2928 |
| Gab_nae | Ant commercial buildings | 0.5198 | 0.1280 |
| Gab_fi | Ant fisheries-related buildings | 0.1306 | 0.0688 |
| Gab_a | AA built-up | 0.6714 | 0.1228 |
| Vmag_a | AA regulated magazine | 0.0874 | 0.0218 |
| Vei_b_a | AA road | 0.5517 | 0.1708 |
| Sti_a | AA trail, path | 0.0588 | 0.1076 |
| Setr_s | Ant summer mountain pasture | 0.3080 | -0.1338 |
| Sefr_a | AA old buildings | 0.5921 | 0.0536 |
| Rd_anl_a | AA reindeer husbandry facilities | -0.0926 | 0.0066 |
| Build_a | AA built-up area | 0.3381 | 0.0680 |
| Lled_a | AA power lines | 0.5006 | 0.1651 |
| Cul_u_s | Number of cultural heritage sites outdoors | 0.1169 | 0.0299 |
| Cul_t_s | number of technical heritage sites | 0.1534 | 0.0744 |
| Cul_k_s | number of church ruins | 0.2630 | 0.1392 |
| Cul_b_s | number of ancient rock art sites | 0.0953 | 0.0170 |
| Cul_a_s | number of archaeological heritage sites | 0.4490 | 0.2126 |
| Abygg_a | AA large buildings | 0.3441 | 0.0707 |
| Arfull_a | AA arable land | 0.5948 | 0.1608 |
| Arover_a | AA surface cultivated land | 0.3975 | 0.1925 |

We interpret GNMDS axis 2 as related to variation from coastal to inland geo-ecological properties, as expressed by the variables distance to coast, inverse island size, bedrock and number of lakes (Table 24). This gradient was also related to the gradient from boreal to alpine landscapes revealed by GNMDS axis 1.

*Conclusion*. Results of ordination and correlation analyses support the separation of fine-sediment plains (IF) from other inland plains (IX). Based on ordination analyses and interpretation, we identified the following 6 CLG candidates in the three functional variables categories G, U and A (note that most CLG candidates have a weak definition basis) for the IX major-type candidate:

(G1) Bedrock (BG; Axis 2): Bplu_a (0.3845)
(G2) Freshwater lake properties (IP; Axis 2): Inns_s (0.3306)
(G3) Coastal position (KA; Axis 2): Discoast (0.3255), Oyst_i (0.3032)
(U1) Vegetation (forest) cover (incl. the gradient from mountain to lowland; SkP; Axis 1/2): Arbar_a (0.5631), AlpA (0.5235), Arbla_a (0.5056), Sn_frisk (0.4994), Sn_torr (0.4941), Araaf_a (0.4935), Sn_flekk (0.4642), Arlov_a (0.4468).
(A1) Infrastructure (OI; Axis 1): Gab_a (0.6714), IfIu (0.6366), Sefr_a (0.5921), Vei_b_a (0.5517), Gab_nae (0.5598), Cul_a_s (0.4490)
(A2) Degree of agricultural intensity (JP; Axis 1): Arfull_a (0.5948), JI (0.5853), Arover_a (0.3975)

*Comments:* The U variables form a ‘chain’ of correlated variables that are strongly (|τ| > 0.4) correlated with the abundance of open heath and alpine areas (Sn_frisk and/or AlpA), which are in turn strongly correlated (|τ| > 0.4874) with each other. Coniferous forest and boreal heaths (Arbar_a and Bohei_a), which are weakly negatively correlated with each other, make up the extremes of this group. This group is therefore regarded as one CLG candidate. Boreal heath (Bohei_a), which has the strongest correlation with GNMDS axis 2 among potential ‘members’ in a group for open areas, |τ| = 0.3610, was left out because more than 8 variables were at least quite strongly correlated (|τ| > 0.4) with this axis.

#### Inland valleys (ID)

A total of 1012 OUs were affiliated with the inland valleys (ID) major-type candidate. The three axes of the 3-dimensional GNMDS ordination of the ID1012 data subset were confirmed by the corresponding DCA axes (Table 25). Gradient lengths of the second and third GNMDS axes were 0.760× and 0.664× the length of GNMDS axis 1, respectively. The GNMDS ordination had Procrustes SS = 0.1796 and one unstable OU. No visible artefacts could be seen in the ordination diagrams (Figs 80 and 81), but a tendency for reduced density towards the periphery of the ordination space was observed. Gradient lengths of the first axes of both ordinations indicate existence of a strong major gradient in the ID1012 data subset (Table 25). Correlations between key and analytic variables on one hand, and GNMDS axis 1 scores on the other (Table 26), supported by vector diagrams (Fig. 81) and isoline diagrams (Figs 82–86), established a very strong relationship between the first axes and a gradient in vegetation cover from alpine to boreal OUs (Fig. 84), along which the intensity of land use (infrastructure and agricultural land use; Figs 82 and 83, respectively) increased. The isoline diagrams show that OUs with medium and high agricultural land-use intensity form a well-delimited group in the lower right corner of the ordination diagram (Fig. 83; JP > 1). The amount of infrastructure (Fig. 82) followed the same pattern although in a more gradual manner, increasing from the upper left (mountains) to the lower right (lowlands) in the ordination diagram.

**Fig. 80.**
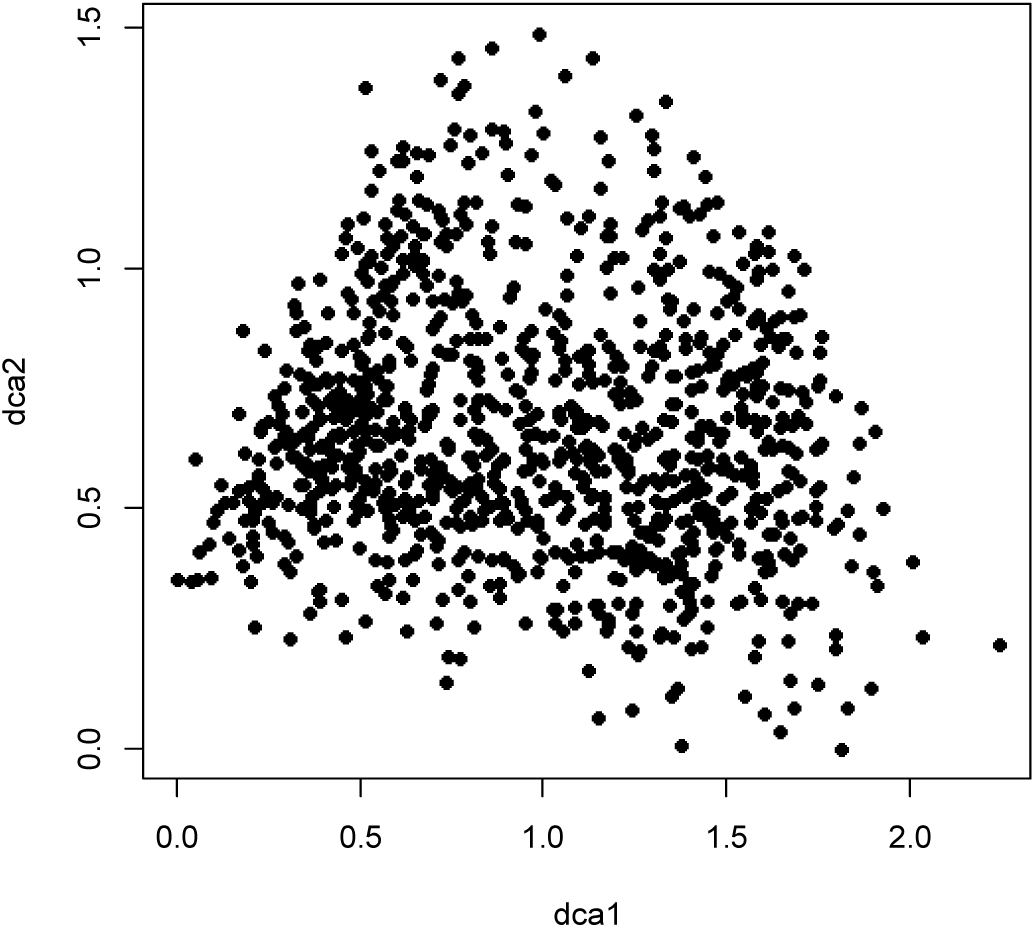
DCA ordination of the subset ID1012 (1012 observation units in inland valleys), axes 1 and 2, scaled in standard deviation (S.D.) units.

**Figs. 81–86.**
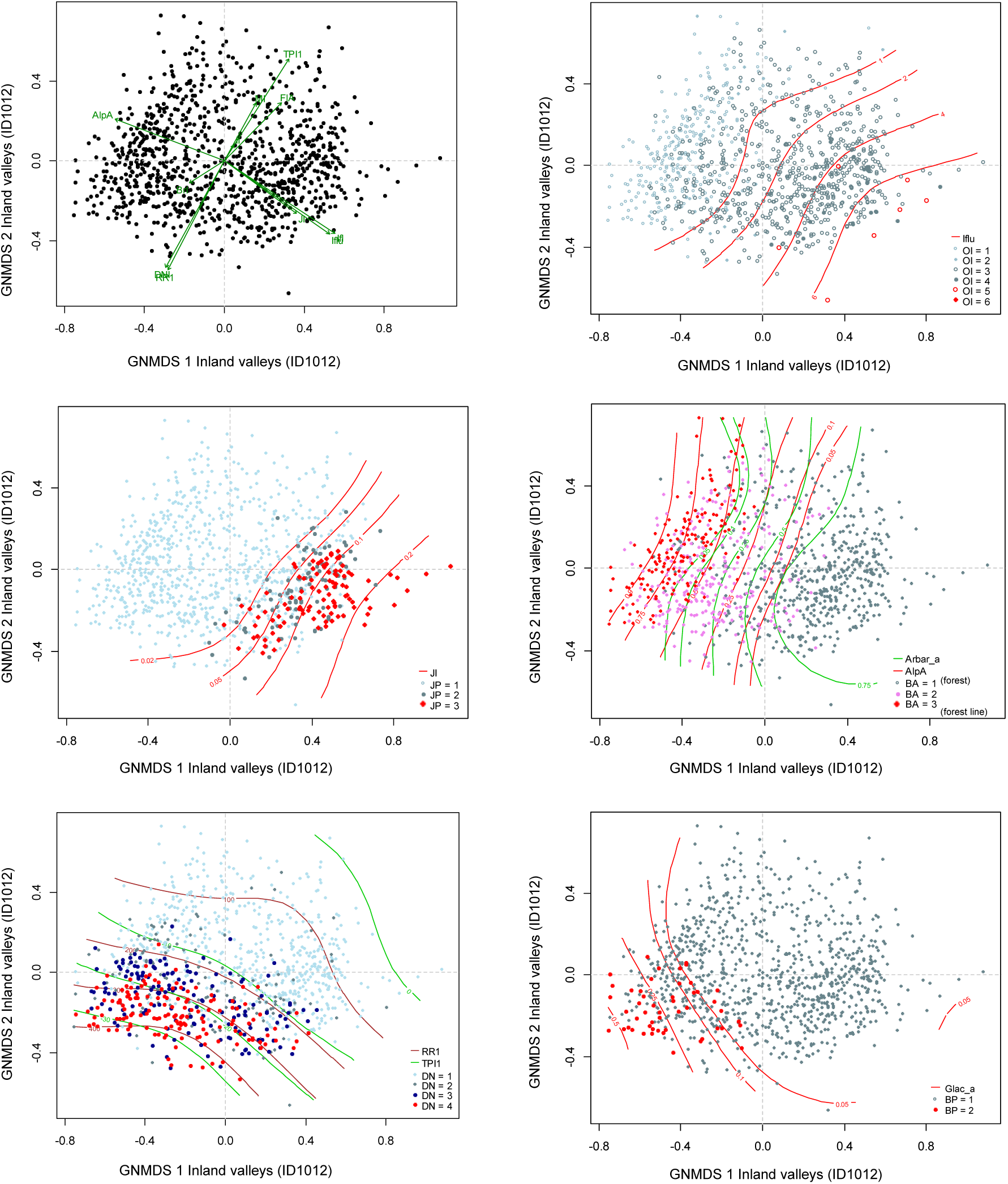
Distribution of observation units with specific properties in the GNMDS ordination diagram, axes 1 and 2, for the inland valleys data set (ID1012). (81) Selected variables are represented by green arrows pointing in the direction of maximum increase of the variable (relative lengths of arrows are proportional to the correlation between the recorded variable and the ordination axis). Variable names are abbreviated according to Tables 4 and 5. (82) Isolines (red) for the infrastructure index (Iflu). OI = stepwise variation in the amount of infrastructure from low (OI 1–2) via medium (OI 3–4) to high amount of infrastructure, including towns and/or cities (OI 5–6). (83) Isolines (red) for agricultural land- use intensity (JI). JP = stepwise variation in agricultural land-use intensity from low (JP 1) via medium (JP2) to high (JP 3). (84) Isolines (red) for the proportion of OU area situated above the forest line (AlpA). BA = stepwise variation along the gradient from boreal to alpine landscapes from areas dominated by forest (BA 1) via areas located at the forest line (BA 2) to mountainous areas above the forest line (BA 3). (85) Isolines for relative relief (RR1; brown lines) and terrain position index (TPI1; green lines). DN = valley shape: DN 1 = open valley; DN 2 = relatively open valley; DN 3 = relatively narrow and steep valley; DN 4 = narrow and steep-sided valley. (86) Isolines (red) for the proportion of the OU covered by glaciers (BrI). BP1 = absence of glacier, BP2 = presence of glacier.

**Table 25.** Kendall’s rank correlation coefficients (τ), eigenvalues and gradient lengths for parallel DCA- and GNMDS-ordinations of the subset ID1012, Inland valleys. S.D. units = standard deviation units; H.C. units = half-change units.

|  | GNMDS axis 1 | GNMDS axis 2 | GNMDS axis 3 | – |
| --- | --- | --- | --- | --- |
| Strongest correlated DCA-axis | DCA 1 | DCA 2 | DCA 3 | DCA 4 |
| $\tau$ (with strongest correlated DCA-axis) | 0.8943 | 0.6450 | 0.4169 | – |
| DCA-axis eigenvalue | 0.1793 | 0.0664 | 0.0474 | 0.0425 |
| DCA-axis gradient length (S.D. units) | 2.2410 | 1.4898 | 1.2891 | 1.2601 |
| GNMDS-axis gradient length (H.C. units) | 1.8338 | 1.3946 | 1.2171 | – |

**Table 26.**
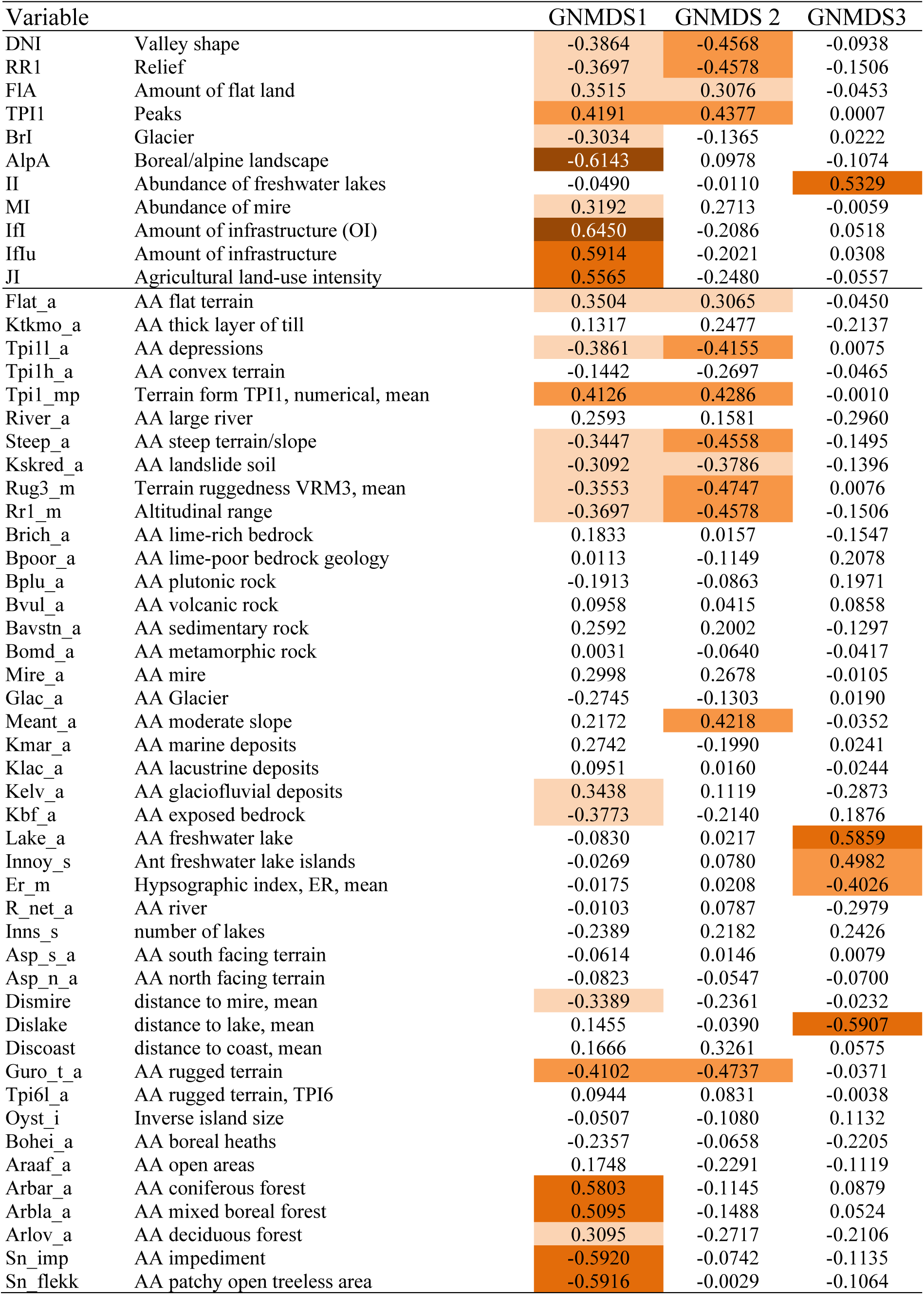
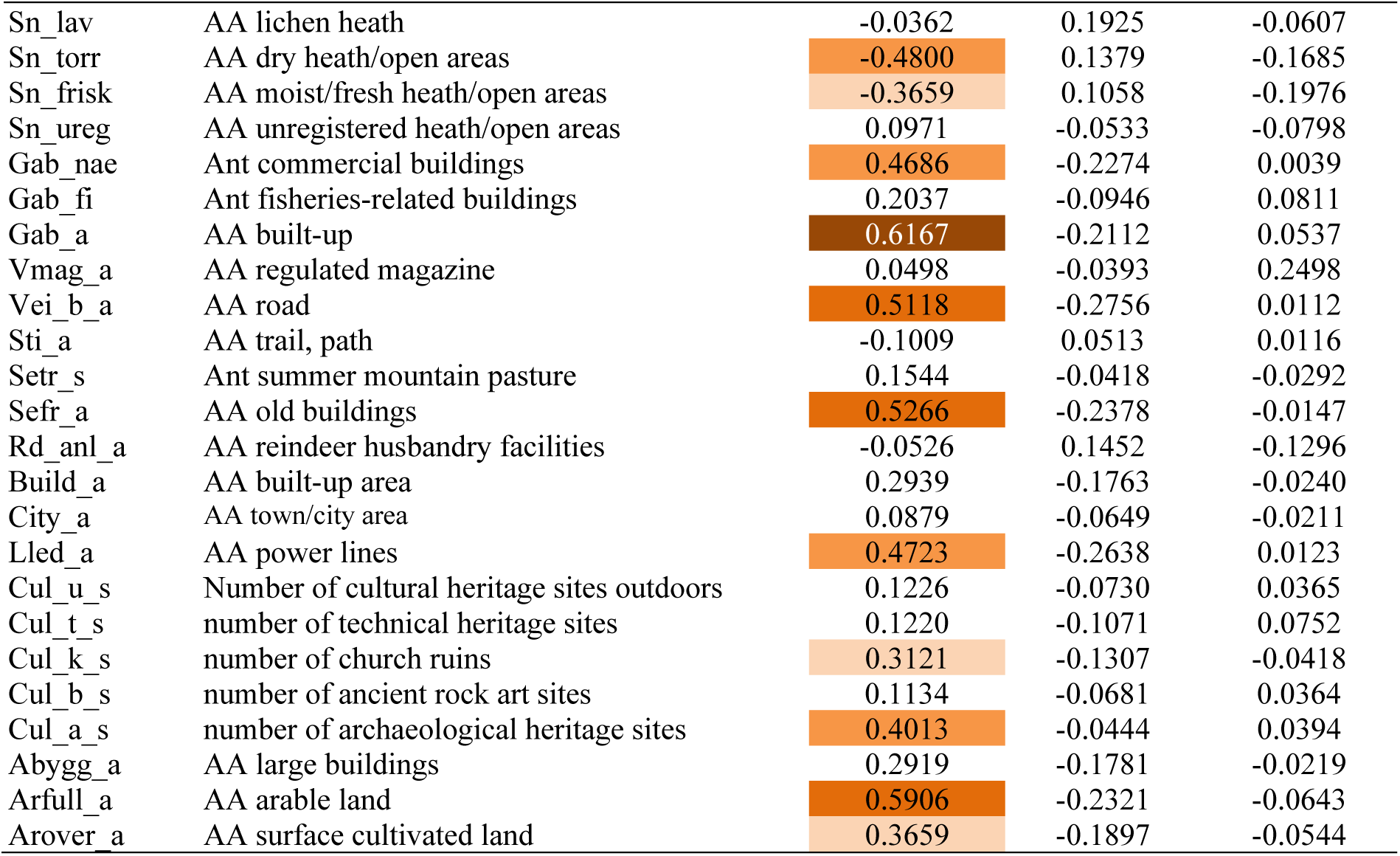
Kendall’s rank correlation coefficients (τ) between GNMDS ordination axes 1–3 for the inland valleys data subset (ID1012) and the 11 primary key variables (rows 1–11) and the 67 analytic variables used to characterise the 1012 observation units.

GNMDS axis 2 was clearly related to terrain form; three key variables and seven analytical variables that expressed terrain variables had correlation coefficients |τ| > 0.4 with this axis (Table 26; Fig. 81). The vector diagram (Fig. 81) shows that the material can be arranged in two very distinct variable groups; (i) the gradient from above to below the treeline (i.e. from mountains to mainly forested lowlands), along which the amount of infrastructure and agricultural land-use intensity increased; and (ii) relative relief and other terrain-form variables. OUs with presence of glacier (Fig. 86; BA = 2) occurred in the lower left corner in the space spanned by GNMDS ordination axes 1 and 2, associated with high relief. OUs with glacier did, however, not form a separate group but occurred intermixed with high-relief OUs without glaciers. Thus, glacier presence apparently varies within the ‘high-relief end’ of the relief gradient.

GNMDS axis 3 was strongly correlated with variables that expressed presence and areal cover of freshwater lakes (Table 26; Figs 87–88) although OUs characterised by lakes did not segregate clearly from other OUs along GNMDS axis 3 (Fig. 88).

**Figs. 87–88.**
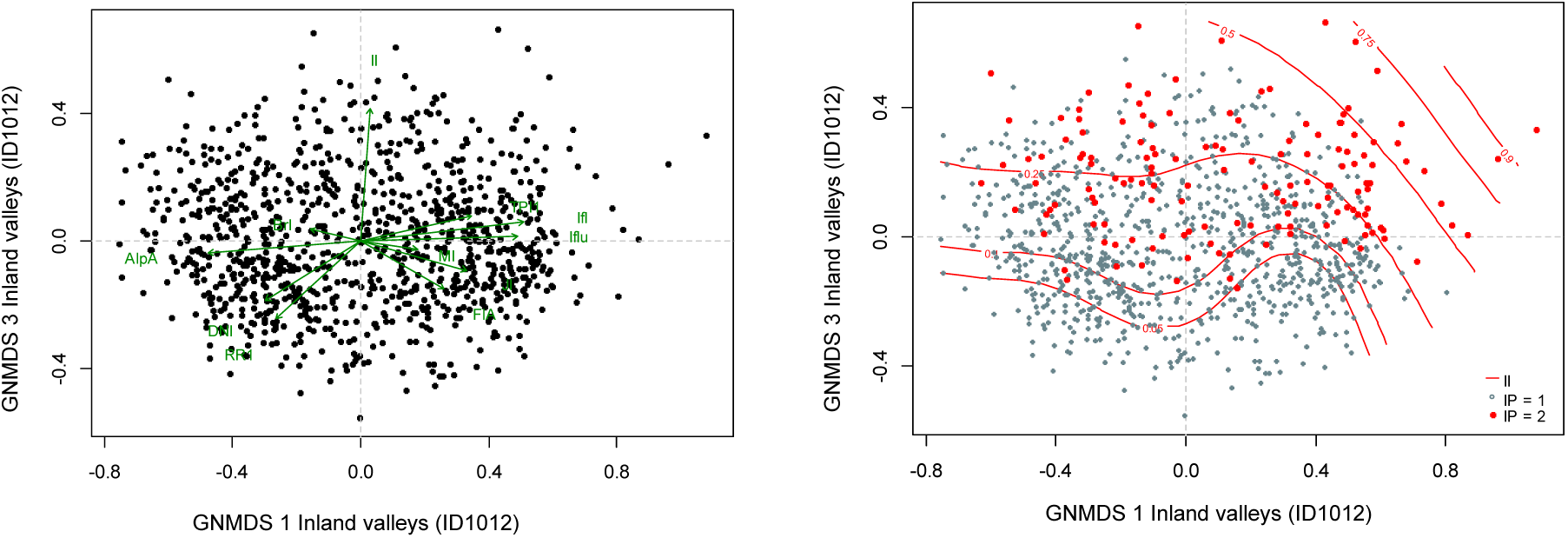
Distribution of observation units with specific properties in the GNMDS ordination diagram, axes 1 and 3, for the inland valleys data set (ID1012). (87) Selected variables represented by green arrows pointing in the direction of maximum increase of the variable (relative lengths of arrows are proportional to the correlation between the recorded variable and the ordination axis). Variable names are abbreviated according to Tables 4 and 5. (88) Isolines (red) for the lake index (II). IP = abundance of freshwater lakes (IP1 = low; IP2 = high).

Since the proportions of areas of both glacial deposits and bare rock were correlated with GNMDS axis 1, we plotted the distribution of the 1018 OUs on primary soil classes JK1. A clear, but not strong, relationship between soil category (JK1) and positions in the ordination diagram can be seen in Fig. 89. The distribution of soil types was related to relief; areas with thick layers of till are associated with low relative relief, while bare rock and landslide soils are correlated with high relative relief. The only soil type that was clearly associated with the CLG from mountain to lowland was, not unexpectedly, the abundance of marine sediments (11 OUs in the lower right corner of the ordination diagram).

**Fig. 89.**
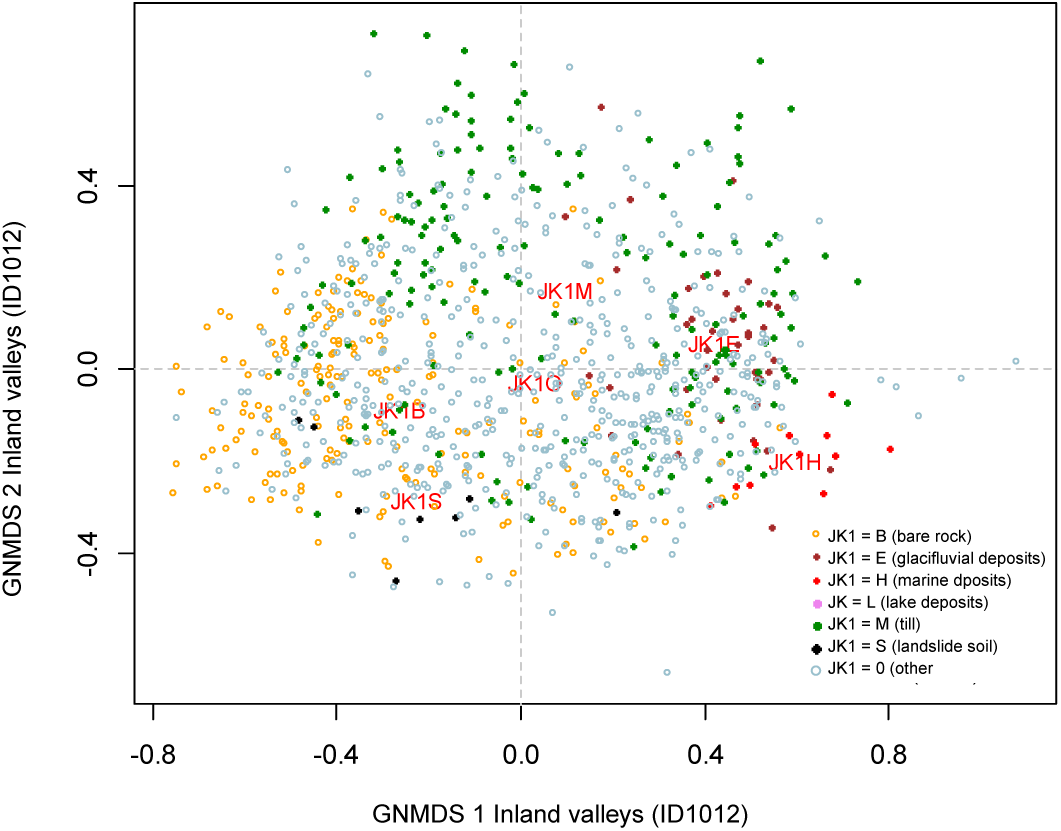
Distribution of observation units with specific soil/sediment properties in the GNMDS ordination diagram, axes 1 and 2, for the inland valleys data set (ID1012). The centroid of observation units belonging to each class is indicated by red text. Primary soil/sediment classes (JK1): B = exposed bedrock (> 50% of area covered by exposed bedrock); E = glacifluvial deposits; H = marine deposits; M = thick layer of till; 0 = no soil/sediment class (> 50% not assigned to specific sediment/soil type).

Based on ordination analysis and interpretation, we identified the following 8 CLG candidates in the three functional variable categories G, U and A (note that most CLG candidates have a weak definition basis):

(G1) Relief (valley form) (RE; GNMDS axis 2/1): Rug3_m (0.4747), Guro_t_a (0.4737) Steep_a (0.4588), Rr1_m (0.4578), DNI (0.4568), TPI1 (0.4377), Meant_a (0.4218), TPI1l_a (0.4155), TPI1_mp (0.4286)
(G2) Freshwater lake properties (IP; GNMDS axis 3/1): Dislake (0.5907), Lake_a (0.5859), II (0.5329), Innoy_s (0.4982), Er_m (0.4026)
(G3) Soil (JA; GNMDS axis 1): Kbf_a (0.3773), Kelv_a (0.3438); JK1 (2 classes + ‘background’)
(G4) Abundance of mire (MP; GNMDS axis 1): Dismire (0.3389), MI (0.3192) (G5) Glacier (BP; GNMDS axis 1): BrI (0.3034)
(U1) Vegetation (forest) cover (SkP; GNMDS axis 1): AlpA (0.6143), Sn_imp (0.5920), Sn_flekk (0.5916), Arbar_a (0.5803), Arbla_a (0.5095), Sn_torr (0.4800)
(A1) Infrastructure (OI; GNMDS axis 1): Gab_a (0.6167), IfIu (0.5914), Sefr_a (0.5166), Vei_b_a (0.5118), Lled_a (0.4723), Gab_nae (0.4686), Cul_a_s (0.4013)
(A2) Degree of agricultural land-use intensity (JP; GNMDS axis 1): Arfull_a (0.5906), JI (0.5563), Arover_a (0.3659)

*Comments:* The hypsographic index (Er_m) included in CLG-candidate G2 was correlated with GNMDS axis 3 as well as with lake variables and was assigned to this CLG candidate despite the pairwise correlations were weak [τ (Er_m, Lake_a) = –0.5129]. The amount of freshwater lakes and abundance of mires (G2 and G4, respectively) make up two separate CLG candidates as evident from the lack of a relationship: τ (Dismire, Lake_a) = 0.0678. Soil, expressed as a factor-type variable (G3), was included among hCLG candidates with classes for dominance of bare rock (B) and of glacial sediments (E). These two dominant soil types were well represented in the data material and the centroids of both were placed clearly off the origin in the ordination diagram (Fig. 89). Glacier (G5) is included as a separate CLG candidate because the analysis variable Glac_a was not strongly correlated with relief [τ (Glac_a, RR1) = 0.2289].

#### Inland hills and mountains (IA)

A total of 1618 OUs were affiliated with the inland valleys (IA) major-type candidate. The three axes in the 3-dimensional GNMDS ordination of the IA1618 data subset were confirmed by corresponding DCA axes (Table 27). The GNMDS ordination had Procrustes SS = 0.3208 and no unstable OU. No visible artefacts could be seen in the ordination diagrams (Figs 90 and 91), but a tendency for reduced density towards the periphery of the ordination space was observed. The cloud of points in the DCA ordination diagram (axes 1 and 2) had a distinctive triangular structure. Gradient lengths of the first axes of GNMDS and DCA ordinations indicated existence of a strong first axis (Table 27), while the relative length of both of GNMDS axes 2 and 3 were > 0.8× the length of GNMDS axis 1.

**Fig. 90.**
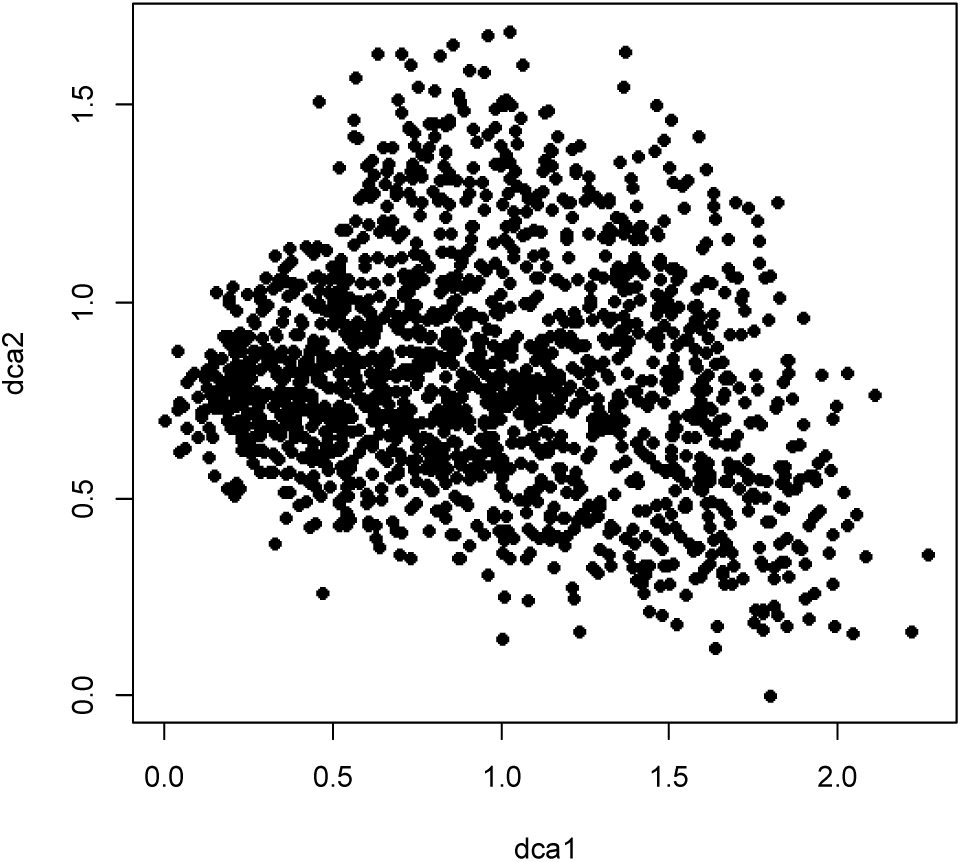
DCA ordination of the subset IA1618 (1618 observation units in inland hills and mountains), axes 1 and 2, scaled in standard deviation (S.D.) units.

**Figs. 91–96.**
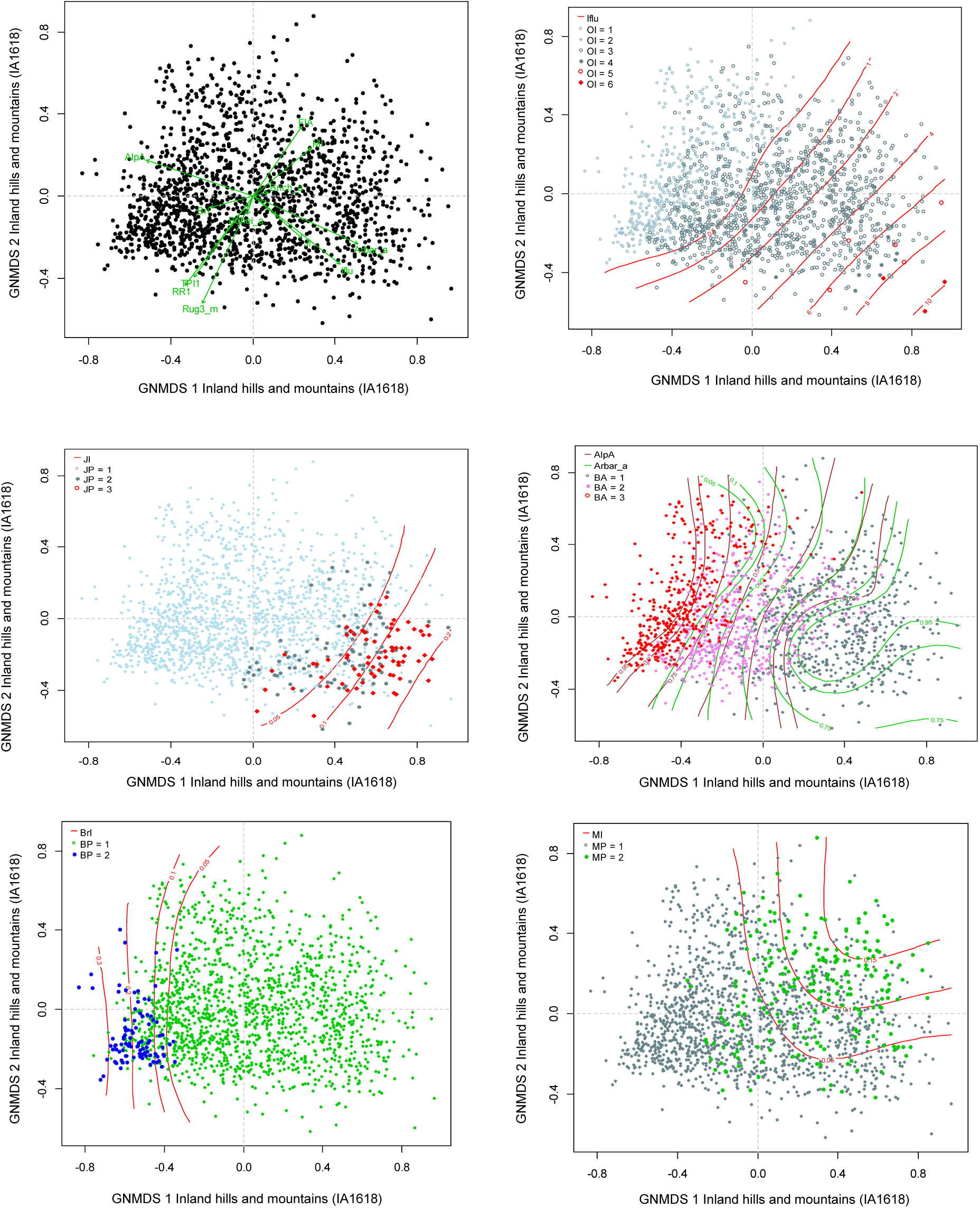
Distribution of observation units with specific properties in the GNMDS ordination diagram, axes 1 and 2, for the inland hills and mountains data set (IA1618). (91) Selected variables are represented by green arrows pointing in the direction of maximum increase of the variable (relative lengths of arrows are proportional to the correlation between the recorded variable and the ordination axis). Variable names are abbreviated according to Tables 4 and 5. (92) Isolines (red) for the infrastructure index (Iflu). OI = stepwise variation in the amount of infrastructure from low (OI 1–2) via medium (OI 3–4) to high amount of infrastructure, including towns and/or cities (OI 5–6). (93) Isolines (red) for agricultural land-use intensity (JI). JP = stepwise variation in agricultural land-use intensity from low (JP 1) via medium (JP2) to high (JP 3). (94) Isolines for the proportion of OU area situated above the forest line (AlpA; brown lines) and coniferous forest (Arbar_a; green lines). BA = stepwise variation along the gradient from boreal to alpine landscapes from areas dominated by forest (BA 1) via areas located at the forest line (BA 2) to mountainous areas above the forest line (BA 3). (95) Isolines (red) for the proportion of the OU covered by glaciers (BrI). BP1 = absence of glacier, BP2 = presence of glacier. (96) Isolines for the mire index (MI). MP1 = low abundance of mire; MP2 = high abundance of mire.

**Table 27.** Kendall’s rank correlation coefficients (τ), eigenvalues and gradient lengths for parallel DCA- and GNMDS-ordinations of the subset IA1618, Inland hills and mountains. S.D. units = standard deviation units; H.C. units = half-change units.

|  | GNMDS axis 1 | GNMDS axis 2 | GNMDS axis 3 | – |
| --- | --- | --- | --- | --- |
| Strongest correlated DCA-axis | DCA 1 | DCA 2 | DCA 3 | DCA 4 |
| $\tau$ (with strongest correlated DCA-axis) | 0.8884 | 0.7025 | 0.5634 | – |
| DCA-axis eigenvalue | 0.1975 | 0.0840 | 0.0580 | 0.0441 |
| DCA-axis gradient length (S.D. units) | 2.2627 | 1.6898 | 1.4818 | 1.3459 |
| GNMDS-axis gradient length (H.C. units) | 1.7949 | 1.4962 | 1.5590 | – |

Correlations between key and analytic variables on one hand, and GNMDS axis 1 scores on the other (Table 28), supported by vector diagrams (Fig. 91) and isoline diagrams (Figs 92–96), revealed a first axis that was very strongly related to a bio- and geo-ecological gradient in landscape element composition from mountains to lowlands, that also included variation in human land-use intensity. Along this composite gradient, the relative relief of the terrain decreased, the fractional area covered by forest and mire increased and the amount of infrastructure (including agricultural land-use intensity) increased sharply (Fig. 91). OUs with medium and high agricultural land-use intensity form a clearly differentiated group in the lower right corner of the ordination diagram (Fig. 93; JP > 1). The amount of infrastructure followed the same pattern but in a more gradual manner, increasing from the upper left (mountains) to the lower right (lowlands) in the diagram of GNMDS-axes 1 and 2 (Fig. 92). Vegetation cover followed the same pattern: OUs from alpine areas (BA = 3, Alp_a > 0.75), with high values for exposed bedrock (Kbf_a) and open treeless areas (Sn_flekk) prevailed at low GNMDS axis 1 values while coniferous (Arbar_a) and mixed (Arbla_a) boreal forests (BA = 1) prevailed at the high-score end of this GNMDS axis (Table 28; Figs 91 and 94).

**Table 28.**
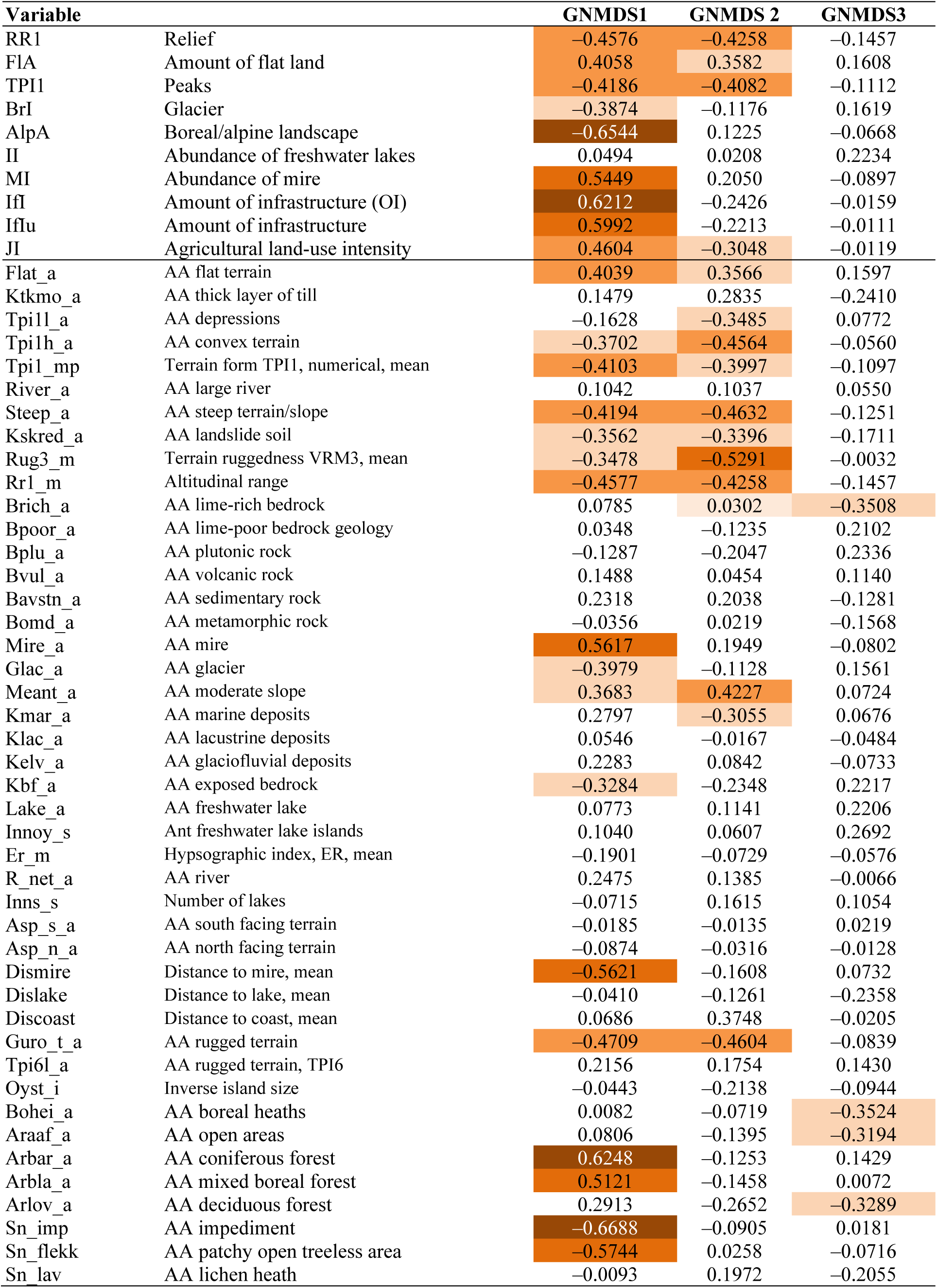
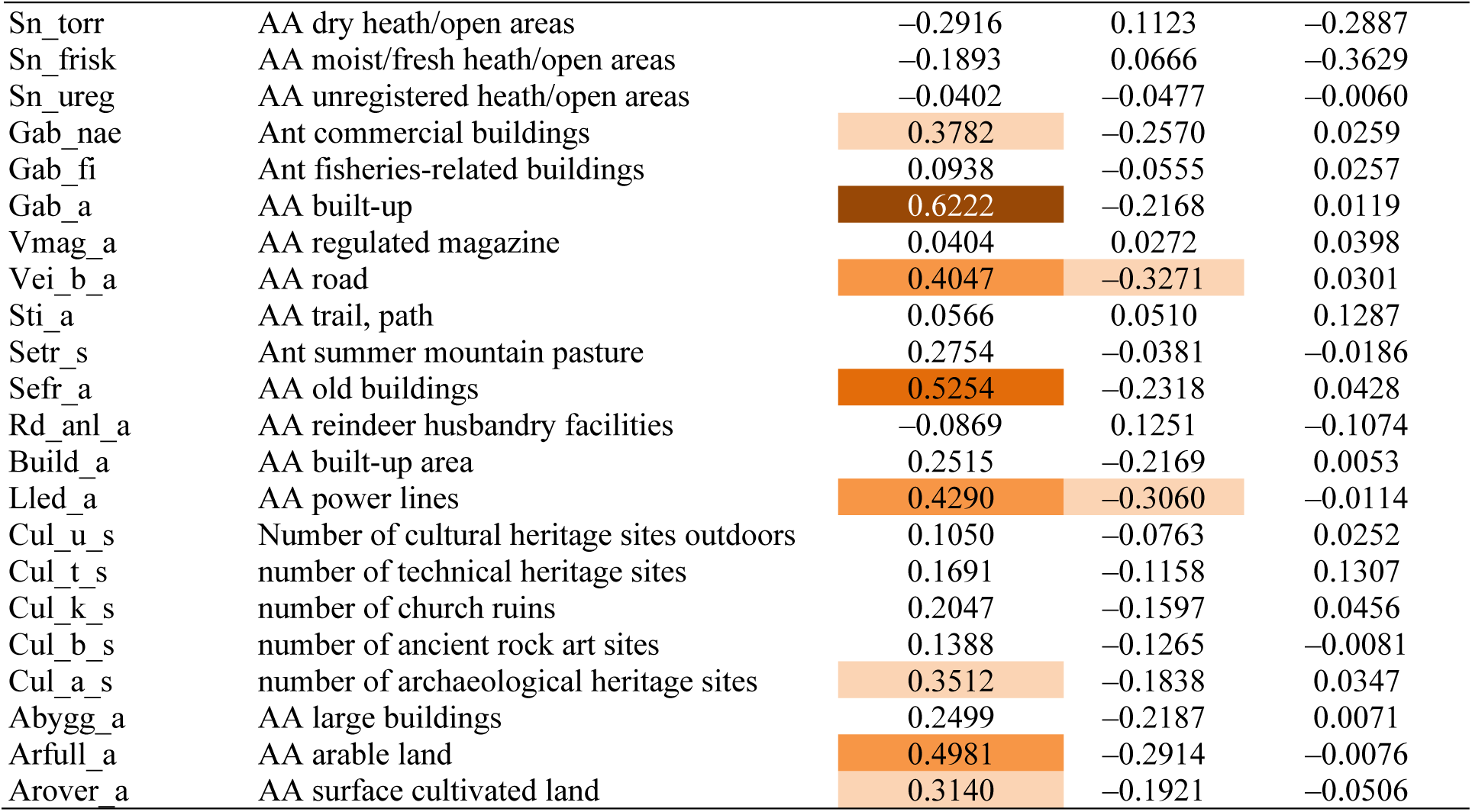
Kendall’s rank correlation coefficients (τ) between GNMDS ordination axes 1–3 for the data subset inland hills and mountains (IA 1618) and the 10 primary key variables (rows 1–10) and the 66 analytic variables used to characterise the 1618 observation units.

| Variable |  | GNMDS1 | GNMDS 2 | GNMDS3 |
| --- | --- | --- | --- | --- |
| RR1 | Relief | −0.4576 | −0.4258 | −0.1457 |
| FIA | Amount of flat land | 0.4058 | 0.3582 | 0.1608 |
| TPI1 | Peaks | −0.4186 | −0.4082 | −0.1112 |
| BrI | Glacier | −0.3874 | −0.1176 | 0.1619 |
| AlpA | Boreal/alpine landscape | −0.6544 | 0.1225 | −0.0668 |
| II | Abundance of freshwater lakes | 0.0494 | 0.0208 | 0.2234 |
| MI | Abundance of mire | 0.5449 | 0.2050 | −0.0897 |
| IfI | Amount of infrastructure (OI) | 0.6212 | −0.2426 | −0.0159 |
| IfIu | Amount of infrastructure | 0.5992 | −0.2213 | −0.0111 |
| JI | Agricultural land-use intensity | 0.4604 | −0.3048 | −0.0119 |
| Flat_a | AA flat terrain | 0.4039 | 0.3566 | 0.1597 |
| Ktkmo_a | AA thick layer of till | 0.1479 | 0.2835 | −0.2410 |
| Tpi1l_a | AA depressions | −0.1628 | −0.3485 | 0.0772 |
| Tpi1h_a | AA convex terrain | −0.3702 | −0.4564 | −0.0560 |
| Tpi1_mp | Terrain form TPI1, numerical, mean | −0.4103 | −0.3997 | −0.1097 |
| River_a | AA large river | 0.1042 | 0.1037 | 0.0550 |
| Steep_a | AA steep terrain/slope | −0.4194 | −0.4632 | −0.1251 |
| Kskred_a | AA landslide soil | −0.3562 | −0.3396 | −0.1711 |
| Rug3_m | Terrain ruggedness VRM3, mean | −0.3478 | −0.5291 | −0.0032 |
| Rr1_m | Altitudinal range | −0.4577 | −0.4258 | −0.1457 |
| Brich_a | AA lime-rich bedrock | 0.0785 | 0.0302 | −0.3508 |
| Bpoor_a | AA lime-poor bedrock geology | 0.0348 | −0.1235 | 0.2102 |
| Bplu_a | AA plutonic rock | −0.1287 | −0.2047 | 0.2336 |
| Bvul_a | AA volcanic rock | 0.1488 | 0.0454 | 0.1140 |
| Bavstn_a | AA sedimentary rock | 0.2318 | 0.2038 | −0.1281 |
| Bomd_a | AA metamorphic rock | −0.0356 | 0.0219 | −0.1568 |
| Mire_a | AA mire | 0.5617 | 0.1949 | −0.0802 |
| Glac_a | AA glacier | −0.3979 | −0.1128 | 0.1561 |
| Meant_a | AA moderate slope | 0.3683 | 0.4227 | 0.0724 |
| Kmar_a | AA marine deposits | 0.2797 | −0.3055 | 0.0676 |
| Klac_a | AA lacustrine deposits | 0.0546 | −0.0167 | −0.0484 |
| Kelv_a | AA glaciofluvial deposits | 0.2283 | 0.0842 | −0.0733 |
| Kbf_a | AA exposed bedrock | −0.3284 | −0.2348 | 0.2217 |
| Lake_a | AA freshwater lake | 0.0773 | 0.1141 | 0.2206 |
| Innoy_s | Ant freshwater lake islands | 0.1040 | 0.0607 | 0.2692 |
| Er_m | Hypsographic index, ER, mean | −0.1901 | −0.0729 | −0.0576 |
| R_net_a | AA river | 0.2475 | 0.1385 | −0.0066 |
| Inns_s | Number of lakes | −0.0715 | 0.1615 | 0.1054 |
| Asp_s_a | AA south facing terrain | −0.0185 | −0.0135 | 0.0219 |
| Asp_n_a | AA north facing terrain | −0.0874 | −0.0316 | −0.0128 |
| Dismire | Distance to mire, mean | −0.5621 | −0.1608 | 0.0732 |
| Dislake | Distance to lake, mean | −0.0410 | −0.1261 | −0.2358 |
| Discoast | Distance to coast, mean | 0.0686 | 0.3748 | −0.0205 |
| Guro_t_a | AA rugged terrain | −0.4709 | −0.4604 | −0.0839 |
| Tpi6l_a | AA rugged terrain, TPI6 | 0.2156 | 0.1754 | 0.1430 |
| Oyst_i | Inverse island size | −0.0443 | −0.2138 | −0.0944 |
| Bohei_a | AA boreal heaths | 0.0082 | −0.0719 | −0.3524 |
| Araaf_a | AA open areas | 0.0806 | −0.1395 | −0.3194 |
| Arbar_a | AA coniferous forest | 0.6248 | −0.1253 | 0.1429 |
| Arbla_a | AA mixed boreal forest | 0.5121 | −0.1458 | 0.0072 |
| Arlov_a | AA deciduous forest | 0.2913 | −0.2652 | −0.3289 |
| Sn_imp | AA impediment | −0.6688 | −0.0905 | 0.0181 |
| Sn_flekk | AA patchy open treeless area | −0.5744 | 0.0258 | −0.0716 |
| Sn_lav | AA lichen heath | −0.0093 | 0.1972 | −0.2055 |
| Sn_torr | AA dry heath/open areas | -0.2916 | 0.1123 | -0.2887 |
| Sn_frisk | AA moist/fresh heath/open areas | -0.1893 | 0.0666 | -0.3629 |
| Sn_ureg | AA unregistered heath/open areas | -0.0402 | -0.0477 | -0.0060 |
| Gab_nae | Ant commercial buildings | 0.3782 | -0.2570 | 0.0259 |
| Gab_fi | Ant fisheries-related buildings | 0.0938 | -0.0555 | 0.0257 |
| Gab_a | AA built-up | 0.6222 | -0.2168 | 0.0119 |
| Vmag_a | AA regulated magazine | 0.0404 | 0.0272 | 0.0398 |
| Vei_b_a | AA road | 0.4047 | -0.3271 | 0.0301 |
| Sti_a | AA trail, path | 0.0566 | 0.0510 | 0.1287 |
| Setr_s | Ant summer mountain pasture | 0.2754 | -0.0381 | -0.0186 |
| Sefr_a | AA old buildings | 0.5254 | -0.2318 | 0.0428 |
| Rd_anl_a | AA reindeer husbandry facilities | -0.0869 | 0.1251 | -0.1074 |
| Build_a | AA built-up area | 0.2515 | -0.2169 | 0.0053 |
| Lled_a | AA power lines | 0.4290 | -0.3060 | -0.0114 |
| Cul_u_s | Number of cultural heritage sites outdoors | 0.1050 | -0.0763 | 0.0252 |
| Cul_t_s | number of technical heritage sites | 0.1691 | -0.1158 | 0.1307 |
| Cul_k_s | number of church ruins | 0.2047 | -0.1597 | 0.0456 |
| Cul_b_s | number of ancient rock art sites | 0.1388 | -0.1265 | -0.0081 |
| Cul_a_s | number of archaeological heritage sites | 0.3512 | -0.1838 | 0.0347 |
| Abygg_a | AA large buildings | 0.2499 | -0.2187 | 0.0071 |
| Arfull_a | AA arable land | 0.4981 | -0.2914 | -0.0076 |
| Arover_a | AA surface cultivated land | 0.3140 | -0.1921 | -0.0506 |

Isolines for the proportion of lime-rich bedrock (Brich_a; Fig. 98) show a similar, but weaker pattern. OUs with glacier (Fig. 95; BP = 2) were restricted to the low-score end of GNMDS axis 1, where they occurred intermixed with OUs with glacier. Unlike in inland valleys, OUs with glacier occurred in inland hills and mountains without any relationship to terrain variation (i.e. along GNMDS axis 2).

GNMDS axis 2 was clearly related to (residual variation in) relief. Variation in the abundance of mires followed the variation in relief (Fig. 96). OUs with distinct peaks (Fig 97; TP = 2) to some extent followed the same pattern as glaciers and occurred intermixed with other OUs with high relief at low GNMDS axis 2 (and GNMDS axis 1) scores.

**Figs. 97–98.**
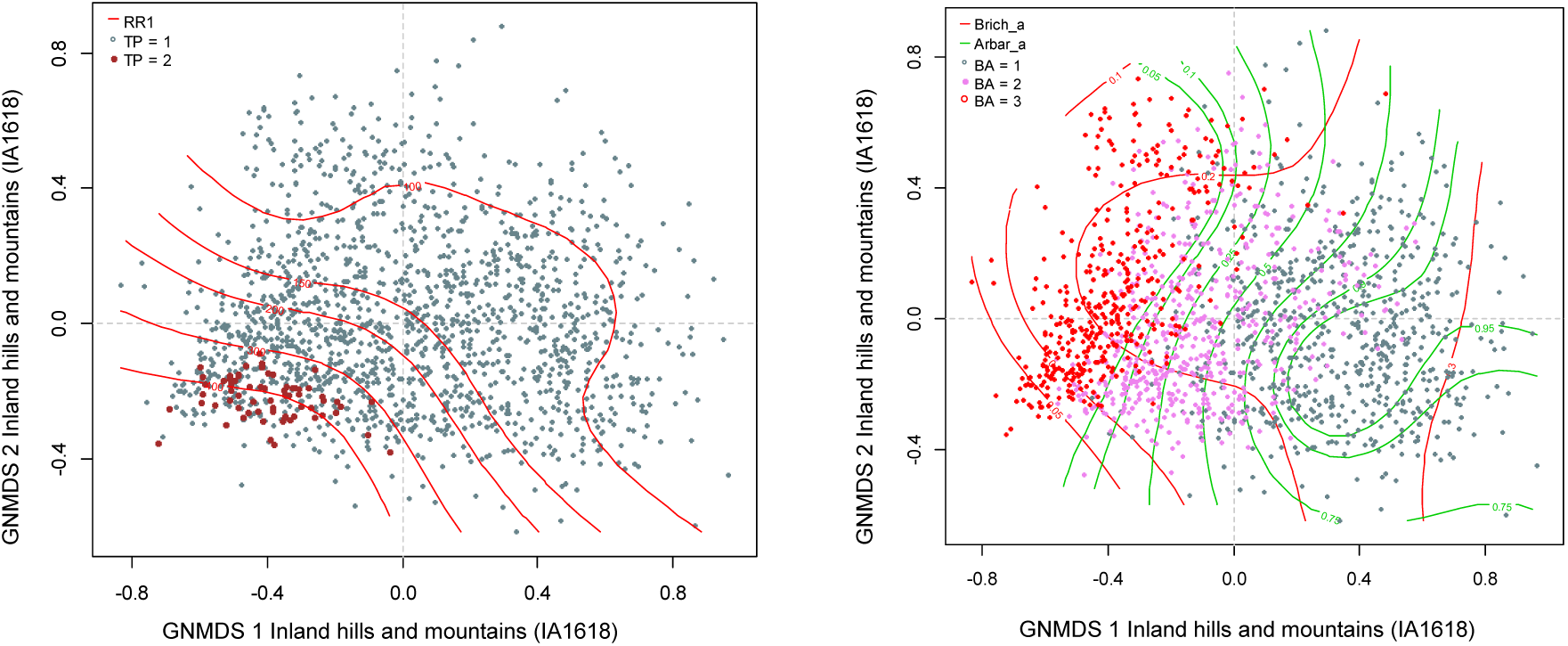
Distribution of observation units with specific properties in the GNMDS ordination diagram, axes 1 and 2, for the inland hills and mountains data set (IA1618). (97) Isolines (red) for relative relief (RR1). TP = presence (2) or absence (1) of peaks. (98) Isolines for the fraction of OU area covered by lime-rich bedrock (Brich_a; red lines) and coniferous forest (Arbar_a; green lines). BA = stepwise variation along the gradient from boreal to alpine landscapes from areas dominated by forest (BA 1) via areas located at the forest line (BA 2) to mountainous areas above the forest line (BA 3).

**Figs. 99–100.**
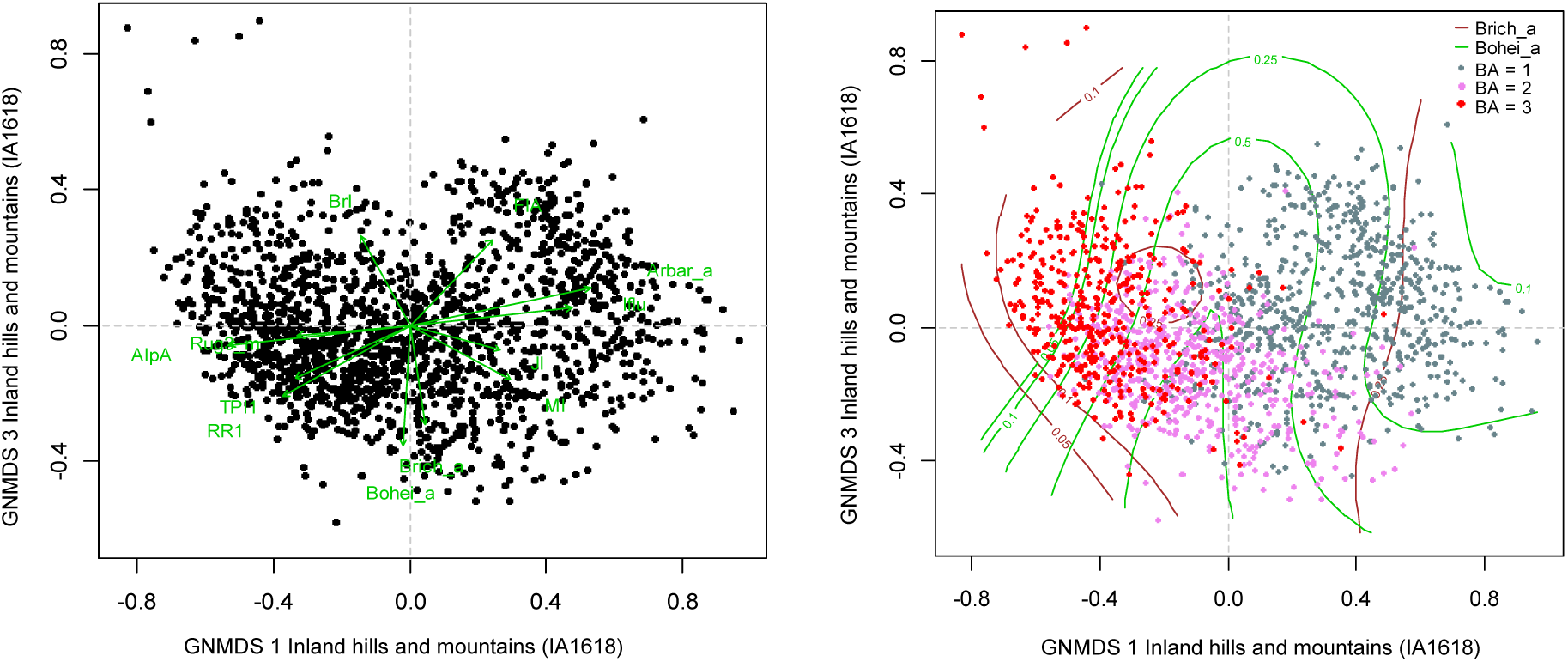
Distribution of observation units with specific properties in the GNMDS ordination diagram, axes 1 and 3, for the inland hills and mountains data set (IA1618). Variable names are abbreviated according to Tables 4 and 5. (99) Selected variables represented by green arrows pointing in the direction of maximum increase of the variable (relative lengths of arrows are proportional to the correlation between the recorded variable and the ordination axis). (100) Isolines for the fraction of OU area covered by lime-rich bedrock (Brich_a; brown lines) and boreal heath (Bohei_a; green lines). BA = stepwise variation along the gradient from boreal to alpine landscapes from areas dominated by forest (BA 1) via areas located at the forest line (BA 2) to mountainous areas above the forest line (BA 3).

GNMDS axis 3 was correlated with few analysis variables, and only weakly so. No clear, easily interpretable pattern was observed along this axis, which identified six OUs with outlier positions in the upper left corner of the ordination diagram (low scores for GNMDS axis 1 and high scores for GNMDS axis 3).

Based on ordination analyses and interpretation, we identified the following 8 CLG candidates in the three functional variables categories G, U and A (thematically slightly marginal variables are highlighted in grey; the absolute value of the correlation coefficient with the relevant GNMDS axis in parentheses):

(G1) Relief (Valley Cutting) (RE; GNMDS axis 1/2): Rug3_m (0.5291), Guro_t_a (0.4709), Steep_a (0.4632), RR1 (0.4576), Tpi1h_a (0.4564), Meant_a (0.4227), TPI1 (0.4186), Tpi1_mp (0.4103), FlA (0.4058)
(G2) Abundance of mire (MP; GNMDS axis 1): Dismire (0.5621), Mire_a (0.5617), MI (0.5449)
(G3) Glacier (BP; GNMDS axis 1) Glac_a (0.3979), BrI (0.3874)
(U1) Vegetation (forest) cover (SkP; GNMDS axis 1): Sn_imp (0.6688), AlpA (0.6544), Arbar_a (0.6248), Sn_flekk (0.5744), Arbla_a (0.5121)
(U2) Open area cover (AP; GNMDS axis 3): Bohei_a (0.3524), Araaf_a (0.3194) (U3) Geological richness (GR; GNMDS axis 3) Brich_a (0.3508)
(A1) Infrastructure (OI; GNMDS axis 1): Gab_a (0.6222), IfIu (0.5992), Sefr_a (0.5254), Lled_a (0.4290), Vei_b_a (0.4047)
(A2) Degree of agricultural intensity (JP; GNMDS axis 1): Arfull_a (0.5906), JI (0.4604), Arover_a (0.3659)

*Comments:* For U1, the value of the correlation coefficient τ (Arlov_a, Bohei_a) = 0.3142 shows that deciduous forest (Arlov_a) is not part of the same group as boreal heath. Since other vegetation-cover variables were correlated with GNMDS axis 1, and the correlation between Arlov_a and the other forest variables was weak, Arlov_a is not accepted as separate CLG candidate. U2 consists of the two variables open areas (Araaf_a) and boreal heaths (Bohei_a), which were connected by τ (Araaf_a, Bohei_a) = 0.4331. U2 and U3 form two separate groups because the geological lime-richness and boreal heaths were uncorrelated (τ (Brich_a, Bohei_a) = 0.0240.

### ANALYSES FOR ESTABLISHMENT AND STANDARDISED SEGMENTATION OF A PARSIMONIOUS SET OF CLGS FOR EACH MAJOR TYPE

#### Coastal plains (KS)

The analyses for identification of independent significant analytic variables for each CLG candidate by forward selection from the *n* = 84 variables using RDA show that all nine CLG candidates satisfied the requirement for explaining at least 6% of the variation (Table 29). The parsimonious set of CLGs, found by RDA with forward selection among the CLG candidates separately for each functional variable category, contained 5 CLGs (Table 29). The properties of the five CLGs and their associated (ranked primary) ONVs are described below on the basis of Tables 30–34 and Figs 101–113.

**Fig. 101.**
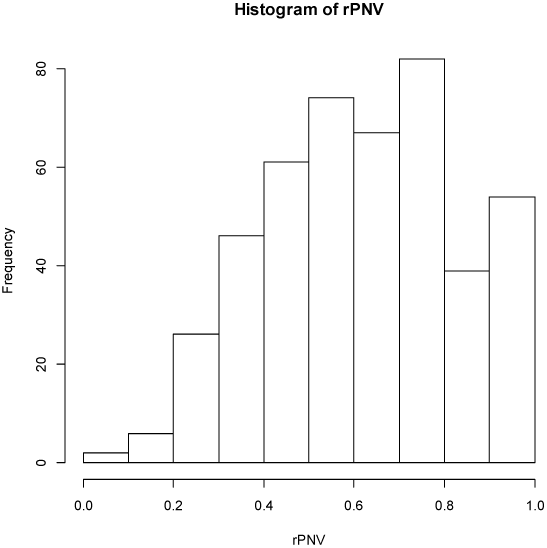
Major type coastal plains (KS): candidate complex-landscape gradient (CLG) 1, inner-outer coast (IYK): distribution of OUs on the ranged but not rescaled orthogonal key variable (ONV).

**Figs. 102–104.**
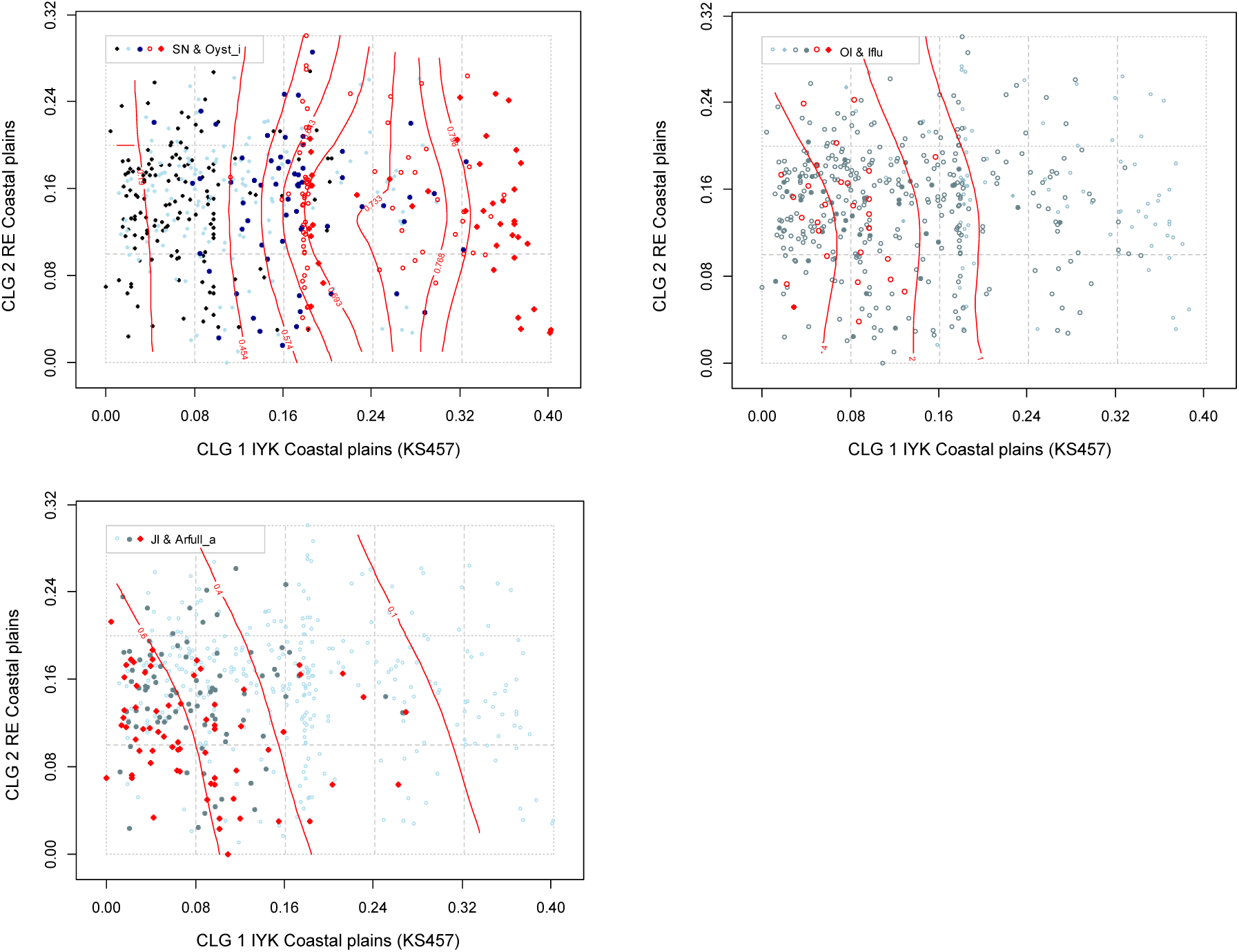
Major type coastal plains (KS): distribution of 457 observation units (KS457 data subset) along rescaled orthogonal key variables (rONVs) for complex landscape gradients CLG 1 (inner–outer coast; IYK) and CLG 2 (relief; RE). Symbols show affiliation of OUs to levels of relevant landscape-gradient (LG) variables, operationalised as ordered factor variables in accordance with Table 3. Isolines are given for relevant primary key variables (Table 4) and analytic variables (Table 5).

**Fig. 105.**
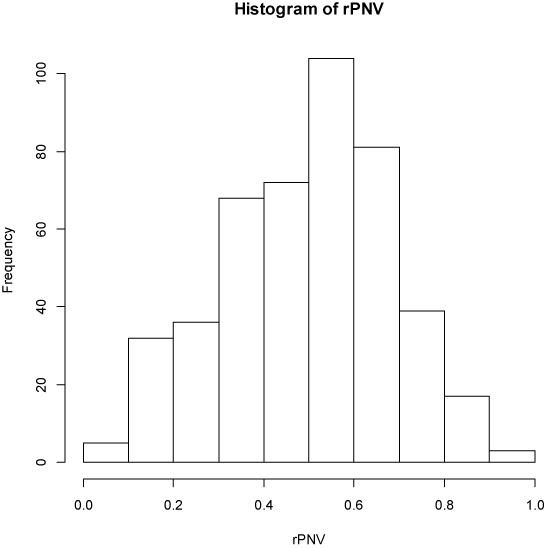
Major type coastal plains (KS): candidate complex-landscape gradient (CLG) 2, relief (RE): distribution of OUs on the ranged but not rescaled orthogonal key variable (ONV).

**Figs. 106–107.**
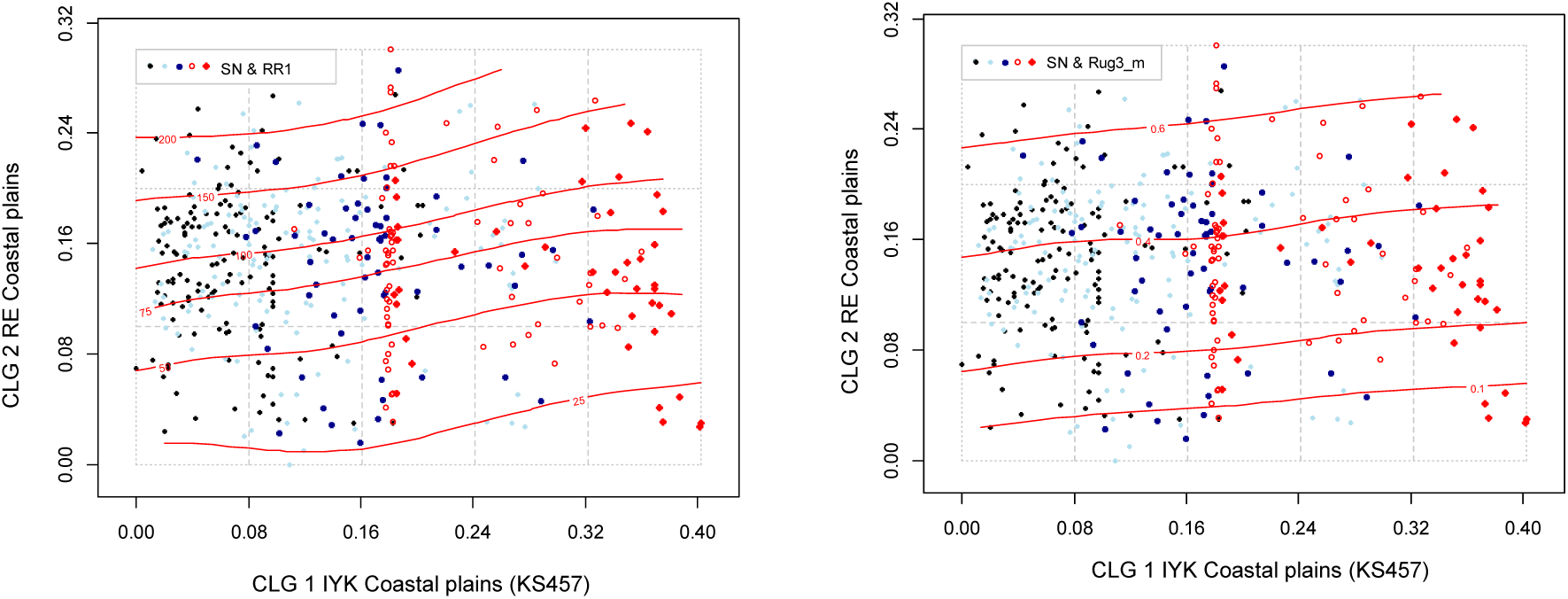
Major type coastal plains (KS): distribution of 457 observation units (KS457 data subset) along rescaled orthogonal key variables (rONVs) for complex landscape gradients CLG 1 (inner–outer coast; IYK) and CLG 2 (relief; RE). Symbols show affiliation of OUs to levels of relevant landscape-gradient (LG) variables, operationalised as ordered factor variables in accordance with Table 3. Isolines are given for relevant primary key variables (Table 4) and analytic variables (Table 5).

**Fig. 108.**
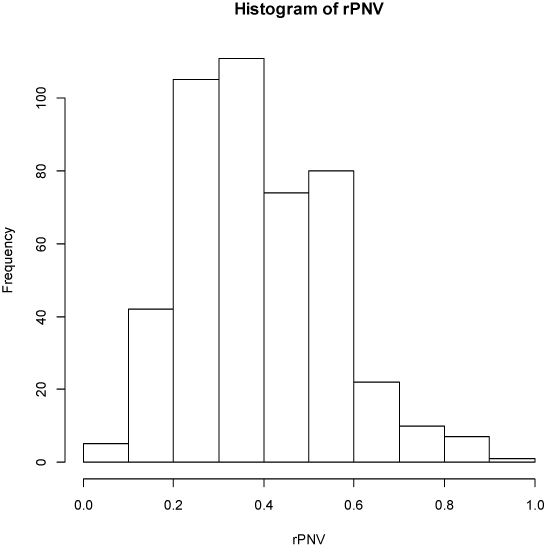
Major type coastal plains (KS): candidate complex-landscape gradient (CLG) 3, abundance of wetlands (VP): distribution of OUs on the ranged but not rescaled orthogonal key variable (ONV).

**Figs. 109–110.**
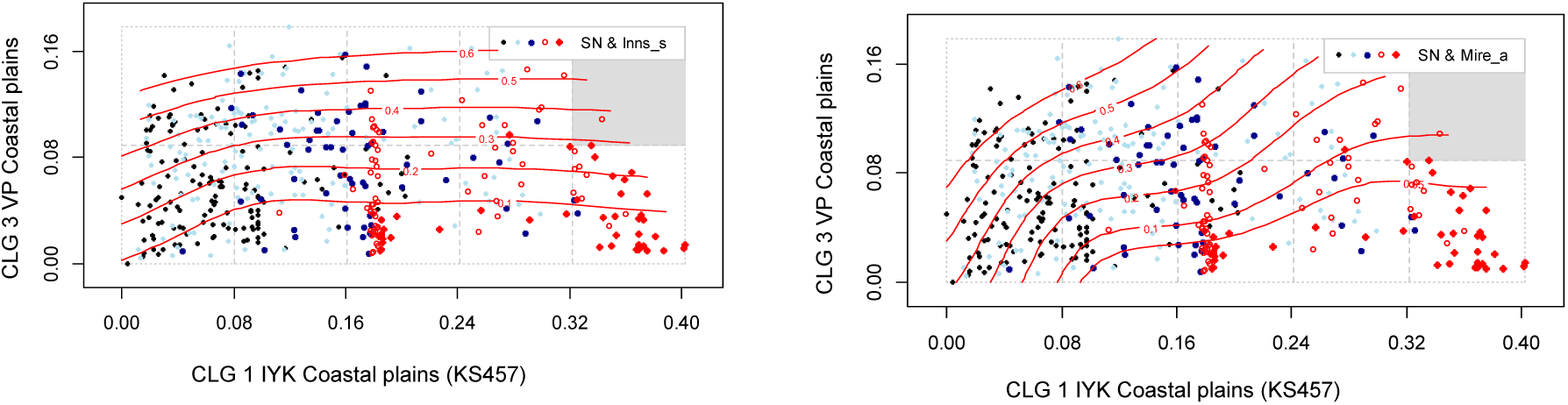
Major type coastal plains (KS): distribution of 457 observation units (KS457 data subset) along rescaled orthogonal key variables (rONVs) for complex landscape gradients CLG 1 (inner–outer coast; IYK) and CLG Abundance of wetlands; VP). Symbols show affiliation of OUs to levels of relevant landscape-gradient (LG) variables, operationalised as ordered factor variables in accordance with Table 3. Isolines are given for relevant primary key variables (Table 4) and analytic variables (Table 5).

**Fig. 111.**
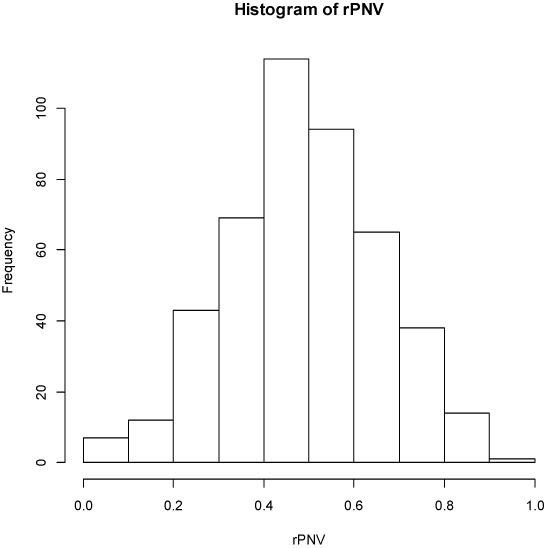
Major type coastal plains (KS): candidate complex-landscape gradient (CLG) 4, forest cover): distribution of OUs on the ranged but not rescaled orthogonal key variable (ONV).

**Fig. 112.**
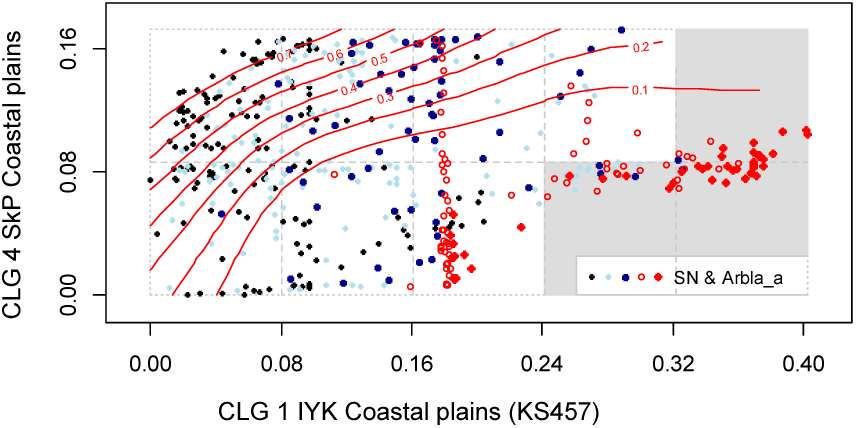
Major type coastal plains (KS): distribution of 457 observation units (KS457 data subset) along rescaled orthogonal key variables (rONVs) for complex landscape gradients CLG 1 (inner–outer coast; IYK) and CLG 4 forest cover; SkP). Symbols show affiliation of OUs to levels of relevant landscape-gradient (LG) variables, operationalised as ordered factor variables in accordance with Table 3. Isolines are given for relevant primary key variables (Table 4) and analytic variables (Table 5).

**Fig. 113.**
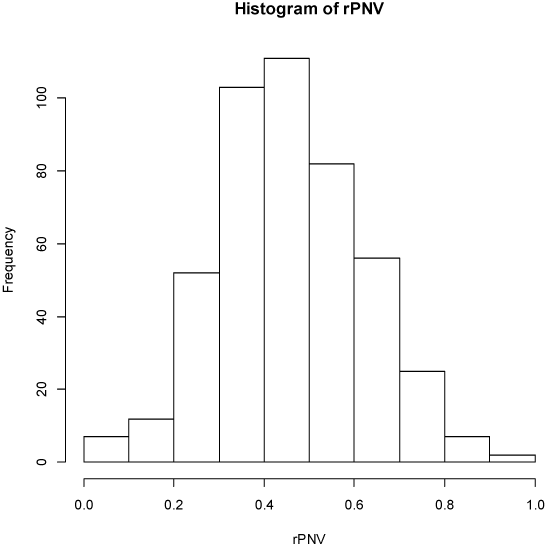
Major type coastal plains (KS): candidate complex-landscape gradient (CLG) 5, amount of infrastructure (OI): distribution of OUs on the ranged but not rescaled orthogonal key variable (ONV).

**Table 29.**
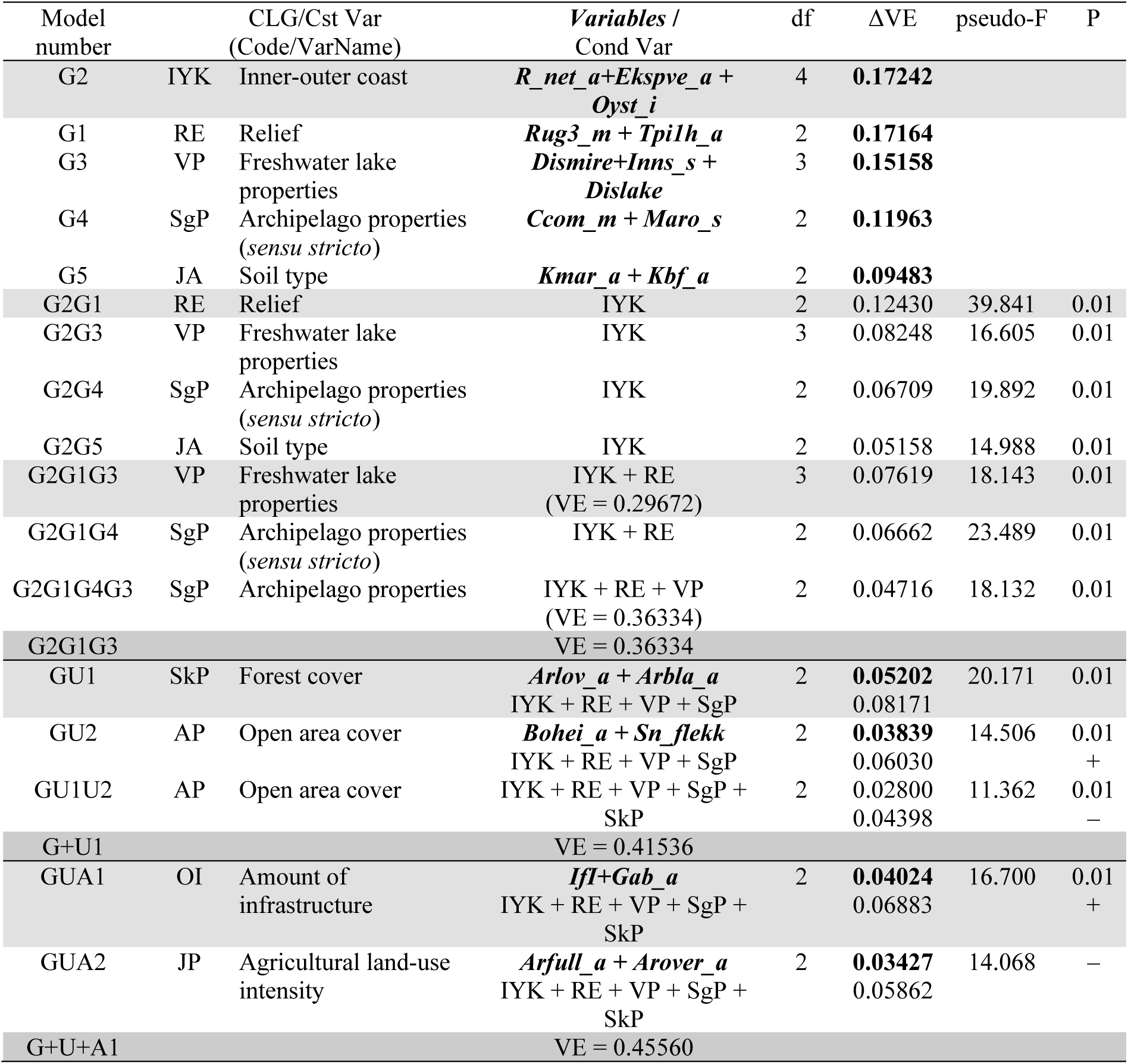
Identification of CLG candidates in coastal plains by forward selection of variables using RDA. Results of tests of single CLG candidates are shown in bold-face types. The best model at each step in the selection process is highlighted using light grey shading while the best model after testing all CLG candidates in a functional variable category is highlighted using darker grey shading. Abbreviations and explanations: Var = variable; Cst Var = constraining variable; Cond Var = conditioning variable(s); df = degrees of freedom; VE = variation explained (expressed as fraction of the total variation); ΔVE = additional variation explained; pseudo-F and P = F-statistic used in the test of the null hypothesis that adding the variable(s) does not contribute more to explaining variation in landscape-element composition than a random variable, with the associated P value. Variable names are abbreviated according to Tables 4 and 5. For further explanation, se Material and methods chapter.

**Table 30.** Major type coastal plains (KS): key properties* and segmentation** of candidate complex-landscape gradient G1, inner-outer coast (IYK), as represented by its orthogonal key variable (ONV). OU = observation unit; Threshold = upper limit of class (RSC = rescaling). PD unit = Proportional dissimilarity unit. Segment threshold shows position of standard segment borders relative to initial segments.

|  | Initial CLG segment |  |  |  |  |  |  |  |
| --- | --- | --- | --- | --- | --- | --- | --- | --- |
|  | 1 | 2 | 3 | 4 | 5 | 6 | 7 | 8 |
| Number of OUs | 25 | 48 | 59 | 77 | 73 | 82 | 37 | 56 |
| ONV threshold before RSC | 0.2789 | 0.3820 | 0.4850 | 0.5888 | 0.6952 | 0.7969 | 0.8976 | 1 |
| ONV threshold after RSC | 0.0544 | 0.1113 | 0.1855 | 0.2742 | 0.4170 | 0.5224 | 0.6858 | 1 |
| ONV threshold, in PD units | 0.0219 | 0.0448 | 0.0747 | 0.1104 | 0.1679 | 0.2103 | 0.2761 | 0.4026 |
| Segment threshold |  |  |  | 12 | 23 |  | 34 | 45 |

\*Variation explained by variable group: 0.1724, by PNV and ONV: 1158
Gradient length estimates: Method I: 0.4411; Method II: 0.1948; Method III: 0.4026
\*\* Tentative number of standard segments = 5
Number of OUs per standard segment: 144 – 128 – 107 – 39 – 35

**Table 31.**
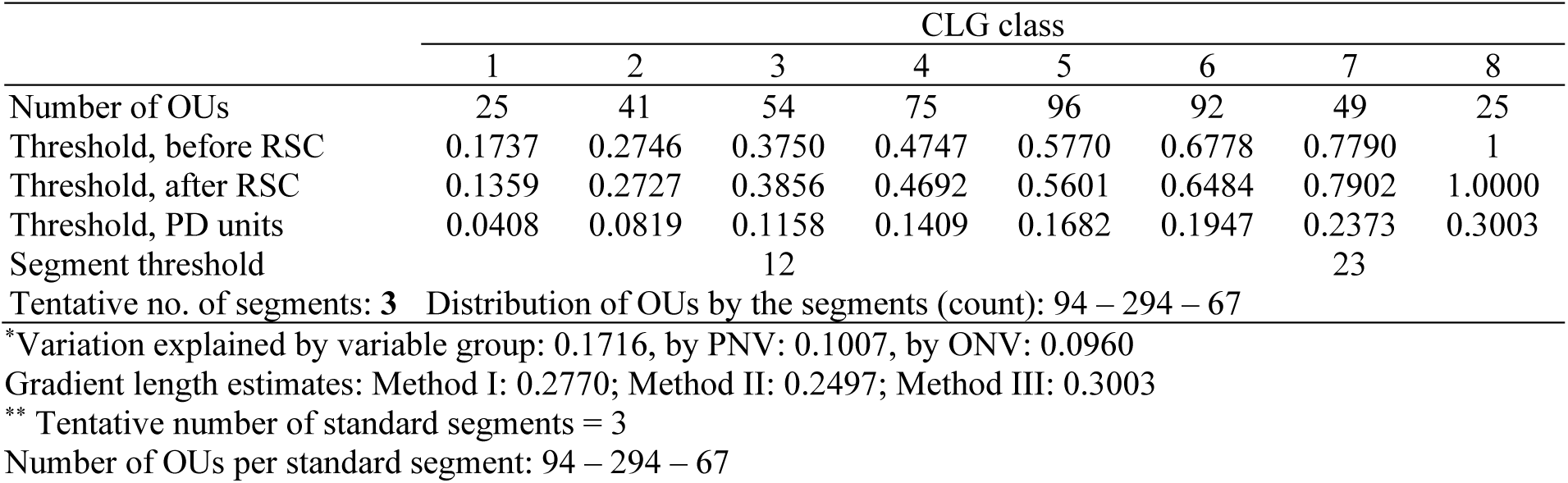
Major type coastal plains (KS): key properties* and segmentation** of candidate complex-landscape gradient G2, relief (RE), as represented by its orthogonal key variable (ONV). OU = observation unit; Threshold = upper limit of class (RSC = rescaling). PD unit = Proportional dissimilarity unit. Segment threshold shows position of standard segment borders relative to initial segments.

**Table 32.**
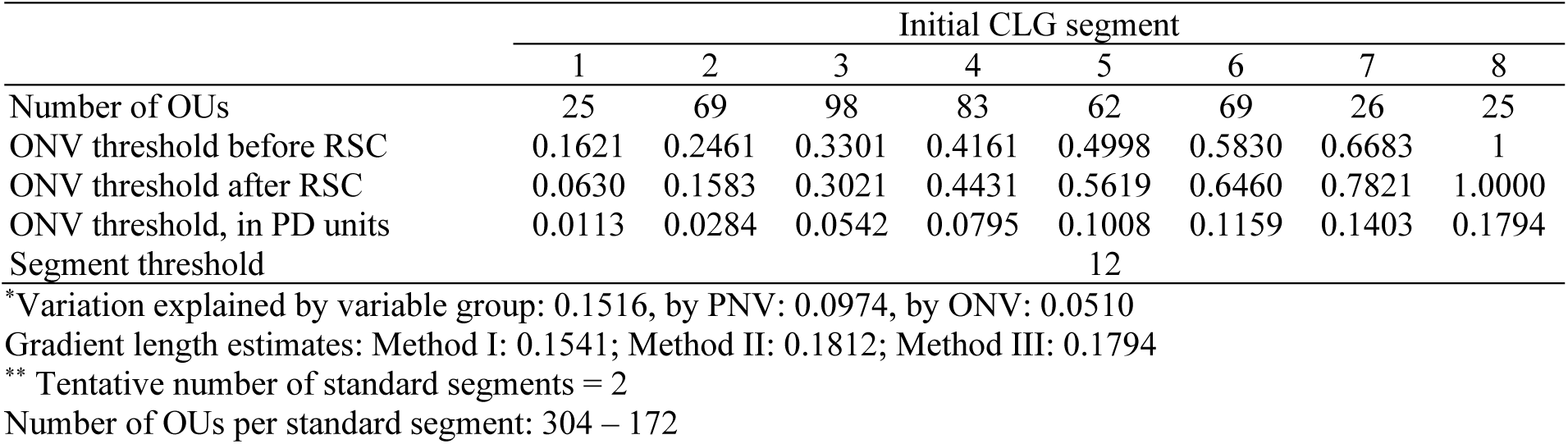
Major type coastal plains (KS): key properties* and segmentation** of candidate complex-landscape gradient G3, abundance of wetlands (VP), as represented by its orthogonal key variable (ONV). OU = observation unit; Threshold = upper limit of class (RSC = rescaling). PD unit = Proportional dissimilarity unit. Segment threshold shows position of standard segment borders relative to initial segments.

**Table 33.**
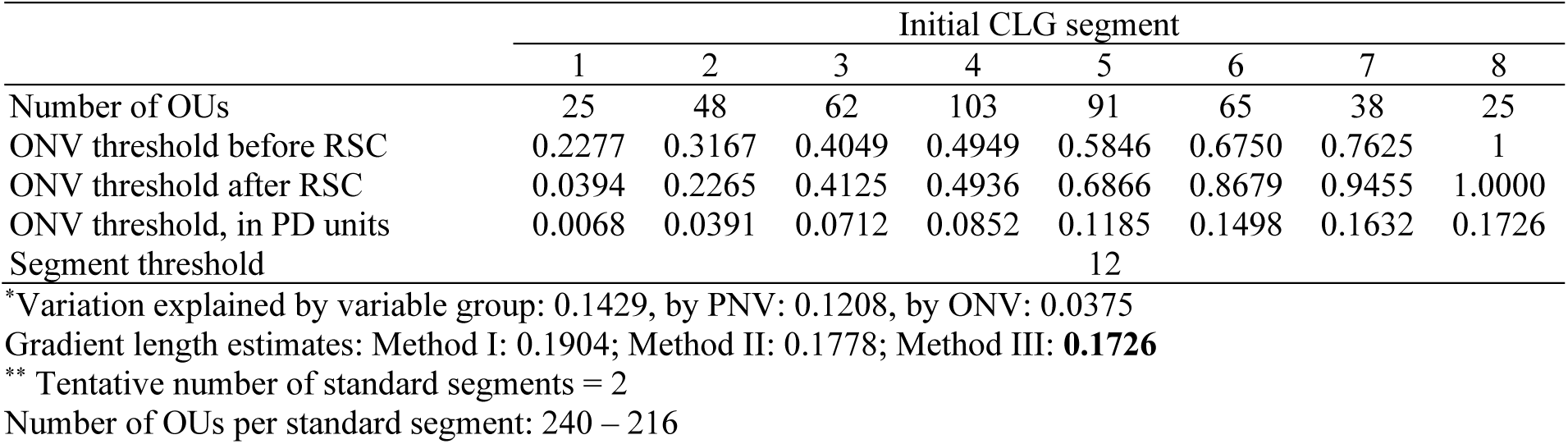
Major type coastal plains (KS): key properties* and segmentation** of candidate complex-landscape gradient U1 Forest cover (SkP), as represented by its orthogonal key variable (ONV). OU = observation unit; Threshold = upper limit of class (RSC = rescaling). PD unit = Proportional dissimilarity unit. Segment threshold shows position of standard segment borders relative to initial segments.

**Table 34.**
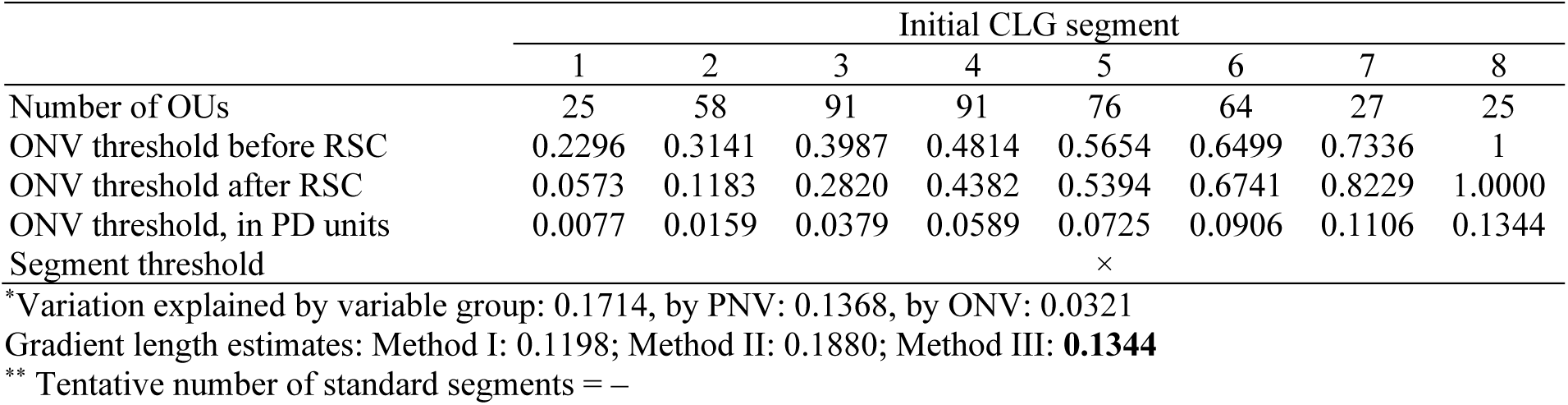
Major type coastal plains (KS): key properties* and segmentation** of candidate complex-landscape gradient A1, amount of infrastructure (OI), as represented by its orthogonal key variable (ONV). OU = observation unit; Threshold = upper limit of class (RSC = rescaling). PD unit = Proportional dissimilarity unit. Segment threshold shows position of standard segment borders relative to initial segments.

##### CLG 1 – (G1) Inner-outer coast: IYK (Table 30, Figs 101–104)

The three variables included in the ONV that represents CLG 1, inner-outer coast (IYK), were: (1) number of rivers (R_net_a); (2) areal coverage of exposed coast (Ekspve_a); and (3) inverse island size (Oyst_i). This geo-ecological CLG had an estimated gradient length of 0.4026/0.08 = 5.033 EDU–L (ecodiversity distance units in landscapes) and was, accordingly, divided into 5 standard segments (Table 30). The distribution of OUs along this ONV (in its original scaling) was left-skewed (Fig. 101).

This CLG expresses variation in coastal landscapes from areas with ‘inland properties’ on the inner side of larger islands, hardly exposed to the harsh conditions of the open sea (protected inner coast), to the outer coast, directly exposed to the actions of wind, waves and ocean currents, i.e. strongly wave-exposed outer coast (Figs 102–104). The intermediate segments are termed: moderately protected coast, moderately wave-exposed coast; and wave-exposed outer coast.

Differences in landscape-element composition among OUs from the outer coast (relatively small islands, SN-classes 3–5; cf. Fig. 102, Table 30), which was represented in the data subset by relatively few OUs, contributed strongly to gradient length. The gradient length estimate is therefore uncertain.

A closer examination of variation along the rescaled ONV for this CLG (Fig. 102) motivated for using the following logarithmic sequence of (largest) island sizes as criteria for separating standard segments 4 and 5, 3 and 4, 2 and 3, respectively: 1 km²; 10 km^2^; 100 km². The border between standard segments 1 and 2 was less distinct in terms of island vs. mainland properties.

Infrastructure and agricultural land use followed CLG 1 IYK closely (Figs 103–104), suggesting that most of the variation in infrastructure and agricultural land-use intensity was also captured by IYK. With few exceptions the outer coast (IYK 4–5) lacked landscape elements typical of agricultural land use and had very limited infrastructure. Land-use intensity (the amount of infrastructure) increased most strongly at the transition from IYK·3 to IYK·2.

##### CLG 2 – (G2) Relief in coastal plains: RE (Table 31, Figs 105–107)

The two variables included in the ONV that represents CLG 2, relief (RE), were: (1) mean terrain ruggedness (Rug3_m) and (2) areal coverage of convex terrain (Tpi1h_a). This geo-ecological CLG had an estimated gradient length of 0.3003/0.08 = 3.754 EDU–L units and was, accordingly, divided into 3 standard segments (Table 31). The distribution of OUs along this ONV (in its original scaling) was slightly left-skewed (Fig. 105).

This CLG expresses variation in terrain-form within coastal plains from (relatively) flat coastal plains (via moderately rugged coastal plains) to rugged coastal plains, often with cliffs or remnant peaks. The CLG had good linearity, with an even distribution of OUs along the CLG (Figs 106–107). A slight increase in PD between adjacent initial CLG segments in both directions from the middle of the candidate CLG indicates decreasing degree of generalisation towards gradient endpoints (Table 31). A clear relationship between relative relief (RR1) and the ONV was found: RR1 values of 50 m and 125 m corresponded to transitions between RE standard segments 1–2, and 2–3, respectively. Inspection of OUs (results not shown) revealed presence of residual mountains (Norwegian geomorphological term = ‘*restfjell*’) in a large majority of OUs with RE·3. The isolines for mean terrain ruggedness (Rug3_m; Fig. 107) run parallel with the ONV_RE, in accordance with Rug3_m being the single variable that was most strongly correlated with ONV_RE.

##### CLG 3 – (G3) Abundance of wetlands: VP (Table 32, Figs 108–110)

The three variables included in the ONV that represents CLG 3, abundance of wetlands (VP), were: (1) mean distance to mire (Dismire); (2) number of lakes (Inns_s); and (3) mean distance to lake (Dislake). This geo-ecological CLG had an estimated gradient length of 0.1794/0.08 = 2.243 EDU–L units and was, accordingly, divided into 2 standard segments (Table 32). The distribution of OUs along this ONV (in its original scaling) was right-skewed (Fig. 108).

This CLG expresses variation in the areal cover of wetlands (including mires) and the abundance of small lakes and tarns, which are often associated with wetlands. The two segments are characterised by low to medium abundance (VP·1) and high abundance (VP·2), respectively, of wetlands and associated ecosystems (Figs 109–110).

We identified some nonlinearity in initial segments 6–7 (cf. Table 32). The increase in PD between adjacent initial CLG segments from mid-gradient towards the gradient’s upper end (i.e. with increasing abundance of lakes and wetlands) indicates decreasing generalisation. This suggests that there is some variation in lake occurrence also on small islands (Fig. 108), while mires are more or less completely confined to larger islands and the mainland (Fig. 109). The variation along this CLG is most clearly expressed by the number of lakes, as indicated by isolines for the Inns_s variable in Fig. 109.

##### CLG 4 – (U1) Forest cover: SkP (Table 33, Figs 111–112)

The two variables included in the ONV that represents CLG 4, forest cover (SkP), were: (1) areal coverage of deciduous forest (Arlov) and (2) areal coverage of mixed forest (Arbla_a). This bio-ecological CLG had an estimated gradient length of 0.1726/0.08 = 2.158 EDU–L units and was, accordingly, divided into 2 standard segments (Table 33). The distribution of OUs along this ONV (in its original scaling) was unimodal and symmetric (Fig. 111).

This CLG expresses variation in forest cover from bare rock without or with sparse forest cover, e.g. or heaths (SkP·1) to forested areas (SkP·2) (Fig. 112).

The increase in PD between adjacent initial CLG segments from mid-gradient towards the lower end of the gradient (open landscapes) indicates decreasing degree of generalisation towards this gradient endpoint (Table 33). Variation in the proportion of OU area covered with forest (e.g. coniferous forest cover; Arbla_a; Fig. 112) is primarily found on the mainland and on the largest islands.

##### CLG 5 – (A1) Amount of infrastructure: OI (Table 34, Figs 103 and 113)

The two variables included in the ONV that represents CLG 5, amount of infrastructure (OI), were: (1) the infrastructure index (IfI) and (2) areal coverage of built-up areas (Gab_a). This land-use related CLG had an estimated gradient length of 0.1344/0.08 = 1.68 EDU–L units and was, accordingly, not divided into standard segments (Table 34). The distribution of OUs along this ONV (in its original scaling) was unimodal and symmetric.

This CLG expresses variation in human impact as reflected in man-made infrastructure, expressed by the abundances of buildings, roads and other visible traces of human activity. Low PD values show that there is little residual variation attributable to infrastructure left after extraction of CLGs 1–4. The weak foundation for OI as an independent CLG in coastal plains motivated for not dividing this CLG into standard segments.

##### Tentative division of coastal plains into minor types

Correlations coefficients between ONVs and ordination axes, primary key variables and analysis variables were calculated as a basis for operationalisation of the CLGs (Table 35). The theoretical number of gradient combinations (tentative minor types) is 5 × 3 × 2 × 2 (× 1) = 60. Of these, 51 were represented in the total data set (Table 36); 31 of which with > 4 OUs (i.e. > 1% of the KS457 data subset). Several figures (e.g. Figs 109 and 112) indicate that the variation within IYK·5 (to some extent also IYK·4) along the other CLGs is too small to allow further division. If this holds true, some minor-type candidates should be merged regardless of the number of OUs assigned to them.

**Table 35.** Correlations between the five ONVs and ordination axes (rows 1–4), primary key variables (rows 6–13), and analytic variables in major landscape type coastal plains (KS).

| Variable |  | IYK | RE | VP | SkP | OI |
| --- | --- | --- | --- | --- | --- | --- |
| GNMDS4_1 |  | −0.6416 | −0.0317 | 0.0349 | 0.2579 | 0.2136 |
| GNMDS4_2 |  | −0.0924 | 0.6587 | 0.0168 | −0.0673 | −0.0769 |
| GNMDS4_3 |  | −0.0076 | −0.0361 | 0.3764 | −0.1063 | 0.0104 |
| GNMDS4_4 |  | 0.0087 | −0.1271 | −0.3181 | −0.0447 | −0.0196 |
| RR1 | Relief | −0.1548 | 0.6536 | −0.0097 | −0.0482 | 0.0159 |
| TPI1 | Peaks | 0.0867 | 0.1654 | 0.0400 | −0.0258 | 0.0268 |
| AlpA | Boreal/alpine landscape | −0.0083 | 0.0801 | 0.0199 | −0.0454 | −0.0279 |
| II | Abundance of freshwater lakes | −0.3320 | 0.0274 | 0.4148 | 0.0687 | 0.0382 |
| MI | Abundance of mire | −0.3851 | −0.0295 | 0.3473 | 0.0085 | −0.0690 |
| Ifl | Amount of infrastructure (OI) | −0.4246 | −0.0743 | −0.0226 | 0.2989 | 0.4557 |
| Iflu |  | −0.5562 | −0.0445 | 0.0537 | 0.2859 | 0.3304 |
| JI | Agricultural land-use intensity | −0.4721 | −0.1223 | −0.0832 | 0.2196 | 0.2282 |
| Gab_nae | Ant commercial buildings | −0.4072 | −0.0787 | 0.0197 | 0.2859 | 0.3556 |
| Gab_fi | Ant fisheries-related buildings | −0.2141 | −0.0335 | 0.1301 | 0.0991 | 0.3197 |
| Gab_a | AA built-up | −0.4944 | −0.0223 | 0.0297 | 0.2978 | 0.4034 |
| Flat_a | AA flat terrain | 0.1395 | −0.5609 | −0.0075 | −0.0167 | −0.0203 |
| Vmag_a | AA regulated magazine | −0.0537 | −0.0012 | −0.0215 | 0.0209 | −0.0006 |
| Vei_b_a | AA road | −0.4865 | −0.0760 | 0.0255 | 0.1925 | 0.2563 |
| Ktkmo_a | AA thick layer of till | −0.2119 | −0.0317 | −0.0720 | 0.0688 | 0.0710 |
| Tpil1_a | AA depressions | −0.1790 | 0.5801 | 0.0171 | 0.0026 | −0.0020 |
| Tpilh_a | AA convex terrain | −0.0511 | 0.5878 | 0.0361 | 0.0044 | 0.0233 |
| Tpil_mp | Terrain form TPI1, numerical, mean | 0.0845 | 0.1573 | 0.0431 | −0.0229 | 0.0314 |
| Stroms_a | AA strong ocean current | 0.0346 | −0.0476 | 0.0191 | −0.0680 | 0.0952 |
| Stromn_a | AA normal ocean current | 0.4913 | −0.0634 | −0.0049 | −0.1577 | −0.0635 |
| Stroml_a | AA weak ocean current | −0.1220 | 0.0876 | −0.0779 | 0.0883 | −0.0283 |
| River_a | AA large river | −0.1784 | −0.1681 | 0.0489 | −0.0137 | 0.0325 |
| Sti_a | AA trail, path | −0.0493 | −0.0302 | −0.0465 | 0.1058 | 0.0885 |
| Steep_a | AA steep terrain/slope | −0.1386 | 0.6922 | 0.0086 | −0.0303 | 0.0103 |
| Kskred_a | AA landslide soil | 0.0176 | 0.1359 | 0.0011 | 0.0389 | −0.0310 |
| Setr_s | Ant summer mountain pasture | −0.1332 | 0.0073 | 0.0137 | 0.1063 | 0.0431 |
| Sefr_a | AA old buildings | −0.3666 | −0.0199 | 0.0452 | 0.2638 | 0.4048 |
| Rug3_m | Terrain ruggedness VRM3, mean | −0.0936 | 0.8604 | 0.0582 | −0.0024 | 0.0180 |
| Rr1_m | Altitudinal range | −0.1548 | 0.6536 | −0.0097 | −0.0481 | 0.0158 |
| Brich_a | AA lime-rich bedrock | −0.0125 | 0.0725 | −0.0686 | 0.0170 | −0.0937 |
| Bpoor_a | AA lime-poor bedrock geology | −0.0214 | 0.0583 | 0.0453 | 0.1217 | 0.0804 |
| Rd_anl_a | AA reindeer husbandry facilities | −0.1274 | 0.0477 | −0.0196 | −0.0068 | −0.0815 |
| Bplu_a | AA plutonic rock | −0.0270 | −0.0191 | 0.0717 | 0.0556 | 0.0936 |
| Bvul_a | AA volcanic rock | −0.1186 | −0.0079 | 0.0777 | 0.0879 | 0.0271 |
| Bavstn_a | AA sedimentary rock | −0.1403 | −0.1051 | −0.0525 | −0.0057 | 0.0764 |
| Bomd_a | AA metamorphic rock | 0.0045 | 0.1014 | 0.0395 | −0.0126 | −0.0708 |
| Land_a | AA terrestrial area | −0.5479 | −0.0163 | 0.1023 | 0.2151 | 0.0728 |
| Build_a | AA built-up area | −0.2685 | −0.0861 | 0.0170 | 0.2376 | 0.3011 |
| City_a | AA town/city area | −0.1061 | −0.0438 | −0.0353 | 0.0554 | 0.0942 |
| Mire_a | AA mire | −0.4023 | −0.0098 | 0.3666 | 0.0384 | −0.0297 |
| Meant_a | AA moderate slope | −0.1357 | 0.3625 | 0.0151 | 0.0523 | 0.0085 |
| Maro_s | Ant marine islands | 0.1956 | 0.0298 | 0.2434 | 0.0799 | 0.0203 |
| Kmar_a | AA marine deposits | −0.3165 | −0.1839 | −0.0782 | 0.1497 | 0.1333 |
| Lled_a | AA power lines | −0.5100 | −0.0101 | 0.0689 | 0.2673 | 0.2308 |
| Klac_a | AA lacustrine deposits | −0.0361 | −0.0681 | −0.0670 | −0.0260 | −0.0045 |
| Kelv_a | AA glaciofluvial deposits | −0.1852 | −0.0246 | −0.0899 | 0.0906 | −0.0351 |
| Kbf_a | AA exposed bedrock | 0.4453 | 0.1769 | −0.0108 | −0.0929 | −0.1538 |
| Lake_a | AA freshwater lake | −0.3187 | 0.0146 | 0.4959 | 0.0794 | −0.0277 |
| Innoy_s | Ant freshwater lake islands | −0.2394 | 0.0438 | 0.2960 | 0.0956 | 0.0060 |
| Er_m | Hypsographic index, ER, mean | 0.1600 | 0.0304 | 0.0957 | −0.0941 | 0.0381 |
| R_net_a | AA river | −0.6889 | 0.0308 | 0.1020 | 0.0673 | 0.0654 |
| Ekspve_a | AA exposed coast | 0.5410 | -0.0032 | -0.0158 | -0.0142 | -0.0407 |
| Ekspmo_a | AA slightly exposed coast | 0.3526 | -0.0174 | 0.1029 | -0.1196 | -0.0275 |
| Ekspbe_a | AA slightly protected coast | -0.0806 | 0.0067 | -0.1111 | -0.0937 | 0.0411 |
| Inns_s | number of lakes | -0.1552 | -0.0120 | 0.6803 | 0.0261 | 0.0058 |
| Cul_u_s | number of cultural heritage sites outdoors | -0.1598 | -0.0663 | -0.0095 | 0.1499 | 0.1010 |
| Cul_t_s | number of technical heritage sites | -0.1365 | 0.0029 | 0.0166 | 0.1020 | 0.1695 |
| Cul_m_s | number of marine cultural heritage sites | -0.0918 | 0.1424 | 0.0657 | 0.1869 | 0.1991 |
| Cul_k_s | number of church ruins | -0.1703 | -0.0618 | 0.0244 | 0.2272 | 0.2494 |
| Cul_b_s | number of ancient rock art sites | -0.1568 | -0.0886 | -0.0892 | 0.1057 | 0.0613 |
| Cul_a_s | number of archaeological heritage sites | -0.3371 | -0.0326 | 0.0452 | 0.2715 | 0.2476 |
| Cre_b_a | AA steep coast | -0.0892 | 0.4157 | 0.0223 | -0.0400 | 0.0120 |
| Cre_f_a | AA flat coast | 0.1241 | -0.4471 | 0.0928 | 0.0029 | 0.0251 |
| Crug3_m | coastal ruggedness, VRM3, mean | -0.0356 | 0.6560 | -0.0373 | -0.0184 | -0.0080 |
| Crug9_m | coastal ruggedness, coarse scale, VRM9, mean | -0.0801 | 0.6597 | -0.0103 | -0.0218 | 0.0056 |
| Cr3_u_a | AA rugged coast | -0.0305 | 0.5620 | 0.0527 | 0.0055 | 0.0706 |
| Cr3_r_a | AA smooth/flat coast | -0.0670 | -0.0733 | 0.3221 | 0.1746 | 0.1433 |
| Ccom_m | coastal complexity | -0.0739 | 0.1156 | 0.3379 | 0.1551 | 0.1382 |
| C_kk_a | AA complex coastline | 0.0146 | 0.0188 | 0.3501 | 0.1467 | 0.0895 |
| C_ek_a | AA simple coastline Gab_fi | -0.0961 | 0.0898 | -0.1386 | -0.0549 | 0.0313 |
| Bohei_a | AA boreal heaths | -0.3307 | 0.0122 | 0.3352 | 0.0122 | 0.1222 |
| Asp_s_a | AA south facing terrain | 0.0202 | -0.0339 | 0.0204 | 0.0452 | -0.0053 |
| Asp_n_a | AA north facing terrain | 0.1413 | 0.0253 | 0.0750 | -0.0631 | -0.0469 |
| Abygg_a | AA large buildings | -0.2705 | -0.0884 | 0.0174 | 0.2353 | 0.3044 |
| Araaf_a | AA open areas | -0.2311 | -0.0163 | 0.3107 | 0.1529 | 0.1101 |
| Arbar_a | AA coniferous forest | -0.4631 | 0.0568 | -0.0104 | 0.3532 | 0.1385 |
| Arbla_a | AA mixed boreal forest | -0.4375 | 0.0438 | -0.0604 | 0.5176 | 0.0264 |
| Arfull_a | AA arable land | -0.4576 | -0.1258 | -0.0685 | 0.2421 | 0.2395 |
| Arlov_a | AA deciduous forest | -0.4881 | 0.0457 | 0.0357 | 0.4430 | 0.0073 |
| Arover_a | AA surface cultivated land | -0.2500 | -0.0255 | 0.1675 | 0.1242 | 0.2240 |
| Dismire | distance to mire, mean | 0.4457 | -0.0582 | -0.3491 | -0.0238 | 0.0011 |
| Dislake | distance to lake, mean | 0.3470 | -0.0601 | -0.4977 | -0.0218 | -0.0024 |
| Discoast | distance to coast, mean | 0.3766 | -0.0114 | 0.1459 | -0.0915 | -0.0437 |
| Guro_t_a | AA rugged terrain | -0.0874 | 0.5134 | 0.0650 | -0.0236 | 0.0034 |
| Tpi6l_a | AA rugged terrain, TPI6 | -0.1951 | 0.0610 | -0.0675 | 0.0782 | -0.0361 |
| Sn_imp | AA impediment | 0.1797 | 0.1324 | 0.0557 | -0.0969 | -0.0840 |
| Sn_flekk | AA patchy open treeless area | 0.1029 | 0.1623 | 0.3152 | -0.1636 | -0.0667 |
| Sn_lav | AA lichen heath | -0.0606 | -0.0099 | -0.0437 | -0.0343 | 0.0794 |
| Sn_torr | AA dry heath/open areas | -0.0717 | 0.0689 | 0.4291 | -0.1118 | -0.0297 |
| Sn_frisk | AA moist/fresh heath/open areas | -0.0869 | 0.0128 | 0.2905 | -0.1276 | 0.1004 |
| Sn_ureg | AA unregistered heath/open areas | -0.2128 | -0.1371 | -0.0478 | 0.2381 | 0.1860 |
| Oyst_i | Inverse island size | 0.5539 | -0.0164 | -0.0853 | -0.0521 | -0.0697 |

**Table 36.** Realised gradient combinations (unique combinations of segments, i.e. tentative minor landscape types) and number of observation units for each combination in coastal plains (KS).

| Tentative<br>minor<br>type<br>number | Unique<br>combinations<br>of segments | No.<br>of<br>OUs |
| --- | --- | --- |
| 1 | 1111 | 4 |
| 2 | 1112 | 14 |
| 3 | 1121 | 2 |
| 4 | 1122 | 1 |
| 5 | 1211 | 32 |
| 6 | 1212 | 38 |
| 7 | 1221 | 18 |
| 8 | 1222 | 18 |
| 9 | 1311 | 6 |
| 10 | 1312 | 2 |
| 11 | 1321 | 4 |
| 12 | 1322 | 3 |
| 13 | 2111 | 13 |
| 14 | 2112 | 5 |
| 15 | 2121 | 10 |
| 16 | 2122 | 6 |
| 17 | 2211 | 20 |
| 18 | 2212 | 28 |
| 19 | 2221 | 11 |
| 20 | 2222 | 20 |
| 21 | 2311 | 7 |
| 22 | 2312 | 6 |
| 23 | 2321 | 3 |
| 24 | 2322 | 3 |
| 25 | 3111 | 11 |
| 26 | 3112 | 1 |
| 27 | 3121 | 5 |
| 28 | 3211 | 24 |
| 29 | 3212 | 21 |
| 30 | 3221 | 8 |
| 31 | 3222 | 10 |
| 32 | 3311 | 11 |
| 33 | 3312 | 1 |
| 34 | 3321 | 3 |
| 35 | 3322 | 6 |
| 36 | 4111 | 2 |
| 37 | 4112 | 2 |
| 38 | 4121 | 1 |
| 39 | 4122 | 5 |
| 40 | 4211 | 9 |
| 41 | 4212 | 8 |
| 42 | 4221 | 10 |
| 43 | 4311 | 5 |
| 44 | 4321 | 1 |
| 45 | 4322 | 1 |
| 46 | 5111 | 1 |
| 47 | 5112 | 8 |
| 48 | 5122 | 1 |
| 49 | 5211 | 16 |
| 50 | 5212 | 8 |
| 51 | 5311 | 4 |

#### Coastal fjords (KF)

The analyses for identification of independent significant analytic variables for each CLG candidate by forward selection from the *n* = 84 variables using RDA show that seven of the eight CLG candidates satisfied the requirement for explaining at least 6% of the variation (Table 37). The parsimonious set of CLGs, found by RDA with forward selection among the CLG candidates separately for each functional variable category, contained 4 CLGs (Table 37). The properties of the four CLGs and their associated (ranked primary) ONVs are described below on the basis of Tables 37–43 and Figs 114–122.

**Fig. 114.**
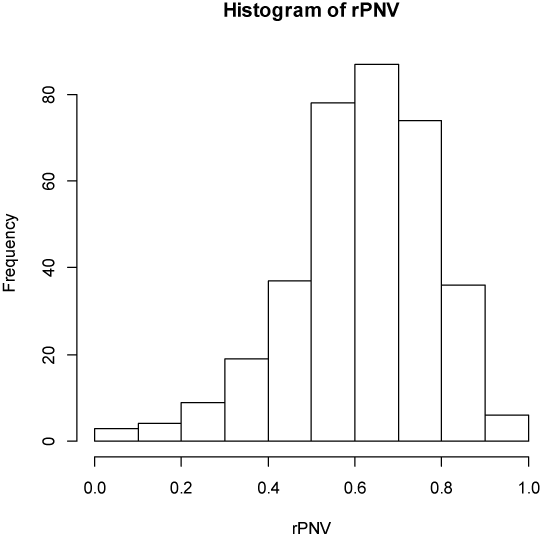
Major type coastal fjords (KF): candidate complex-landscape gradient (CLG) 1, relief (RE): distribution of OUs on the ranged but not rescaled orthogonal key variable (ONV).

**Fig. 115.**
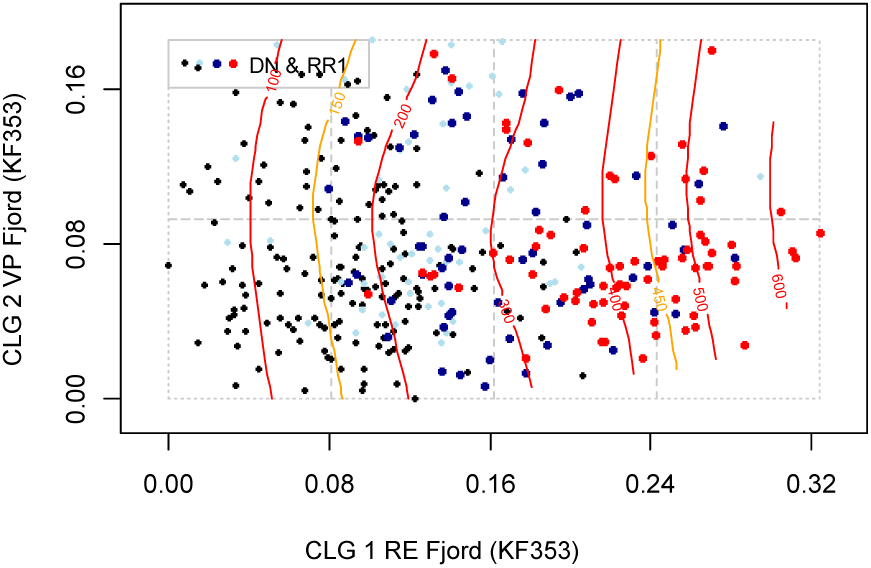
Major type coastal fjords (KF): distribution of 353 observation units (KF353 data subset) along rescaled orthogonal key variables (rONVs) for complex landscape gradients CLG 1 (relief; RE) and CLG 2 (abundance of wetlands; VP). Symbols show affiliation of OUs to levels of relevant landscape-gradient (LG) variables, operationalised as ordered factor variables in accordance with Table 3. Isolines are given for relevant primary key variables (Table 4) and analytic variables (Table 5).

**Fig. 116.**
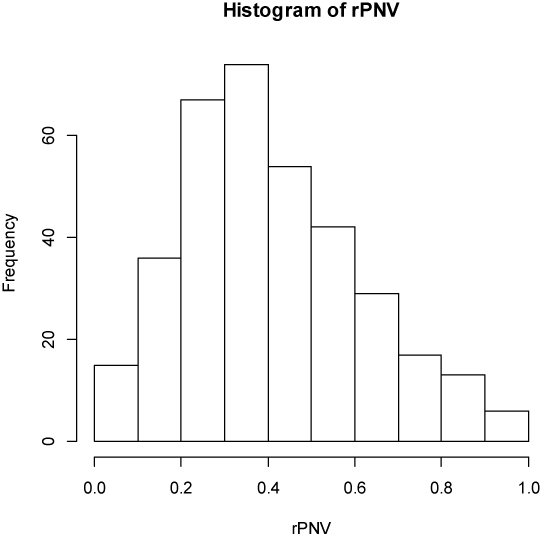
Major type coastal fjords (KF): candidate complex-landscape gradient (CLG) 2, abundance of lakes and wetlands (VP): distribution of OUs on the ranged but not rescaled orthogonal key variable (ONV).

**Fig. 117–118.**
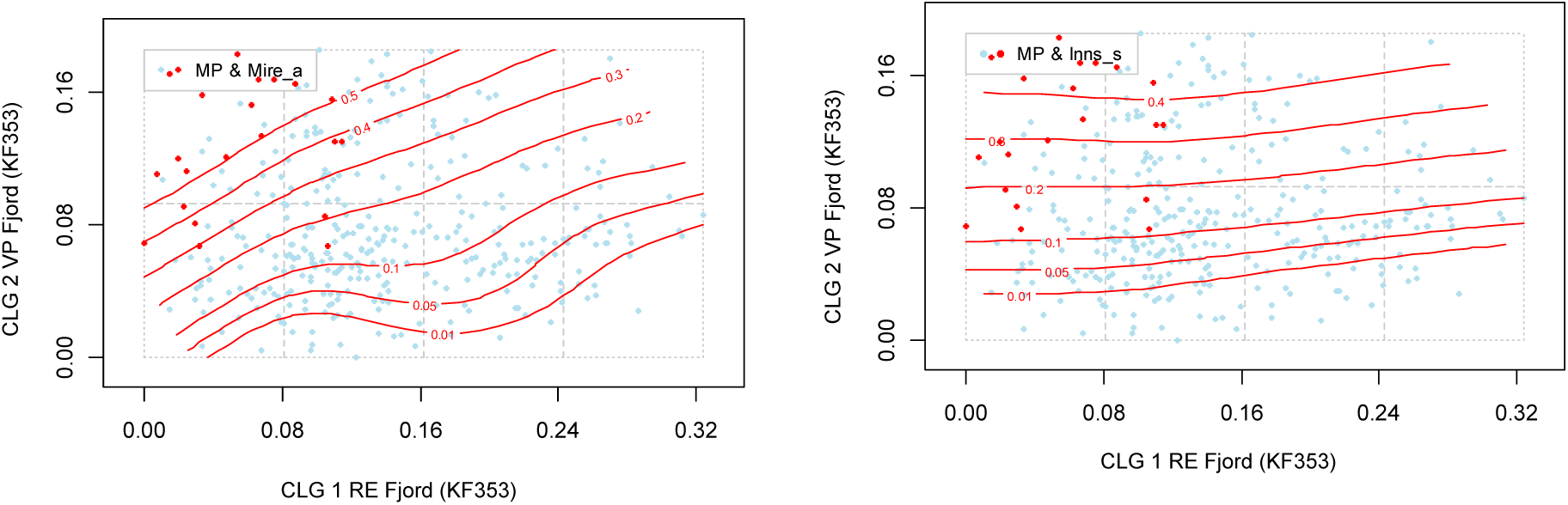
Major type coastal fjords (KF): distribution of 353 observation units (KF353 data subset) along rescaled orthogonal key variables (rONVs) for complex landscape gradients CLG 1 (relief; RE) and CLG 2 (abundance of wetlands; VP). Symbols show affiliation of OUs to levels of relevant landscape-gradient (LG) variables, operationalised as ordered factor variables in accordance with Table 3. Isolines are given for relevant primary key variables (Table 4) and analytic variables (Table 5).

**Fig. 119.**
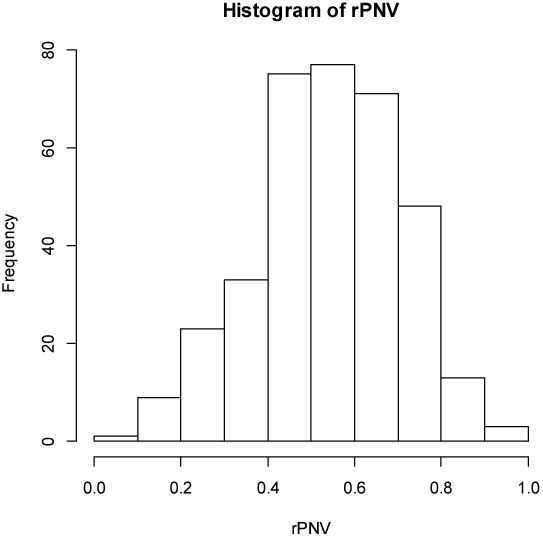
Major type coastal fjords (KF): candidate complex-landscape gradient (CLG) 3, open area cover (AP): distribution of OUs on the ranged but not rescaled orthogonal key variable (ONV).

**Fig. 120.**
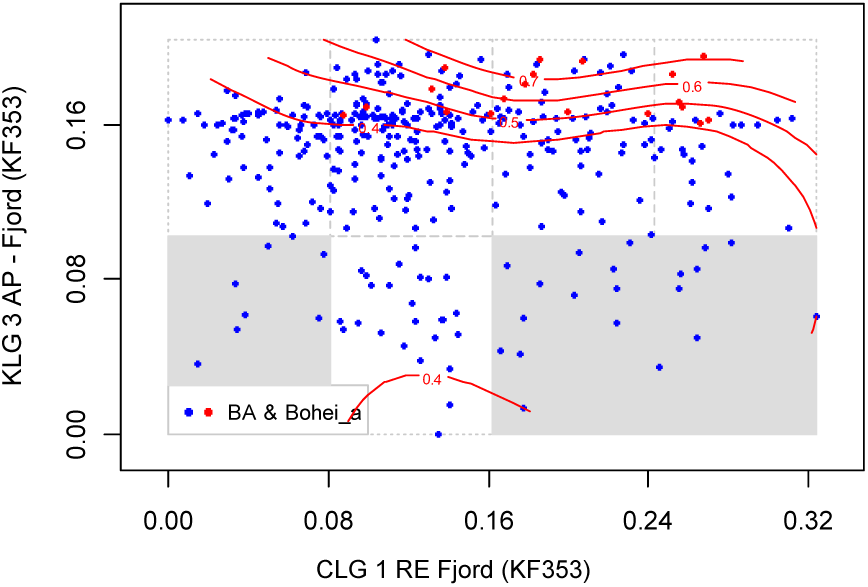
Major type coastal fjords (KF): distribution of 353 observation units (KF353 data subset) along rescaled orthogonal key variables (rONVs) for complex landscape gradients CLG 1 (relief; RE) and CLG 3 (open area cover; AP). Symbols show affiliation of OUs to levels of relevant landscape-gradient (LG) variables, operationalised as ordered factor variables in accordance with Table 3. Isolines are given for relevant primary key variables (Table 4) and analytic variables (Table 5).

**Fig. 121.**
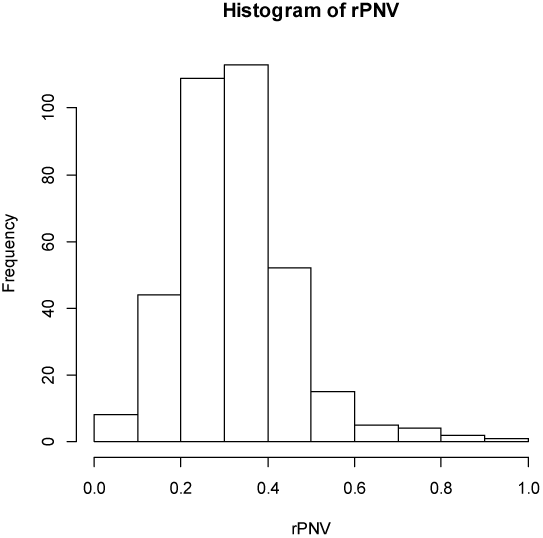
Major type coastal fjords (KF): candidate complex-landscape gradient (CLG) 4, amount of infrastructure (OI): distribution of OUs on the ranged but not rescaled orthogonal key variable (ONV).

**Fig. 122.**
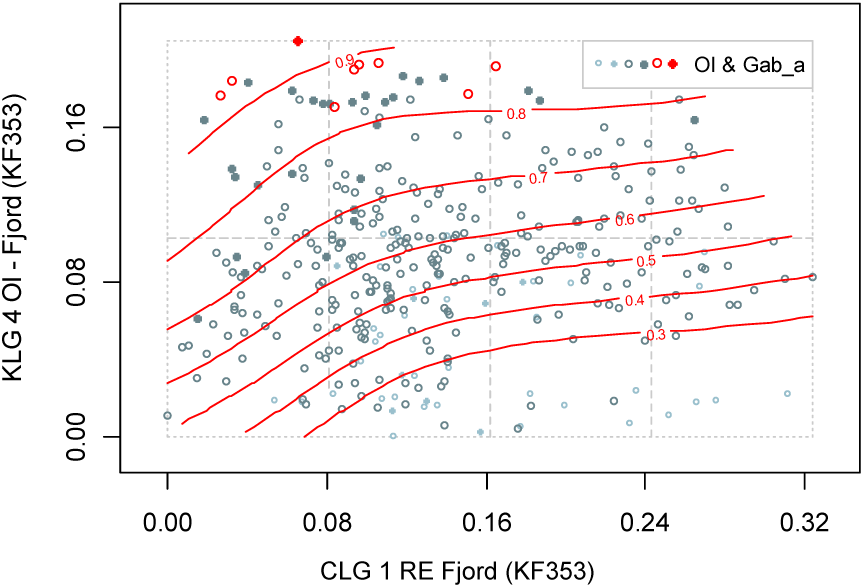
Major type coastal fjords (KF): distribution of 353 observation units (KF353 data subset) along rescaled orthogonal key variables (rONVs) for complex landscape gradients CLG 1 (relief; RE) and CLG 4 (amount of infrastructure; OI). Symbols show affiliation of OUs to levels of relevant landscape-gradient (LG) variables, operationalised as ordered factor variables in accordance with Table 3. Isolines are given for relevant primary key variables (Table 4) and analytic variables (Table 5).

**Table 37.** Identification of CLG candidates in coastal fjords by forward selection of variables using RDA. Results of tests of single CLG candidates are shown in bold-face types. The best model at each step in the selection process is highlighted using light grey shading while the best model after testing all CLG candidates in a functional variable category is highlighted using darker grey shading. Var = variable; Cst Var = constraining variable; Cond Var = conditioning variable(s); df = degrees of freedom; VE = variation explained (expressed as fraction of the total variation); ΔVE = additional variation explained; pseudo-F and P = F-statistic used in the test of the null hypothesis that adding the variable(s) do not contribute more to explaining variation in landscape-element composition than a random variable, with the associated P value. Variable names are abbreviated according to Tables 4 and 5. For further explanation, se Material and methods chapter.

| Model number | | CLG/Cst Var (Code/VarName) | Variables / Cond Var | df | ΔVE | pseudo-F | P |
| --- | --- | --- | --- | --- | --- | --- | --- |
| G1 | RE | Relief | ***Crug9_m + Kskred_a + Rug3_m + RRI*** | 4 | **0.21519** |  |  |
| G2 | VP | Abundance of wetlands | ***Mire_a + Inns_s*** | 2 | **0.10266** |  |  |
| G3 | KX | Coastal exposure | ***R_net_a + Stroml_a*** | 2 | **0.09495** |  |  |
| G1G2 | VP | Abundance of wetlands | RE | 2 | 0.07178 | 17.417 | 0.01 |
| G1G3 | KX | Coastal exposure | RE | 2 | 0.05906 | 14.078 | 0.01 |
| G1G2 |  |  | VE = 0.28657 |  |  |  |  |
| GU1 | AP | Open area cover | ***Bohei_a + Kbf_a + Araaf_a + Sn_imp*** | 4 | **0.08279** | 11.232 | 0.01 |
|  |  |  | RE + VP |  | 0.11604 |  |  |
| GU2 | AP | Forest cover | ***Arbar_a + Arlov_a*** | 2 | **0.05875** | 15.444 | 0.01 |
|  |  |  | RE + VP |  | 0.96370 |  |  |
| GU1U2 | A | Open area cover | RE + VP + AP | 2 | 0.03337 | 9.505 | 0.01 |
|  |  |  |  |  | 0.05287 |  |  |
| G+U1 |  |  | VE = 0.36977 |  |  |  |  |
| GUA1 | OI | Amount of infrastructure | ***Iflu+Build_a+Gab_a*** | 3 | **0.07504** | 15.271 | 0.01 |
|  |  |  | RE + VP + AP |  | 0.11888 |  | + |
| GUA2 | JP | Agricultural land-use intensity | ***Arfull_a*** | 1 | **0.02984** | 16.945 | – |
|  |  |  | RE + VP + AP |  | 0.04728 |  |  |
| G+U+A1 |  |  | VE = 0.44481 |  |  |  |  |

**Table 38.**
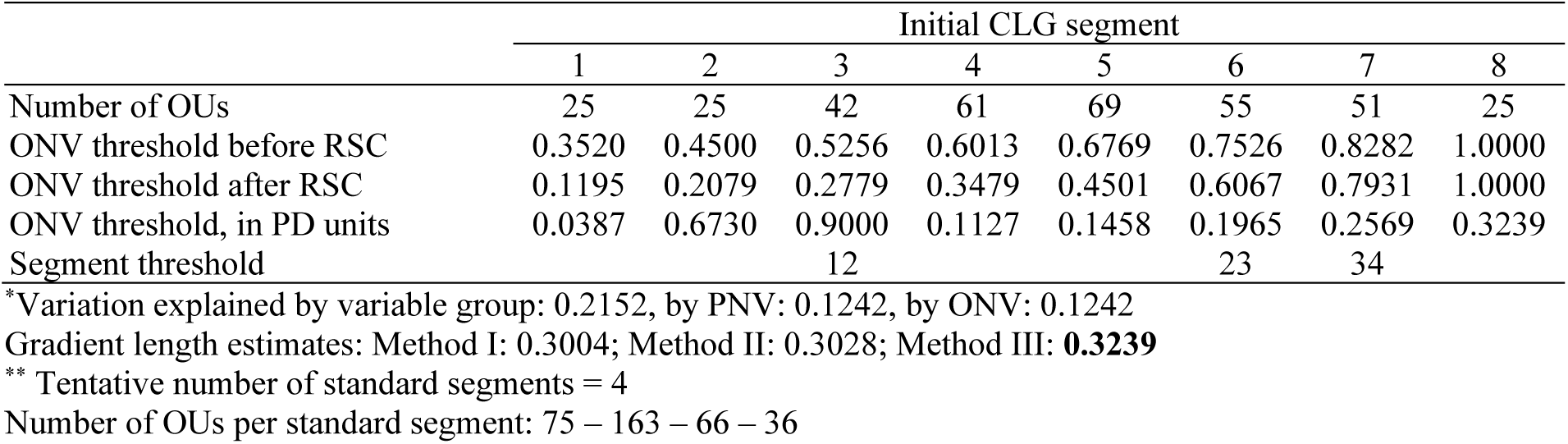
Major type coastal fjords (KF): key properties* and segmentation** of candidate complex-landscape gradient G1, relief (RE), as represented by its orthogonal key variable (ONV). OU = observation unit; Threshold = upper limit of class (RSC = rescaling). PD unit = Proportional dissimilarity unit. Segment threshold shows position of standard segment borders relative to initial segments.

|  | Initial CLG segment |  |  |  |  |  |  |  |
| --- | --- | --- | --- | --- | --- | --- | --- | --- |
|  | 1 | 2 | 3 | 4 | 5 | 6 | 7 | 8 |
| Number of OUs | 25 | 25 | 42 | 61 | 69 | 55 | 51 | 25 |
| ONV threshold before RSC | 0.3520 | 0.4500 | 0.5256 | 0.6013 | 0.6769 | 0.7526 | 0.8282 | 1.0000 |
| ONV threshold after RSC | 0.1195 | 0.2079 | 0.2779 | 0.3479 | 0.4501 | 0.6067 | 0.7931 | 1.0000 |
| ONV threshold, in PD units | 0.0387 | 0.6730 | 0.9000 | 0.1127 | 0.1458 | 0.1965 | 0.2569 | 0.3239 |
| Segment threshold |  |  | 12 |  |  | 23 | 34 |  |

\*Variation explained by variable group: 0.2152, by PNV: 0.1242, by ONV: 0.1242
Gradient length estimates: Method I: 0.3004; Method II: 0.3028; Method III: **0.3239**
\*\* Tentative number of standard segments = 4
Number of OUs per standard segment: 75 – 163 – 66 – 36

**Table 39.**
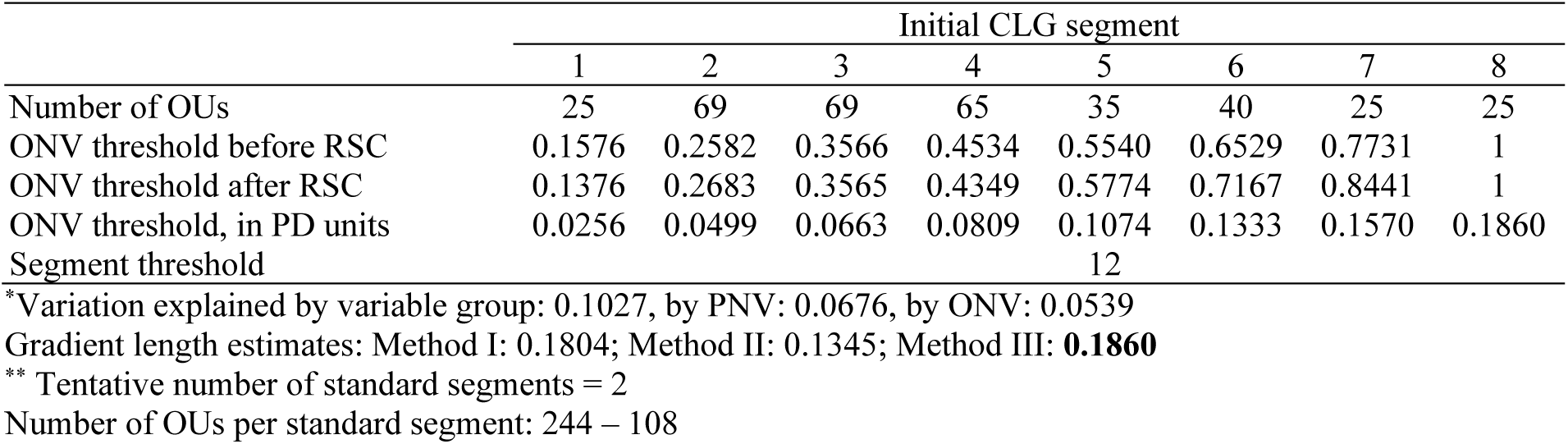
Major type coastal fjords (KF): key properties* and segmentation** of candidate complex-landscape gradient G2, abundance of lakes and wetlands (VP), as represented by its orthogonal key variable (ONV). OU = observation unit; Threshold = upper limit of class (RSC = rescaling). PD unit = Proportional dissimilarity unit. Segment threshold shows position of standard segment borders relative to initial segments.

**Table 40.**
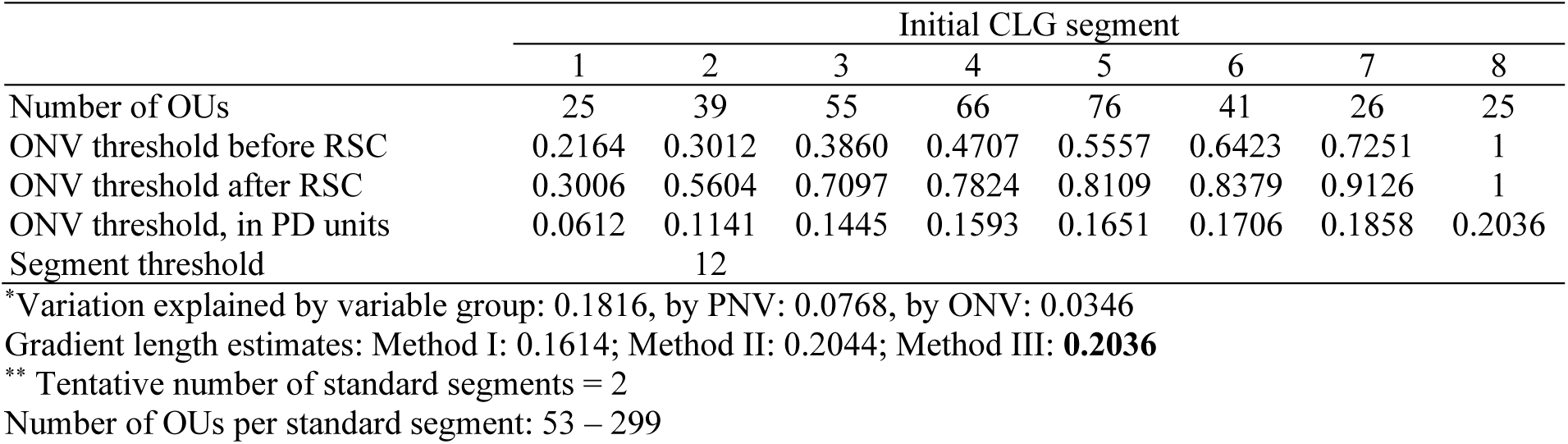
Major type coastal fjords (KF): key properties* and segmentation** of candidate complex-landscape gradient U1, open area cover (AP), as represented by its orthogonal key variable (ONV). OU = observation unit; Threshold = upper limit of class (RSC = rescaling). PD unit = Proportional dissimilarity unit. Segment threshold shows position of standard segment borders relative to initial segments.

**Table 41.**
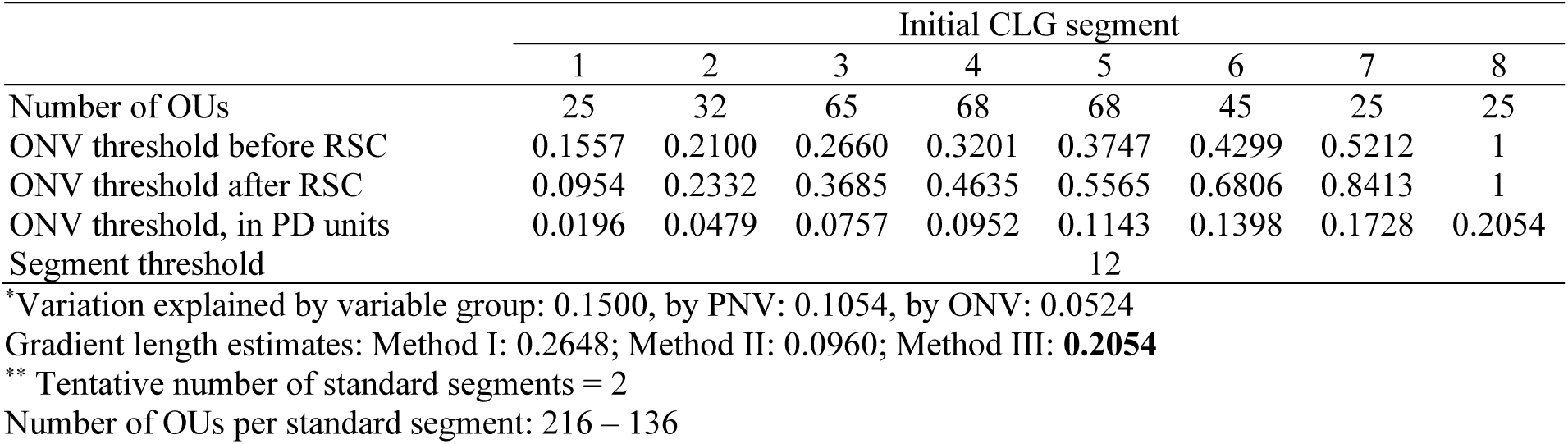
Major type coastal fjords (KF): key properties* and segmentation** of candidate complex-landscape gradient A1, amount of infrastructure (OI), as represented by its orthogonal key variable (ONV). OU = observation unit. Threshold = upper limit of class (RSC = rescaling). PD unit = Proportional dissimilarity unit. Segment threshold shows position of standard segment borders relative to initial segments.

**Table 42.** Correlations between the four ONVs and ordination axes (rows 1–4), primary key variables (rows 6–15), and analysis variables in major landscape type coastal fjords (KF).

| Variable |  | RE | VP | AP | OI |
| --- | --- | --- | --- | --- | --- |
| GNMDS 1 |  | –0.3684 | –0.1461 | –0.2633 | 0.3109 |
| GNMDS 2 |  | 0.2248 | 0.3029 | 0.0297 | 0.1741 |
| GNMDS 3 |  | –0.4474 | 0.2770 | 0.2249 | –0.1571 |
| DNI | Valley shape | 0.6755 | 0.0485 | 0.0173 | 0.0421 |
| RR1 | Relief | 0.6984 | 0.0241 | 0.0016 | 0.0430 |
| TPI1 | Peaks | –0.5798 | –0.0576 | –0.0347 | 0.0026 |
| Br1 | Glacier | 0.0598 | 0.0057 | 0.0980 | –0.0603 |
| AlpA | Boreal/alpine landscape | 0.1569 | 0.2450 | 0.3728 | –0.0394 |
| II | Abundance of freshwater lakes | –0.0051 | 0.3218 | 0.0637 | 0.0545 |
| MI | Abundance of mire | –0.2441 | 0.4311 | 0.0312 | –0.0114 |
| If1 | Amount of infrastructure | –0.2717 | –0.2419 | –0.1921 | 0.3642 |
| If1u | Amount of infrastructure | –0.1816 | –0.0186 | –0.2101 | 0.4616 |
| JI | Agricultural land-use intensity | –0.2160 | –0.0892 | –0.1879 | 0.3355 |
| Gab_nae | Ant commercial buildings | –0.1750 | –0.1159 | –0.1875 | 0.4437 |
| Gab_fi | Ant fisheries-related buildings | –0.0626 | –0.0135 | –0.0307 | 0.2818 |
| Gab_a | AA built-up | –0.2049 | –0.0981 | –0.1937 | 0.5038 |
| Flat_a | AA flat terrain | –0.4696 | 0.0015 | 0.0438 | –0.0144 |
| Vei_b_a | AA regulated magazine | –0.1523 | –0.0593 | –0.2043 | 0.3866 |
| Ktkmo_a | AA road | –0.0354 | 0.0612 | –0.0151 | 0.0972 |
| Tpill_a | AA thick layer of till | 0.6097 | 0.0420 | 0.0406 | –0.0044 |
| Tpilh_a | AA depressions | 0.3569 | –0.0668 | –0.0771 | 0.0478 |
| Tpil_mp | AA convex terrain | –0.5680 | –0.0552 | –0.0327 | 0.0036 |
| Stroms_a | Terrain form TPI1, numerical, mean | –0.0084 | 0.0803 | –0.0360 | –0.0235 |
| Stromn_a | AA normal ocean current | –0.0802 | –0.0872 | 0.0813 | –0.0725 |
| Stroml_a | AA weak ocean current | –0.0275 | –0.1855 | –0.0144 | –0.0792 |
| River_a | AA large river | –0.1235 | 0.0774 | –0.1035 | 0.1957 |
| Sti_a | AA trail, path | 0.0190 | –0.0437 | –0.0027 | 0.0797 |
| Steep_a | AA steep terrain/slope | 0.6530 | –0.0349 | –0.0374 | 0.0260 |
| Kskred_a | AA landslide soil | 0.3027 | 0.0106 | 0.0057 | 0.0283 |
| Setr_s | Ant summer mountain pasture | –0.0409 | 0.0149 | –0.1206 | 0.1777 |
| Sefr_a | AA old buildings | –0.0658 | –0.1383 | –0.2264 | 0.4192 |
| Rug3_m | Terrain ruggedness VRM3, mean | 0.7625 | –0.0007 | –0.0121 | 0.0211 |
| Rr1_m | Altitudinal range | 0.6984 | 0.0241 | 0.0016 | 0.0430 |
| Brich_a | AA lime-rich bedrock | –0.1779 | –0.0258 | –0.0536 | 0.0753 |
| Bpoor_a | AA lime-poor bedrock geology | 0.1830 | –0.0101 | –0.1239 | 0.0874 |
| Rd_anl_a | AA reindeer husbandry facilities | 0.0039 | 0.1447 | 0.1162 | –0.0238 |
| Bplu_a | AA plutonic rock | 0.1134 | 0.0710 | 0.0692 | –0.0303 |
| Bvul_a | AA volcanic rock | 0.0322 | –0.0453 | –0.0217 | 0.0538 |
| Bavstn_a | AA sedimentary rock | –0.2143 | –0.0673 | –0.0304 | 0.0855 |
| Bomd_a | AA metamorphic rock | –0.0434 | –0.0537 | –0.0620 | 0.0405 |
| Land_a | AA terrestrial area | 0.1016 | 0.2953 | 0.0037 | 0.1712 |
| Build_a | AA built-up area | –0.1631 | –0.1603 | –0.1181 | 0.4572 |
| City_a | AA town/city area | –0.0875 | –0.0284 | –0.0129 | 0.1451 |
| Mire_a | AA mire | –0.2298 | 0.4917 | 0.0487 | 0.0320 |
| Glac_a | AA glacier | 0.0690 | 0.0157 | 0.0879 | –0.0391 |
| Meant_a | AA moderate slope | –0.4560 | 0.0382 | 0.0339 | 0.0107 |
| Maro_s | Ant marine islands | 0.0152 | –0.0957 | 0.0093 | 0.0081 |
| Kmar_a | AA marine deposits | –0.4089 | –0.0183 | –0.0453 | 0.0258 |
| Lled_a | AA power lines | –0.1055 | 0.0253 | –0.1762 | 0.3309 |
| Klac_a | AA lacustrine deposits | 0.0406 | 0.0236 | –0.0165 | 0.0578 |
| Kelv_a | AA glaciofluvial deposits | –0.0141 | –0.0080 | 0.0170 | –0.0168 |
| Kbf_a | AA exposed bedrock | 0.1183 | –0.0765 | 0.2251 | –0.0816 |
| Lake_a | AA freshwater lake | –0.0883 | 0.4881 | –0.0184 | –0.0367 |
| Innoy_s | Ant freshwater lake islands | –0.0130 | 0.1910 | 0.0052 | 0.0394 |
| Er_m | Hypsographic index, ER, mean | –0.0510 | –0.1225 | –0.0960 | 0.0132 |
| R_net_a | AA river | 0.0870 | 0.3018 | –0.0493 | 0.1654 |
| Ekspve_a | AA exposed coast | –0.0076 | 0.0236 | 0.1214 | –0.0774 |
| Ekspmo_a | AA slightly exposed coast | -0.0056 | -0.0144 | 0.1670 | -0.1055 |
| Ekspe_a | AA slightly protected coast | -0.0971 | -0.1861 | -0.0066 | -0.1221 |
| Inns_s | Ant lakes | -0.1022 | 0.6960 | 0.0159 | -0.0120 |
| Cul_u_s | Ant cultural heritage sites outdoors | -0.0862 | -0.0476 | 0.0467 | 0.1111 |
| Cul_t_s | Ant technical heritage sites | 0.0098 | 0.0157 | -0.1022 | 0.2343 |
| Cul_m_s | Ant marine cultural heritage sites | 0.0905 | -0.1409 | -0.1097 | 0.1705 |
| Cul_k_s | Ant church ruins | -0.1058 | -0.0380 | -0.0655 | 0.3963 |
| Cul_b_s | Ant ancient rock art sites | -0.0423 | -0.0389 | -0.0438 | 0.0499 |
| Cul_a_s | Ant archaeological heritage sites | -0.2222 | -0.0372 | -0.0580 | 0.2495 |
| Cre_b_a | AA steep coast | 0.5958 | 0.0024 | 0.0125 | 0.0313 |
| Cre_f_a | AA flat coast | -0.2641 | 0.0647 | 0.0371 | -0.0394 |
| Crug3_m | coastal ruggedness, VRM3, mean | 0.6677 | -0.0412 | -0.0344 | 0.0366 |
| Crug9_m | coastal ruggedness, coarse scale, VRM9, mean | 0.8576 | 0.0245 | -0.0207 | 0.0332 |
| Cr3_u_a | AA rugged coast | 0.5610 | 0.0219 | -0.0438 | 0.0870 |
| Cr3_r_a | AA smooth/flat coast | -0.3207 | 0.1408 | 0.0135 | 0.1090 |
| Ccom_m | coastal complexity | 0.1033 | 0.0871 | 0.0044 | 0.1011 |
| C_kk_a | AA complex coastline | -0.0296 | 0.0845 | -0.0496 | 0.0727 |
| C_ek_a | AA simple coastline Gab_fi | 0.2011 | 0.0183 | 0.0870 | 0.0717 |
| Bohei_a | AA boreal heaths | 0.0430 | 0.4261 | 0.3190 | 0.0231 |
| Asp_s_a | AA south facing terrain | 0.0045 | 0.0097 | -0.0401 | 0.0265 |
| Asp_n_a | AA north facing terrain | 0.0182 | 0.0118 | 0.0634 | 0.0145 |
| Abygg_a | AA large buildings | -0.1690 | -0.1556 | -0.1225 | 0.4503 |
| Araaf_a | AA open areas | -0.0255 | 0.0936 | -0.0379 | 0.0385 |
| Arbar_a | AA coniferous forest | 0.0314 | -0.2555 | -0.3195 | 0.2152 |
| Arbla_a | AA mixed boreal forest | 0.0754 | -0.2078 | -0.2864 | 0.1769 |
| Arfull_a | AA arable land | -0.1823 | -0.0927 | -0.2167 | 0.3242 |
| Arlov_a | AA deciduous forest | -0.0112 | 0.0403 | -0.2476 | 0.1175 |
| Arover_a | AA surface cultivated land | -0.0639 | 0.0009 | -0.0844 | 0.2060 |
| Dismire | distance to mire, mean | 0.1710 | -0.2334 | 0.0880 | -0.0027 |
| Dislake | distance to lake, mean | -0.0584 | -0.3611 | -0.0703 | -0.0106 |
| Discoast | distance to coast, mean | -0.0256 | 0.2223 | 0.0717 | -0.1207 |
| Guro_t_a | AA rugged terrain | 0.7817 | 0.0631 | 0.0078 | 0.0280 |
| Tpi6l_a | AA rugged terrain, TPI6 | -0.1027 | 0.0206 | 0.0423 | -0.0768 |
| Sn_imp | AA impediment | 0.1750 | 0.2834 | 0.5276 | 0.0418 |
| Sn_flekk | AA patchy open treeless area | 0.1581 | 0.2859 | 0.2861 | -0.0882 |
| Sn_lav | AA lichen heath | 0.0565 | -0.0101 | -0.0540 | -0.0191 |
| Sn_torr | AA dry heath/open areas | 0.1051 | 0.3030 | 0.2348 | -0.0792 |
| Sn_frisk | AA moist/fresh heath/open areas | 0.0206 | 0.3753 | 0.2924 | -0.0462 |
| Sn_ureg | AA unregistered heath/open areas | -0.1610 | 0.0068 | -0.0290 | 0.2252 |
| Oyst_i | Inverse island size | -0.0535 | 0.1327 | 0.0793 | -0.1530 |

**Table 43.** Realised gradient combinations (unique combinations of segments, i.e. tentative minor landscape types) and number of observation units for each combination in coastal fjords (KF).

| Tentative<br>minor<br>type<br>number | Unique<br>combinations<br>of segments | No. of OUs |
| --- | --- | --- |
| 1 | 1111 | 2 |
| 2 | 1112 | 1 |
| 3 | 1121 | 27 |
| 4 | 1122 | 14 |
| 5 | 1211 | 3 |
| 6 | 1212 | 1 |
| 7 | 1221 | 13 |
| 8 | 1222 | 8 |
| 9 | 2111 | 12 |
| 10 | 2112 | 4 |
| 11 | 2121 | 61 |
| 12 | 2122 | 43 |
| 13 | 2211 | 8 |
| 14 | 2212 | 1 |
| 15 | 2221 | 25 |
| 16 | 2222 | 17 |
| 17 | 3111 | 7 |
| 18 | 3112 | 2 |
| 19 | 3121 | 24 |
| 20 | 3122 | 24 |
| 21 | 3211 | 1 |
| 22 | 3212 | 2 |
| 23 | 3221 | 15 |
| 24 | 3222 | 4 |
| 25 | 4111 | 4 |
| 26 | 4112 | 3 |
| 27 | 4121 | 13 |
| 28 | 4122 | 5 |
| 29 | 4212 | 1 |
| 30 | 4221 | 6 |
| 31 | 4222 | 2 |

##### CLG 1 – (G1) Relief in fjords: RE (Table 38, Figs 114–115)

The four variables included in the ONV that represents CLG 1, relief (RE), were: (1) mean coastal ruggedness, coarse scale (VRM9); (2) areal coverage of landslide soil; (3) mean terrain ruggedness (Rug3_m); and (4) relief (RR1). This geo-ecological CLG had an estimated gradient length of 0.3239/0.08 = 4.049 EDU–L units and was, accordingly, divided into 4 standard segments (Table 38). The distribution of OUs along this ONV (in its original scaling) was left-skewed (Fig. 114).

This CLG expresses variation in terrain form within coastal fjord landscapes as expressed by the depth/width ratio of the valley/fjord relative to its surroundings; from wide fjord via open fjord to narrow, deep and steep-sided fjord. The CLG had good linearity, with an even distribution of OUs along the CLG, but with diminishing degree of generalization towards the gradient’s upper end (high relief; Fig. 115). Isolines for RR1 values of 150 m, 300 m and 450 m (Fig. 115) corresponded closely to the transitions between RE standard segments 1–2, 2–3 and 3–4, respectively.

##### CLG 2 – (G2) Abundance of wetlands: VP (Table 39, Figs 116–118)

The two variables included in the ONV that represents CLG 2, abundance of wetlands (VP) were: (1) areal coverage of mire (Mire_a) and (2) number of lakes (Inns_s). This geo-ecological CLG had an estimated gradient length of 0.1860/0.08 = 2.325 EDU–L units and was, accordingly, divided into 2 standard segments (Table 39). The distribution of OUs along this ONV (in its original scaling) was right-skewed (Fig. 116).

This CLG expresses variation in the fraction of an OU’s area that was covered by wetland systems (predominantly mires) and in the abundance of small lakes and tarns which are often associated with wetlands: from low to medium abundance to high abundance of mires and tarns (Figs 117–118). Variation along this CLG was found for OUs with low relative relief (Figs 117–118). Isolines for number of lakes (Inns_s) ran almost exactly parallel with the CLG (Fig. 118). The CLG VP in fjords also captured variation in vegetation cover; this CLG was more strongly correlated with the fractional cover of boreal heaths (Table 42; Bohei_a; τ = 0.4261) than was the orthogonalised gradient AP which expresses open area cover (Table 42; AP and Bohei_a: τ = 0.3190). The variation along this CLG is most clearly expressed by number of lakes (Inns_s).

##### CLG 3 – (U1) Open area cover: AP (Table 40, Figs 119–120)

The four variables included in the ONV that represents CLG 3, open area cover (AP), were the fractions of OU area covered by: (1) boreal heath (Bohei_a); (2) exposed bedrock (Kbf_a); (3) open areas (Araaf_a); and (4) ‘impediment’ (i.e. areas not suitable for agriculture or forestry; Sn_imp), respectively. This bio-ecological CLG had an estimated gradient length of 0.2036/0.08 = 2.545 EDU–L units and was, accordingly, divided into 2 standard segments (Table 40). The distribution of OUs along this ONV (in its original scaling) was slightly left-skewed (Fig. 119).

This CLG (Fig. 120) expresses variation in the fraction of OUs covered by open land, from areas with low open area cover (i.e. forests) to predominately open areas without or with sparse forest cover. The linearity of this CLG was generally low (Table 40). CLG AP separated a relatively small group of low-score OUs from the majority of OUs which were placed in segment 2. Only three variables had correlations coefficients |τ| > 0.3 with CLG AP (Table 42): areal coverage of impediment (Sn_imp; τ = 0.5276), areal coverage of boreal heaths (Bohei_a; τ = 0.3190) and relative abundance of boreal vs. alpine landscapes (AlpA; τ = 0.3728). The CLG AP explained variation in vegetation cover not accounted for by CLG 1, RE. Based on the weak correlation between primary key variable AlpA and this CLG, we regard the foundation for AP as an independent CLG in fjords as relatively weak.

##### CLG 4 – (A1) Amount of infrastructure: OI (Table 41, Figs 121–122)

The two variables included in the ONV that represents CLG 4, amount of infrastructure (OI), were: (1) the amount of infrastructure (IfIu) and (2) areal coverage of built-up areas (Build_a and Gab_a). This land-use related CLG had an estimated gradient length of 0.2054/0.08 = 2.567 EDU–L units and was, accordingly, divided into 2 standard segments (Table 40). The distribution of OUs along this ONV (in its original scaling) was right skewed (Fig. 121).

This CLG expresses variation in the abundance of buildings and other infrastructure from low and intermediate to settlement (village, small town or city; Fig. 122). The CLG has good linearity with decreasing degree of generalisation towards both gradient ends (Table 41, Fig. 122). The gradient explains residual variation related to infrastructure not accounted for by the first axes. Variation was found primarily in wide and open fjords. The CLG is correlated with variables expressing variation in agricultural land-use intensity (e.g. the key variable agricultural land-use intensity, JI; τ = 0.3355) and has a very low (negative) correlation with the ONV that expresses the open-area cover (AP; CLG 3). Eventual removal of CLG AP from accepted CLGs will therefore have minimal effect on CLG amount of infrastructure. Amount of infrastructure (OI) was only weakly (negatively) correlated with CLG 1 Relief (e.g. IfIu has τ = –0.1816 with CLG 1 RE). This substantiates amount of infrastructure as a strong orthogonalised CLG.

##### Tentative division of coastal fjords into minor types

Correlations between ONVs and ordination axes, primary key variables, and analysis variables were calculated as a basis for operationalisation of the CLGs (Table 42). The theoretical number of gradient combinations (tentative minor types) is 4 × 2 × 2 × 2 = 32. Of these, 31 were represented in the total data set (Table 43), 20 of which with > 3 OUs (i.e. > 1% of the KF353 data subset). The most commonly encountered combination in our data set was 2121: i.e. open fjords (RE·2) with low abundance of wetlands (VP·1), high open area cover (AP·2) and low amount of infrastructure (OI·1).

#### Inland fine-sediment plains (IS)

The analyses for identification of independent significant analytic variables for each CLG candidate by forward selection from the *n* = 84 variables using RDA (Table 44) show that five out of nine CLG candidates satisfied the relaxed criterion for accepting CLG candidates set to 8% of the variation (as opposed to 6% for the other major types). Use of a relaxed criterion was motivated by the small data set (118 OUs). For the RDA analysis with forward selection of variables within each functional variable category, undertaken to find a parsimonious set of CLGs, the CLG candidate soil type (JA) was subjectively chosen as CLG 1. The reason for this was (1) the sharp separation of the two dominant soil type classes in the ordination diagram, and (2) that the variation explained by relief (RE) was inflated by the large number of intercorrelated topographic variables in the data set. The parsimonious set of CLGs contained 4 CLGs, while abundance of mire (MP) was very close to fulfil the requirement. The properties of the four CLGs and their associated (ranked primary) ONVs are described below on the basis of Tables 45–48 and Figs 123–128.

**Fig. 123.**
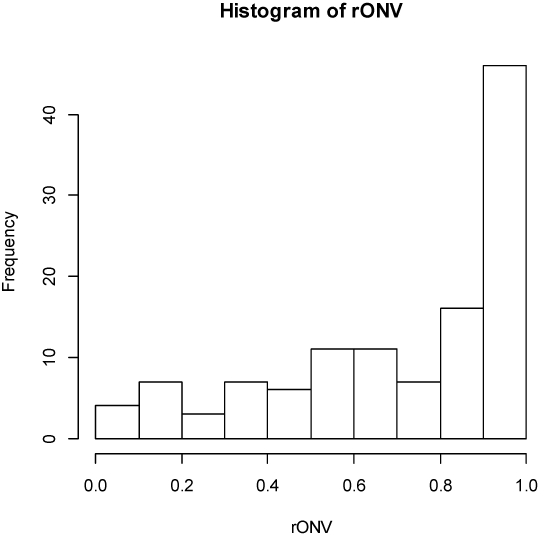
Inland fine-sediment plains (IF): candidate complex-landscape gradient (CLG) 1, soil type (JA): distribution of OUs on the ranged but not rescaled orthogonal key variable (ONV).

**Fig. 124.**
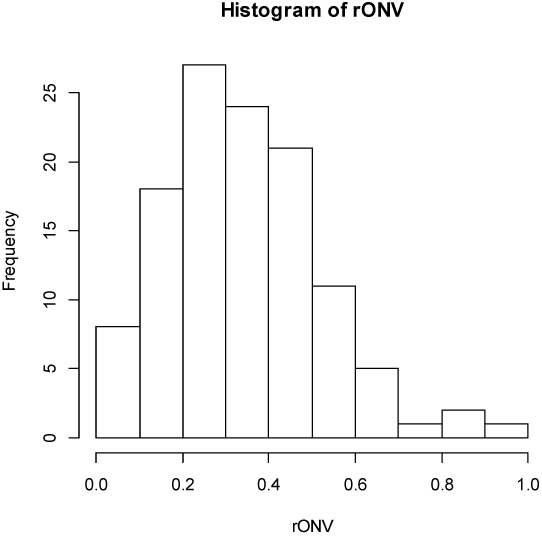
Inland fine-sediment plains (IF): candidate complex-landscape gradient (CLG) 2, relief (RE): distribution of OUs on the ranged but not rescaled orthogonal key variable (ONV).

**Fig. 125.**
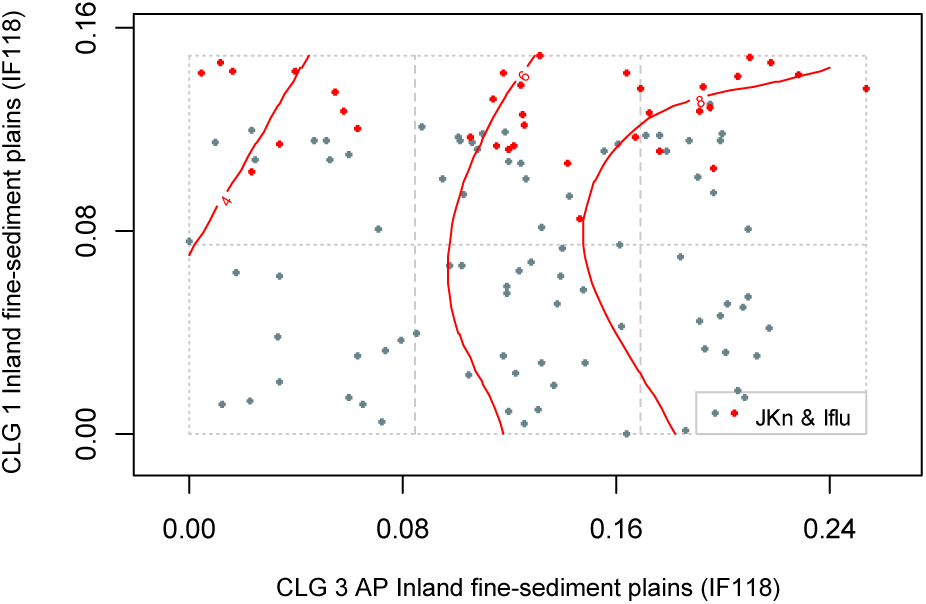
Major type inland fine-sediment plains (IF): distribution of 118 observation units (IF118 data subset) along rescaled orthogonal key variables (rONVs) for complex landscape gradients CLG 3 (open area cover; AP) and CLG 4 (amount of infrastructure; OI). Symbols show affiliation of OUs to levels of relevant landscape-gradient (LG) variables, operationalised as ordered factor variables in accordance with Table 3. Isolines are given for relevant primary key variables (Table 4) and analytic variables (Table 5).

**Fig. 126.**
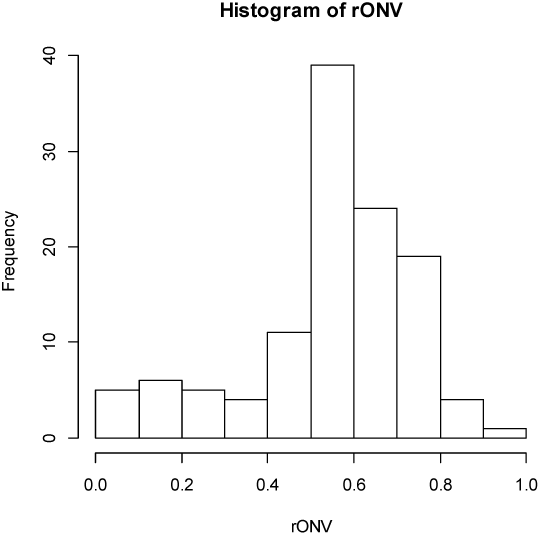
Inland fine-sediment plains (IF): candidate complex-landscape gradient (CLG) 3, open area cover (AP): distribution of OUs on the ranged but not rescaled orthogonal key variable (ONV).

**Fig. 127.**
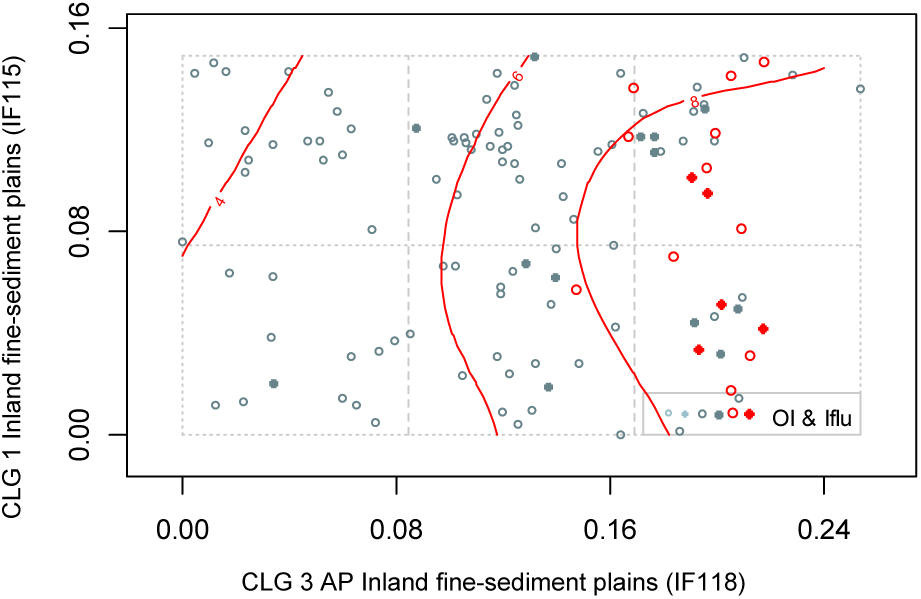
Major type inland fine-sediment plains (IF): distribution of 118 observation units (IF118 data subset) along rescaled orthogonal key variables (rONVs) for complex landscape gradients CLG 3 (open area cover; AP) and CLG 4 (amount of infrastructure; OI). Symbols show affiliation of OUs to levels of relevant landscape-gradient (LG) variables, operationalised as ordered factor variables in accordance with Table 3. Isolines are given for relevant primary key variables (Table 4) and analytic variables (Table 5).

**Fig. 128.**
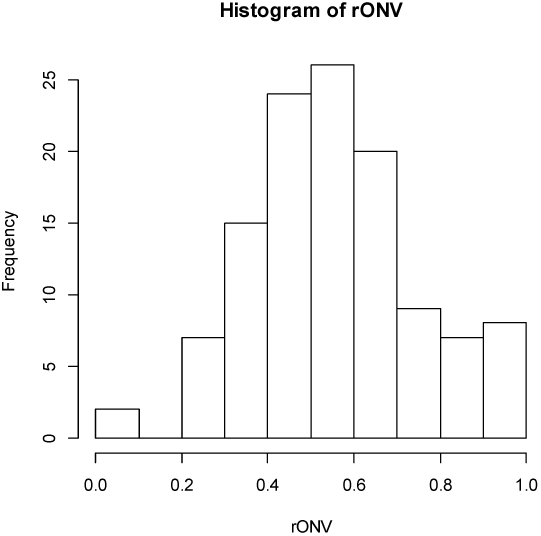
Inland fine sediment plains (IF): candidate complex-landscape gradient (CLG) 4, amount of infrastructure (OI): distribution of OUs on the ranged but not rescaled orthogonal key variable (ONV).

**Table 44.** Identification of CLG candidates in inland fine-sediment plains (IF) by forward selection of variables using RDA. Results of tests of single CLG candidates are shown in bold-face types. The best model at each step in the selection process is highlighted using light grey shading while the best model after testing all CLG candidates in a functional variable category is highlighted using darker grey shading. Abbreviations and explanations: Var = variable; Cst Var = constraining variable; Cond Var = conditioning variable(s); df = degrees of freedom; VE = variation explained (expressed as fraction of the total variation); ΔVE = additional variation explained; pseudo-F and P = F-statistic used in the test of the null hypothesis that adding the variable(s) do not contribute more to explaining variation in landscape-element composition than a random variable, with the associated P value. Variable names are abbreviated according to Tables 4 and 5. For further explanation, se Material and methods chapter. Note that the CLG candidate G1 was selected *a priori*.

| Model number | CLG/Cst Var (Code/VarName) | Variables / Cond Var | df | ΔVE | pseudo-F | P |
| --- | --- | --- | --- | --- | --- | --- |
| G4 | RE Relief | ***RR1 + Tpi1h_a*** | 2 | **0.15520** |  |  |
| G1 | JA Soil type | ***Kbf_a + Kelv_a*** | 2 | **0.10388** |  |  |
| G3 | VP Abundance of wetlands | Dismire |  | **0.08734** |  |  |
| G4G1 | RE Relief | ***RR1 + Tpi1h_a***<br>JA | 2 | 0.13210 | 9.767 | 0.01 |
| G3G1 | MP Abundance of mire | ***Dismire***<br>JA | 1 | 0.08425 | 11.830 | 0.01 |
| G3G4G1 | MP Abundance of mire | ***Dismire***<br>JA + RE | 1 | 0.07809 | 12.751 | 0.01 |
| G1G3 |  | VE = 0.23598 |  |  |  |  |
| GU1 | AP Open area cover | ***Araaf_a + Sn_ureg***<br>JA + RE | 2 | **0.10891**<br>0.14255 | 9.229 | 0.01 |
| G+U1 |  | VE = 0.34489 |  |  |  |  |
| GUA1 | OI Amount of infrastructure | ***Iflu***<br>JA + SkP + AP | 1 | **0.06053**<br>0.09240 | 22.991 | 0.01<br>+ |
| G+U1+A1 |  | VE = 0.40542 |  |  |  |  |

**Table 45.**
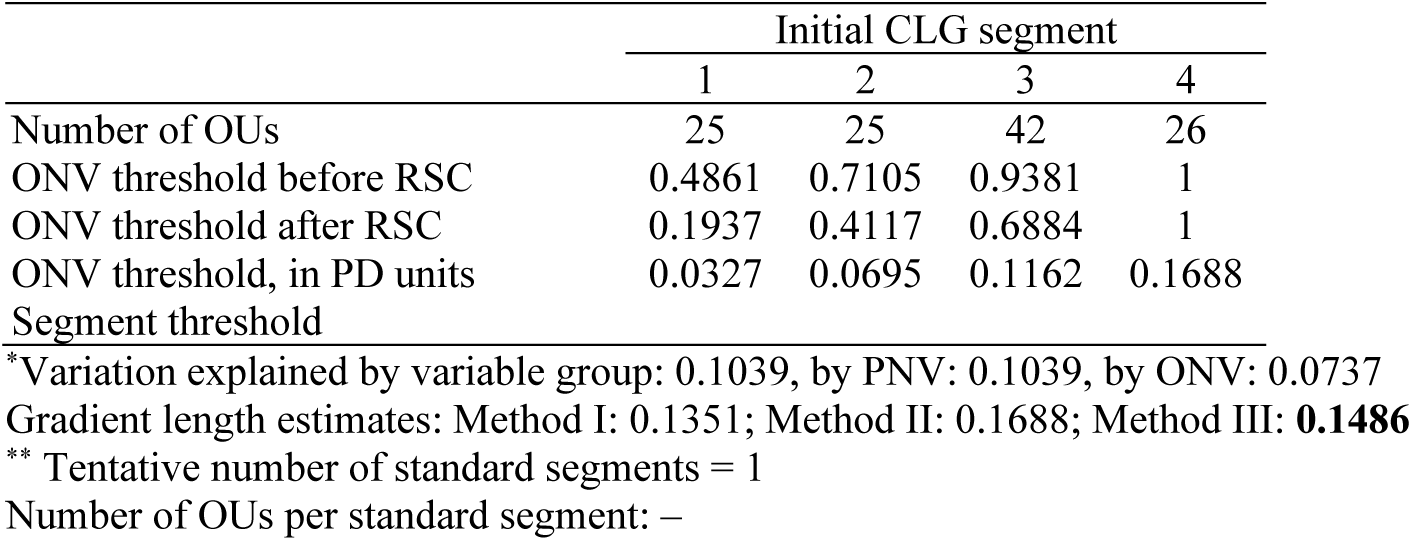
Major type inland fine-sediment plains (IF): key properties* and segmentation** of candidate G1, complex-landscape gradient soil type (JA), as represented by its orthogonal key variable (ONV). OU = observation unit; Threshold = upper limit of class (RSC = rescaling). PD unit = Proportional dissimilarity unit. Segment threshold shows position of standard segment borders relative to initial segments.

|  | Initial CLG segment |  |  |  |
| --- | --- | --- | --- | --- |
|  | 1 | 2 | 3 | 4 |
| Number of OUs | 25 | 25 | 42 | 26 |
| ONV threshold before RSC | 0.4861 | 0.7105 | 0.9381 | 1 |
| ONV threshold after RSC | 0.1937 | 0.4117 | 0.6884 | 1 |
| ONV threshold, in PD units | 0.0327 | 0.0695 | 0.1162 | 0.1688 |
| Segment threshold |  |  |  |  |

\*Variation explained by variable group: 0.1039, by PNV: 0.1039, by ONV: 0.0737
Gradient length estimates: Method I: 0.1351; Method II: 0.1688; Method III: **0.1486**
\*\* Tentative number of standard segments = 1
Number of OUs per standard segment: –

**Table 46.**
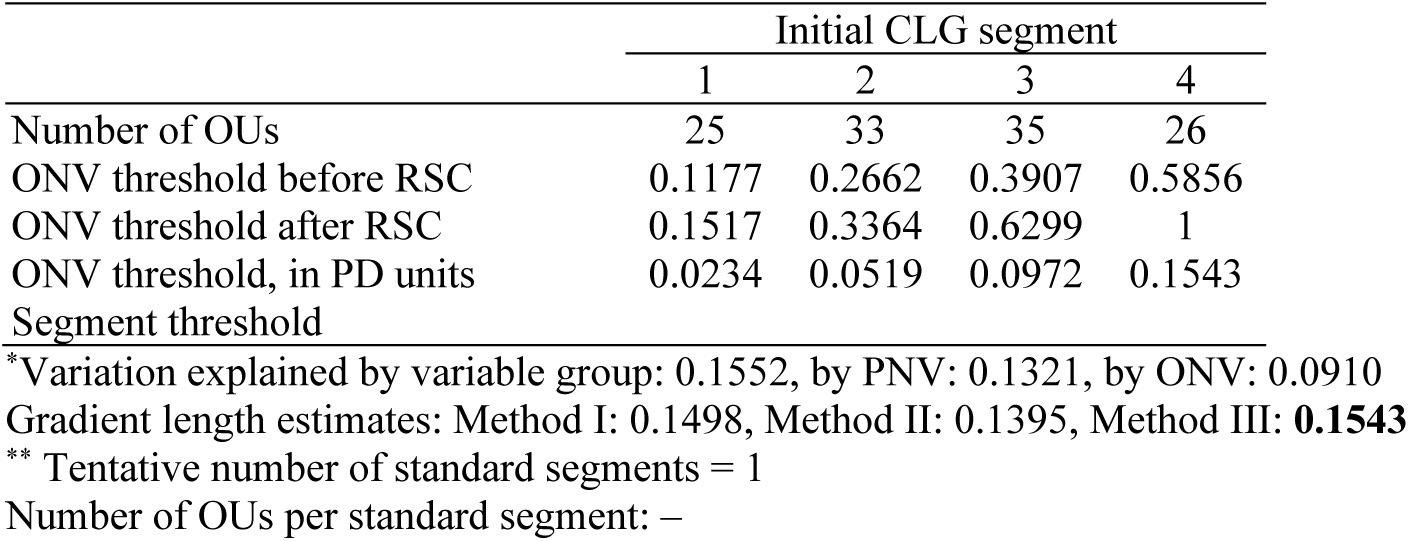
Major type inland fine-sediment plains (IF): key properties* and segmentation** of candidate complex-landscape gradient G2, relief (RE), as represented by its orthogonal key variable (ONV). OU = observation unit; Threshold = upper limit of class (RSC = rescaling). PD unit = Proportional dissimilarity unit. Segment threshold shows position of standard segment borders relative to initial segments.

**Table 47.**
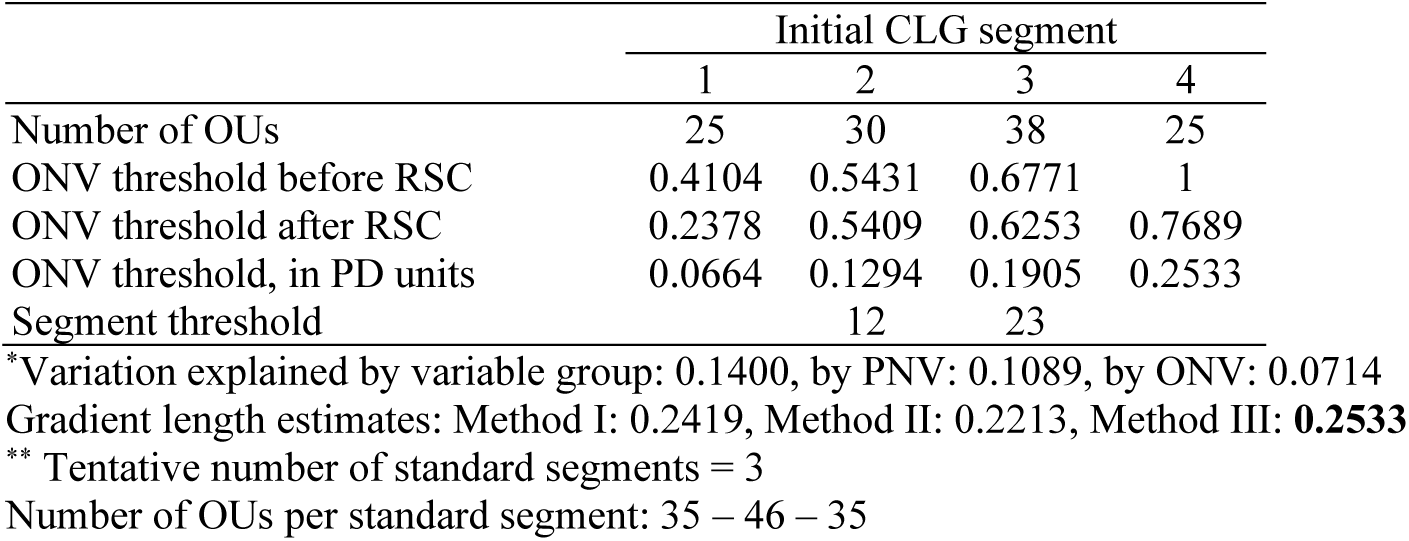
Major type inland fine-sediment plains (IF): key properties* and segmentation** of candidate complex-landscape gradient U1, open areas (open area cover, AP), as represented by its orthogonal key variable (ONV). OU = observation unit; Threshold = upper limit of class (RSC = rescaling). PD unit = Proportional dissimilarity unit. Segment threshold shows position of standard segment borders relative to initial segments.

**Table 48.**
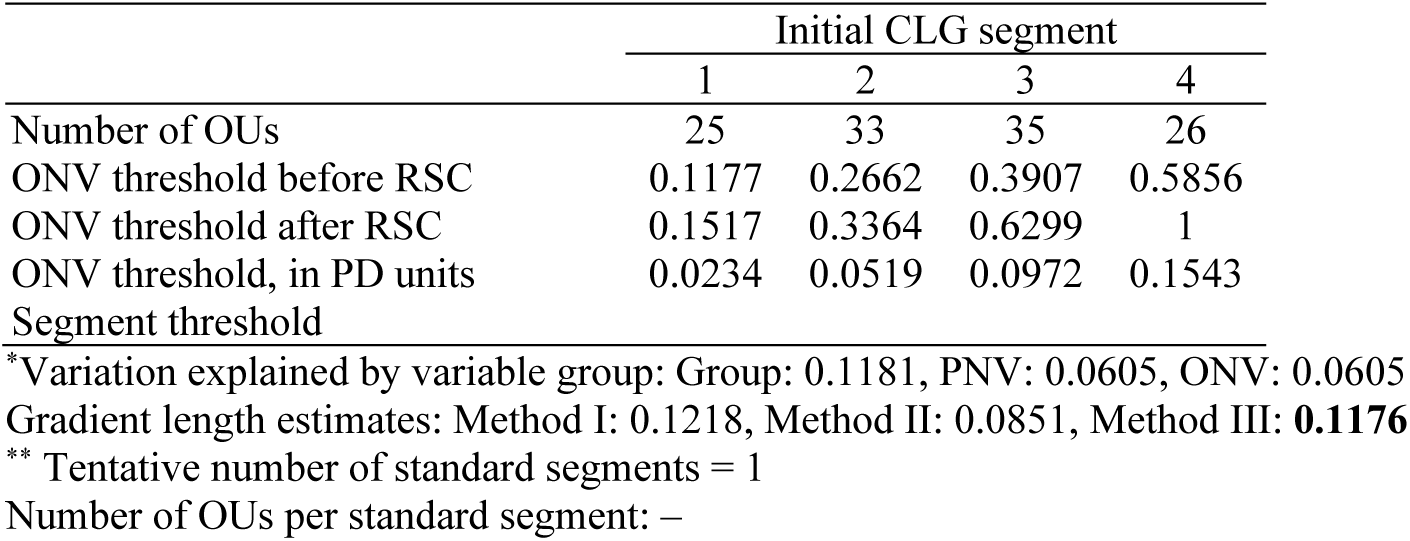
Major type inland fine sediment plains (IF): key properties* and segmentation** of candidate complex-landscape gradient A1, amount of infrastructure (OI), as represented by its orthogonal key variable (ONV). OU = observation unit; Threshold = upper limit of class (RSC = rescaling). PD unit = Proportional dissimilarity unit. Segment threshold shows position of standard segment borders relative to initial segments.

##### CLG 1 – (G1) Soil type: JA (Table 38, Figs 123)

The two variables included in the ONV that represents CLG 1, soil type (JA), were the areal coverage of: (1) exposed bedrock (Kbf_a) and (2) glaciofluvial deposits (Kelv_a). This geo-ecological CLG had an estimated gradient length of 0.1486/0.08 = 1.858 EDU–L units and was, accordingly, not divided into standard segments (Table 45). The distribution of OUs along this ONV (in its original scaling) was left-skewed (Fig. 123).

This CLG expresses variation associated with primary soil category in inland fine-sediment plains, separating fine-sediment plains dominated by marine deposits from plains dominated by glaciofluvial sediments. However, despite the clear separation of the two soil types in the ordination diagram (Fig. 69), the total difference in landscape characteristics between the two classes was not large enough (as estimated by Method III) to motivate for a division of this CLG into two segments.

##### CLG 2 – (G2) Relief in inland fine-sediment plains: RE (Table 46, Figs 124–125)

The two variables included in CLG 2, relief (RE), were: (1) relief (RR1) and (2) areal coverage of convex terrain (Tpi1h_a). This geo-ecological CLG had an estimated gradient length of 0.1543/0.08 = 1.929 EDU–L units and was, accordingly, not divided into standard segments (Table 38). The distribution of OUs along this ONV (in its original scaling) was right-skewed (Fig. 124).

While this ONV expresses variation associated with relief within inland fine-sediment plains, the total variation in landscape characteristics was insufficient (as estimated by Method III) to motivate for a division of this CLG into two segments.

##### CLG 3 – (U1) Open area cover: AP (Table 47, Fig. 127)

The two variables included in the ONV that represents CLG 3, open area cover (AP), were the fractions of OU area covered by: (1) open areas (Araaf_a) and (2) unregistered heath/open areas (Sn_ureg). This bio-ecological CLG had an estimated gradient length of 0.2533/0.08 = 3.166 EDU–L units and was, accordingly, divided into 3 standard segments (Table 47). The distribution of OUs along this ONV (in its original scaling) was slightly left-skewed (Fig. 127).

This CLG expresses variation in vegetation cover and corresponding variation in land-use intensity, displaying variation in open-area cover (AP·1) (Fig. 127). The gradient was most strongly correlated with areal cover of open heath (Sn_frisk; τ = 0.7750) and unregistered heath/open areas (Sn_ureg; τ = 0.5224). GNMDS axis 1 was correlated with several features related to human land-use intensity (Table 49) that are difficult to interpret.

**Table 49.** Correlations between the four ONVs and ordination axes (rows 1–2), primary key variables (rows 3–12), and analytic variables in major landscape type inland fine-sediment plains (IF).

| Variable |  | sONV JA | sONV AP | sONV RE | sONV OI |
| --- | --- | --- | --- | --- | --- |
| GNMDS 1 |  | -0.3175 | 0.3788 | 0.0886 | 0.0381 |
| GNMDS 2 |  | -0.5001 | -0.3533 | 0.1790 | -0.1118 |
| RR1 | Relief | -0.3329 | -0.0018 | 0.8091 | -0.1066 |
| FIA | Amount of flat land | 0.3481 | -0.0038 | -0.5466 | 0.0829 |
| TPI1 | Peaks | -0.1194 | 0.0670 | -0.1162 | -0.0536 |
| AlpA | Boreal/alpine landscape | 0.0857 | -0.0857 | 0.1303 | -0.0969 |
| II | Abundance of freshwater lakes | -0.1535 | 0.1427 | 0.0310 | -0.0655 |
| MI | Abundance of mire | 0.0700 | -0.2661 | -0.0141 | -0.0823 |
| IIf | Amount of infrastructure (OI) | -0.0051 | 0.4670 | 0.0585 | 0.2178 |
| Iflu | Amount of infrastructure | -0.0313 | 0.4819 | 0.0508 | 0.1939 |
| JI | Agricultural land-use intensity | 0.1736 | -0.0346 | -0.1233 | 0.2770 |
| JKn | Soil type | 0.3473 | -0.0028 | -0.5463 | 0.0827 |
| Flat_a | AA flat terrain | 0.0587 | -0.1364 | 0.1151 | 0.0346 |
| Ktkmo_a | AA thick layer of till | -0.2394 | 0.0054 | 0.5869 | -0.0828 |
| Tpi1l_a | AA depressions | -0.4163 | 0.0070 | 0.4525 | -0.1215 |
| Tpi1h_a | AA convex terrain | -0.0942 | 0.0807 | -0.0965 | -0.0548 |
| Tpi1_mp | Terrain form TPI1, numerical, mean | 0.2098 | -0.0712 | 0.0801 | -0.0068 |
| River_a | AA large river | -0.2806 | 0.0051 | 0.6383 | -0.1153 |
| Steep_a | AA steep terrain/slope | -0.0129 | 0.0387 | 0.1517 | -0.0975 |
| Kskred_a | AA landslide soil | -0.3768 | 0.0117 | 0.5691 | -0.1212 |
| Rug3_m | Terrain ruggedness VRM3, mean | -0.3329 | -0.0018 | 0.8091 | -0.1066 |
| Rr1_m | Altitudinal range | -0.1232 | 0.1779 | 0.2652 | -0.1614 |
| Brich_a | AA lime-rich bedrock | -0.0791 | -0.1647 | -0.2655 | 0.1737 |
| Bpoor_a | AA lime-poor bedrock geology | -0.3271 | -0.0059 | -0.0210 | 0.0209 |
| Bplu_a | AA plutonic rock | 0.0336 | 0.0296 | 0.1462 | -0.0382 |
| Bvul_a | AA volcanic rock | 0.0050 | 0.1607 | 0.2885 | -0.1157 |
| Bavstn_a | AA sedimentary rock | 0.0119 | -0.1779 | -0.0245 | 0.1191 |
| Bomd_a | AA metamorphic rock | 0.0134 | -0.2758 | -0.0080 | -0.0347 |
| Mire_a | AA mire | -0.3108 | 0.0294 | 0.5949 | -0.0882 |
| Meant_a | AA moderate slope | -0.1725 | 0.0458 | 0.0394 | 0.0251 |
| Kmar_a | AA marine deposits | -0.0144 | 0.0256 | -0.0440 | -0.0240 |
| Klac_a | AA lacustrine deposits | 0.6984 | 0.0176 | -0.1125 | 0.1346 |
| Kelv_a | AA glaciofluvial deposits | -0.1497 | 0.0427 | 0.0138 | -0.1097 |
| Lake_a | AA freshwater lake | -0.1941 | 0.0165 | -0.0683 | -0.0188 |
| Innoy_s | Ant freshwater lake islands | 0.2384 | 0.0948 | -0.1387 | 0.0434 |
| Er_m | Hypsographic index, ER, mean | 0.0231 | -0.1449 | 0.2196 | 0.0231 |
| R_net_a | AA river | -0.1102 | -0.0221 | 0.0188 | -0.0138 |
| Inns_s | Number of lakes | 0.1713 | 0.0436 | -0.1029 | -0.0107 |
| Asp_s_a | AA south facing terrain | -0.1209 | -0.1251 | 0.0902 | -0.0142 |
| Asp_n_a | AA north facing terrain | 0.0254 | 0.2600 | 0.0178 | 0.0451 |
| Dismire | distance to mire, mean | 0.1440 | -0.0218 | 0.0170 | 0.0881 |
| Dislake | distance to lake, mean | 0.3723 | -0.1254 | -0.1433 | 0.1378 |
| Discoast | distance to coast, mean | -0.0934 | -0.0866 | 0.2624 | -0.1955 |
| Guro_t_a | AA rugged terrain | 0.0650 | -0.1136 | 0.2838 | 0.0540 |
| Tpi6l_a | AA rugged terrain, TPI6 | -0.1355 | -0.1225 | 0.0073 | -0.1400 |
| Oyst_i | Inverse island size | -0.7295 | 0.0363 | 0.1053 | -0.0766 |
| Kbf_a | AA exposed bedrock | -0.0745 | 0.1986 | -0.1255 | 0.0019 |
| Bohei_a | AA boreal heaths | 0.0765 | -0.1103 | -0.0327 | -0.0045 |
| Arbar_a | AA coniferous forest | 0.1634 | 0.1223 | 0.2274 | 0.0471 |
| Arbla_a | AA mixed boreal forest | 0.1350 | 0.2012 | 0.1760 | -0.0354 |
| Arlov_a | AA deciduous forest | 0.0857 | -0.0857 | 0.1303 | -0.0969 |
| Sn_imp | AA impediment | -0.0311 | -0.1274 | 0.0942 | -0.1211 |
| Sn_flekk | AA patchy open treeless area | 0.0584 | -0.1081 | -0.0442 | -0.0166 |
| Sn_lav | AA lichen heath | -0.0295 | -0.1808 | 0.0957 | -0.1153 |
| Sn_torr | AA dry heath/open areas | -0.0779 | -0.1424 | 0.0511 | -0.0838 |
| Sn_frisk | AA moist/fresh heath/open areas | 0.0257 | 0.7750 | -0.0689 | -0.0631 |
| Sn_ureg | AA unregistered heath/open areas | -0.0191 | 0.5224 | 0.0262 | 0.1117 |
| Gab_nae | Ant commercial buildings | -0.0023 | 0.0619 | -0.0115 | 0.0886 |
| Gab_fi | Ant fisheries-related buildings | -0.0212 | 0.4546 | 0.0676 | 0.2219 |
| Gab_a | AA built-up | -0.1403 | 0.0228 | -0.2038 | -0.0984 |
| Vmag_a | AA regulated magazine | 0.1229 | 0.3508 | -0.0181 | 0.1816 |
| Veib_a | AA road | -0.1270 | 0.2098 | 0.2641 | -0.0039 |
| Sti_a | AA trail, path | -0.0092 | 0.0534 | -0.0148 | -0.0104 |
| Setr_s | Ant summer mountain pasture | 0.0664 | 0.4185 | 0.0681 | 0.1584 |
| Sefr_a | AA old buildings | 0.1475 | -0.1467 | 0.0418 | -0.0118 |
| Rd_anl_a | AA reindeer husbandry facilities | -0.0421 | 0.4608 | 0.0796 | 0.1794 |
| Build_a | AA built-up area | -0.0140 | 0.2145 | 0.2180 | -0.1530 |
| City_a | AA town/city area | 0.0200 | 0.0228 | -0.0278 | 0.2400 |
| Lled_a | AA power lines | -0.0863 | 0.2933 | 0.1764 | -0.0441 |
| Cul_u_s | Ant cultural heritage sites outdoors | -0.0936 | 0.2366 | 0.1803 | -0.0551 |
| Cul_t_s | Ant technical heritage sites | -0.0265 | 0.3677 | 0.0373 | 0.0390 |
| Cul_k_s | Ant church ruins | -0.1924 | 0.1867 | 0.1376 | 0.0060 |
| Cul_b_s | Ant ancient rock art sites | -0.0664 | 0.2318 | -0.0241 | -0.0168 |
| Cul_a_s | Ant archaeological heritage sites | -0.0360 | 0.4654 | 0.0835 | 0.1768 |
| Abygg_a | AA large buildings | 0.1266 | 0.4848 | 0.0454 | -0.0989 |
| Araaf_a | AA open areas | 0.2257 | 0.0224 | -0.0646 | 0.2286 |
| Arfull_a | AA arable land | -0.1280 | 0.1563 | 0.0143 | -0.0255 |
| Arover_a | AA surface cultivated land | -0.3329 | -0.0018 | 0.8091 | -0.1066 |

Correlations also with reindeer husbandry facilities; (Rd_anl_a; τ = 0.4848); number of fisheries-related buildings (Gab_fi; τ = 0.4546); summer mountain pastures (Setr_s; τ = 0.4185); and regulated magazines (Vmag_a; τ = 0.3508) may indicate that the axis also captures regional variation, i.e. from southern to northern Norway.

##### CLG 4 – (A1) Amount of infrastructure: OI (Table 48, Fig. 128)

The single variable included in the ONV that represents CLG 4, amount of infrastructure (OI), was amount of infrastructure (IfIu). This land-use related CLG had an estimated gradient length of 0.1176/0.08 = 1.47 EDU–L units and was, accordingly, not divided into standard segments (48). The distribution of OUs along this ONV (in its original scaling) was slightly right-skewed (Fig. 128).

This CLG expresses variation in amount of buildings and other infrastructure in inland fine-sediment plains, but the total variation in landscape characteristics was insufficient (as estimated by Method III) to motivate for a division of this CLG into segments.

##### Tentative division of inland fine-sediment plains into minor types

Correlations between ONVs and ordination axes, primary key variables, and analysis variables were calculated as a basis for operationalisation of the CLGs (Table 49). The theoretical number of combinations of standard segments is basically only 3 because only one CLG reached the threshold for segmentation of 2 EDU–L units. However, the clear separation of the two primary soil categories along CLG 1 supported recognition of 6 gradient combinations based upon division of each AP segment into two (sub)segments by dominant soil type (JA). All 6 combinations of AP interval and soil type were represented in the IS118 data subset (Table 50).

**Table 50.** Realised gradient combinations (unique combinations of segments, i.e. tentative minor landscape types) and number of observation units for each combination in inland fine-sediment plains (IF).

| Tentative<br>minor<br>type<br>number | Unique<br>combinations<br>of segments | No. of OUs |
| --- | --- | --- |
| 1 | 1a | 16 |
| 2 | 1b | 15 |
| 3 | 2a | 10 |
| 4 | 2b | 15 |
| 5 | 3a | 36 |
| 6 | 3b | 22 |

#### Inland plains without dominance of fine sediments (‘other inland plains’; IX)

The analyses for identification of independent significant analytic variables for each CLG candidate by forward selection from the *n* = 65 variables using RDA show that five of the six CLG candidates satisfied the requirement for explaining at least 6% of the variation (Table 51). The parsimonious set of CLGs, found by RDA with forward selection among the CLG candidates separately for each functional variable category, contained 4 CLGs (Table 51). The properties of the five CLGs and their associated (ranked primary) ONVs are described below on the basis of Tables 52–55 and Figs 129–137.

**Fig. 129.**
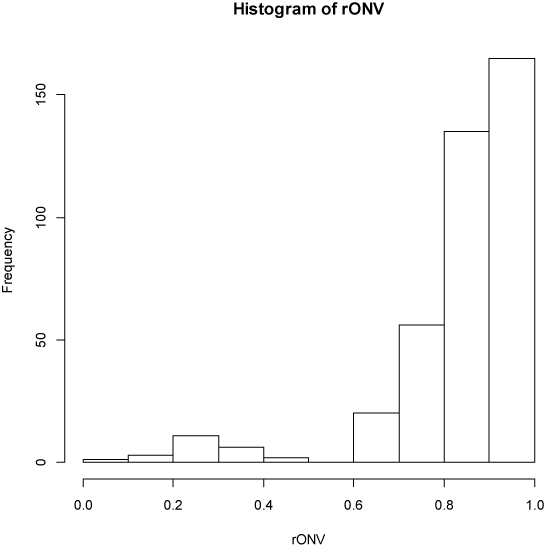
Other inland plains (IX): candidate complex-landscape gradient (CLG) 1, distance to coast (KA): distribution of OUs on the ranged but not rescaled orthogonal key variable (ONV).

**Table 51.**
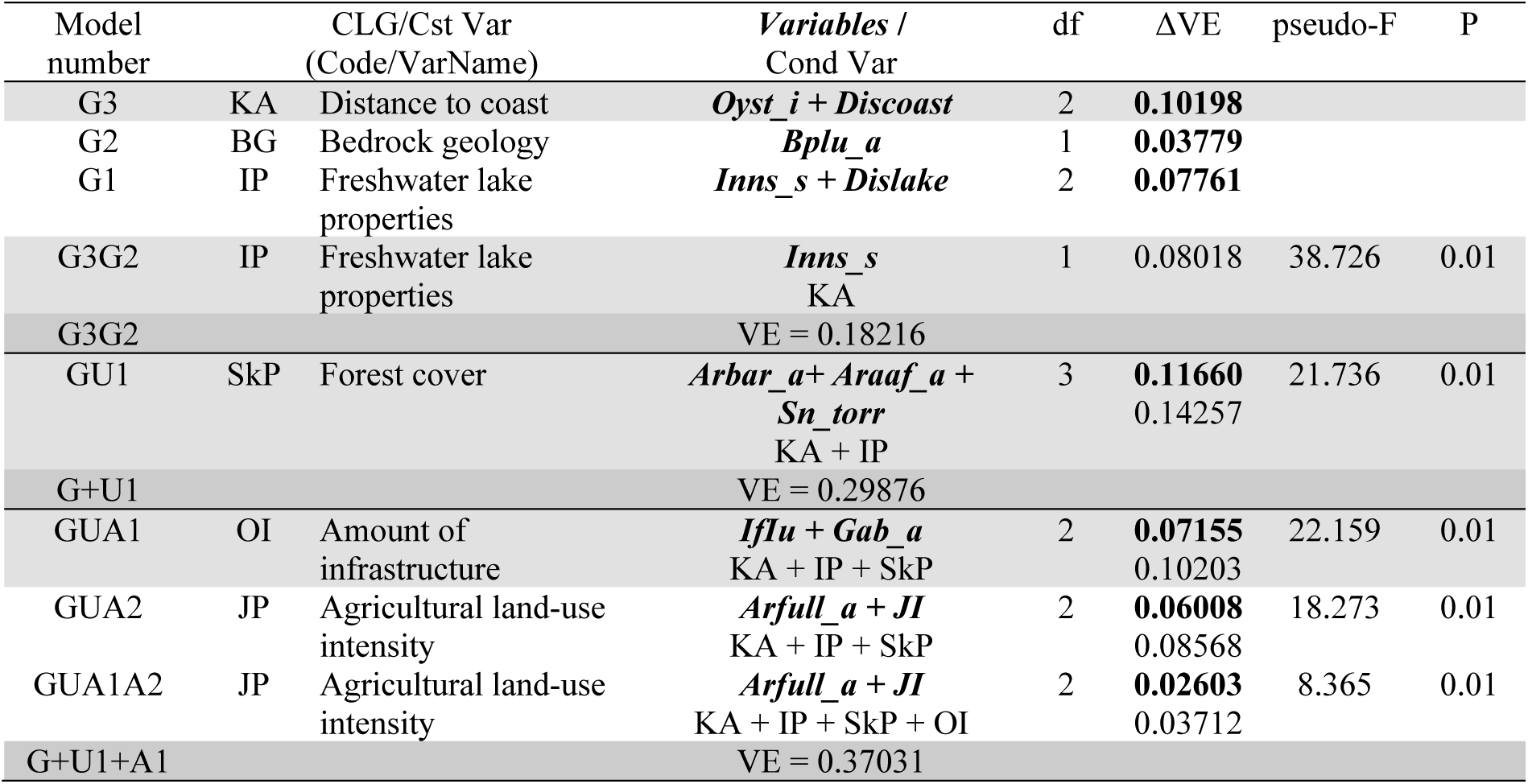
Identification of CLG candidates in other inland plains (IX) by forward selection of variables using RDA. Results of tests of single CLG candidates are shown in bold-face types. The best model at each step in the selection process is highlighted using light grey shading while the best model after testing all CLG candidates in a functional variable category is highlighted using darker grey shading. Abbreviations and explanations: Var = variable; Cst Var = constraining variable; Cond Var = conditioning variable(s); df = degrees of freedom; VE = variation explained (expressed as fraction of the total variation); ΔVE = additional variation explained; pseudo-F and P = F-statistic used in the test of the null hypothesis that adding the variable(s) do not contribute more to explaining variation in landscape-element composition than a random variable, with the associated P value. Variable names are abbreviated according to Tables 4 and 5. For further explanation, se Material and methods chapter.

**Table 52.**
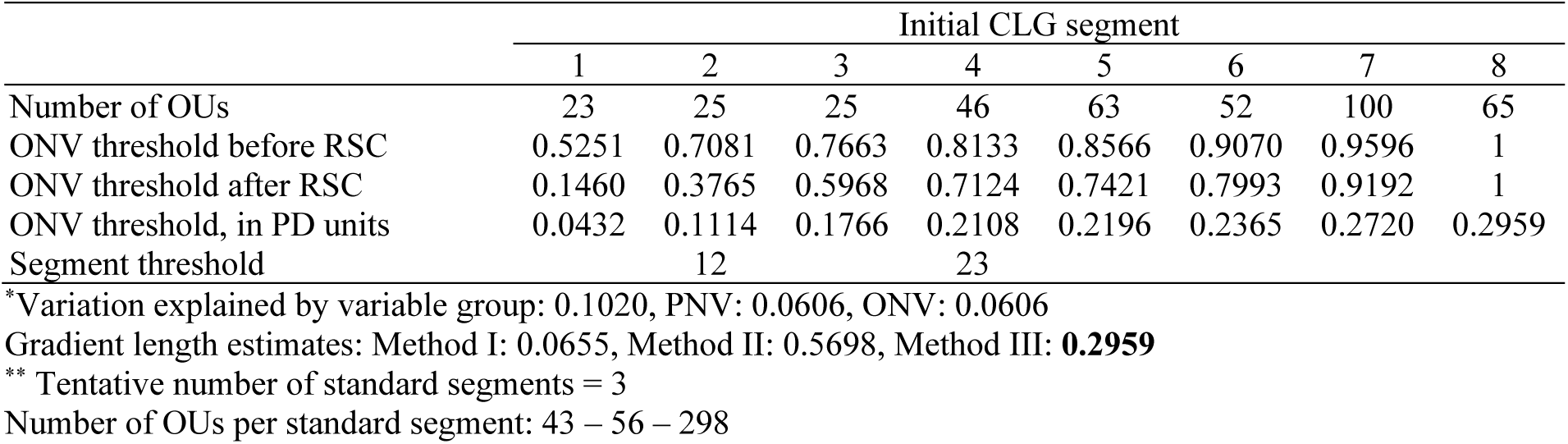
Major type other inland plains (IX): key properties* and segmentation** of candidate complex-landscape gradient G1, distance to coast (KA), as represented by its orthogonal key variable (ONV). OU = observation unit; Threshold = upper limit of class (RSC = rescaling). PD unit = Proportional dissimilarity unit. Segment threshold shows position of standard segment borders relative to initial segments.

**Table 53.** Major type other inland plains (IX): key properties* and segmentation** of candidate complex-landscape gradient G2, freshwater lake properties (IP), as represented by its orthogonal key variable (ONV). OU = observation unit; Threshold = upper limit of class (RSC = rescaling). PD unit = Proportional dissimilarity unit. Segment threshold shows position of standard segment borders relative to initial segments.

|  | Initial CLG segment |  |  |  |  |  |  |  |
| --- | --- | --- | --- | --- | --- | --- | --- | --- |
|  | 1 | 2 | 3 | 4 | 5 | 6 | 7 | 8 |
| Number of OUs | 25 | 32 | 35 | 64 | 66 | 90 | 62 | 25 |
| ONV threshold before RSC | 0.1666 | 0.2601 | 0.3555 | 0.4496 | 0.5434 | 0.6386 | 0.7319 | 1 |
| ONV threshold after RSC | 0.0817 | 0.1709 | 0.3198 | 0.4897 | 0.6277 | 0.7834 | 0.9139 | 1 |
| ONV threshold, in PD units | 0.0263 | 0.0550 | 0.1029 | 0.1576 | 0.2020 | 0.2521 | 0.2941 | 0.3218 |
| Segment threshold |  |  | 12 |  | 23 | 34 |  |  |

\*Variation explained by variable group: Group: 0.0776, PNV: 0.0776, ONV: 0.0767
Gradient length estimates: Method I: Method I: 0.2654, Method II: 0.3842, Method III: **0.3218**
\*\* Tentative number of standard segments = 4
Number of OUs per standard segment: 75 – 84 – 131 – 106

**Table 54.** Major type other inland plains (IX): key properties* and segmentation** of candidate complex-landscape gradient U1 forest cover (SkP), as represented by its orthogonal key variable (ONV). OU = observation unit; Threshold = upper limit of class (RSC = rescaling). PD unit = Proportional dissimilarity unit. Segment threshold shows position of standard segment borders relative to initial segments.

|  | Initial CLG segment |  |  |  |  |  |  |  |
| --- | --- | --- | --- | --- | --- | --- | --- | --- |
|  | 1 | 2 | 3 | 4 | 5 | 6 | 7 | 8 |
| Number of OUs | 25 | 47 | 99 | 58 | 48 | 46 | 51 | 25 |
| ONV threshold before RSC | 0.2090 | 0.3088 | 0.4065 | 0.5075 | 0.6076 | 0.7079 | 0.8077 | 1 |
| ONV threshold after RSC | 0.2416 | 0.4131 | 0.5259 | 0.6783 | 0.7954 | 0.8437 | 0.9102 | 1 |
| ONV threshold, in PD units | 0.1116 | 0.1908 | 0.2429 | 0.3133 | 0.3674 | 0.3897 | 0.4204 | 0.4619 |
| Segment threshold | 12 |  | 23 | 34 |  | 45 |  |  |

\*Variation explained by variable group: Group: 0.1688, PNV: 0.1089, ONV: 0.0720
Gradient length estimates: Method I: 0.3550, Method II: 0.4593, Method III: **0.4619**
\*\* Tentative number of standard segments = 5
Number of OUs per standard segment: 20 – 47 – 130 – 81 – 117

**Table 55.** Major type other inland plains (IX): key properties* and segmentation** of candidate complex-landscape gradient A1, amount of infrastructure (OI), as represented by its orthogonal key variable (ONV). OU = observation unit; Threshold = upper limit of class (RSC = rescaling). PD unit = Proportional dissimilarity unit. Segment threshold shows position of standard segment borders relative to initial segments.

|  | Initial CLG segment |  |  |  |  |  |  |  |
| --- | --- | --- | --- | --- | --- | --- | --- | --- |
|  | 1 | 2 | 3 | 4 | 5 | 6 | 7 | 8 |
| Number of OUs | 25 | 36 | 43 | 83 | 108 | 54 | 25 | 25 |
| ONV threshold before RSC | 0.1549 | 0.2194 | 0.2853 | 0.3481 | 0.4117 | 0.4759 | 0.5812 | 1 |
| ONV threshold after RSC | 0.0239 | 0.0481 | 0.1209 | 0.1747 | 0.2435 | 0.3376 | 0.5997 | 1 |
| ONV threshold, in PD units | 0.0083 | 0.0167 | 0.0420 | 0.0607 | 0.0846 | 0.1173 | 0.2084 | 0.3475 |
| Segment threshold |  |  |  |  | 12 |  | 23 | 34 |

\*Variation explained by variable group: 0.1742, PNV: 0.1395, ONV: 0.0530
Gradient length estimates: Method I: 0.2434, Method II: 0.4028, Method III: 0.3475
\*\* Tentative number of standard segments = 4
Number of OUs per standard segment: 298 – 65 – 18 – 15

##### CLG 1 – (G1) Distance to coast: KA (Table 51, Figs 129–130)

The two variables included in the ONV that represents CLG 1, distance to coast (KA), were: (1) inverse island size (Oyst_i); and (2) distance to coast (Discoast). This geo-ecological CLG had an estimated gradient length of 0.2959/0.08 = 3.699 EDU–L units and was, accordingly, divided into 3 standard segments (Table 52). The distribution of OUs along this ONV (in its original scaling) was strongly left-skewed (Fig. 129).

This CLG separated a ‘coastal group’ with 23 OUs (segment 1; values before rescaling < 0.52) from an inland group (the rest). The variation along KA was strongly non-linear, indicating presence of little systematic variation. Accordingly, we interpret this CLG as an indication that inland plains close to the coast with presence of marine deposits, should be kept separate from other inland plains (Fig. 130). Several of the OUs from inland plains close to the coast were located on islands, but lacking coastline. Except for the correlation with marine deposits (Table 56; Kmar_a; τ = –0.3076), this CLG had no strong correlations with other analysis variables than those already included in the CLG. While Table 52 shows the tentative division of the CLG into three segments suggested by the analyses, the interpretation motivates for a division into two segments only. A critical assessment of the basis for segmentation of this CLG is required prior to practical implementation in a landscape-type system.

**Fig. 130.**
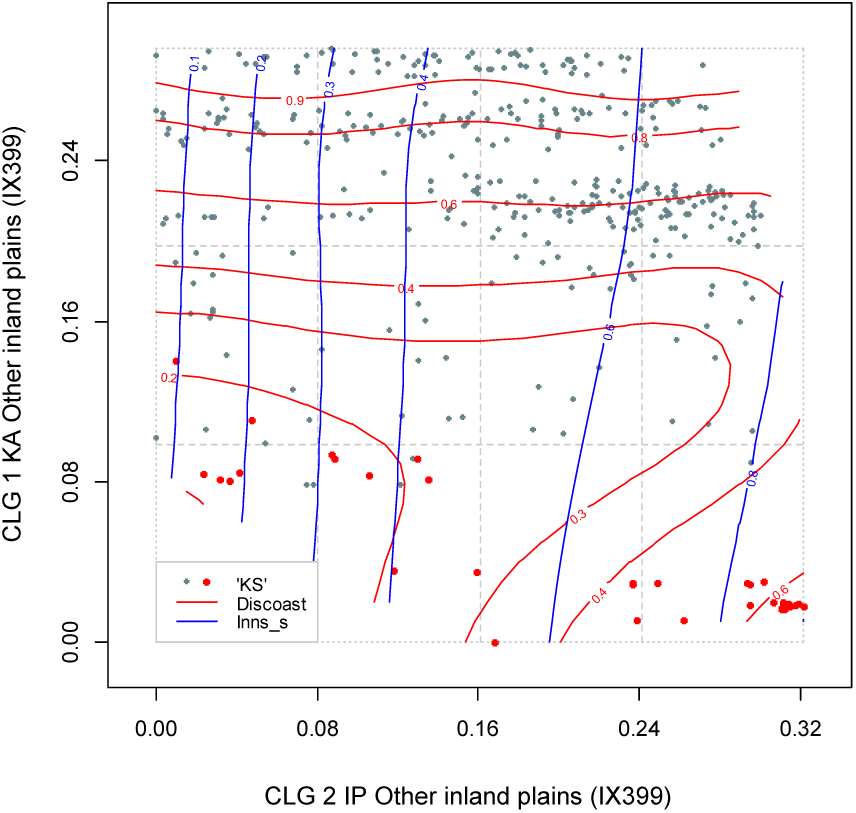
Major type other inland plains (IX): distribution of 399 observation units (IX399 data subset) along rescaled orthogonal key variables (rONVs) for complex landscape gradients CLG 1 (distance to coast; KA) and CLG 2 (freshwater lake properties; IP). Symbols show affiliation of OUs to levels of relevant landscape-gradient (LG) variables, operationalised as ordered factor variables in accordance with Table 3. Isolines are given for relevant primary key variables (Table 4) and analytic variables (Table 5).

**Table 56.**
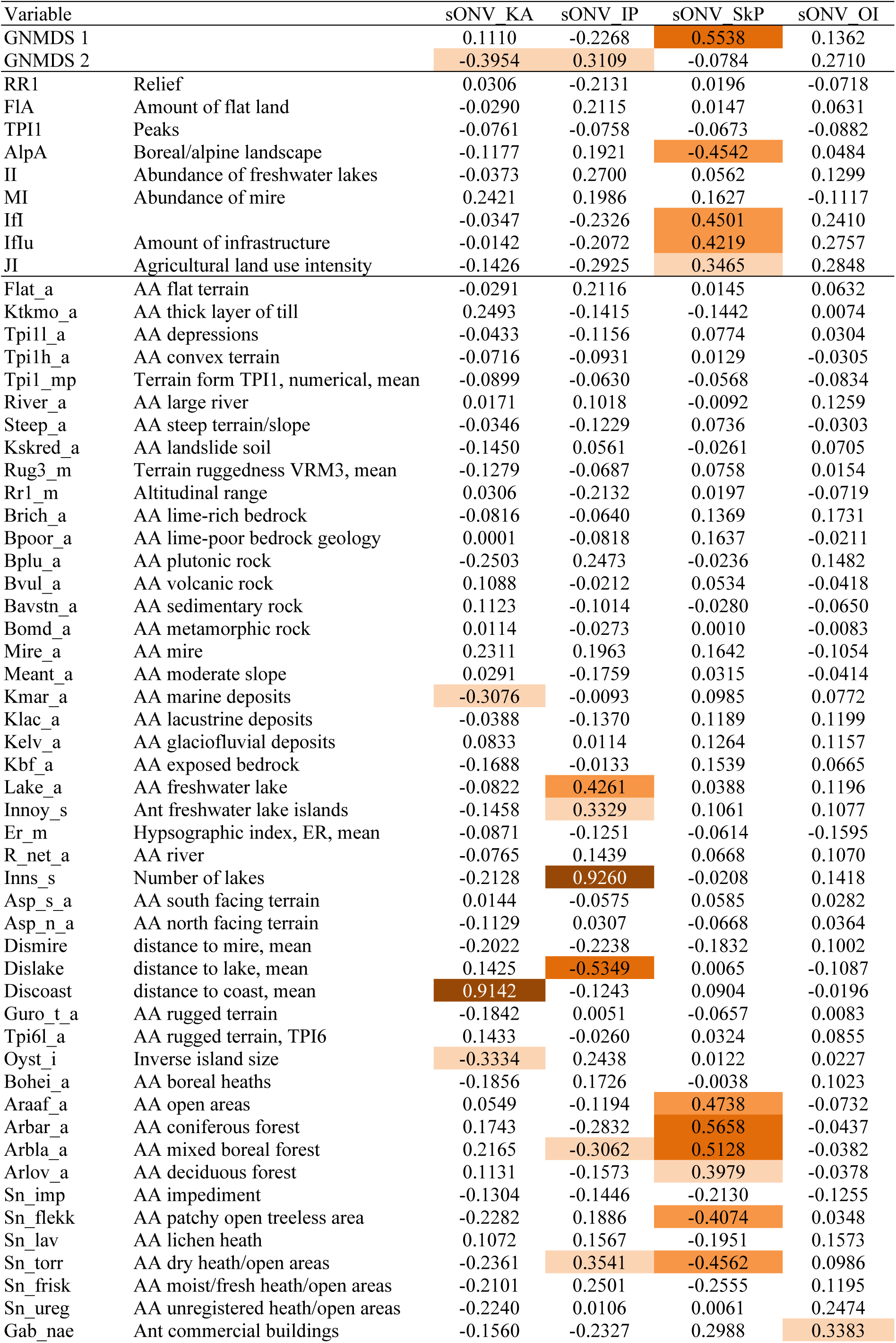
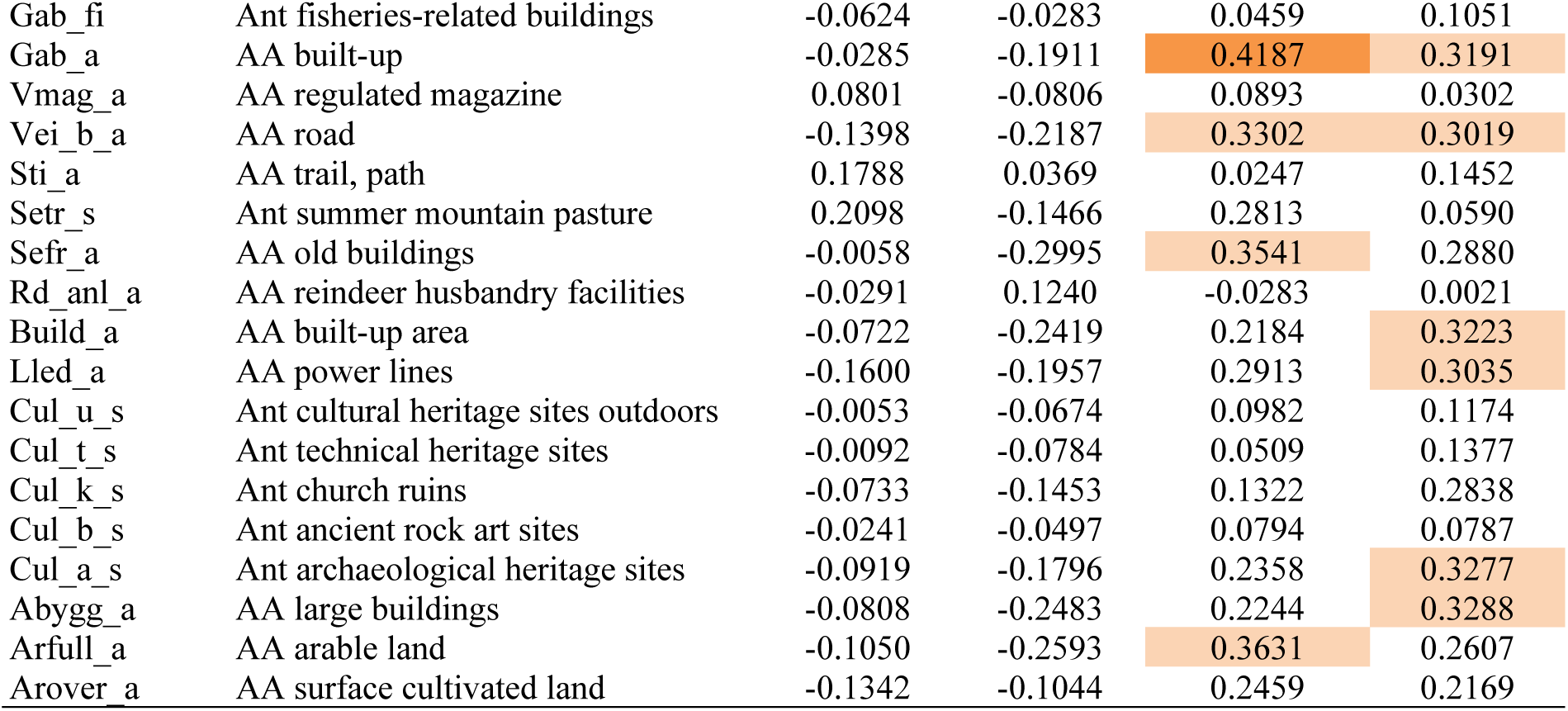
Correlations between the four ONVs and ordination axes (rows 1–2), primary key variables (rows 3–11), and analytic variables in major landscape type other inland plains (IX).

| Variable |  | sONV KA | sONV IP | sONV SkP | sONV OI |
| --- | --- | --- | --- | --- | --- |
| GNMDS 1 |  | 0.1110 | -0.2268 | 0.5538 | 0.1362 |
| GNMDS 2 |  | -0.3954 | 0.3109 | -0.0784 | 0.2710 |
| RR1 | Relief | 0.0306 | -0.2131 | 0.0196 | -0.0718 |
| FlA | Amount of flat land | -0.0290 | 0.2115 | 0.0147 | 0.0631 |
| TPI1 | Peaks | -0.0761 | -0.0758 | -0.0673 | -0.0882 |
| AlpA | Boreal/alpine landscape | -0.1177 | 0.1921 | -0.4542 | 0.0484 |
| II | Abundance of freshwater lakes | -0.0373 | 0.2700 | 0.0562 | 0.1299 |
| MI | Abundance of mire | 0.2421 | 0.1986 | 0.1627 | -0.1117 |
| IfI |  | -0.0347 | -0.2326 | 0.4501 | 0.2410 |
| IfIu | Amount of infrastructure | -0.0142 | -0.2072 | 0.4219 | 0.2757 |
| JI | Agricultural land use intensity | -0.1426 | -0.2925 | 0.3465 | 0.2848 |
| Flat_a | AA flat terrain | -0.0291 | 0.2116 | 0.0145 | 0.0632 |
| Ktkmo_a | AA thick layer of till | 0.2493 | -0.1415 | -0.1442 | 0.0074 |
| Tpi1l_a | AA depressions | -0.0433 | -0.1156 | 0.0774 | 0.0304 |
| Tpi1h_a | AA convex terrain | -0.0716 | -0.0931 | 0.0129 | -0.0305 |
| Tpi1_mp | Terrain form TPI1, numerical, mean | -0.0899 | -0.0630 | -0.0568 | -0.0834 |
| River_a | AA large river | 0.0171 | 0.1018 | -0.0092 | 0.1259 |
| Steep_a | AA steep terrain/slope | -0.0346 | -0.1229 | 0.0736 | -0.0303 |
| Kskred_a | AA landslide soil | -0.1450 | 0.0561 | -0.0261 | 0.0705 |
| Rug3_m | Terrain ruggedness VRM3, mean | -0.1279 | -0.0687 | 0.0758 | 0.0154 |
| Rr1_m | Altitudinal range | 0.0306 | -0.2132 | 0.0197 | -0.0719 |
| Brich_a | AA lime-rich bedrock | -0.0816 | -0.0640 | 0.1369 | 0.1731 |
| Bpoor_a | AA lime-poor bedrock geology | 0.0001 | -0.0818 | 0.1637 | -0.0211 |
| Bplu_a | AA plutonic rock | -0.2503 | 0.2473 | -0.0236 | 0.1482 |
| Bvul_a | AA volcanic rock | 0.1088 | -0.0212 | 0.0534 | -0.0418 |
| Bavstn_a | AA sedimentary rock | 0.1123 | -0.1014 | -0.0280 | -0.0650 |
| Bomd_a | AA metamorphic rock | 0.0114 | -0.0273 | 0.0010 | -0.0083 |
| Mire_a | AA mire | 0.2311 | 0.1963 | 0.1642 | -0.1054 |
| Meant_a | AA moderate slope | 0.0291 | -0.1759 | 0.0315 | -0.0414 |
| Kmar_a | AA marine deposits | -0.3076 | -0.0093 | 0.0985 | 0.0772 |
| Klac_a | AA lacustrine deposits | -0.0388 | -0.1370 | 0.1189 | 0.1199 |
| Kelv_a | AA glaciofluvial deposits | 0.0833 | 0.0114 | 0.1264 | 0.1157 |
| Kbf_a | AA exposed bedrock | -0.1688 | -0.0133 | 0.1539 | 0.0665 |
| Lake_a | AA freshwater lake | -0.0822 | 0.4261 | 0.0388 | 0.1196 |
| Innoy_s | Ant freshwater lake islands | -0.1458 | 0.3329 | 0.1061 | 0.1077 |
| Er_m | Hypsographic index, ER, mean | -0.0871 | -0.1251 | -0.0614 | -0.1595 |
| R_net_a | AA river | -0.0765 | 0.1439 | 0.0668 | 0.1070 |
| Inns_s | Number of lakes | -0.2128 | 0.9260 | -0.0208 | 0.1418 |
| Asp_s_a | AA south facing terrain | 0.0144 | -0.0575 | 0.0585 | 0.0282 |
| Asp_n_a | AA north facing terrain | -0.1129 | 0.0307 | -0.0668 | 0.0364 |
| Dismire | distance to mire, mean | -0.2022 | -0.2238 | -0.1832 | 0.1002 |
| Dislake | distance to lake, mean | 0.1425 | -0.5349 | 0.0065 | -0.1087 |
| Discoast | distance to coast, mean | 0.9142 | -0.1243 | 0.0904 | -0.0196 |
| Guro_t_a | AA rugged terrain | -0.1842 | 0.0051 | -0.0657 | 0.0083 |
| Tpi6l_a | AA rugged terrain, TPI6 | 0.1433 | -0.0260 | 0.0324 | 0.0855 |
| Oyst_i | Inverse island size | -0.3334 | 0.2438 | 0.0122 | 0.0227 |
| Bohei_a | AA boreal heaths | -0.1856 | 0.1726 | -0.0038 | 0.1023 |
| Araaf_a | AA open areas | 0.0549 | -0.1194 | 0.4738 | -0.0732 |
| Arbar_a | AA coniferous forest | 0.1743 | -0.2832 | 0.5658 | -0.0437 |
| Arbla_a | AA mixed boreal forest | 0.2165 | -0.3062 | 0.5128 | -0.0382 |
| Arlov_a | AA deciduous forest | 0.1131 | -0.1573 | 0.3979 | -0.0378 |
| Sn_imp | AA impediment | -0.1304 | -0.1446 | -0.2130 | -0.1255 |
| Sn_flekk | AA patchy open treeless area | -0.2282 | 0.1886 | -0.4074 | 0.0348 |
| Sn_lav | AA lichen heath | 0.1072 | 0.1567 | -0.1951 | 0.1573 |
| Sn_torr | AA dry heath/open areas | -0.2361 | 0.3541 | -0.4562 | 0.0986 |
| Sn_frisk | AA moist/fresh heath/open areas | -0.2101 | 0.2501 | -0.2555 | 0.1195 |
| Sn_ureg | AA unregistered heath/open areas | -0.2240 | 0.0106 | 0.0061 | 0.2474 |
| Gab_nae | Ant commercial buildings | -0.1560 | -0.2327 | 0.2988 | 0.3383 |
| Gab_fi | Ant fisheries-related buildings | -0.0624 | -0.0283 | 0.0459 | 0.1051 |
| Gab_a | AA built-up | -0.0285 | -0.1911 | 0.4187 | 0.3191 |
| Vmag_a | AA regulated magazine | 0.0801 | -0.0806 | 0.0893 | 0.0302 |
| Vei_b_a | AA road | -0.1398 | -0.2187 | 0.3302 | 0.3019 |
| Sti_a | AA trail, path | 0.1788 | 0.0369 | 0.0247 | 0.1452 |
| Setr_s | Ant summer mountain pasture | 0.2098 | -0.1466 | 0.2813 | 0.0590 |
| Sefr_a | AA old buildings | -0.0058 | -0.2995 | 0.3541 | 0.2880 |
| Rd_anl_a | AA reindeer husbandry facilities | -0.0291 | 0.1240 | -0.0283 | 0.0021 |
| Build_a | AA built-up area | -0.0722 | -0.2419 | 0.2184 | 0.3223 |
| Lled_a | AA power lines | -0.1600 | -0.1957 | 0.2913 | 0.3035 |
| Cul_u_s | Ant cultural heritage sites outdoors | -0.0053 | -0.0674 | 0.0982 | 0.1174 |
| Cul_t_s | Ant technical heritage sites | -0.0092 | -0.0784 | 0.0509 | 0.1377 |
| Cul_k_s | Ant church ruins | -0.0733 | -0.1453 | 0.1322 | 0.2838 |
| Cul_b_s | Ant ancient rock art sites | -0.0241 | -0.0497 | 0.0794 | 0.0787 |
| Cul_a_s | Ant archaeological heritage sites | -0.0919 | -0.1796 | 0.2358 | 0.3277 |
| Abygg_a | AA large buildings | -0.0808 | -0.2483 | 0.2244 | 0.3288 |
| Arfull_a | AA arable land | -0.1050 | -0.2593 | 0.3631 | 0.2607 |
| Arover_a | AA surface cultivated land | -0.1342 | -0.1044 | 0.2459 | 0.2169 |

##### CLG 2 – (G2) Freshwater lake properties: IP (Table 53, Figs 131–132)

The single variable that represented CLG 2, freshwater lake properties (IP), was number of lakes (Inns_s). CLG 2 had an estimated gradient length of 0.3218/0.08 = 4.022 EDU–L units and was, accordingly, divided into 4 standard segments (Table 53). The distribution of OUs along this ONV (in its original scaling) was slightly left-skewed, with few OUs representing the high-score end of the rONV (Fig. 131).

**Fig. 131.**
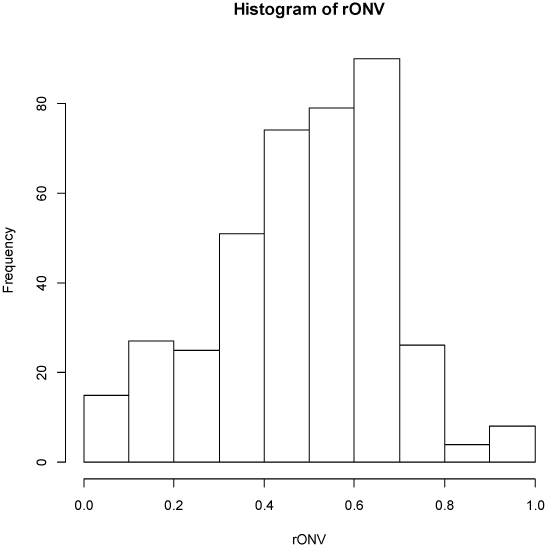
Other inland plains (IX): candidate complex-landscape gradient (CLG) 2, freshwater lake properties (IP): distribution of OUs on the ranged but not rescaled orthogonal key variable (ONV).

**Fig. 132.**
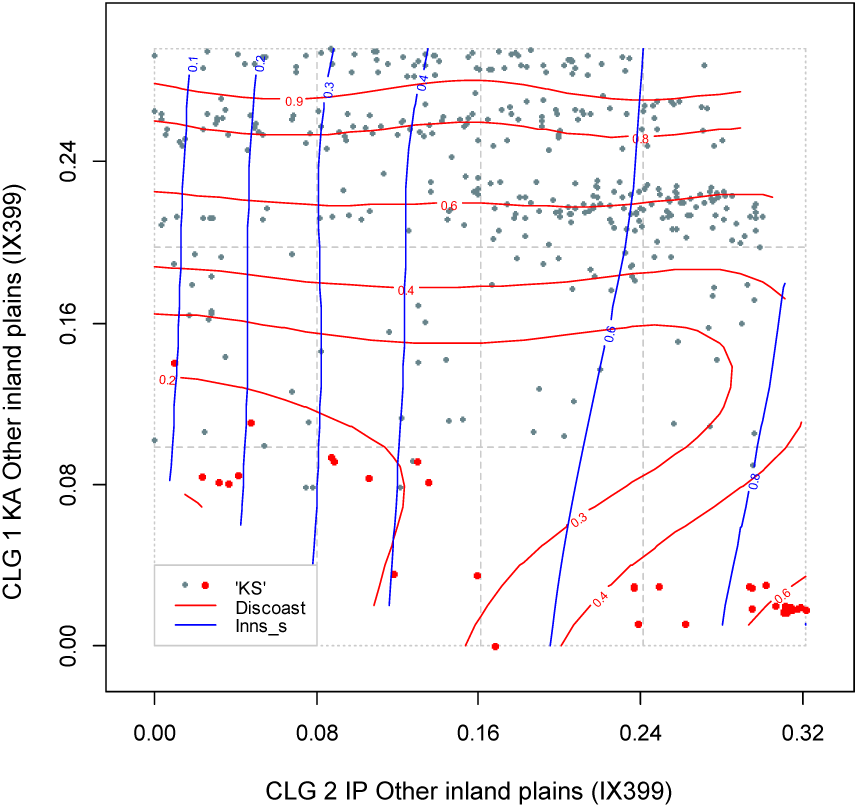
Major type other inland plains (IX): distribution of 399 observation units (IX399 data subset) along rescaled orthogonal key variables (rONVs) for complex landscape gradients CLG 1 (distance to coast; KA) and CLG 2 (freshwater lake properties; IP). Symbols show affiliation of OUs to levels of relevant landscape-gradient (LG) variables, operationalised as ordered factor variables in accordance with Table 3. Isolines are given for relevant primary key variables (Table 4) and analytic variables (Table 5).

This CLG expresses variation within inland plains in the abundance of small lakes and tarns (which are often associated with wetland systems); from low or medium abundance to high abundance. This CLG was negatively correlated with mean distance to lake (Dislake; τ = –0.5349) and mixed boreal forest (Arbla_a; τ = –0.3062), and positively correlated with areal coverage of freshwater lakes (Lake_a; τ = 0.4261), number of freshwater lake islands (Innoy_s; τ = 0.3329) and areal cover of dry heath/open areas (Sn_torr; τ = 0.3541). All of these variables represent landscape properties typical of mountain plateaus (Norwegian: ‘fjellvidde’) with a high abundance of small lakes.

While the gradient-length estimate (Table 53) suggests a tentative division of the CLG into four segments, we question the basis for such a fine division. Most likely, some variation in infrastructure, perhaps also other characteristics, which are correlated with lake abundance, contribute to the large estimated gradient length. It is first and foremost the presence of many small lakes that define the ‘end’ of this CLG.

##### CLG 3 – (U1) Forest cover: SkP (Table 54, Figs 133–134)

The three variables included in the ONV that represents CLG 3, forest cover (SkP), were the areal coverage of: (1) coniferous forest (Arbar_a); (2) open areas (Araaf_a); and (3) open areas with dry heath (Sn_torr). CLG 3, SkP, had an estimated gradient length of 0.4619/0.08 = 5.774 EDU–L units and was, accordingly, divided into 5 standard segments (Table 54). The distribution of OUs along this ONV (in its original scaling) was slightly right-skewed (Fig. 133).

**Fig. 133.**
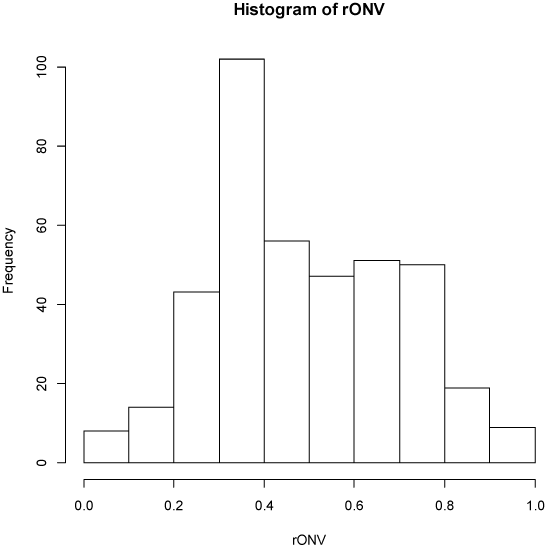
Other inland plains (IX): candidate complex-landscape gradient (CLG) 3, forest cover (SkP): distribution of OUs on the ranged but not rescaled orthogonal key variable (ONV).

This CLG was strongly correlated with the key variable amount of alpine areas (Alp_a; τ = 0.5538) and six analytical variables related to vegetation cover (Table 56). Accordingly, this CLG expresses variation in vegetation cover within inland plains, from barren mountain plains without or with sparse vegetation cover to forested or potentially forested plains below the climatic forest line (Fig. 134). The intermediate segments consist of plains with open mountain heaths and boreal heath-dominated areas below the climatic forest line, kept open as a result of historical land use by logging and grazing. The CLG was also strongly correlated with key variables for the amount of infrastructure (Ifl; τ = 0.4501) and agricultural land-use intensity (JI; τ = 0.3465), and had correlation coefficients τ > 0.3 with four land-use and building abundance variables (Table 56).

**Fig. 134.**
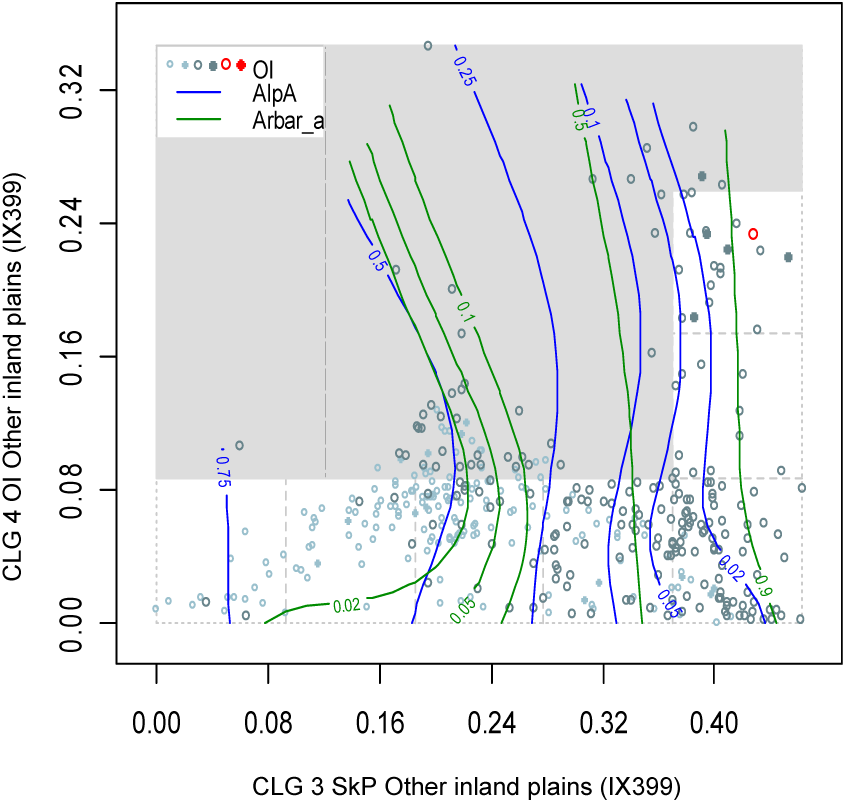
Major type inland other inland plains (IX): distribution of 399 observation units (IX399 data subset) along rescaled orthogonal key variables (rONVs) for complex landscape gradients CLG 3 (forest cover; SkP) and CLG 4 (land use intensity; OI). Symbols show affiliation of OUs to levels of relevant landscape-gradient (LG) variables, operationalised as ordered factor variables in accordance with Table 3. Isolines are given for relevant primary key variables (Table 4) and analytic variables (Table 5).

Although the gradient-length estimate (Table 54) suggests a tentative division of the CLG into five segments, the basis for such a fine division is questionable.

##### CLG 4 – (A1) Amount of infrastructure: OI (Table 55, Figs 135–137)

The two variables included in the ONV that represents CLG 4, amount of infrastructure (OI), were: (1) the amount of infrastructure (IfIu) and (2) areal coverage of built-up areas (Gab_a). This land-use related CLG had an estimated gradient length of 0.3475/0.08 = 4.344 EDU–L units and was, accordingly, divided into 4 standard segments (Table 55). The distribution of OUs along this ONV (in its original scaling) was right-skewed and leptokurtic (Fig. 135).

**Fig. 135.**
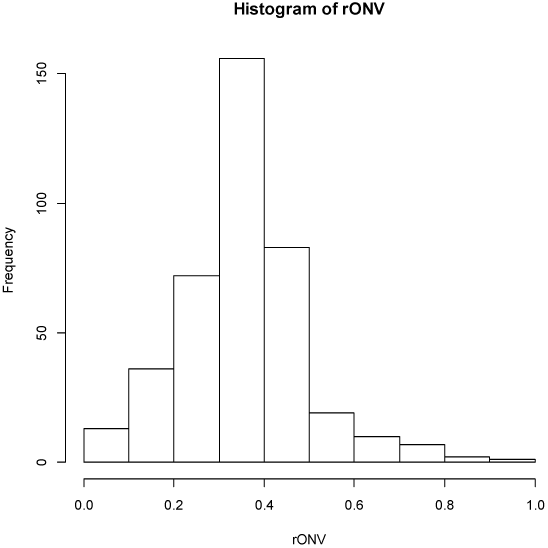
Other inland plains (IX): candidate complex-landscape gradient (CLG) 4, land use intensity (OI): distribution of OUs on the ranged but not rescaled orthogonal key variable (ONV).

This CLG separates the few OUs in other inland plains with a high amount of buildings and other infrastructure from other OUs. Most of the variation related to density of infrastructure occurred in the lowlands (Fig. 136). Variables that express variation in agricultural land-use intensity were relatively weakly correlated with this CLG (Table 56; τ < 0.3), but seemed to follow the variation in the amount of infrastructure (Fig. 137).

**Figs. 136–137.**
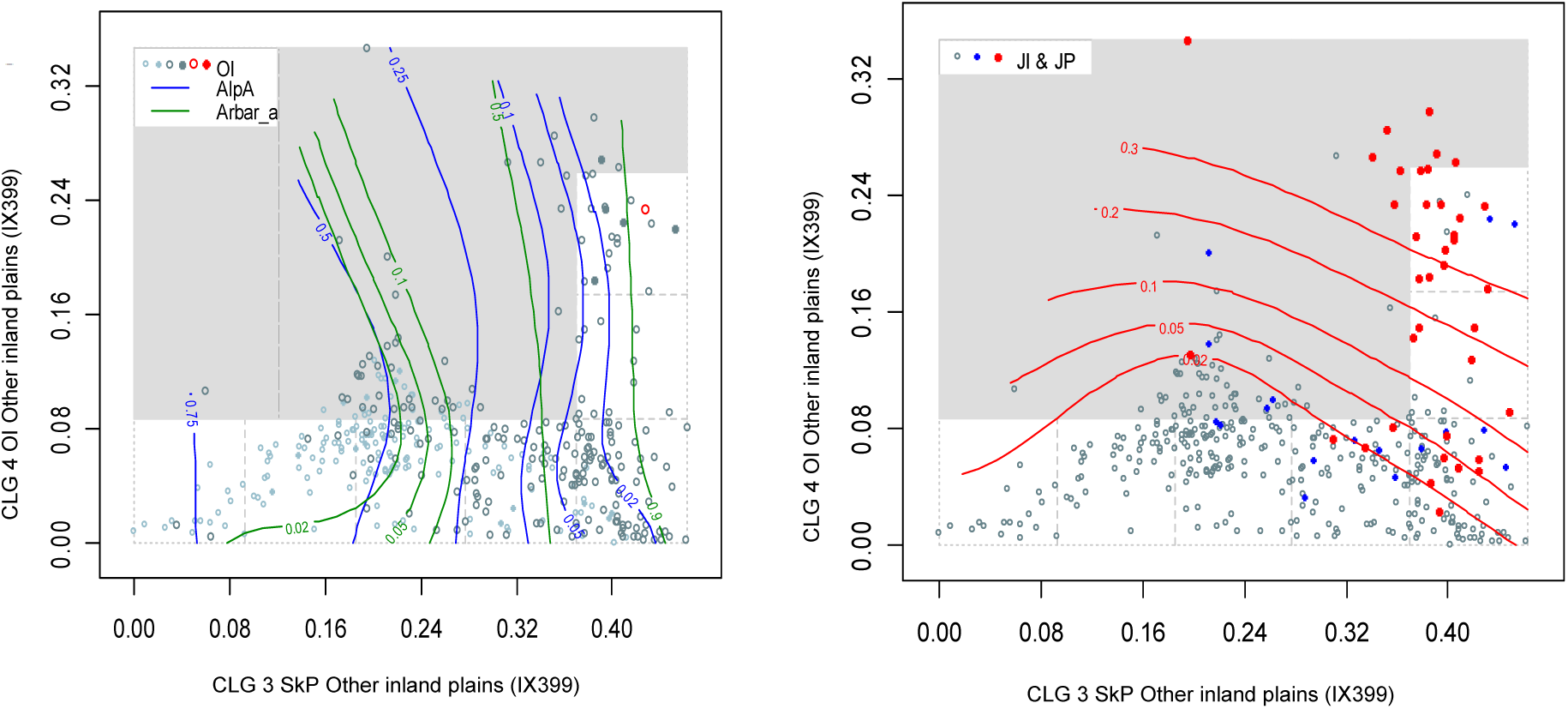
Major type other inland plains (IX): distribution of 399 observation units (IX399 data subset) along rescaled orthogonal key variables (rONVs) for complex landscape gradients CLG 3 (forest cover; SkP) and CLG 4 (land use intensity; OI). Symbols show affiliation of OUs to levels of relevant landscape-gradient (LG) variables, operationalised as ordered factor variables in accordance with Table 3. Isolines are given for relevant primary key variables (Table 4) and analytic variables (Table 5).

Although the gradient-length estimate (Table 55) suggests a tentative division of the CLG into four segments, the basis for such a fine division can be questioned. The heterogeneous distribution of OUs in the ‘landscape space’ spanned by the CLGs SkP and OI (Figs 136–137) indicate that the suggested 4 segments can be reduced to 3 without significant loss of information.

##### Tentative division of other inland plains into minor types

Correlations between ONVs and ordination axes, primary key variables, and analysis variables were calculated as a basis for operationalisation of the CLGs (Table 56). The theoretical number of combinations of intervals along gradients is 3 × 4 × 5 × 4 = 240. Of these, 75 were represented in the total data set (Table 57); 34 of which with > 4 OUs (> 1% of the IX399 data subset). By far the most commonly encountered combination in our data set was 2121, i.e. inland plains distant from the coast with low abundance of freshwater lakes, heath or boreal heath vegetation, low amount of infrastructure and low agricultural land-use intensity. This combination of segments represents typical properties for mountain plateaus (in Norwegian: ‘fjellvidde’) in Norway.

**Table 57.**
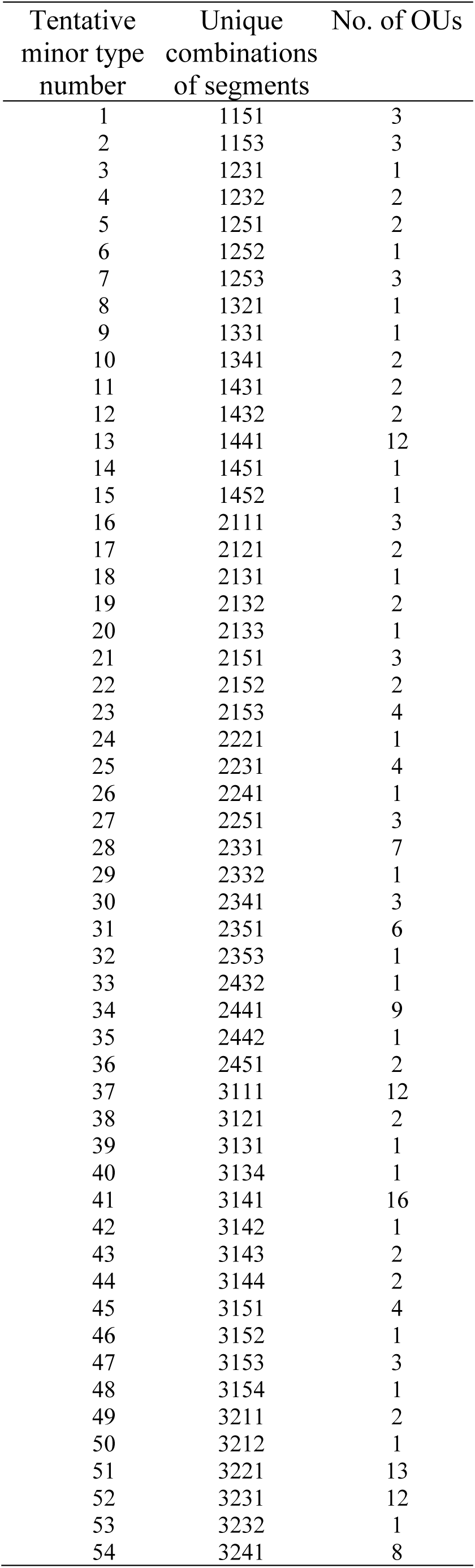
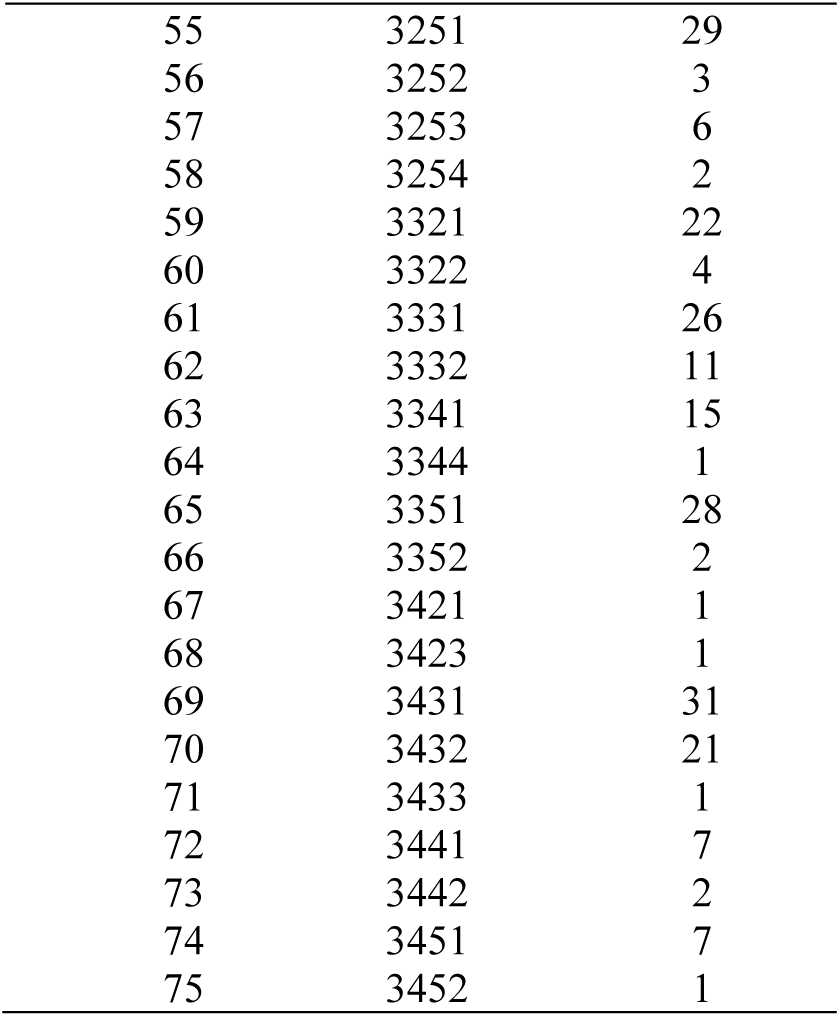
Realised gradient combinations (unique combinations of segments, i.e. tentative minor landscape types) and number of observation units for each combination in other inland plains (IX).

| Tentative<br>minor type<br>number | Unique<br>combinations<br>of segments | No. of OUs |
| --- | --- | --- |
| 1 | 1151 | 3 |
| 2 | 1153 | 3 |
| 3 | 1231 | 1 |
| 4 | 1232 | 2 |
| 5 | 1251 | 2 |
| 6 | 1252 | 1 |
| 7 | 1253 | 3 |
| 8 | 1321 | 1 |
| 9 | 1331 | 1 |
| 10 | 1341 | 2 |
| 11 | 1431 | 2 |
| 12 | 1432 | 2 |
| 13 | 1441 | 12 |
| 14 | 1451 | 1 |
| 15 | 1452 | 1 |
| 16 | 2111 | 3 |
| 17 | 2121 | 2 |
| 18 | 2131 | 1 |
| 19 | 2132 | 2 |
| 20 | 2133 | 1 |
| 21 | 2151 | 3 |
| 22 | 2152 | 2 |
| 23 | 2153 | 4 |
| 24 | 2221 | 1 |
| 25 | 2231 | 4 |
| 26 | 2241 | 1 |
| 27 | 2251 | 3 |
| 28 | 2331 | 7 |
| 29 | 2332 | 1 |
| 30 | 2341 | 3 |
| 31 | 2351 | 6 |
| 32 | 2353 | 1 |
| 33 | 2432 | 1 |
| 34 | 2441 | 9 |
| 35 | 2442 | 1 |
| 36 | 2451 | 2 |
| 37 | 3111 | 12 |
| 38 | 3121 | 2 |
| 39 | 3131 | 1 |
| 40 | 3134 | 1 |
| 41 | 3141 | 16 |
| 42 | 3142 | 1 |
| 43 | 3143 | 2 |
| 44 | 3144 | 2 |
| 45 | 3151 | 4 |
| 46 | 3152 | 1 |
| 47 | 3153 | 3 |
| 48 | 3154 | 1 |
| 49 | 3211 | 2 |
| 50 | 3212 | 1 |
| 51 | 3221 | 13 |
| 52 | 3231 | 12 |
| 53 | 3232 | 1 |
| 54 | 3241 | 8 |
| 55 | 3251 | 29 |
| 56 | 3252 | 3 |
| 57 | 3253 | 6 |
| 58 | 3254 | 2 |
| 59 | 3321 | 22 |
| 60 | 3322 | 4 |
| 61 | 3331 | 26 |
| 62 | 3332 | 11 |
| 63 | 3341 | 15 |
| 64 | 3344 | 1 |
| 65 | 3351 | 28 |
| 66 | 3352 | 2 |
| 67 | 3421 | 1 |
| 68 | 3423 | 1 |
| 69 | 3431 | 31 |
| 70 | 3432 | 21 |
| 71 | 3433 | 1 |
| 72 | 3441 | 7 |
| 73 | 3442 | 2 |
| 74 | 3451 | 7 |
| 75 | 3452 | 1 |

#### Inland valleys (ID)

The analyses for identification of independent significant analytic variables for each CLG candidate by forward selection from the *n* = 67 variables using RDA show that six of the eight CLG candidates satisfied the requirement for explaining at least 6% of the variation. The parsimonious set of CLGs, found by RDA with forward selection among the CLG candidates separately for each functional variable category, contained 4 CLGs (Table 58). The properties of the four CLGs and their associated (ranked primary) ONVs are described below on the basis of Tables 59–64 and Figs 138–148.

**Fig. 138.**
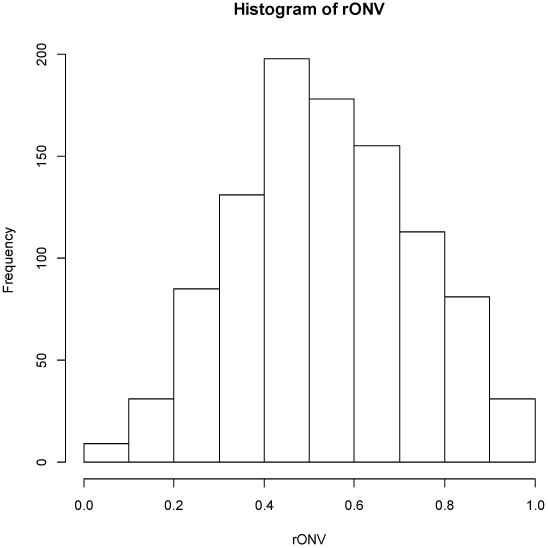
Inland valleys (ID): candidate complex-landscape gradient (CLG) 1, relief (RE): distribution of OUs on the ranged but not rescaled orthogonal key variable (ONV).

**Table 58.**
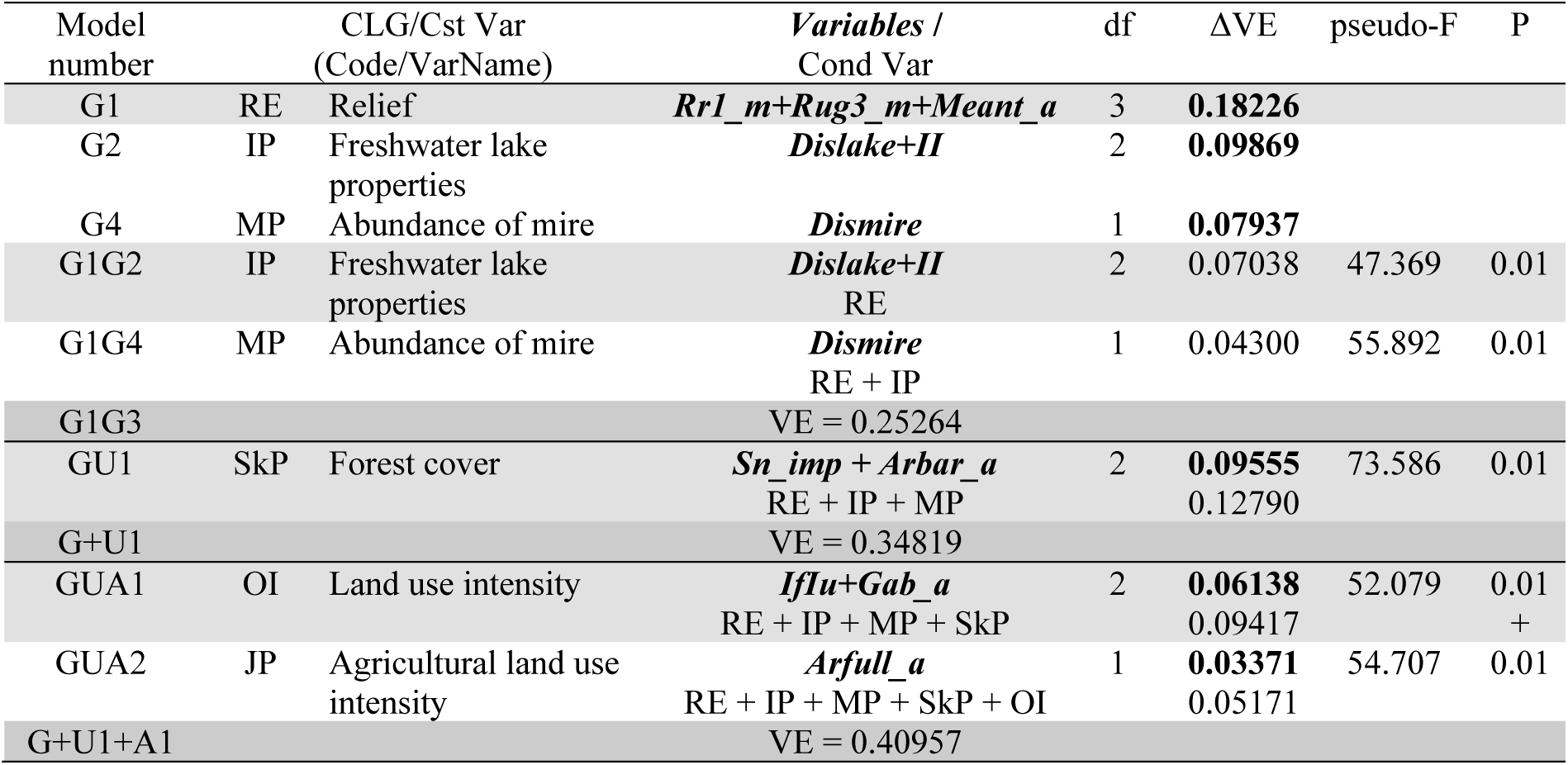
Identification of CLG candidates in inland valleys by forward selection of variables using RDA. Results of tests of single CLG candidates are shown in bold-face types. The best model at each step in the selection process is highlighted using light grey shading while the best model after testing all CLG candidates in a functional variable category is highlighted using darker grey shading. Abbreviations and explanations: Var = variable; Cst Var = constraining variable; Cond Var = conditioning variable(s); df = degrees of freedom; VE = variation explained (expressed as fraction of the total variation); ΔVE = additional variation explained; pseudo-F and P = F-statistic used in the test of the null hypothesis that adding the variable(s) do not contribute more to explaining variation in landscape-element composition than a random variable, with the associated P value. Variable names are abbreviated according to Tables 4 and 5. For further explanation, se Material and methods chapter.

**Table 59.** Major type inland valleys (ID): key properties* and segmentation** of candidate complex-landscape gradient G1, relief (RE), as represented by its orthogonal key variable (ONV). OU = observation unit; Threshold = upper limit of class (RSC = rescaling). PD unit = Proportional dissimilarity unit. Segment threshold shows position of standard segment borders relative to initial segments.

|  | Initial CLG segment |  |  |  |  |  |  |  |
| --- | --- | --- | --- | --- | --- | --- | --- | --- |
|  | 1 | 2 | 3 | 4 | 5 | 6 | 7 | 8 |
| Number of OUs | 25 | 84 | 160 | 223 | 217 | 146 | 112 | 45 |
| ONV threshold before RSC | 0.1703 | 0.2886 | 0.4074 | 0.5259 | 0.6454 | 0.7629 | 0.8806 | 1 |
| ONV threshold after RSC | 0.1878 | 0.3458 | 0.4632 | 0.5801 | 0.6908 | 0.8029 | 0.9119 | 1 |
| ONV threshold, in PD units | 0.0998 | 0.1837 | 0.2461 | 0.3082 | 0.3670 | 0.4266 | 0.4845 | 0.5313 |
| Segment threshold | 12 | 23 |  | 34 |  | 45 | 56 |  |

\*Variation explained by variable group: 0.1823, PNV: 0.1343, ONV: 0.1343
Gradient length estimates: Method I: 0.4923, Method II: 0.4924, Method III: 0.5313
\*\* Tentative number of standard segments = 6
Number of OUs per standard segment: 22 – 79 – 236 – 321 – 224 – 135

**Table 60.** Major type inland valleys (ID): key properties* and segmentation** of candidate complex-landscape gradient G2, freshwater lake properties (IP), as represented by its orthogonal key variable (ONV). OU = observation unit; Threshold = upper limit of class (RSC = rescaling). PD unit = Proportional dissimilarity unit. Segment threshold shows position of standard segment borders relative to initial segments.

|  | Initial CLG segment |  |  |  |  |  |  |  |
| --- | --- | --- | --- | --- | --- | --- | --- | --- |
|  | 1 | 2 | 3 | 4 | 5 | 6 | 7 | 8 |
| Number of OUs | 25 | 57 | 119 | 198 | 201 | 224 | 163 | 25 |
| ONV threshold before RSC | 0.2906 | 0.3829 | 0.4765 | 0.5703 | 0.6635 | 0.7567 | 0.8499 | 1 |
| ONV threshold after RSC | 0.0825 | 0.1871 | 0.2944 | 0.3718 | 0.4715 | 0.6253 | 0.8089 | 1 |
| ONV threshold, in PD units | 0.0312 | 0.0708 | 0.1114 | 0.1407 | 0.1784 | 0.2366 | 0.3061 | 0.3784 |
| Segment threshold |  |  | 12 |  |  | 23 | 34 |  |

\*Variation explained by variable group: 0.0987, PNV: 0.0633, ONV: 0.0558
Gradient length estimates: Method I: 0.3823, Method II: 0.3588, Method III: 0.3784
\*\* Tentative number of standard segments = 4
Number of OUs per standard segment: 151 – 489 – 292 – 77

**Table 61.**
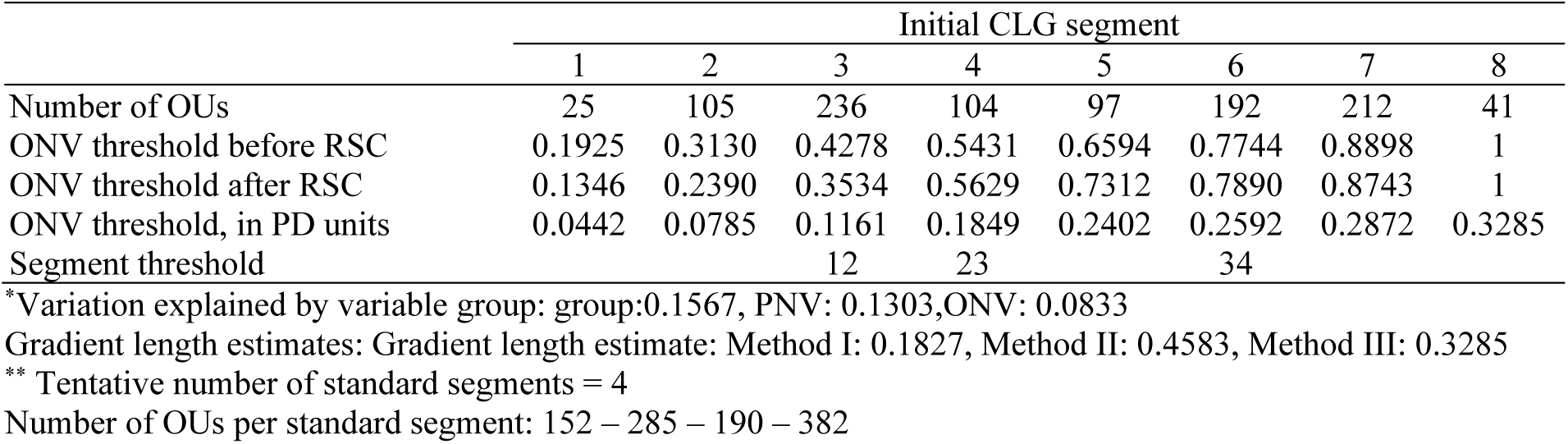
Major type inland valleys (ID): key properties* and segmentation** of candidate complex-landscape gradient U1, forest cover (SkP), as represented by its orthogonal key variable (ONV). OU = observation unit; Threshold = upper limit of class (RSC = rescaling). PD unit = Proportional dissimilarity unit. Segment threshold shows position of standard segment borders relative to initial segments.

|  | Initial CLG segment |  |  |  |  |  |  |  |
| --- | --- | --- | --- | --- | --- | --- | --- | --- |
|  | 1 | 2 | 3 | 4 | 5 | 6 | 7 | 8 |
| Number of OUs | 25 | 105 | 236 | 104 | 97 | 192 | 212 | 41 |
| ONV threshold before RSC | 0.1925 | 0.3130 | 0.4278 | 0.5431 | 0.6594 | 0.7744 | 0.8898 | 1 |
| ONV threshold after RSC | 0.1346 | 0.2390 | 0.3534 | 0.5629 | 0.7312 | 0.7890 | 0.8743 | 1 |
| ONV threshold, in PD units | 0.0442 | 0.0785 | 0.1161 | 0.1849 | 0.2402 | 0.2592 | 0.2872 | 0.3285 |
| Segment threshold |  |  | 12 | 23 |  | 34 |  |  |

\*Variation explained by variable group: group:0.1567, PNV: 0.1303, ONV: 0.0833
Gradient length estimates: Gradient length estimate: Method I: 0.1827, Method II: 0.4583, Method III: 0.3285
\*\* Tentative number of standard segments = 4
Number of OUs per standard segment: 152 – 285 – 190 – 382

**Table 62.**
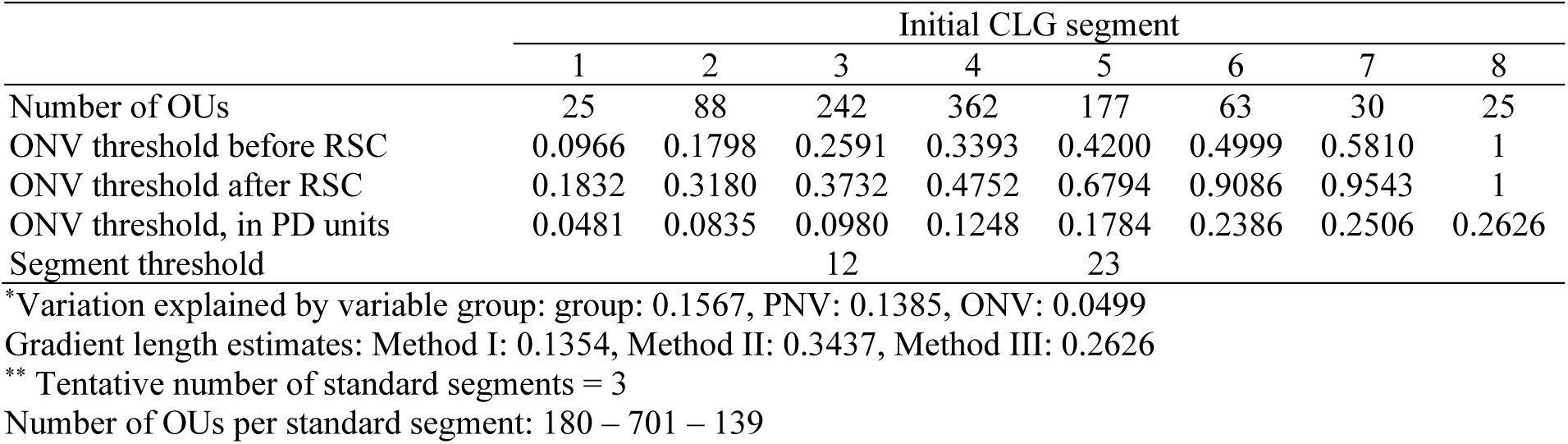
Major type inland valleys (ID): key properties* and segmentation** of candidate complex-landscape gradient A1, land use intensity (OI), as represented by its orthogonal key variable (ONV). OU = observation unit; Threshold = upper limit of class (RSC = rescaling). PD unit = Proportional dissimilarity unit. Segment threshold shows position of standard segment borders relative to initial segments.

**Table 63.** Correlations between the four ONVs and ordination axes (rows 1–3), primary key variables (rows 4–14), and analytic variables in major landscape type inland valleys (ID).

| Variable |  | sONV RE | sONV IP | sONV SkP | sONV OI |
| --- | --- | --- | --- | --- | --- |
| GNMDS 1 |  | -0.3791 | -0.2275 | 0.4101 | 0.1170 |
| GNMDS 2 |  | -0.5108 | 0.0853 | -0.2437 | -0.2086 |
| GNMDS3 |  | -0.0670 | 0.4405 | 0.1486 | 0.0441 |
| DNI | Valley shape | 0.7914 | 0.0167 | 0.0093 | 0.0062 |
| RR1 | Relief | 0.8020 | -0.0025 | 0.0227 | 0.0129 |
| FLA | Amount of flat land | -0.5008 | -0.1710 | 0.0316 | 0.0454 |
| TPI1 | Peaks | -0.7444 | -0.0964 | 0.0001 | 0.0037 |
| BrI | Glacier | 0.2446 | 0.0496 | -0.2083 | -0.0063 |
| AlpA | Boreal/alpine landscape | 0.2197 | 0.1491 | -0.5309 | -0.0531 |
| II | Abundance of freshwater lakes | -0.0434 | 0.4619 | 0.0098 | 0.0657 |
| MI | Abundance of mire | -0.3359 | -0.0973 | 0.1813 | -0.1776 |
| IfI | Amount of infrastructure (OI) | -0.1746 | -0.1764 | 0.4204 | 0.3883 |
| IfIu | Amount of infrastructure | -0.1577 | -0.1865 | 0.3885 | 0.4386 |
| JI | Agricultural land use intensity | -0.1528 | -0.2715 | 0.3382 | 0.3503 |
| Flat_a | AA flat terrain | -0.4990 | -0.1702 | 0.0317 | 0.0453 |
| Ktkmo_a | AA thick layer of till | -0.1949 | -0.0969 | -0.0636 | -0.0426 |
| Tpil1_a | AA depressions | 0.7127 | 0.0962 | 0.0201 | 0.0145 |
| Tpil1h_a | AA convex terrain | 0.4108 | -0.1098 | 0.0552 | 0.0288 |
| Tpil1_mp | Terrain form TPI1, numerical, mean | -0.7331 | -0.0948 | 0.0002 | 0.0047 |
| River_a | AA large river | -0.1874 | -0.2694 | 0.0657 | 0.0461 |
| Steep_a | AA steep terrain/slope | 0.8023 | -0.0583 | 0.0365 | -0.0052 |
| Kskred_a | AA landslide soil | 0.5077 | 0.0355 | -0.0640 | 0.0111 |
| Rug3_m | Terrain ruggedness VRM3, mean | 0.7850 | 0.0031 | 0.0202 | 0.0006 |
| Rr1_m | Altitudinal range | 0.8020 | -0.0025 | 0.0227 | 0.0129 |
| Brich_a | AA lime-rich bedrock | -0.1431 | -0.0769 | 0.0586 | 0.0684 |
| Bpoor_a | AA lime-poor bedrock geology | 0.0781 | 0.0770 | 0.1411 | -0.0120 |
| Bplu_a | AA plutonic rock | 0.1259 | 0.1285 | -0.0608 | 0.0152 |
| Bvul_a | AA volcanic rock | -0.0811 | 0.0142 | 0.0701 | 0.0520 |
| Bavstn_a | AA sedimentary rock | -0.2559 | -0.1755 | -0.0305 | 0.0229 |
| Bomd_a | AA metamorphic rock | 0.0539 | 0.0359 | 0.0944 | -0.0064 |
| Mire_a | AA mire | -0.3164 | -0.0926 | 0.1663 | -0.1675 |
| Glac_a | AA Glacier | 0.2238 | 0.0365 | -0.1935 | -0.0158 |
| Meant_a | AA moderate slope | -0.5420 | -0.0054 | -0.0080 | -0.0146 |
| Kmar_a | AA marine deposits | -0.1152 | -0.1460 | 0.1443 | 0.1495 |
| Klac_a | AA lacustrine deposits | -0.1072 | -0.0483 | -0.0271 | 0.0055 |
| Kelv_a | AA glaciofluvial deposits | -0.1927 | -0.3023 | 0.0911 | 0.0832 |
| Kbf_a | AA exposed bedrock | 0.3452 | 0.1940 | -0.0776 | 0.0203 |
| Lake_a | AA freshwater lake | -0.0562 | 0.5429 | 0.0038 | 0.0601 |
| Innoy_s | Ant freshwater lake islands | -0.0839 | 0.3714 | 0.0519 | 0.0327 |
| Er_m | Hypsographic index, ER, mean | 0.0635 | -0.3584 | -0.0366 | -0.0880 |
| R_net_a | AA river | 0.0301 | -0.2325 | 0.0089 | -0.0954 |
| Inns_s | Number of lakes | -0.0429 | 0.3561 | -0.1129 | -0.0454 |
| Asp_s_a | AA south facing terrain | 0.0609 | 0.0090 | -0.0214 | 0.0080 |
| Asp_n_a | AA north facing terrain | 0.1021 | 0.0051 | -0.0133 | -0.0141 |
| Dismire | distance to mire, mean | 0.3149 | 0.0842 | -0.2450 | 0.1540 |
| Dislake | distance to lake, mean | 0.0193 | -0.6275 | -0.0161 | -0.0450 |
| Discoast | distance to coast, mean | -0.2867 | 0.0277 | 0.0414 | -0.0306 |
| Guro_t_a | AA rugged terrain | 0.7797 | 0.0162 | -0.0396 | -0.0044 |
| Tpi6l_a | AA rugged terrain, TPI6 | -0.1127 | 0.0371 | 0.0705 | 0.0248 |
| Oyst_i | Inverse island size | 0.0918 | 0.1022 | 0.0355 | 0.0236 |
| Bohei_a | AA boreal heaths | 0.1713 | 0.0148 | -0.1539 | 0.0031 |
| Araaf_a | AA open areas | 0.0577 | -0.1143 | 0.2355 | 0.0671 |
| Arbar_a | AA coniferous forest | -0.1624 | -0.1825 | 0.6459 | -0.0138 |
| Arbla_a | AA mixed boreal forest | -0.1410 | -0.1553 | 0.5467 | 0.0226 |
| Arlov_a | AA deciduous forest | 0.0511 | -0.2331 | 0.2812 | 0.0576 |
| Sn_imp | AA impediment | 0.3558 | 0.1137 | -0.4927 | 0.0481 |
| Sn_flekk | AA patchy open treeless area | 0.2901 | 0.1384 | -0.4291 | -0.0258 |
| Sn_lav | AA lichen heath | -0.1383 | -0.0012 | -0.1341 | -0.0295 |
| Sn_torr | AA dry heath/open areas | 0.1363 | 0.0887 | -0.3933 | -0.1110 |
| Sn_frisk | AA moist/fresh heath/open areas | 0.1331 | 0.0711 | -0.2919 | -0.0796 |
| Sn_ureg | AA unregistered heath/open areas | -0.0820 | -0.0856 | -0.0422 | 0.1664 |
| Gab_nae | Ant commercial buildings | -0.1214 | -0.1609 | 0.3040 | 0.3925 |
| Gab_fi | Ant fisheries-related buildings | -0.0709 | 0.0036 | 0.1288 | 0.1704 |
| Gab_a | AA built-up | -0.1659 | -0.1674 | 0.4103 | 0.3957 |
| Vmag_a | AA regulated magazine | -0.0579 | 0.1950 | 0.0011 | 0.1762 |
| Vei_b_a | AA road | -0.1210 | -0.2032 | 0.3450 | 0.3783 |
| Sti_a | AA trail, path | 0.0372 | 0.0655 | -0.0788 | 0.0722 |
| Setr_s | Ant summer mountain pasture | -0.0080 | -0.0993 | 0.1418 | 0.0140 |
| Sefr_a | AA old buildings | -0.1151 | -0.2348 | 0.3849 | 0.2991 |
| Rd_anl_a | AA reindeer husbandry facilities | -0.0871 | -0.0005 | -0.1220 | -0.0369 |
| Build_a | AA built-up area | -0.1090 | -0.1535 | 0.1133 | 0.3179 |
| City_a | AA town/city area | -0.0642 | -0.0627 | -0.0032 | 0.0980 |
| Lled_a | AA power lines | -0.0988 | -0.1556 | 0.3094 | 0.4313 |
| Cul_u_s | Ant cultural heritage sites outdoors | -0.0801 | -0.0427 | 0.0071 | 0.0983 |
| Cul_t_s | Ant technical heritage sites | -0.0074 | -0.0305 | 0.1138 | 0.1248 |
| Cul_k_s | Ant church ruins | -0.1118 | -0.1645 | 0.1443 | 0.2790 |
| Cul_b_s | Ant ancient rock art sites | -0.0590 | -0.0246 | 0.0461 | 0.0938 |
| Cul_a_s | Ant archaeological heritage sites | -0.2199 | -0.1290 | 0.2048 | 0.2429 |
| Abygg_a | AA large buildings | -0.1058 | -0.1526 | 0.1125 | 0.3155 |
| Arfull_a | AA arable land | -0.1634 | -0.2688 | 0.3702 | 0.3195 |
| Arover_a | AA surface cultivated land | -0.0716 | -0.1484 | 0.2659 | 0.2513 |

**Table 64.**
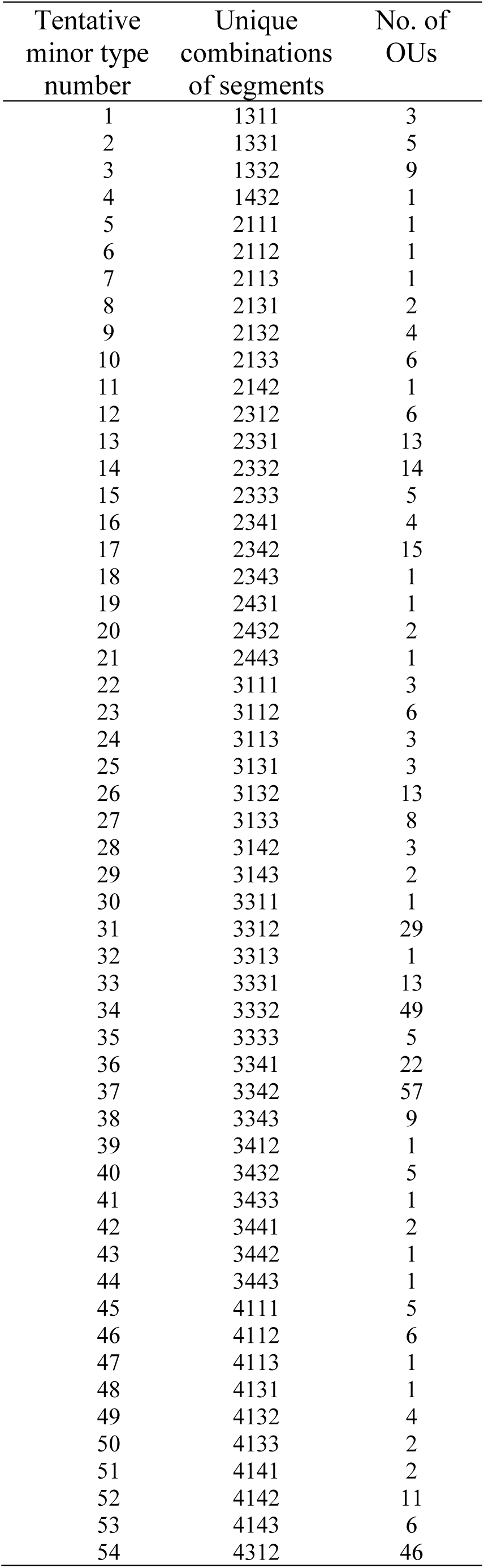
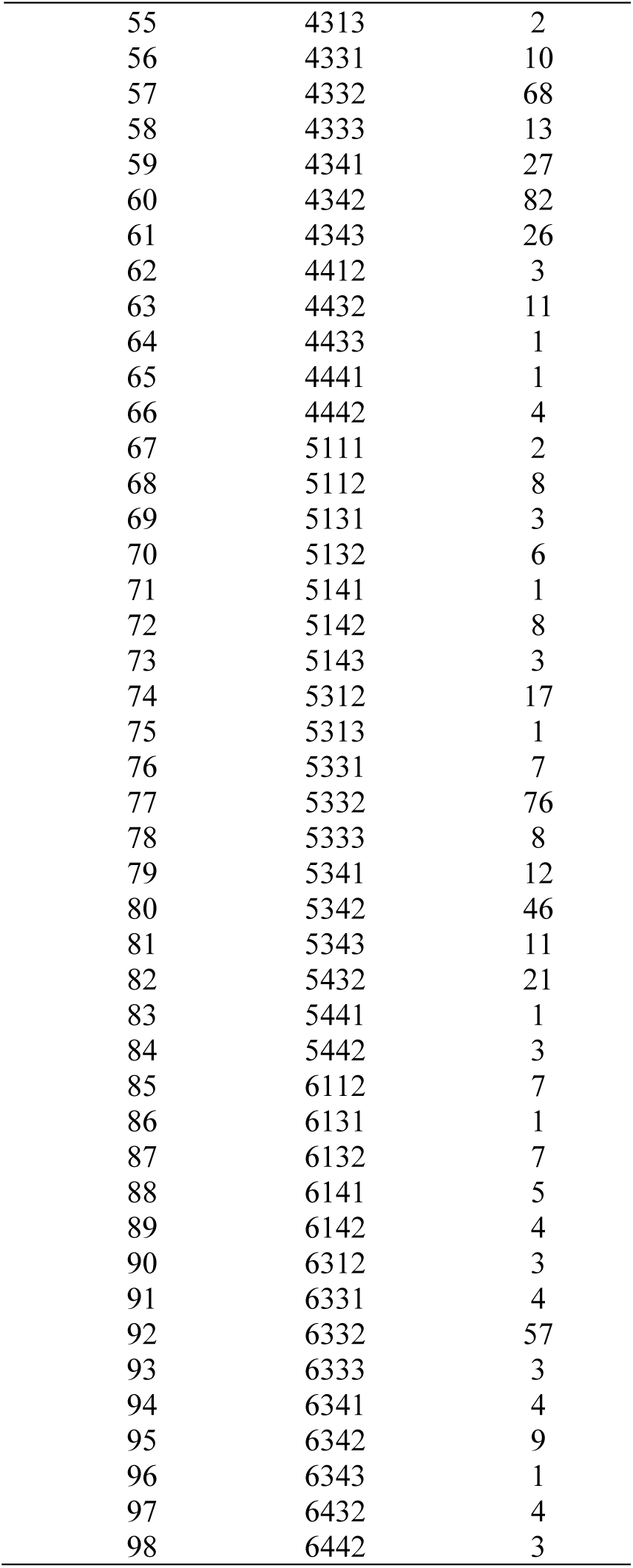
Realised gradient combinations (unique combinations of segments, i.e. tentative minor landscape types) and number of observation units for each combination in inland valleys (ID).

| Tentative<br>minor type<br>number | Unique<br>combinations<br>of segments | No. of<br>OUs |
| --- | --- | --- |
| 1 | 1311 | 3 |
| 2 | 1331 | 5 |
| 3 | 1332 | 9 |
| 4 | 1432 | 1 |
| 5 | 2111 | 1 |
| 6 | 2112 | 1 |
| 7 | 2113 | 1 |
| 8 | 2131 | 2 |
| 9 | 2132 | 4 |
| 10 | 2133 | 6 |
| 11 | 2142 | 1 |
| 12 | 2312 | 6 |
| 13 | 2331 | 13 |
| 14 | 2332 | 14 |
| 15 | 2333 | 5 |
| 16 | 2341 | 4 |
| 17 | 2342 | 15 |
| 18 | 2343 | 1 |
| 19 | 2431 | 1 |
| 20 | 2432 | 2 |
| 21 | 2443 | 1 |
| 22 | 3111 | 3 |
| 23 | 3112 | 6 |
| 24 | 3113 | 3 |
| 25 | 3131 | 3 |
| 26 | 3132 | 13 |
| 27 | 3133 | 8 |
| 28 | 3142 | 3 |
| 29 | 3143 | 2 |
| 30 | 3311 | 1 |
| 31 | 3312 | 29 |
| 32 | 3313 | 1 |
| 33 | 3331 | 13 |
| 34 | 3332 | 49 |
| 35 | 3333 | 5 |
| 36 | 3341 | 22 |
| 37 | 3342 | 57 |
| 38 | 3343 | 9 |
| 39 | 3412 | 1 |
| 40 | 3432 | 5 |
| 41 | 3433 | 1 |
| 42 | 3441 | 2 |
| 43 | 3442 | 1 |
| 44 | 3443 | 1 |
| 45 | 4111 | 5 |
| 46 | 4112 | 6 |
| 47 | 4113 | 1 |
| 48 | 4131 | 1 |
| 49 | 4132 | 4 |
| 50 | 4133 | 2 |
| 51 | 4141 | 2 |
| 52 | 4142 | 11 |
| 53 | 4143 | 6 |
| 54 | 4312 | 46 |
| 55 | 4313 | 2 |
| 56 | 4331 | 10 |
| 57 | 4332 | 68 |
| 58 | 4333 | 13 |
| 59 | 4341 | 27 |
| 60 | 4342 | 82 |
| 61 | 4343 | 26 |
| 62 | 4412 | 3 |
| 63 | 4432 | 11 |
| 64 | 4433 | 1 |
| 65 | 4441 | 1 |
| 66 | 4442 | 4 |
| 67 | 5111 | 2 |
| 68 | 5112 | 8 |
| 69 | 5131 | 3 |
| 70 | 5132 | 6 |
| 71 | 5141 | 1 |
| 72 | 5142 | 8 |
| 73 | 5143 | 3 |
| 74 | 5312 | 17 |
| 75 | 5313 | 1 |
| 76 | 5331 | 7 |
| 77 | 5332 | 76 |
| 78 | 5333 | 8 |
| 79 | 5341 | 12 |
| 80 | 5342 | 46 |
| 81 | 5343 | 11 |
| 82 | 5432 | 21 |
| 83 | 5441 | 1 |
| 84 | 5442 | 3 |
| 85 | 6112 | 7 |
| 86 | 6131 | 1 |
| 87 | 6132 | 7 |
| 88 | 6141 | 5 |
| 89 | 6142 | 4 |
| 90 | 6312 | 3 |
| 91 | 6331 | 4 |
| 92 | 6332 | 57 |
| 93 | 6333 | 3 |
| 94 | 6341 | 4 |
| 95 | 6342 | 9 |
| 96 | 6343 | 1 |
| 97 | 6432 | 4 |
| 98 | 6442 | 3 |

##### CLG 1 – (G1) Relief: RE (Table 59, Figs 138–139)

The three variables included in the ONV that represents CLG 1, relief (RE), were: (1) relief (Rr1_m); (2) mean terrain ruggedness (Rug3_m); and (3) areal coverage of moderate slope (Meant_a). This geo-ecological CLG 1, relief (RE), had an estimated gradient length of 0.5313/0.08 = 6.641 EDU–L (ecodiversity distance in landscapes) units and was, accordingly, divided into 6 standard segments (Table 59). The distribution of OUs along this ONV (in its original scaling) was almost symmetric (Fig. 138).

This CLG expresses variation in valley shape, as expressed by the depth/width ratio of the valley relative to its surroundings; from wide valleys via open valleys to narrow and steep-sided valleys. The CLG has good linearity, but the degree of generalisation is reduced towards the low-score end of the gradient, i.e. segments 1 and 2 (of 6), corresponding to relative relief RR < 100 m.

Segments 1 and 2 contained relatively few OUs compared to the other segments (Fig. 139), suggesting that these OUs are atypical for the major type. This opens for the possibility that these OUs are transitional to the major types inland plains and inland mountains. They may, however, also result from inconsistencies or uncertainties in the OU delineation process. Segments 5 and 6 are characterised by the same threshold values as landscape-gradient segments DN2/DN3 and DN3/DN4, respectively, in the ‘pilot Nordland project’ (see Appendix S1).

**Fig. 139.**
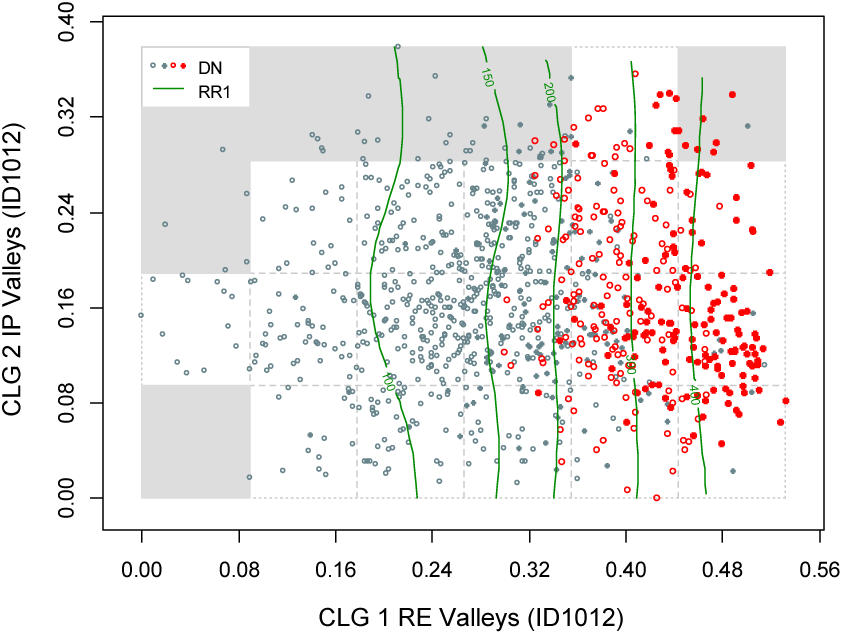
Major type inland valleys (ID): distribution of 1012 observation units (ID1012 data subset) along rescaled orthogonal key variables (rONVs) for complex landscape gradients CLG 1 (relief; RE) and CLG 2 (freshwater lake properties; IP). Symbols show affiliation of OUs to levels of relevant landscape-gradient (LG) variables, operationalised as ordered factor variables in accordance with Table 3. Isolines are given for relevant primary key variables (Table 4) and analytic variables (Table 5).

This CLG apparently captures all variation related to valley form and relief in the major type. Three key variables and 6 analysis variables had correlations with |τ| > 0.6 with the sONV. The strong signal from RE was also correlated with variation in areal coverage of: moderate slope (from flat to steep terrain; τ = –0.5420), landslide soil (τ = 0.5077), exposed bedrock (Kbf_a; τ = 0.3452) and mires (Mire_a; τ = –0. 3164). The number of strongly correlated variables and large total amount of variation captured by this CLG explains the large gradient-length estimate.

##### CLG 2 – (G2) Freshwater lake properties: IP (Table 60, Figs 140–141)

The two variables included in the ONV that represents CLG 2, freshwater lake properties (IP), were: (1) mean distance to lake (Dislake) and (2) abundance of freshwater lakes (II). This CLG had an estimated gradient length of 0.3784/0.08 = 4.73 EDU–L units and was, accordingly, divided into 4 standard segments (Table 60). The distribution of OUs along this ONV (in its original scaling) was left-skewed (Fig. 140). The CLG has a good linearity and reduced degree of generalization towards gradient endpoints.

**Fig. 140.**
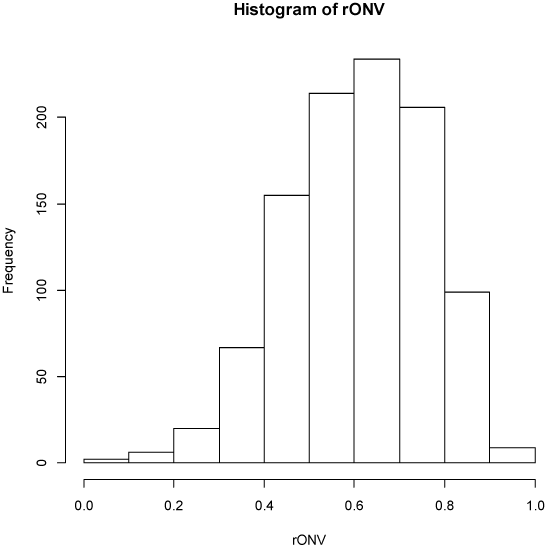
Inland valleys (ID): candidate complex-landscape gradient (CLG) 2, abundance of lakes (VP): distribution of OUs on the ranged but not rescaled orthogonal key variable (ONV).

**Fig. 141.**
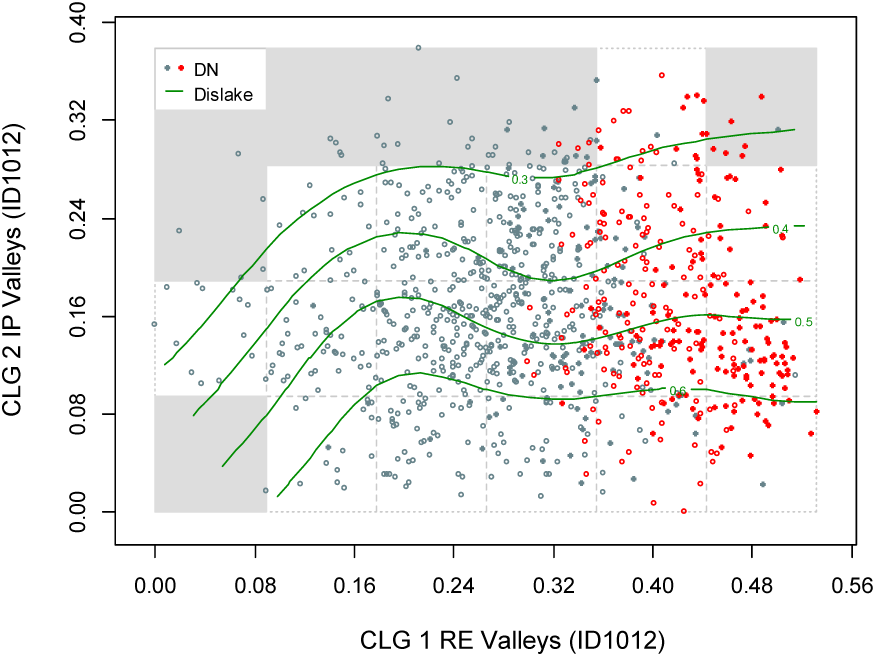
Major type inland valleys (ID): distribution of 1012 observation units (ID1012 data subset) along rescaled orthogonal key variables (rONVs) for complex landscape gradients CLG 1 (relief; RE) and CLG 2 (freshwater lake properties; IP). Symbols show affiliation of OUs to levels of relevant landscape-gradient (LG) variables, operationalised as ordered factor variables in accordance with Table 3. Isolines are given for relevant primary key variables (Table 4) and analytic variables (Table 5).

This CLG expresses variation within valleys from landscapes without lakes or with small lakes to valley landscapes with large lakes; the typical ‘inland fjords’. Four analytical variables related to hydrology were strongly (|τ| > 0.3) correlated with this CLG (Table 63): areal coverage of freshwater lake (Lake_a; τ = 0.5429); number of freshwater lake islands (Innoy_s; τ = 0.3714); areal coverage of river (R_net_a; τ = –0.2325); and number of lakes (Inns_s; τ = 0.3561). The only variables not directly related to lake occurrence that were correlated with this CLG were areal coverage of glaciofluvial deposits (Kelv_a; τ = –0.3023) and the mean of the hypsographic index, ER (Er_m; τ = –0.3584).

##### CLG 3 – (U2) Forest cover: SkP (Table 61, Figs 142–145)

The two variables included in the ONV that represents CLG 3, forest cover (SkP), were: (1) areal coverage of impediment (i.e. areas not suitable for agriculture or forestry; Sn_imp) and (2) areal coverage of coniferous forest (Arbar_a). This bio-ecological CLG had an estimated gradient length of 0.3285/0.08 = 4.106 EDU–L units and was, accordingly, divided into 4 standard segments (Table 61). The distribution of OUs along this ONV (in its original scaling) was bimodal and left-skewed (Fig. 142). The linearity was low. The noteworthy ‘gap’ in landscape-property composition in the middle segments, giving rise to the bimodal frequency distribution, was probably due to a clear-cut differentiation between mountains and forested lowlands.

**Fig. 142.**
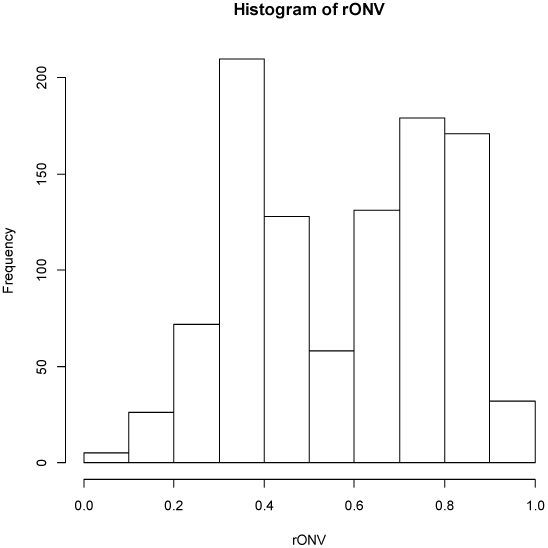
Inland valleys (ID): candidate complex-landscape gradient (CLG) 3, forest cover (SkP): distribution of OUs on the ranged but not rescaled orthogonal key variable (ONV).

**Fig. 143–145.**
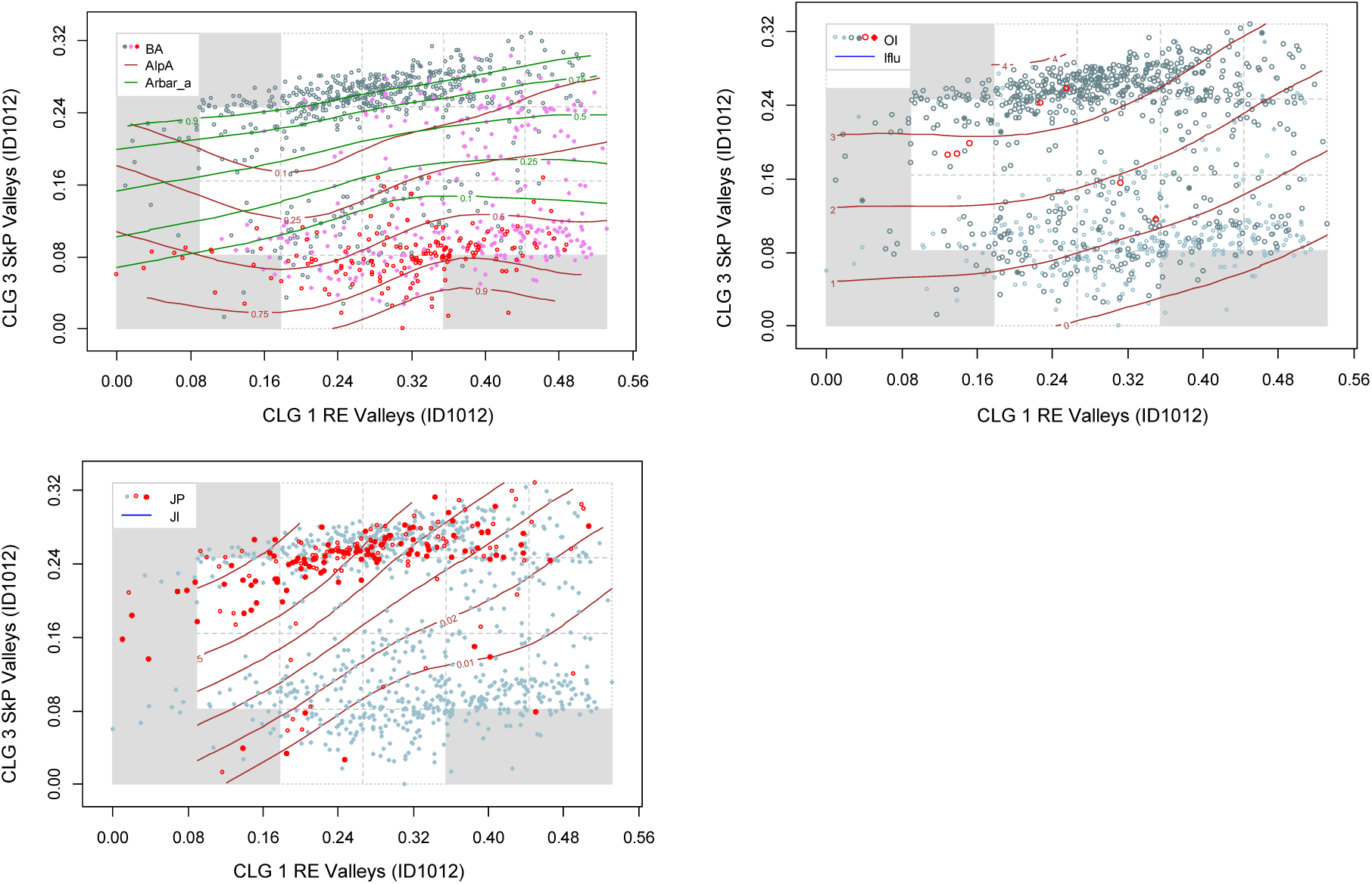
Major type inland valleys (ID): distribution of 1012 observation units (ID1012 data subset) along rescaled orthogonal key variables (rONVs) for complex landscape gradients CLG 1 (relief; RE) and CLG 3 (forest cover; SkP). Symbols show affiliation of OUs to levels of relevant landscape-gradient (LG) variables, operationalised as ordered factor variables in accordance with Table 3. Isolines are given for relevant primary key variables (Table 4) and analytic variables (Table 5).

Four vegetation variables that express vegetation cover were correlated with this CLG (Table 63): the areal coverage of mixed boreal forest (Arbla_a; τ = 0.5467); dry heath/open areas (Sn_torr; τ = –0.3933); patchy open treeless areas (Sn_flekk; τ = –0.4291); and boreal/alpine landscapes (AlpA; τ = –0.5309). In addition, this CLG captured most of the variation in infrastructure and agricultural land-use intensity (Fig. 144–145), with nine land use variables correlated at |τ| > 0.3 with the CLG (Table 63). Fig. 144 indicates that agricultural land-use intensity was non-linearly related to this CLG.

##### CLG 4 – (A3) Amount of infrastructure: OI (Table 62, Figs 146–148)

The two variables included in the ONV that represents CLG 4, amount of infrastructure, were: (1) the infrastructure index (IfI) and (2) areal coverage of built-up areas (Gab_a). This land-use related CLG had an estimated gradient length of 0.2626/0.08 = 3.282 EDU–L units and was, accordingly, divided into 3 standard segments (Table 62). The distribution of OUs along this ONV (in its original scaling) was right-skewed (Fig. 146). Non-linearity occurred towards the high-score end of the gradient; the landscape element composition of segment 8 was more similar to that of segments 6 than with that of segment 7.

**Fig. 146.**
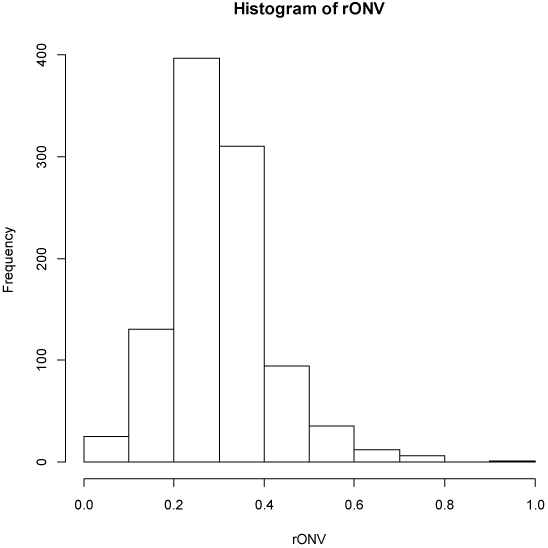
Inland valleys (ID): candidate complex-landscape gradient (CLG) 4, land use intensity (OI): distribution of OUs on the ranged but not rescaled orthogonal key variable (ONV).

**Fig. 147–148.**
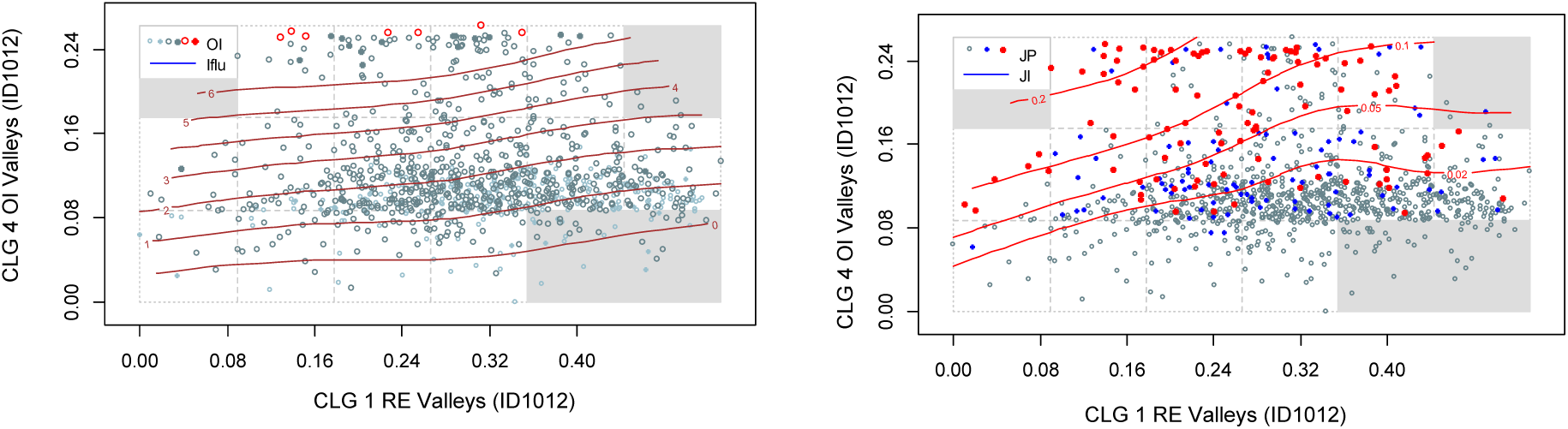
Major type inland valleys (ID): distribution of 1012 observation units (ID1012 data subset) along rescaled orthogonal key variables (rONVs) for complex landscape gradients CLG 1 (relief; RE) and CLG 4 (amount of infrastructure; OI). Symbols show affiliation of OUs to levels of relevant landscape-gradient (LG) variables, operationalised as ordered factor variables in accordance with Table 3. Isolines are given for relevant primary key variables (Table 4) and analytic variables (Table 5).

This CLG expresses residual variation in human land-use not captured by the other gradients and not related to the gradient from lowlands to mountains (CLG 3, SkP). This pattern clearly emerges from the six analytical human land-use related variables with |τ| > 0.3 with this CLG (Table 63), which include buildings, built-up areas and other technical infrastructure (roads, power lines), and by the weak correlation of the CLG with the boreal-alpine variable (AlpA; τ = –0.0531).

##### Tentative division of inland valleys into minor types

Correlations between ONVs and ordination axes, primary key variables, and analysis variables were calculated as a basis for operationalisation of the CLGs (Table 63). The theoretical number of combinations of intervals along gradients is 6 × 4 × 4 × 3 = 288. Of these, 98 were represented in the total data set (Table 64), 24 of which with > 11 OUs (> 1% of the ID1012 data subset).

#### Inland hills and mountains (IA)

The analyses for identification of independent significant analytic variables for each CLG candidate by forward selection from the *n* = 66 variables using RDA show that five of the eight CLG candidates satisfied the requirement for explaining at least 6% of the variation (Table 65). The parsimonious set of CLGs, found by RDA with forward selection among the CLG candidates separately for each functional variable category, contained 3 CLGs (Table 65). The properties of the five CLGs and their associated (ranked primary) ONVs are described below on the basis of Tables 66–68 and Figs 149–156.

**Fig. 149.**
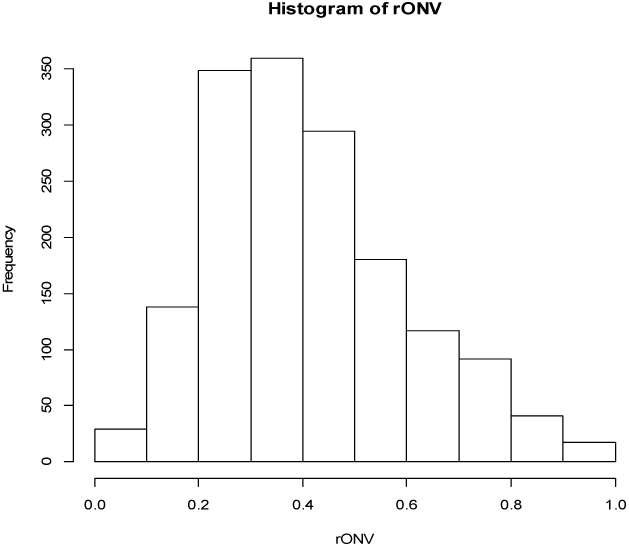
Inland hills and mountains (IA): candidate complex-landscape gradient (CLG) 1, relief (RE): distribution of OUs on the ranged but not rescaled orthogonal key variable (ONV).

**Table 65.**
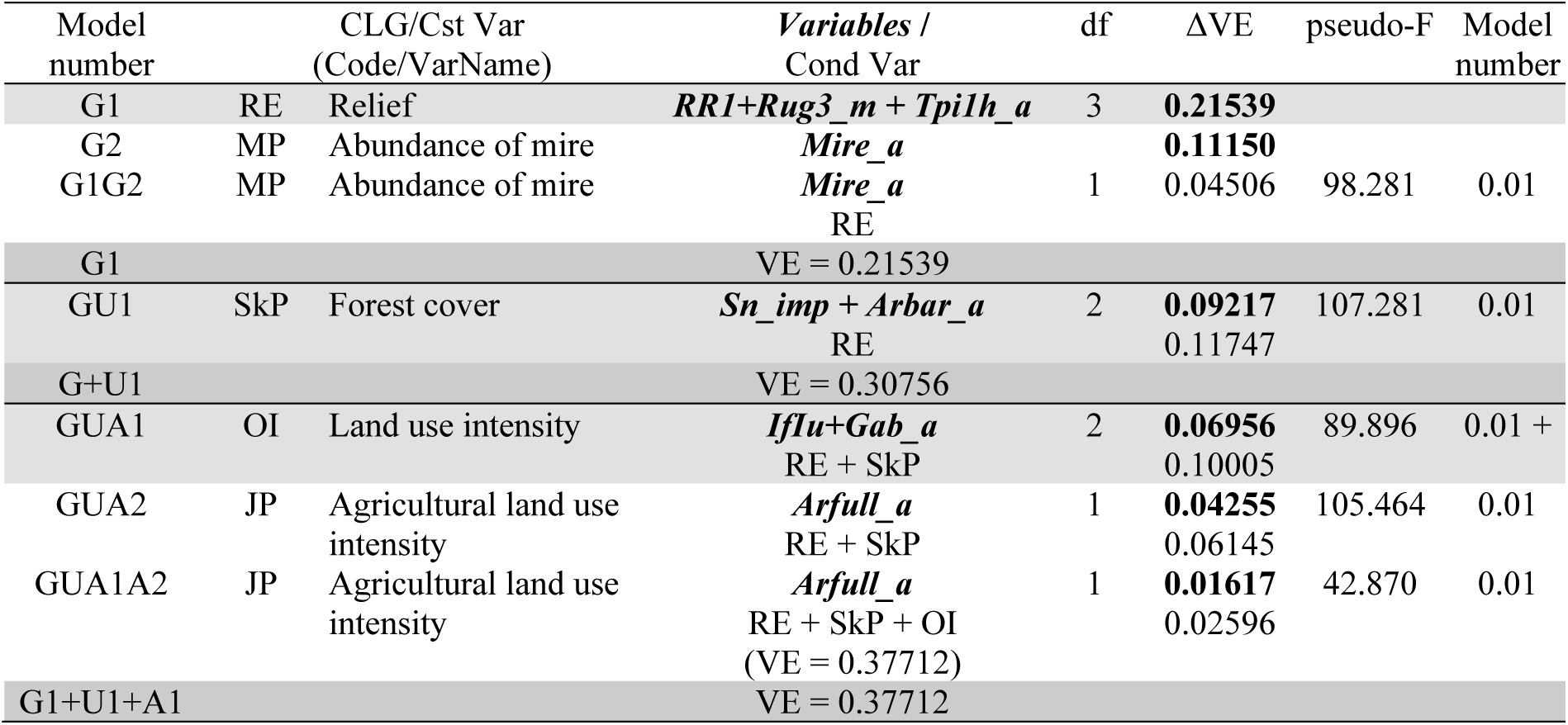
Identification of CLG candidates in inland hills and mountains by forward selection of variables using RDA. Results of tests of single CLG candidates are shown in bold-face types. The best model at each step in the selection process is highlighted using light grey shading while the best model after testing all CLG candidates in a functional variable category is highlighted using darker grey shading. Abbreviations and explanations: Var = variable; Cst Var = constraining variable; Cond Var = conditioning variable(s); df = degrees of freedom; VE = variation explained (expressed as fraction of the total variation); ΔVE = additional variation explained; pseudo-F and P = F-statistic used in the test of the null hypothesis that adding the variable(s) do not contribute more to explaining variation in landscape-element composition than a random variable, with associated P value. Variable names are abbreviated according to Tables 4 and 5. For further explanation, se Material and methods chapter.

**Table 66.** Major type inland hills- and mountains (IA): key properties* and segmentation** of candidate complex-landscape gradient G1, Relief (RE), as represented by its orthogonal key variable (ONV). OU = observation unit; Threshold = upper limit of class (RSC = rescaling). PD unit = Proportional dissimilarity unit. Segment threshold shows position of standard segment borders relative to initial segments.

|  | Initial CLG segment |  |  |  |  |  |  |  |
| --- | --- | --- | --- | --- | --- | --- | --- | --- |
|  | 1 | 2 | 3 | 4 | 5 | 6 | 7 | 8 |
| Number of OUs | 42 | 275 | 447 | 391 | 217 | 137 | 84 | 25 |
| ONV threshold before RSC | 0.1212 | 0.2465 | 0.3693 | 0.4931 | 0.6165 | 0.7394 | 0.8729 | 1 |
| ONV threshold after RSC | 0.1398 | 0.2732 | 0.4413 | 0.6236 | 0.7611 | 0.8710 | 0.9472 | 1 |
| ONV threshold, in PD units | 0.0776 | 0.1516 | 0.2449 | 0.3461 | 0.4224 | 0.4834 | 0.5257 | 0.5550 |
| Segment threshold |  | 12 | 23 | 34 | 45 | 56 |  |  |

\*Variation explained by variable group: 0.1823, PNV: 0.1584, ONV: 0.1584
Gradient length estimates: Method I: 0.5148, Method II: 0.5655, Method III: 0.5550
\*\* Tentative number of standard segments = 6
Number of OUs per standard segment: 100 – 376 – 411 – 332 – 238 – 156

**Table 67.**
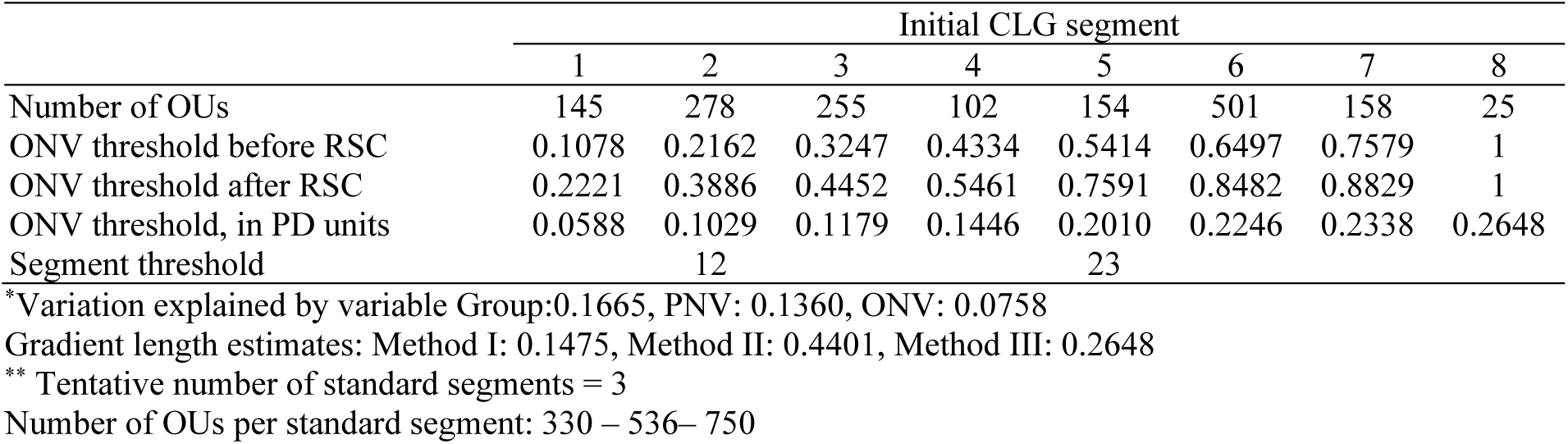
Major type inland hills and mountains (IA): key properties* and segmentation** of candidate complex-landscape gradient U1, forest cover (SkP), as represented by its orthogonal key variable (ONV). OU = observation unit; Threshold = upper limit of class (RSC = rescaling). PD unit = Proportional dissimilarity unit. Segment threshold shows position of standard segment borders relative to initial segments.

**Table 68.**
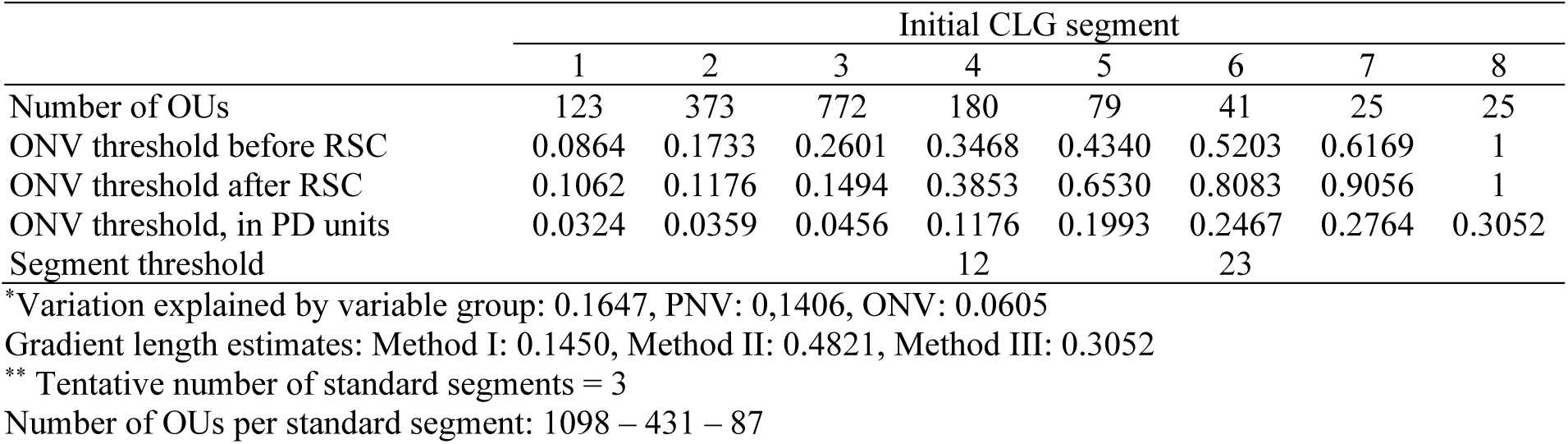
Major type inland hills and mountains (IA): key properties* and segmentation** of candidate complex-landscape gradient A1, land use intensity (OI), as represented by its orthogonal key variable (ONV). OU = observation unit; Threshold = upper limit of class (RSC = rescaling). PD unit = Proportional dissimilarity unit. Segment threshold shows position of standard segment borders relative to initial segments.

##### CLG 1 – (G1) Relief: RE (Table 66, Figs 149–150)

The three variables included in the ONV that represents CLG 1, relief (RE), were: (1) relief (RR1); (2) mean terrain ruggedness (Rug3_m); and (3) areal coverage of convex terrain (Tpi1h_a). This geo-ecological CLG had an estimated gradient length of 0.5550/0.08 = 6.938 EDU–L units and was, accordingly, divided into 6 standard segments (Table 66). The distribution of OUs along this ONV (in its original scaling) was right-skewed (Fig. 149).

This CLG expresses variation in terrain shape within inland hill and mountain landscapes; from depressions in hills and mountains, via undulating inland hills and mountains with low relief to steep and rugged hills and mountains (typical ‘alpine landscapes’). The CLG has good linearity, but weaker degree of generalisation at the lower end of the gradient, i.e. segment 1, which is characterised by relative relief RR < 100 m. The relatively few OUs in segment 1 (Fig. 150) may suggest that this segment represents a transition towards the major type inland plains.

**Fig. 150.**
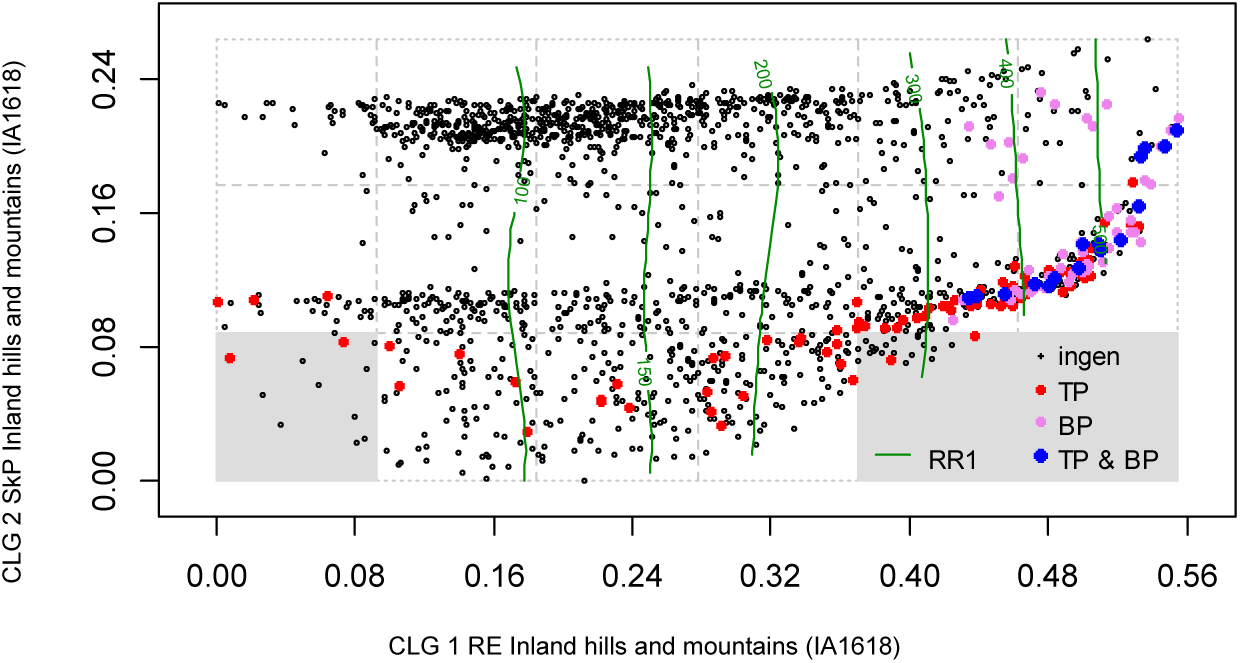
Major type inland hills and mountains (IA): distribution of 1618 observation units (IA1618 data subset) along rescaled orthogonal key variables (rONVs) for complex landscape gradients CLG 1 (relief; RE) and CLG 2 (forest cover; SkP). Symbols show affiliation of OUs to levels of relevant landscape-gradient (LG) variables, operationalised as ordered factor variables in accordance with Table 3. Isolines are given for relevant primary key variables (Table 4) and analytic variables (Table 5).

A total of 11 variables had |τ| > 0.6 with sONV_RE; the strongest correlation was obtained for relief (Rr1_m; τ = 0.8606; Table 69). This was the strongest correlation observed for a combination of a CLG and a variable in our material, showing that relief controls most of the variation in landscape properties in inland hills and mountains. The fundamental role of relief in this major type is further substantiated by the large number of strongly correlated variables, the high total amount of variation captured by this CLG and its large estimated gradient length. Mire abundance (MI; τ = –0.3778) was negatively correlated with this CLG, following the same pattern as relief. Neither the amount of infrastructure nor the gradient from lowland to mountain follow the relief gradient (Table 69; Fig. 156). Glaciers show clear concentration to the steepest terrain (BP; Fig. 150), almost completely confined to segments 5 and 6. Exceptions do, however, occur. The analyses did not identify presence of glaciers as a CLG on its own, most probably because of the few glacier-related variables included in our data set. Landscapes with glaciers and/or sharp peaks/pinnacles had a high relative relief within short distances (TP; Fig. 150) and RR > 400 m. OUs with these characteristics were, however, not clearly separated from other OUs with RR > 400 and hence do not quality for separation as a CLG on its own. This CLG has noticeable correlations with variables related to presence or absence of vegetation cover, i.e. expressing areal coverage of impediment (Sn_imp; τ = 0.4109); mire (Mire_a; τ = –0.3856); exposed bedrock (Kbf_a; τ = 0.3272) and patchy open treeless area (Sn_flekk; τ = 0.3105). Variables related to vegetation cover were, however, more strongly correlated with CLG 2, SkP (Table 69).

**Table 69.** Correlations between the three ONVs and ordination axes (rows 1–3), primary key variables (rows 4–14), and analysis variables in major landscape type inland hills and mountains (IA).

| Variable |  | sONV RE | sONV SkP | sONV OI |
| --- | --- | --- | --- | --- |
| GNMDS 1 |  | −0.4397 | 0.4438 | −0.0256 |
| GNMDS 2 |  | −0.4717 | −0.2288 | −0.2601 |
| GNMDS3 |  | −0.1063 | −0.0361 | −0.0623 |
| DNI | Valley shape | −0.0405 | 0.0361 | 0.0338 |
| RR1 | Relief | 0.8606 | 0.0150 | 0.0795 |
| FIA | Amount of flat land | −0.6991 | −0.0180 | −0.0535 |
| TPI1 | Peaks | 0.7660 | −0.0231 | 0.0912 |
| BrI | Glacier | 0.2456 | −0.1817 | 0.0779 |
| AlpA | Boreal/alpine landscape | 0.2582 | −0.4916 | 0.0559 |
| II | Abundance of freshwater lakes | −0.1019 | −0.0997 | −0.0407 |
| MI | Abundance of mire | −0.3778 | 0.3205 | −0.1864 |
| IfI | Amount of infrastructure (OI) | −0.1976 | 0.4211 | 0.2296 |
| IfIu | Amount of infrastructure | −0.2037 | 0.3998 | 0.2499 |
| JI | Agricultural land use intensity | −0.1582 | 0.3199 | 0.3085 |
| Flat_a | AA flat terrain | −0.6961 | −0.0180 | −0.0536 |
| Ktkmo_a | AA thick layer of till | −0.2120 | 0.0536 | 0.0196 |
| Tpi1l_a | AA depressions | 0.3508 | −0.0173 | 0.0188 |
| Tpi1h_a | AA convex terrain | 0.7973 | −0.0096 | 0.0946 |
| Tpi1_mp | Terrain form TPI1, numerical, mean | 0.7517 | −0.0227 | 0.0892 |
| River_a | AA large river | −0.1926 | −0.0610 | 0.0238 |
| Steep_a | AA steep terrain/slope | 0.8435 | 0.0185 | 0.0817 |
| Kskred_a | AA landslide soil | 0.5056 | −0.0536 | 0.0901 |
| Rug3_m | Terrain ruggedness VRM3, mean | 0.7172 | −0.0186 | 0.0695 |
| Rr1_m | Altitudinal range | 0.8606 | 0.0150 | 0.0795 |
| Brich_a | AA lime-rich bedrock | −0.0210 | 0.0516 | 0.0361 |
| Bpoor_a | AA lime-poor bedrock geology | 0.0506 | 0.0751 | −0.0111 |
| Bplu_a | AA plutonic rock | 0.1624 | −0.0313 | −0.0026 |
| Bvul_a | AA volcanic rock | −0.1094 | 0.0832 | 0.0081 |
| Bavstn_a | AA sedimentary rock | −0.2504 | 0.1018 | −0.0137 |
| Bomd_a | AA metamorphic rock | 0.0289 | −0.0299 | 0.0151 |
| Mire_a | AA mire | −0.3856 | 0.3116 | −0.1821 |
| Glac_a | AA glacier | 0.2493 | −0.1935 | 0.0859 |
| Meant_a | AA moderate slope | −0.6161 | 0.0156 | −0.0934 |
| Kmar_a | AA marine deposits | −0.0919 | 0.1692 | 0.1410 |
| Klac_a | AA lacustrine deposits | −0.0551 | 0.0286 | 0.0708 |
| Kelv_a | AA glaciofluvial deposits | −0.2236 | 0.0773 | 0.0682 |
| Kbf_a | AA exposed bedrock | 0.3272 | −0.1260 | −0.0379 |
| Lake_a | AA freshwater lake | −0.1599 | −0.1213 | −0.1225 |
| Innoy_s | Ant freshwater lake islands | −0.1367 | −0.0380 | −0.1096 |
| Er_m | Hypsographic index, ER, mean | 0.2598 | −0.0309 | 0.0506 |
| R_net_a | AA river | −0.2160 | 0.0956 | −0.1232 |
| Inns_s | Number of lakes | −0.0240 | −0.1599 | −0.2405 |
| Asp_s_a | AA south facing terrain | 0.0274 | −0.0370 | 0.0135 |
| Asp_n_a | AA north facing terrain | 0.0873 | −0.0536 | −0.0510 |
| Dismire | Distance to mire, mean | 0.3454 | −0.3295 | 0.1785 |
| Dislake | Distance to lake, mean | 0.1231 | 0.1404 | 0.1659 |
| Discoast | Distance to coast, mean | −0.2045 | 0.0024 | −0.1427 |
| Guro_t_a | AA rugged terrain | 0.7347 | −0.0665 | 0.0783 |
| Tpi6l_a | AA rugged terrain, TPI6 | −0.3436 | −0.0010 | −0.0515 |
| Oyst_i | Inverse island size | 0.1747 | −0.0009 | 0.0449 |
| Bohei_a | AA boreal heaths | 0.1200 | 0.0342 | −0.0799 |
| Araaf_a | AA open areas | 0.1174 | 0.1222 | −0.0517 |
| Arbar_a | AA coniferous forest | −0.2314 | 0.6676 | −0.1161 |
| Arbla_a | AA mixed boreal forest | −0.1723 | 0.5178 | −0.1075 |
| Arlov_a | AA deciduous forest | 0.0471 | 0.2837 | 0.0606 |
| Sn_imp | AA impediment | 0.4109 | −0.4512 | 0.1436 |
| Sn_flekk | AA patchy open treeless area | 0.3105 | -0.4137 | 0.0117 |
| Sn_lav | AA lichen heath | -0.0992 | -0.0575 | -0.0291 |
| Sn_torr | AA dry heath/open areas | 0.1317 | -0.2787 | -0.1185 |
| Sn_frisk | AA moist/fresh heath/open areas | 0.1668 | -0.1773 | -0.0583 |
| Sn_ureg | AA unregistered heath/open areas | -0.0033 | -0.0540 | 0.0983 |
| Gab_nae | Ant commercial buildings | -0.1737 | 0.2130 | 0.3436 |
| Gab_fi | Ant fisheries-related buildings | -0.0402 | 0.0571 | 0.0632 |
| Gab_a | AA built-up | -0.2384 | 0.4212 | 0.2170 |
| Vmag_a | AA regulated magazine | -0.0610 | -0.0065 | 0.0259 |
| Vei_b_a | AA road | -0.1573 | 0.2468 | 0.3127 |
| Sti_a | AA trail, path | -0.0714 | -0.0002 | 0.0450 |
| Setr_s | Ant summer mountain pasture | -0.0715 | 0.3152 | 0.0191 |
| Sefr_a | AA old buildings | -0.2135 | 0.3731 | 0.2560 |
| Rd_anl_a | AA reindeer husbandry facilities | -0.0419 | -0.1328 | -0.0142 |
| Build_a | AA built-up area | -0.1384 | 0.1080 | 0.2903 |
| Lled_a | AA power lines | -0.1242 | 0.2713 | 0.3084 |
| Cul_u_s | Ant cultural heritage sites outdoors | -0.0725 | 0.0431 | 0.1173 |
| Cul_t_s | Ant technical heritage sites | -0.1133 | 0.0909 | 0.1341 |
| Cul_k_s | Ant church ruins | -0.1038 | 0.1258 | 0.2054 |
| Cul_b_s | Ant ancient rock art sites | -0.0606 | 0.0620 | 0.1524 |
| Cul_a_s | Ant archaeological heritage sites | -0.1710 | 0.1839 | 0.2620 |
| Abygg_a | AA large buildings | -0.1361 | 0.1067 | 0.2921 |
| Arfull_a | AA arable land | -0.1847 | 0.3496 | 0.2695 |
| Arover_a | AA surface cultivated land | -0.1125 | 0.2272 | 0.2231 |

##### CLG 2 – (U1) Forest cover: SkP (Table 67, Figs 151–153)

The two variables included in the ONV that represents CLG 2, forest cover (SkP), were: (1) areal coverage of impediment (i.e. areas not suitable for agriculture or forestry; Sn_imp) and (2) areal coverage of coniferous forest (Arbar_a). This bio-ecological CLG had an estimated gradient length of 0.2648/0.08 = 3.31 EDU–L units and was, accordingly, divided into 3 standard segments (Table 67). The distribution of OUs along this ONV (in its original scaling) was bimodal and right-skewed (Fig. 151).

**Fig. 151.**
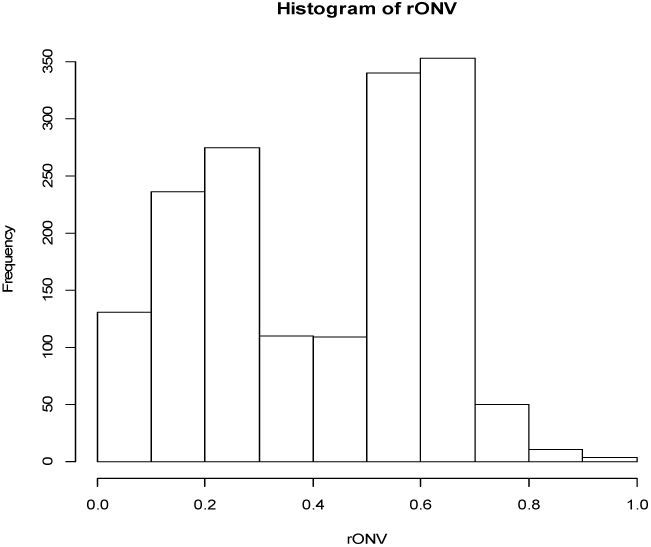
Inland hills and mountains (IA): candidate complex-landscape gradient (CLG) 2, forest cover (SkP): distribution of OUs on the ranged but not rescaled orthogonal key variable (ONV).

This CLG expresses variation in vegetation cover from barren mountains, without or with sparse vegetation cover, to forested or potentially forested areas below the climatic forest line. The intermediate segments include open mountain heaths and boreal heath-dominated areas below the climatic forest line, kept open due to historical land-use (e.g. logging and grazing). Fig. 152 indicates that the bimodal distribution of OUs along this CLG is due to differences in landscape-element composition between mountains and other areas.

**Fig. 152–153.**
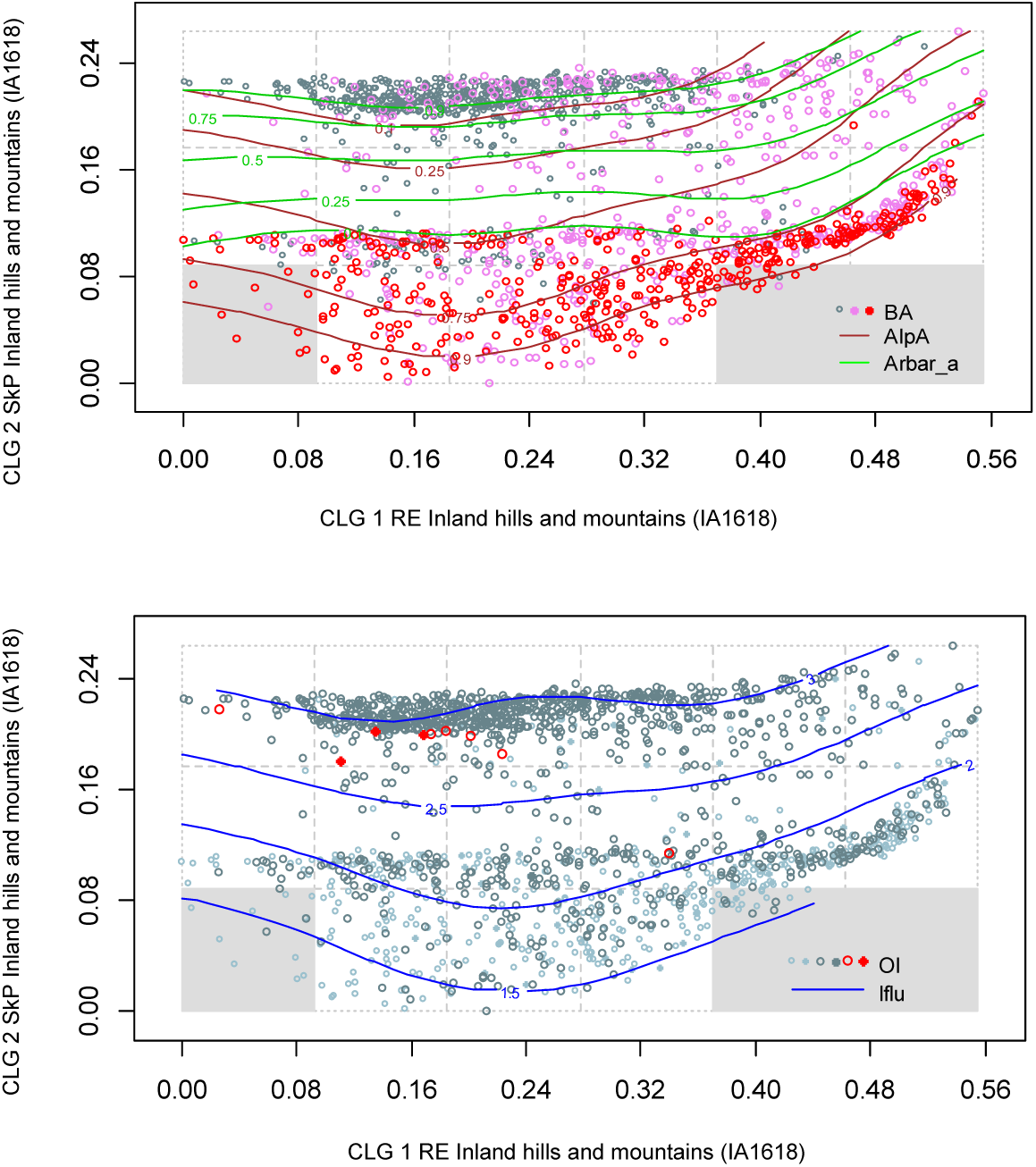
Major type inland hills and mountains (IA): distribution of 1618 observation units (IA1618 data subset) along rescaled orthogonal key variables (rONVs) for complex landscape gradients CLG 1 (relief; RE) and CLG 2 (forest cover; SkP). Symbols show affiliation of OUs to levels of relevant landscape-gradient (LG) variables, operationalised as ordered factor variables in accordance with Table 3. Isolines are given for relevant primary key variables (Table 4) and analytic variables (Table 5).

Correlated bio-ecological variables were (Table 69) the areal coverage of: coniferous forest (Arbar_a; τ = 0.6676); mixed boreal forest (Arbla_a; τ = 0.5178); mean distance to mire (Dismire; τ = –0.3295); patchy open treeless area (Sn_flekk; τ = –0.4137); impediment (Sn_imp; τ = –0.4512); and amount/degree of boreal/alpine landscapes (AlpA; τ = –0.4916). The gradient also expresses variation in human land-use along the gradient from the lowlands to the mountains, as indicated by correlations between the CLG and agricultural land-use intensity (JI; τ = 0.3199) and the number of summer mountain pastures (Setr_s; τ = 0.3152).

##### CLG 3 – (A1) Amount of infrastructure: OI (Table 68, Figs 154–156)

The two variables included in the ONV that represents CLG were: (1) the infrastructure index (IfIu) and (2) areal coverage of built-up areas (Gab_a). This land-use related CLG had an estimated gradient length of 0.3052/0.08 = 3.815 EDU–L units and was, accordingly, divided into 3 standard segments (Table 68). The distribution of OUs along this ONV (in its original scaling) was strongly right-skewed (Fig. 154).

**Fig. 154.**
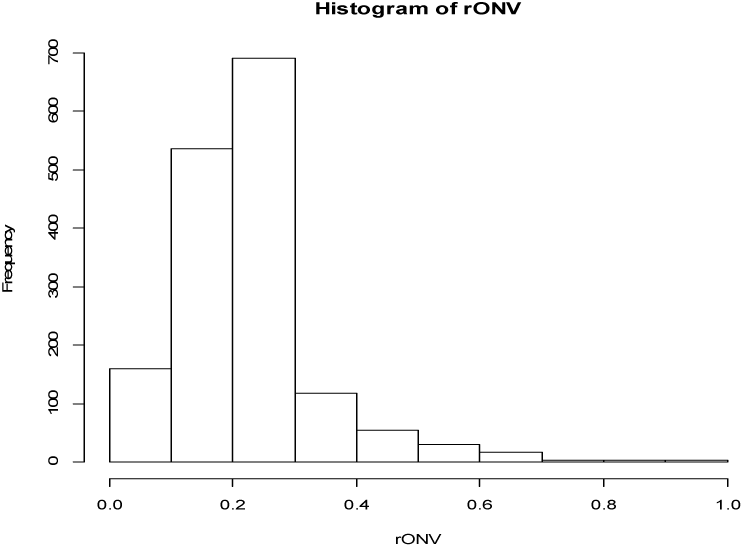
Inland hills and mountains (IA): candidate complex-landscape gradient (CLG) 3, land use intensity (OI): distribution of OUs on the ranged but not rescaled orthogonal key variable (ONV).

This CLG expresses residual variation in the importance of human infrastructure and to some extent also agricultural land-use from low via intermediate to high. The OUs were clustered into three groups, roughly corresponding to the standard segments, on the basis of landscape property composition. The majority of OUs had low amount of infrastructure (Figs 155–156). Except for high scores along this CLG for OUs with large values for the infrastructure index (Fig. 156: red symbols for high density of infrastructure, i.e. cities and towns), this CLG does not clearly distinguish between OUs on the basis of explicit analysis variables, or variable combinations. Table 69 shows that only four variables had noticeable correlations with this CLG: number of commercial buildings (Gab_nae; τ = 0.3436); areal coverage of roads (Vei_b_a; τ = 0.3127), power lines (Lled_a; τ = 0.3084) and agricultural land-use intensity (JI; τ = 0.3085).

**Fig. 155–156.**
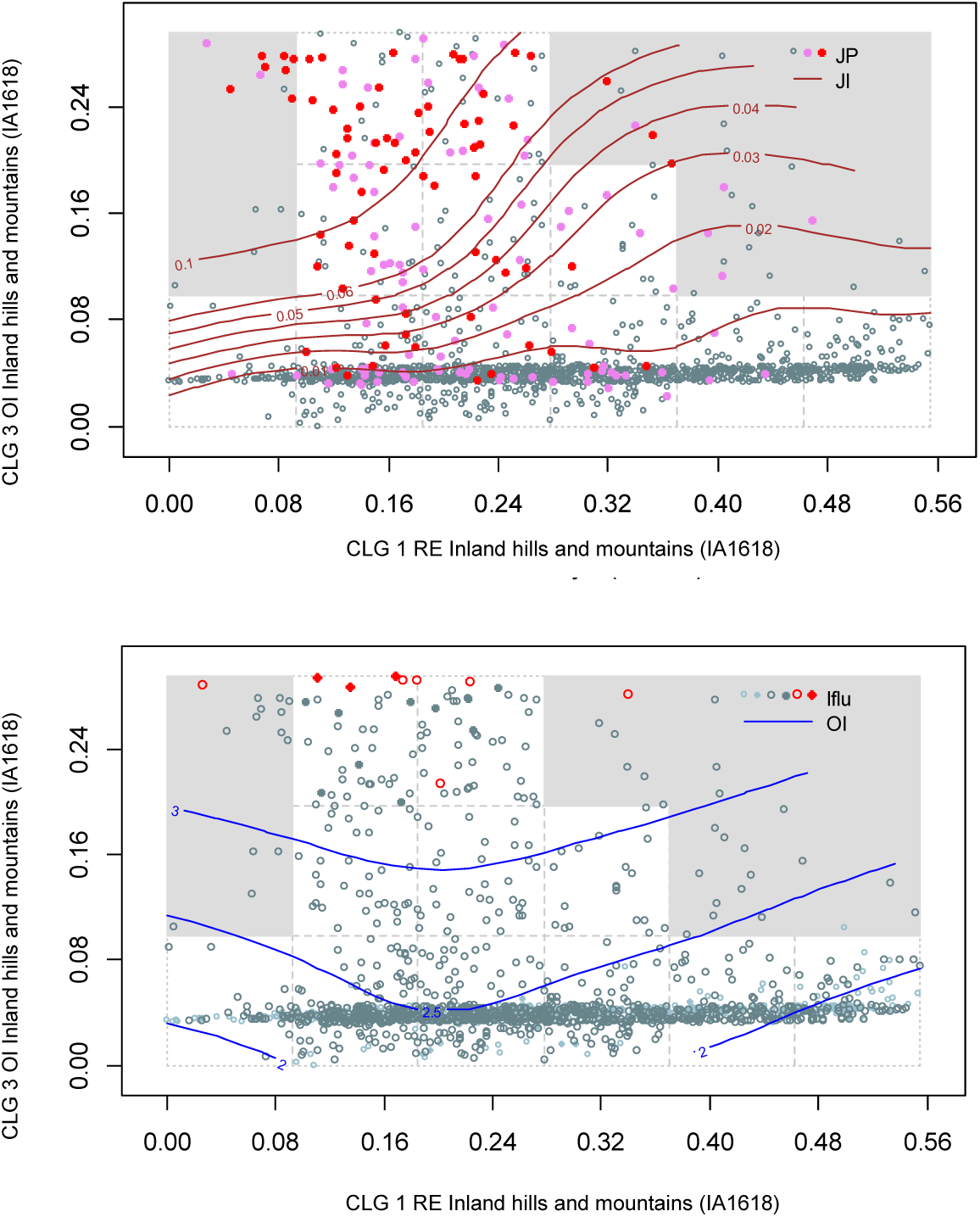
Major type inland hills and mountains (IA): distribution of 1618 observation units (IA1618 data subset) along rescaled orthogonal key variables (rONVs) for complex landscape gradients CLG 1 (relief; RE) and CLG 3 (amount of infrastructure; OI). Symbols show affiliation of OUs to levels of relevant landscape-gradient (LG) variables, operationalised as ordered factor variables in accordance with Table 3. Isolines are given for relevant primary key variables (Table 4) and analytic variables (Table 5).

##### Tentative division of inland hills and mountains into minor types

Correlations between ONVs and ordination axes, primary key variables, and analysis variables were calculated as a basis for operationalisation of the CLGs (Table 69). The theoretical number of combinations of intervals along gradients is 6 × 3 × 3 = 54. Of these, 41 were represented in the total data set (Table 70), 21 of which with 17 OUs or more (i.e. more than 1% of the OUs in the IA1618 dataset; see Table 70).

**Table 70.** Realised gradient combinations (unique combinations of segments, i.e. tentative minor landscape types) and number of observation units for each combination in inland hills and mountains (KS).

| Tentative<br>minor type<br>number | Unique<br>combinations<br>of segments | No. of<br>OUs |
| --- | --- | --- |
| 1 | 111 | 12 |
| 2 | 121 | 24 |
| 3 | 122 | 2 |
| 4 | 123 | 2 |
| 5 | 131 | 19 |
| 6 | 132 | 2 |
| 7 | 133 | 8 |
| 8 | 211 | 75 |
| 9 | 212 | 1 |
| 10 | 221 | 95 |
| 11 | 222 | 6 |
| 12 | 223 | 1 |
| 13 | 231 | 176 |
| 14 | 232 | 28 |
| 15 | 233 | 34 |
| 16 | 311 | 88 |
| 17 | 312 | 4 |
| 18 | 321 | 77 |
| 19 | 322 | 2 |
| 20 | 323 | 3 |
| 21 | 331 | 188 |
| 22 | 332 | 24 |
| 23 | 333 | 37 |
| 24 | 411 | 103 |
| 25 | 412 | 1 |
| 26 | 421 | 66 |
| 27 | 422 | 5 |
| 28 | 423 | 4 |
| 29 | 431 | 131 |
| 30 | 432 | 13 |
| 31 | 433 | 2 |
| 32 | 511 | 20 |
| 33 | 521 | 133 |
| 34 | 522 | 7 |
| 35 | 523 | 3 |
| 36 | 531 | 65 |
| 37 | 532 | 4 |
| 38 | 621 | 110 |
| 39 | 622 | 1 |
| 40 | 631 | 39 |
| 41 | 632 | 3 |

## DISCUSSION

### MAJOR CATEGORIES OF LANDSCAPES

#### The division into coastal and inland landscapes

The analyses reveal very clear patterns of co-ordinated variation in landscape properties and landscape-element composition in Norway, providing both general knowledge about the distribution of landscape elements throughout the country and an empirical basis for the first version of a landscape-type system. We find that the single most important property explaining differences in the composition of landscape elements and features in Norway is the location of the landscape in relation to the coastline. This finding supports construction of a landscape-type hierarchy according to EcoSyst principles (Halvorsen et al. 2020), which is performed by a top-down process, by separating coastal and inland landscapes at the topmost, level of major-type groups.

#### Further division by geomorphological criteria

The statistical analyses of landscape element compositional data provide strong support for a division of the study area into six ‘major landscape types’ by geomorphological criteria: inland hills and mountains; inland valleys; inland fine-sediment plains; other inland plains; coastal plains; and coastal fjords. These major types can be identified either by surface geometry alone or by surface geometry in combination with soil types (inland plains). Back-transformed analysis variable values allow for fine-tuning of criteria for separation between major types based on identified important terrain parameters such as relative relief and proportion of flat terrain. The results of our analyses may thus provide an empirical basis for elaboration of a high-resolution (100×100 m) map of meso-scaled landforms based on consistent criteria.

In a Nordic context, the major types identified by us are recognisable in descriptions (and some coarse-scaled maps) of major geomorphological features that date back to the 19th century (see, e.g. Reusch 1894, Reusch 1905, Rudberg 1960, Gjessing 1978, Klemsdal 1982, Erikstad et al. 2009). Nevertheless, the major types identified in our study differ in several ways from earlier Norwegian geomorphological typologies. Thus Rudberg (1960) identifies 18 ‘morphological type regions’ of which 12 are present in Norway (category numbers according to map in Rudberg [1960]): ‘plains’ (relative height < 20 m); ‘fissure valley landscape’; three classes of ‘undulating hilly land’ with relative height 20–50; 50–100; and > 100 m respectively; ‘monadnock plains’ (i.e. isolated hills or small mountain that rises abruptly from a surrounding plain; present in Finnmark county only); ‘premountain region’; three categories of ‘mountains’ based on relief, including mountain plateaus; ‘strandflat’; and ‘greater faults’. Mattsson et al. (1984) provides a similar but coarser description of 13 (major) landscape types including fjord, strandflat, coastal archipelago, fissure valley landscape, three types of inland plains, and six types of hills and mountains based on relief and location along an elevation gradient. Most, but not all of the types described by Rudberg (1960) and Mattsson et al. (1984), are identified as either major or minor types in our type-system. While our analyses point to geomorphological features as more fundamental for landscape compositional variation than the altitudinal gradient, variation such as between plains along a gradient from the lowland to the mountains in the above-mentioned systems (Norwegian: ‘lavlandssletter’ and ‘fjellvidder’, respectively) are captured by the bio-ecological gradients ‘forest cover’, and hence included in the hierarchy of minor types. Our analyses do not provide support for including ‘fissure valley landscapes’ or ‘coastal archipelago’ (c.f. Rudberg 1960; Mattson et al. 1984) as major types in a consistent system of major landscape types. The fine-grained terrain variation characterising such landscapes will not be captured by morphometric variables derived from a relatively coarse-grained DEM (100×100 m grid cells, derived by interpolation of height data from 20m contour lines). We assume that both fissure valley landscapes and coastal archipelago (i.e. partly submerged fissure valley landscapes), will be recognised by the use of a high-resolution DEM.

Our analyses support the identification of ‘inland valleys’ as a major landscape type.

Inland valleys are widely acknowledged as an important geomorphological feature in geomorphological literature (Gjessing 1978, Sulebak 2007, Erikstad 2009), but not included as a separate type in earlier versions of geomorphological maps covering Norway.

Klemsdal (1982, 2010) identify six primary coastal types in Norway based on morphogenetic criteria (i.e. landform-creating processes): (1) ‘strandflat’; (2) ‘fjord’; (3) ‘fjärd,’ (a Swedish term that refers to submerged valleys that do not have a glacial over-deepened character, often found in Swedish and Southeastern Norwegian coastal landscapes with fissure valleys and ‘skjærgård’); (4) ‘cliff abrasion coast’; (5) ‘flat abrasion coast’; and (6) ‘moraine topography coast’. The major type ‘coastal plains’ identified in our study encompasses the strandflat, the fjärd, the flat abrasion coast and the moraine topography coast in Klemsdal’s typology. Klemsdal’s ‘cliff abrasion coast’ is not identified as a major landscape type on its own in our study. Neither do our analyses reveal distinct patterns of landscape variation that motivate for distinguishing coastal hills and mountains as a separate major type. Contrary to the analyses for inland plains, the ordinations neither reveal patterns of variation along a gradient from coast to inland nor discriminate a coast-near group of landscapes within the major type inland hills and mountains. This result may, however, partly result from the rules for delineation of OUs: ‘coastal hills and mountains’ was represented in the data material with only 3 OUs. In the geomorphological literature, cliff abrasion coast is consistently described as a distinct coastal landform at the scale of our study (Norwegian: ‘klippekyst’; ‘næringer’; see e.g. Gjessing 1977, Klemsdal 1982, Trømborg 2006, Sulebak 2007). We suggest that the distribution of this landform and its co-occurrence with other landforms and landscape elements should be explored further by inclusion of morphometric variables derived from a fine-grained DEM in the analyses. If new analyses still do not provide support for identification of cliff abrasion coast as a separate landscape type, occurrence of this landform can be recognised as a category in the attribute system (see discussion later).

According to EcoSyst principles, a demand on major types is that they differ with respect to the set of complex gradients needed to describe important within-type variation (Halvorsen et al. 2020). This rule motivates for a division of inland plains into two major types: inland fine-sediment plains and other inland plains, based on a clear separation in the ordination diagrams between these two groups with respect to dominant soil type. Nevertheless, since the relatively small number of OUs (118) from inland fine-sediment plains (118 OUs) form a weak basis for identification of major CLGs, we recommend that patterns of landscape-element composition in inland plains are explored further.

The discrepancies between the major types identified by our analyses and earlier Norwegian meso-scale landform typologies may have (at least) three explanations: (1) the major types in our study are identified by similarities in surface geometry (morphometry) without explicitly accounting for the morophogenetic processes responsible for these patterns; (2) finer-scale variation in surface geometry (e.g. gullies, fissures, cliffs, etc.) are not well captured by the relatively coarse-scaled DEM used as the basis for our study; and (3) properties of the data material, such as low representation of OUs from inland plains and coastal hills and mountains or lack of high-quality data for important characteristics such as soil type. On the other hand, the earlier typologies, which express subjective expert opinions, do not provide definitive answers to typification questions. Anyhow, we recommend that the issues raised above are addressed by morphometric studies based on a more detailed DEM, with soil-type data of more consistent quality, and a sufficiently large set of representative observation units.

In a global context, the major landscape types identified in our study largely resemble commonly applied classifications of macro- and meso-scale landforms (e.g. plains, hills and mountains) based on surface geometry, in the tradition of Hammond (1954), Dikau (1989), Brabyn (2005), Karagulle et al. (2017) and Sayre et al. (2019). The major types resemble ‘mesorelief landforms’ in the hierarchical classification of landforms of different sizes by Dikau (1989), exemplified by e.g. valleys and hills with width of the landform units from 10^2^ to 10^4^ m and size of the delineated areas from 10^4^ to 10^8^ m^2^. ‘Mesorelief’ is the level between ‘macrorelief’ (e.g. larger mountain areas such as the Alps or the Rhine Graben) and microrelief (typically identifying landforms such as gullies, dolines, sand dunes and terraces). Sulebak (2007), on the other hand, refer to landforms comparable to the major types in our study as ‘macro-relief’, exemplified by the strandflat, valleys, fjords and large river plains with width of the units from 10 to 10^3^ m and size of the delineated areas from 10^2^ to 10^6^ m², resulting from processes operating over 10^3^–10^6^ years.

### MAIN GRADIENTS OF LANDSCAPE VARIATION

#### Complex gradients of landscape element composition

The analyses show that each major type suggested by the analyses possess a unique set of 2–5 important complex landscape gradients. Furthermore, all major types satisfy the criteria by Halvorsen et al. (2020; Appendix S2.3) that EcoSyst major types have to possess at least one CLG not shared by another major type, and that each major type shall comprise more than one ecodiversity distance unit in landscapes (EDU–L) of variation in landscape-element composition. Although CLGs were identified for each major type independently, many CLGs expressed similar properties across major types. Accordingly, the 24unique CLG variables used to model the proxies representing each major-type specific CLG can be aggregated into 7 ‘universal CLG candidates’ (Table 71). This can be exemplified by the five major-type specific combinations of bio-ecological variables that were used to model the two CLGs ‘open-area cover’ (AP) and ‘forest cover’ (SkP). For all practical purposes, these variable combinations can be interpreted as different expressions of one CLG: ‘vegetation cover’ (the ecological interpretation of this gradient is discussed below). According to the principles of Halvorsen et al. (2016) and Halvorsen et al. (2020), we recommend that ‘universal’ CLGs that can be applied across major types are derived from the major-type specific CLGs whenever possible. The seven ‘universal CLGs’ are discussed separately below.

**Table 71.** The total set of complex landscape gradients (CLGs) identified in this study, including the variables used to model a proxy for each CLG. For each CLG, the following information is given: Universal CLG candidate (I–VII) = group of CLGs expressing similar variation across major landscape types; CLG category (G = geo-ecological; U = bio-ecological; A = land-use related); Code = CLG abbreviation; Major-type(s) [KS = coastal plains; KF = (coastal) fjords, IF = inland fine-sediment plains; IX = other inland plains; ID = (inland) valleys; IA = (inland) hills and mountains]; Analysis variables used to model each CLG (AA = areal coverage of …); Gradient length = gradient length estimate by Method III (se Material and methods section); Segments = number of segments into which the CLG is divided.

| Universal CLG candidate (I–VII) | CLG category | CLG code | CLG | Major type | Analysis variables used to model each CLG | Gradient length | Segments |
| --- | --- | --- | --- | --- | --- | --- | --- |
| I<br>Coast–inland variation | G | IYK | Inner-outer coast | KS | *Number of rivers (R<sub>net_a</sub>) + AA exposed coast (Ekspve_a) + Inverse island size (Oyst_i)* | 0.4026 | 5 |
|  | G | KA | Distance to coast | IX | *Inverse island size (Oyst_i) + distance to coast (Discoast)* | 0.2959 | 2 |
| II<br>Relief | G | RE | Relief | KS | *Mean terrain ruggedness (Rug3_m) + AA convex terrain (Tpi1h_a)* | 0.3003 | 3 |
|  | G | RE | Relief | KF | *Mean coastal ruggedness (coarse scale; Crug9_m) + AA landslide soil (Kskred_a) + Mean terrain ruggedness (Rug3_m) + Relative relief* | 0.3239 | 4 |
|  | G | RE | Relief | IF | *Relative relief (RR1) + AA convex terrain (Tpi1h_a)* | 0.1543 | 1 |
|  | G | RE | Relief | ID | *Altitudinal range (Rr1_m) + Mean terrain ruggedness (Rug3_m) + AA moderate slope (Meant_a)* | 0.5313 | 6 |
|  | G | RE | Relief | IA | *Relative relief (RR1) + Mean terrain ruggedness (Rug3_m) + AA convex terrain (Tpi1h_a)* | 0.555 | 6 |
| III<br>Soil properties | G | JA | Soil type | IF | *AA exposed bedrock (Kbf_a) + AA glaciofluvial deposits (Kelv_a)* | 0.1486 | 1 |
| IV<br>Freshwater lake properties | G | IP | Freshwater lake properties | ID | *Mean distance to lake (Dislake) + Abundance of freshwater lakes(II)* | 0.3784 | 4 |
|  | G | IP | Freshwater lake properties | IX | *Number of lakes (Inns_s)* | 0.3218 | 3 |
| V<br>Abundance of wetlands | G | VP | Abundance of wetlands | KS | *Mean distance to mire (Dismire) + Number of lakes (Inns_s) + Mean distance to lake (Dislake)* | 0.1794 | 2 |
|  | G | VP | Abundance of wetlands | KF | *AA mire (Mire_a) + Number of lakes (Inns_s)* | 0.1860 | 2 |
| VI<br>Vegetation cover | U | SkP | Forest cover | KS | *Areal coverage of deciduous forest (Arlov_a) + AA mixed forest (Arbla_a)* | 0.1726 | 2 |
|  | U | AP | Open-area cover | KF | *AA boreal heaths (Bohei_a) + AA exposed bedrock (Kbf_a) +* | 0.2036 | 2 |

| Universal CLG candidate (I-VII) | CLG category | CLG code | CLG | Major type | Analysis variables used to model each CLG | Gradient length | Segments |
| --- | --- | --- | --- | --- | --- | --- | --- |
|  |  |  |  |  | *AA open areas (Araaf_a ) + AA impediment (Sn_imp)* |  |  |
|  | U | AP | Open-area cover | IF | *AA open areas (Araaf_a ) + AA unregistered heath/open areas (Sn_ureg)* | 0.2533 | 3 |
|  | U | SkP | Forest cover | IX | *AA coniferous forest (Arbar_a ) + AA open areas (Araaf_a ) + AA dry heath/open areas (Sn_torr)* | 0.4619 | 5 |
|  | U | SkP | Forest cover | ID | *AA impediment (Sn_imp ) + AA coniferous forest (Arbar_a)* | 0.3285 | 4 |
|  | U | SkP | Forest cover | IA | *AA impediment (Sn_imp) + AA coniferous forest (Arbar_a)* | 0.2648 | 3 |
| VII<br>Land use | A | OI | Amount of infrastructure | KS | *Amount of infrastructure (IfI) + AA built up areas (Gab_a)* | 0.1344 | – |
|  | A | OI | Amount of infrastructure | KF | *Amount of infrastructure (IfIu ) + AA built-up area (Build_a) + AA built-up(Gab_a)* | 0.2054 | 2 |
|  | A | OI | Amount of infrastructure | IF | *Amount of infrastructure ( IfIu ) + AA built-up (Gab_a)* | 0.1176 | – |
|  | A | OI | Amount of infrastructure | IX | *Amount of infrastructure (IfIu ) + AA built-up area (Build_a)* | 0.3475 | 4 |
|  | A | OI | Amount of infrastructure | ID | *Amount of infrastructure (IfIu ) + AA built-up area (Build_a)* | 0.2626 | 3 |
|  | A | OI | Amount of infrastructure | IA | *Amount of infrastructure (IfIu )* | 0.3052 | 3 |

Furthermore, variation along each universal CLG should subsequently be divided into ‘elementary segments’, i.e. a set of smallest intervals that open for defining major-type adapted segments by terms that can be applied across major types, thus facilitating among major-type comparability.

#### Main geo-ecological landscape gradients in Norway

The geo-ecological CLGs identified in our study express variation in broad structural patterns of abiotic conditions. The analyses reveal that *relief* (RE) is an important geo-ecological CLG in all of the recognised major types except inland plains (which, by definition, has low relative relief). The relief-related CLGs express finer-grained terrain properties than the coarse-grained variation that forms the basis for the division into major landscape types.

Different combinations of variables are used to model the relief gradient in each of the major types. Relief in inland hills and mountains expresses a gradient in steepness and ruggedness from gently undulating to steep and rugged hills and mountains. Relief in valleys and fjords, on the other hand, expresses the depth and width of the valley (or the fjord) relative to its surroundings; from wide, open valleys and fjords to narrow, deep and steep-sided valleys and fjords (see Sulebak 2007). Relief in coastal plains expresses a third, related but slightly different, gradient in topography – ruggedness on a finer scale, i.e. a gradient from flat to rugged coastal plains, the latter often with remnant peaks (in Norwegian: ‘restfjell’).

All of the relief gradients identified in this study are previously recognised and described in the literature (Sulebak 2007). However, to our knowledge they have never been related to other geo- and bio-ecological landscape properties by quantitative analysis at the level and scale of our study. The extensive gradient lengths, the high number of strongly correlated variables and the large fraction of the total variation in landscape properties explained by the relief-related CLGs show that relief controls most of the landscape variation in valleys, fjords and hills and mountains. Our division of the relief gradient in plains, hills and mountains also accords with landform classifications applied at regional, continental or global scale described by, e.g. Hammond (1954), Dikau (1991), Sayre el al. (2014) and Karagulle et al. (2017). All of these typeifications apply subdivisions of major landscape types based on stepwise variation in relief; e.g. plains (from flat to irregular) and hills and mountains (from moderate to very high relief).

Two geo-ecological variables identified in our study express a gradient from coast to inland. The CLG *inner-outer coast* (IYK) is identified as the most important CLG in coastal plains, while the CLG ‘*distance to coast*’ (KA) is identified as the most important gradient in other inland plains. The first CLG explains variation in landscape element composition from the outer coast, directly exposed to the actions of wind, waves and ocean currents towards coastal plains with more typical inland properties. The latter CLG, distance to coast, may be interpreted as a continuation of the former, separating inland plains < 5 km from the coastline from plains further inland. Notably, the analytical variable mean distance to coast (Discoast) was also strongly correlated (τ = –0.45) with the first GNMDS ordination axis in inland fine-sediment plains, although not identified as a separate CLG in this major type. Regional environmental variation in climatic conditions and species distributions from coast to the interior of a landmass is well documented in biogeography worldwide (Lomolino et al. 2017) as well as in Norway (Gjærevoll 1973, Moen 1999, Bakkestuen et al. 2008). Our study shows that this gradient is important for landscape-level variation as well, as expressed by variation in the composition of landscape elements.

All hydrological variables and properties are, as expected, strongly related to landform variation, but the relative importance of various hydrological properties vary among major landscape types. The *abundance of wetlands* (including small lakes and tarns; VP) is identified as an important CLG within landforms with low relief (except inland fine-sediment plains), and within fjords with low relative relief. This CLG identifies areas with high abundance of mires, small lakes and tarns as different from areas with low abundance of these elements. Variation in these landscape elements is a well-known property of boreal and subarctic bioclimatic zones (Keith et al. 2020) and is well documented throughout Norway (Moen 1999, Bryn et al. 2018). The CLG *freshwater lake properties* (IP) in valleys identifies ‘inland fjords’ (Erikstad et al. 2009) and other hydrological features typically occurring in inland valleys, while the CLG freshwater lake properties (IP) identified in other inland plains separates areas with many small lakes from areas with few lakes, thus expressing a different aspect of hydrographic variation.

#### Main bio-ecological landscape gradients in Norway

Our study identifies two bio-ecological CLGs: ‘*open area cover*’ (AP; identified as important in fjords and inland fine sediment plains) and ‘*forest cover*’ (SkP; identified as important in other inland plains, inland valleys and inland hills and mountains). Forest cover is also identified as a CLG-candidate in coastal plains in which, however, the additional variation explained by forest cover after accounting for variation along the gradient ‘inner-outer coast is too small to meet the criterion for inclusion as an important CLG. We interpret ‘open area cover’ and ‘forest cover’ as two different versions of the same complex-gradient, expressing variation in *vegetation cover* from more or less barren mountains, outer skerries etc., without or with sparse vegetation cover, to forested or potentially forested areas. The intermediate segments of the forest-cover gradient which are absent from the open area-cover gradient, comprise open mountain heaths and boreal heath-dominated areas below the climatic forest line, kept open due to historical land-use by logging and grazing (see Bryn et al. 2013). Interestingly, the bio-ecological CLG forest cover (SkP) in other inland plains was characterised by an extensive gradient length and, accordingly, as many as five segments. This is an indication that relief controls and constrains a large amount of the bio-ecological variation in the major types where relief is important, while otherwise this variation has to be attributed directly to a bio-ecological variable which then will comprise more segments.

Landforms such as mountain plateaus and lowland plains are clearly separated by this CLG (e.g. Rudberg 1960, Winsnes et al. 1983; see discussion in the section about major types). We interpret the bio-ecological CLG as a response to abiotic processes that scale up to structural patterns of ‘expressed ecological landscape elements’ at landscape-relevant spatial scales. Examples of such processes are the geomorphological processes responsible for landforms (Sulebak 2007), climate (Bakkestuen et al. 2008), biotic and biogeographic processes including interactions between species (Tilman & Kareiva 1997, Ives & Carpenter 2007), dominance by key species (Paine 1969), dispersal (van Groenendael et al. 2000), landscape-scale consequences of trophic cascades (Ripple et al. 2016) and influence from human land use (Moen 1999). Discussions of the causal order, or relative importance, of factors that determine bio-ecological variation are, however, outside the scope of the current study.

#### The main land-use related landscape gradients in Norway

Variation in the density of buildings, infrastructure and built-up landcover types, here referred to as ‘*amount of infrastructure*’, is identified as an important landscape gradient in all major types. This category comprises gradients in the aggregated outcomes of past and present exploitation of natural resources; typically expressing variation from natural via rural to urban landscapes or from landscapes with extensive or little human land use to landscapes shaped by intensive farming or other harvesting of natural resources. In the initial analyses for identification of CLG candidates, the CLG ‘amount of infrastructure’ is strongly correlated (τ > 0.55) with the first GNMDS axis in all major types. Nevertheless, the analyses for deriving a parsimonious set of CLGs within each major type show that the additional amount of variation explained by amount of infrastructure after the geo-ecological and, eventually, bio-ecological, variation is accounted for, is generally low. These results show that patterns of human land use are strongly determined and controlled by geo-ecological variation. Furthermore, our analyses reveal the broad-scale geo-ecological conditions at which the intensity of human land use is at its highest. Xu et al. (2020) point out that humans, just like other species, have an environmental niche, and demonstrate that humans throughout history have concentrated in a surprisingly narrow subset of Earth’s available climates. Our study indicates potentially interesting relationships between geomorphology and land-use patterns that should be explored further in a paleoecological context.

Agricultural land-use intensity is not identified as an independent CLG in our study, although the major type ‘inland fine-sediment plains’, defined by soil type, to a large extent is characterised by high agricultural land-use intensity. Interestingly, the variation in agricultural land-use intensity consistently follows that of infrastructure. The abundance of agricultural areas increases gradually with the amount of infrastructure to a certain point, before decreasing towards heavily utilised or highly urbanised areas. Like the amount of infrastructure, agricultural land-use intensity is strongly correlated (|τ| > 0.45) with the first GNMDS axis in all major types except inland fine-sediment plains which is characterised by high agricultural land-use intensity throughout.

The fact that human influence is ubiquitous, and affect the composition, structure and function of landscapes throughout the World, is well documented (Vitousek et al. 1997, Ellis et al. 2010). Urban–rural gradients similar to the one identified in our study are previously identified by gradient analysis by e.g. Luck & Wu (2003) and Vizzari et al (2018). Our study also confirms and validates the suggestion by Jakobsson et al. (2020) that the infrastructure index coined by Erikstad et al. (2013) is a good predictor of human influence on the landscape: this is the single variable that best explains total variation in human traces in the landscape. The strong co-variation with agriculture and the weak basis for defining agriculture as an independent CLG, indicate that agricultural land use intensity might be incorporated in the infrastructure index, as exemplified by another recent example of continuous mapping of the rural–urban gradient: the ‘Human footprint index’ (Sanderson et al. 2002).

#### How universal are the complex landscape gradients?

The CLGs identified in this study express variation in the distribution and abundance of observable landscape elements and properties, and do not explicitly address the processes behind these patterns. In this sense, they are parallel to the ‘coenoclines’ of Whittaker (1960), the species compositional gradients, which do not explicitly that the conditioning environmental complex-into account and therefore do not allow direct mechanistic modelling of the processes which give rise to the observed patterns. Nevertheless, the order of historical events and processes that have brought about landscape variation is well established by geological, paleoecological and historical evidence (Delcourt et al. 1982, Birks 1993). The identification of CLGs opens for further studies of the relative magnitude of the drivers behind landscape variation, and the response to these drivers as expressed in landscape element composition. Such knowledge may have potential importance also for future predictions of landscape change.

The total set of CLGs identified as an important component of a parsimonious characterisation of landscape variation, bears resemblance to ‘factors’, ‘components’, and categories commonly applied in biophysical landscape characterisation and mapping. In a review of 54 methods, Simensen et al. (2018) find that the factors most often used for landscape characterisation and mapping are landform (included in 95% of the analysis methods), general land cover (83%), vegetation cover (81%), geology (67%), soil (63%), hydrography (57%), agriculture (54%) and buildings and infrastructure (52%), all of which are included in the set of CLGs identified in our study. Since our study explicitly addresses the physical outcome of the processes that constitute landscape variation (i.e. observable landscape elements), drivers such as climate (included in 44% of the studies in the review) are not among the analysis variables included in our study.

The ‘key factors’ applied by Puschmann (2005) in a structured qualitative description of 45 coarse-scale ‘landscape regions’ and 444 ‘sub-regions’ in Norway are: (1) major landform; (2) small-scale landforms; (3) water and watercourses; (4) vegetation; (5) agriculture; (6) buildings and infrastructure. Together, Puschmann (2005) use these components to qualitatively derive and describe the ‘landscape character’ of each region, sub-region and/or ‘landscape area’, subjectively assigning to each component in an area or a region a value from one to three to indicate the component’s relative importance for the total landscape character. Although capturing much of the same variation as our study does, the scale and subjective procedure makes it difficult to develop this approach further in a more quantitative setting. Our quantitative approach successfully identifies the properties that are commonly regarded important in landscape characterisation and mapping, and, more importantly, provide strong empirical evidence for the relationship between these variables, i.e. the relative importance of each CLG. Thus, the outcome of our study lends strong support to Strand (2011), who argue that quantification of landscapes by use of variables and proxies for subsequent application in GIS-based mapping, opens for consistent landscape-type mapping with a level of detail and at spatial extents unachievable by field-based landscape type mapping.

Gradient analysis at the landscape level is, to a large extent, still unexplored [but see Luck & Wu (2002) and McGarigal & Cushman (2005)]. The ‘universality’ of the CLGs and their potential for model extrapolation in time and space (e.g. to Scandinavia and other boreal regions) is therefore largely unknown. The composite and ‘data-driven’ nature of the CLGs may theoretically limit their potential use for model projections and extrapolations in time and space. However, many of the landscape gradients recognised in our study are probably relevant over vast areas and likely to persist for long times, even in a changing climate (e.g. terrain gradients, coast-inland gradients, gradients in human land-use intensity, gradual changes in vegetation cover, the abundance of lakes, etc.). We suggest that the distribution of ecosystems along complex-gradients in the landscape should be explored across larger areas, as such studies may yield new insights of both theoretical and practical importance.

#### Explained and residual landscape variation – the need for a descriptive attribute system

Like ecodiversity models in general, the landscape space model partitions variation into ‘explained’ variation and residual, ‘unexplained’ or ‘apparently random’ (Økland 1990), variation. A landscape-type system built on CLGs will disregard all variation not explained by the important CLGs. Unless appropriately handled, this will have the unfortunate consequence that the ‘uniqueness’ or idiosyncracies, of landscapes (Phillips 2007) which is expressed in unexplained variation, will be lost. This motivates for a standardised attribute system by which the residual variation can be described in a standardised way (see Halvorsen et al. 2020). No such system has, however, yet been developed for description of landscapes in Norway. We recommend development of an attribute system according to EcoSyst principles, structurally similar to that for ecosystems (see Halvorsen et al. 2020). Based on our study, good candidate variables for inclusion in such an attribute system, are CLG-candidates that ‘did not make it’ to the final set of important CLGs within each major type. An example of such a gradient is ‘agricultural land use-intensity’, which failed to explain enough variation to be identified as a separate, important, CLG, but which is obviously useful for a more fine-grained and precise description of the various aspects of human legacies in the landscape. Furthermore, CLGs identified as important in other major types, can be used for a more detailed assessment, such as vegetation cover in coastal plains. This CLG was strongly correlated with geo-ecological variables, not explaining enough addtional variation to be included as a CLG for coastal landscapes. Moreover, all the 85 variables used for the analyses can in principle be useful as variables in an attribute system, for two at least reasons: (1) because they are specifically selected to describe aspects of the landscape that are considered relevant at the scale and level of our study; and (2) because they result from a pruning process by which strongly correlated variables were omitted. All the 85 analysis variables therefore account for some of the variation in landscape properties not also accounted for by other variables.

Data about the presence, absence and relative abundance of major ecosystem types (e.g. exposed ridge, open fen, mire and swamp forest) and microrelief landforms (e.g. gullies, dolines, sand dunes and terraces) will have to be important components of an attribute system for the landscape level. Although field-survey based mapping of such landscape elements is time-consuming and expensive, alternative, efficient pathways to acquiring information about their spatial distribution may be used for this purpose, such as remote sensing (Pereira et al. 2013, Zarnetske et al. 2019), distribution modelling (Simensen et al. 2020) and morphometric analysis (Jasiewicz & Stepinski 2013).

The tentative landscape-type system built on the results of our study highlights landscape-element composition (the relative performance of different observable objects within each spatial unit), with less emphasis of structure (the patterns of distribution of observable objects within these spatial units) or function (the processes that give rise to the variation). We therefore recommend that landscape structure and function are incorporated as central components in the attribute system. This can be accomplished in many different ways due to the rapid proliferation of available ‘landscape metrics’ for quantification of the spatial configuration of categorical landscape patterns over recent decades (Turner & Gardner 2015, Hesselbarth et al. 2019). Similarly, spatial statistics, including geostatistics, are widely applied in landscape ecology to quantify the spatial structure of continuous data (Dale & Fortin 2014). Many of these metrics are good candidates for inclusion in an attribute systems for description of landscape variation. Examples of structural properties not captured by our analysis are the spatial configuration of discrete land-cover types as expressed by edge length, density, contagion and perimeter-to-area ratios. Relevant metrics for patches of single land-cover types can be exemplified by largest patch index, patch dispersion, patch isolation, connectivity and fragmentation (see Turner & Gardner 2015). Variables that describe disturbances and abrupt changes with landscape-scale effects such as flooding regime, wildfires, logging, insect outbreaks, etc. should also be considered for inclusion in an attribute system at the landscape level (Turner 2010, Ratajczak et al. 2018). The same applies to all variables, properties and categories already include in the EcoSyst attribute system, provided that area-covering data exist and that the variables are relevant at the landscape level of analysis.

#### How fine-grained is an ideal landscape-type system?

We demonstrate that the procedure originally developed by Halvorsen et al. (2016) for dividing major ecosystem types into minor types based on differences in species composition – ‘ecodiversity distance units in ecosystems’ – is applicable also to the landscape level of ecological diversity, with necessary adaptions described in the methods chapter. The definition of the EDU–L (ecodiversity distance units in landscapes; see Halvorsen et al. 2020) determines the level of detail captured at the lowermost hierarchical level in the landscape-type hierarchy. When compositional variation along most CLGs is continuous, no a priori reason exist for selecting a specific EDU–L value to define the edge of an ideal basic-type hypercube in landscape space with important CLGs as axes. Standardised stepwise division of landscape gradients can then, as for LCEs on the ecosystem level, be done by defining one EDU–L unit as a specific threshold value on the PD dissimilarity scale. No other guidelines for how much variation one EDU–L should account for, exist other than practical considerations; the choice of the threshold value for landscape distance will determine the level of detail of the type-system. A high threshold value will result in a coarse-grained type-system with few categories while a low threshold value will provide a fine-grained type-system with many categories. The choice between the two approaches is largely a matter of preference; i.e. whether you are a ‘lumper’ or a ‘splitter’, in the words of George G. Simpson (1945).

With the ambition to balance different considerations, we defined one ecodiversity distance unit in landscapes (EDU–L), i.e. one landscape distance unit, to 0.08 PD units. The number of major type-adjusted segments by definition equals the number of EDU-L between CLG endpoints, rounded down to the nearest whole number (Halvorsen et al. 2020). This means that the gradient length must be at least 2 EDU–L units (> 0.16 PD units) in order for a CLG to be used to divide a major type into basic types, at least 0.24 PD units to provide three segments and a series of three basic types, etc. Experience from the analysis of the inland data sets (especially the inland vallet data set ID1012), which resulted in gradient lengths up to 0.5313 for CLG 1 (Relief), corresponding to > 6 EDU–L, may indicate that the definition ‘1 EDU–L = 0.08 PD units’ results in a type-system that is too fine-grained, i.e. with too many types at the lowest level. An alternative view, supported by the fact that the swarm of OUs becomes ‘thinner’ (shows reduced point density) from mid-points towards endpoints along almost all ordination axes, is that the landscape gradient length estimates are disproportionately heavily influenced by outlier OU. Outliers are OUs that occupy isolated positions in an otrdination diagram (Økland 1990). Outlying OUs may represent: (1) individual OUs with atypical properties for the major type in question; (2) OUs in the transition zone between two adjacent major types; or (3) misclassified OUs, which should actually have been affiliated with other major types. Presence of outlier OUs amplify the effect of lack of generalisation in the extreme segments along the rONVs. The interpretation of our results provides several examples of CLGs for which gradient lengths are likely to have been extended beyond reasonable limits for these two reasons, also showing that this effect is independent of the method used for gradient length estimation. This effect calls for careful manual correction of estimated gradient lengths to a ‘gradient length corrected for atypical OUs, transitions to adjacent major types and lack of generalisation of the analysed data’. Such a correction cannot be automated but will have to be done after careful assessment of:

1. Outliers; ‘extreme’ OUs along the sONVs that should be carefully reviewed with respect to their representativity for normal variation along the current CLG within the major type. In particular, misclassified OUs and OUs that represent atypical combinations of landscape properties should be identified and gradient length estimates (and CLG segment numbers) reduced accordingly.
2. The distribution of OU points along ordination axes; areas along CLGs that are very sparsely populated with points should be downweighed in calculations of ‘corrected gradient length’. This is parallel to the criterion used to avoid inappropriate fragmentation of types at the ecosystem level (Halvorsen et al. 2016: chapter B4c).
3. Disjunctions; large ‘gaps’ in the point distribution should be assessed in order to determine whether the CLG should be recognised as naturally divided in classes (factor variables).

Some outliers in the analyses are represented either by few OUs (e.g. highly urbanised landscapes) or by relatively few variables (e.g. landscapes dominated by glaciers and landscapes totally dominated by agricultural land-use). A truly evidence-based type system requires further analyses of such landscapes.

### POTENTIAL SOURCES OF ERRORS AND UNCERTAINTIES

Any statistical analysis is a product of the available data and the analytic methods used. This study is no exception. Despite strong efforts to make the approach to landscape analysis presented here observer-independent, the results of the analyses depend on the selection of data sources and other subjective choices made during the analytic procedure (Yang 2020). The 85 variables used in the multivariate analyses are derived from different sources that are mapped, collected or modelled at different time-points and by use of different methods, inevitably introducing variation in the quality of the input data. Even though we used variables derived from standardised national mapping programs (e.g. AR 5 land-cover maps, N50 topographical land-cover data, etc.), also data from these basic data sources are of varying quality. This is exemplified by Bryn et al. (2018), who show that the cover of wetlands in the basic Norwegian land-cover map N50 (data also used in our study) is strongly underestimated while the cover of, e.g. forest, has been correct. Furthermore, the lack of area-covering and/or reliable data for potentially important landscape elements and properties is an important source of uncertainty in the analysis.

Although we analysed a broad set of 85 variables, relevant data for several potentially relevant variables were not available at the time of the study. Variables that would have been included in the study if area-covering data were available, can be exemplified by the presence and/or fractional area occupied by major ecosystem types (cf. Bryn et al. 2018) and landscape elements and properties related to historical land use (Fairclough & Herring 2016), former extensive land use in particular (e.g. utilisation of outfield resources, grazing, forestry, etc.). Hence, our study must be considered as a systematisation of existing available knowledge that can be tested and improved in an inductive manner with new and better data available for analyses, in accordance wth the basic philosophy of EcoSyst (Halvorsen et al. 2020). We are confident that future landscape analyses will benefit from rapid development of relevant remote sensing technology, as exemplified by the recently published global maps of physiographic features with fine-grained resolution developed by a joint venture among scientists (e.g. Sayre et al. 2014) and rapid development of new elevation data based on LIDAR technology.

Notably, our analysis is confined to a particular scale. Because most land-surface properties are recognisable at specific spatial scale domains, and/or can be represented by different algorithms and sampling grids, no map computed from a DEM is definitive (Pike et al. 2009). Differences in the delineation of major landforms by different map sources may also result from different levels of generalisation (i.e. size of the circle/moving window used for focal calculations). We assume that analyses based on a high-resolution grid (i.e. pixels < 100×100 m) will improve the overall sensitivity of the terrain analyses.

A potential source of uncertainty related to our statistical analyses is the number of correlated variables representing different, but thematically and/or causally related properties. Although strongly correlated variables (τ > 0.7) were removed prior to the analyses, there is still a risk that related landscape properties represented by many correlated variables (e.g. 18 analytical variables related to buildings and infrastructure and 15 variables related to topography) are disproportionately strongly emphasised in the analyses.

Another possible source of errors and uncertainty in our results is the sensitivity of the analytical results to the definition and spatial delineation of data collection units. The ‘zoning problem’ is an aspect of a well-known problem in the statistical analysis of spatial data known as the ‘modifiable areal unit problem’ (MAUP; Gehlke & Biehl 1934, Openshaw 1984). Jelinski & Wu (1996) demonstrate that, when a given set of areal units is recombined into zones that are of the same size but located differently, variation in data values and, consequently, different conclusions may result. According to Fotheringham & Wong (1991), the effects of MAUP is a bigger problem in multivariate statistical analysis than in univariate or bivariate analysis. We addressed the MAUP problem according to the recommendations of Jelinski & Wu (1996) by applying an ‘optimal’ zoning system, i.e., a system that maximises interzonal variation and minimises intrazonal variation. This was done by delineating observation units by a set of rule-based criteria derived from the ‘pilot Nordland study’. This approach may reduce variation in results of an analysis caused by the MAUP, but has the disadvantage that ‘optimality’ is a subjective criterion, both with respect to definition and to operation. Furthermore, delineation errors may be introduced during the process by which ‘optimal spatial units’ are constructed. This is exemplified by inconsistencies or uncertainties in the OU delineation process, which is mentioned as a possible cause of the ‘thinning’ of OUs towards CLGs ends observed in this study, and by OUs transitional between inland plains and inland mountains. The criterion-based delineation of OUs may also introduce an element of circularity in the analysis. By defining OUs by spatial delineation based on a set of pre-defined criteria, there is a risk that the analyses will identify the same OUs and confirm the criteria used to delineate them. Use of standard sample units of a constant size (e.g. 1×1 km cells) removes the subjective judgement involved in the delineation of irregular sample units, but does not solve the zoning problem (i.e. the infinite number of possible locations of these zones). Furthermore, standardised sample units have the drawback that fundamentally different landforms and/or ecosystems such as mountains and ocean systems will inevitably be included in the same unit (Bunce et al. 1996). A lesson learnt from the pilot study in Nordland (Erikstad et al. 2015, Halvorsen et al. 2016) is that use of standardised sample units of 5×5 km is inappropriate for studying patterns in landscape element composition in a region characterised by high landform diversity and ‘steep’ gradients. We instead suggest that the impacts of the choices of shape and size of sampling units are further explored by simulations and experiments.

## CONCLUSION

Any landscape comprises a unique combination of landscape elements and properties – unlikely to be duplicated at any other place or time. Nevertheless, this study shows that landscape variation is, to a large extent, non-random and highly predictable. Our study demonstrates how geomorphology, ecology and human land use are tightly intertwined and show that predominantly abiotic processes, to a large extent, control and/or constrain biotic processes and variation in human land use. Predictable patterns of co-occurrence exist between landscape elements and other properties of landscapes, and most landscape elements tend to have distinct performance (probability of presence and/or abundance) optima along broad-scale compositional landscape gradients. Based on the results presented here, we suggest that the gradient perspective, so often applied in studies at the ecosystem level, is a conceptual framework appropriate for studies of landscape-level variation. Such studies may contribute to a more coherent understanding of the relationship between connections between non-living and living nature.

Although classification must be considered a tool, not a goal in itself (Økland & Bendiksen 1985), any type-system developed with the aim of representing real systems or processes should be based upon a consistent theoretical framework and the best available empirical evidence. Our study provides, to our knowledge, the most comprehensive quantitative assessment that simultaneously addresses biotic and abiotic variation at the landscape level of ecological diversity in Norway. Although data for some of the major types are sparse and should be explored further (especially inland plains and coastal hills and mountains), we regard our results as sufficiently clear to provide the basis for the first version of an empirically based landscape type map of Norway according to EcoSyst principles (Halvorsen et al. 2020).

## ACKNOWLEDGEMENTS

We thank Vegar Bakkestuen (Norwegian Institute for Nature Research) for assistance with data collection, data processing and initial analyses. We are also grateful to Geir Davidsen (Nordland County Council), Lars Andre Uttakleiv (Aurland Naturverkstad), Inge Lindblom (The Norwegian Institute for Cultural Heritage Research; NIKU), Eiliv Larsen (Geological Survey of Norway; NGU), Wendy Fjellstad (Norwegian Institute of Bioeconomy Research; NIBIO), Håvard Hjermstad-Sollerud and Ola-Mattis Drageset (Directorate of Public Roads), Øyvind Bonesrønning and Arild Lindgaard (Norwegian Biodiversity Information Centre) for inputs, feedback and support during the development of the project.

We also thank Dag Bastholm (Nordland County Council), Ingunn Limstrand and Bjørn Bjørnstad (Norwegian Environment Agency), Liv Kirstine Just-Mortensen, Bjørn Casper Horgen and Kristin Nordli (Ministry of Local Government and Modernisation), Kristin Thorsrud Teien (Ministry of Climate and Environment), Ivar Myklebust and Einar M. Hjorthol (Norwegian Biodiversity Information Centre) for continuous backing and administrative support during the development of the project.

This work was economically supported by the Research Council of Norway (Public Sector PhD scheme), the Norwegian Environment Agency and the Norwegian Biodiversity Information Centre.

## SUPPLEMENTARY MATERIAL

### RULES FOR DELINEATION OF SPATIAL UNITS APPLIED IN THE PILOT STUDY IN NORDLAND

Prior to the study, a pilot study in Nordland county were conducted. Basic methodological characteristics of the pilot study in Nordland landscape analysis was: 1) a rule-based division of the landscape into 258 discrete spatial units; 2) recording of a broadest possible selection of physical landscape attributes within each spatial unit (i.e. 173 landscape variables); and 3) multivariate statistical analyses of the data set in order to identify ‘landscape gradients’, i.e., ‘gradual variation in the presence and/or abundance of landscape elements’. The resulting typology was obtained by dividing the identified, major complex landscape gradients that explained most variation in the composition of landscape elements into segments, and then defining landscape types by combining intervals along several gradients.

An overview of the method is presented in presented in Erikstad et al. (2015), while further details are provided in Norwegian language in Halvorsen et al. (2016). Here, we provide a translation of the criteria used for delineation of spatial landscape units for major types (Appendix 8a; Halvorsen et al. 2016) since they are of particular importance for the reproducibility of this study.

**Delineation of spatial units representing major landscape types**

**Minimum mapping units**

The spatial units (polygons) representing major landscape types have to satisfy the following main criteria:

1. Area > 4 km²
2. Minimum width > 2.5 km (geodesic distance between the outer boundaries of a spatial unit)

**Delineation**

1. Spatial units representing a major-type candidate that satisfy the geomorphological definition of a major landscape type but do not meet the minimum size criteria shall, unless otherwise indicated, be assigned to adjacent spatial units at the best discretion.
2. Exceptions are described in the procedure in Table S1.

**Delineation order:** steps A1 – A7 below (applied strictly, no exceptions)

---

**Table S1.** Delineation rules, major landscape types. Abbreviations: RR1 = relative relief within a 1 km neighbourhood; TPI = terrain position index; TPI(mt) = terrain position index based on marine and terrestrial areas (the complete DEM); TPI(t) = TPI based on terrestrial areas only (marine areas set to meters above sea level = 0). The number after ‘TPI’ refer to the size of the neighbourhood circle for TPI-calculations.

| Step | Task | Main criterion | Additional criteria, special cases, comments |
| --- | --- | --- | --- |
| A1 | Delineating spatial units for valleys and fjords | **Definition and starting point for delineation:**<br>Valley and fjord delineation based on terrain position index (TPI) = 0<br>**Manual correction of upper border** (between fjords/valleys and surrounding hills and mountains):<br>TPI = 0 for (neighbourhood = 6 km; the inflection point between concave fjords and/or valleys and convex hills and/or mountains).<br>In cases where the fjord/valley profile lacks inflection points, the following criteria are applied (in ranked order):<br>1. If the valley/fjord side has an even slope (which in most cases is synonymous with gentle), the borderline is drawn in the middle of the slope, i.e., in the middle of the valley bottom and top of the adjacent hill/mountain.<br>2. In cases where the valley/fjord side is not even, the border is drawn through cliffs, steep sections, etc. as close to the middle of the fjord/valley side as possible. Digital topographic maps are useful for this purpose. | **Comments:**<br>1. **Upper border of spatial units for fjords** shall be corrected by TPI(t) = 0, i.e., TPI in a calculation where all marine areas are set to 0 meters above sea level, instead of TPI(mt) = 0, when the line for TPI(mt) = 0<br>a. crosses the coastline<br>b. is located closer to land than 250 m over a distance of 1 km, and/or<br>c. is located in a continuous spatial unit with relative relief (RR1)>200 m (steep terrain) that crosses the coastline in a length of at least 1 km<br>2. For valleys TPI6(mt) = TPI6(t) |
| A2 | Delineating spatial units for coastal plains | **Definition:**<br>1. Coastal plains contain the ‘Strand flat’ along the coast in the classical (geomorphological meaning of the term, from Rogaland and northward, and<br>2. Other coastal landscapes with similar relief located other places along the coast | **Comments:**<br>1. **Upper border towards hills and mountains above the coastal plains** are set to TPI(mt) = 0, except cases where the line for TPI(mt) = 0<br>a. crosses the coastline<br>b. is located closer to land than 250 m over a distance of 1 km, and/or<br>c. is located in a continuous spatial unit with relative relief (RR1)>200 m (steep terrain) that crosses the coastline in a length of at least 1 km |
|  |  |  | When the criterion in A, B or C is met, the line shall be corrected by use of $TPI(t) = 0$.<br>2. If further corrections are needed (without easily identified inflection points in slopes) the criteria A and B in the procedure for delineation fjords are applied. |
| A3 | Delineating spatial units for residual areas with coastline | Definition:<br>‘Residual areas with coastline’ contain areas with coastline, not assigned to spatial units of the major landscape types fjords/valleys or coastal plains in step A1 and A2.<br>Delineation of spatial units for candidate major types include manual assessment of the minimum mapping unit criterion (2), interpreted as generalised coastline length $> 2.5$ km.<br>1. If the criterion for the minimum mapping unit is met, the spatial unit is assigned to the major type ‘coastal hills and mountains’.<br>2. If the criterion for the minimum mapping unit is not met, and the affiliation to major type is equal on both sides of the spatial unit, the adjacent units are merged together.<br>3. If the criterion for the minimum mapping unit is not met, and the affiliation to major type is different on both sides of the spatial unit, the coast-line segment shall be split and merged by the two adjacent spatial units. The borderline between them shall follow a ridge or another natural terrain divide (i.e. ridge, depression, etc.). Note that the criterion 1 for size of minimum mapping unit is not applied in this step. |  |
| A4 | Separating fjords and valleys | **Main criterion:**<br>The borderline shall follow a ridge or another natural geomorphological structure in the terrain across (perpendicular to) the | **Exceptions:**<br>The main criterion is overruled by continuous spatial units with amount of infrastructure ($OI \geq$ segment 5, i.e. village or city). |
|  |  | valley, maximum 2 km from the end of the fjord ('fjordbotn').<br>**Indicative tool:** map of maximum TPI1. | These spatial units shall not be split between fjords and valleys but shall be assigned as fjords.<br>**Comment:**<br>1. Some places, short valley segments will be generated between the fjord and the surrounding hills and mountains. These segments shall be delineated as valleys when the segment is longer than 2 km. When this criterion is not met, the valley segment shall be joined with the fjord unit. |
| A5 | Separating coastal hills and mountains from inland hills and mountains and inland plains | **Main criterion:**<br>TPI6(mt) = 0 (similar to the criterion for separation between the major types 'fjords' and 'coastal plains'), secondarily TPI(t) = 0.<br>**Manual delineation** is applied in gentle or rugged terrain where TPI = 0 is not applicable. The borderline between coastal hills and mountains and inland hills and mountains shall follow the first natural geomorphological feature (normally a ridge), maximum 2 km from the coast-line.<br>**Indicative tool:** map of maximum TPI1.<br>**Manual correction of lower borderline** (to marine landscapes) is applied with similar criteria. | **Exceptions:**<br>1. Continuous coastal spatial units with amount of infrastructure ($OI \geq 5$, i.e. village or city). These spatial units shall not be split, but be assigned as one spatial unit with affiliation to coastal plains.<br>2. If it is not possible to delineate inner spatial units on islands in line with the general size criteria for hills, mountains and inland, the island shall not be split between inland and coastal landscapes<br>3. If it is not possible to delineate spatial units in line with the general size criteria for hills, mountains and inland plains between a spatial unit for coastal hills and mountains and a spatial unit for inland valleys (i.e., a ridge/hilltop between the valley [perpendicular to the valley direction] and the coastal areas outside), the area in between shall be split between the two spatial units. |
| A6 | Delineating spatial units for steep and rugged hills and mountains within coastal plains | Spatial units for hills, mountains and plains shall be delineated within a spatial units in coastal plains, regardless of the size criterion for minimum mapping units when the criterion for 'steep and rugged hills and mountains' (TP = 2, based on TPI1) is met. | **Comment:**<br>1. Segment A7 represent an exception from the minimum size criterion. |
| A7 | Delineating spatial units for inland plains | **Definition**<br>Inland plains are defined by a combination of properties: | **Comment**<br>**1. Borderline towards higher hills and mountains** is drawn with use |
| | | 1. Relative relief (RR) in 1 km neighbourhoods; $RR1 < 50$ m; | of $TPI(t) = 0$. If further corrections are needed (i.e. if adjacent hillside lack inflection points), the criteria |
|  |  | 2. The plain meets the geomorphological definition of a plain formed in deposits and is not a part of a spatial unit assigned as a valley. | A and B in manual adjustments for upper borderline for fjords and surrounding hills and mountains are applied (A1): |

